# System-wide mapping of neuropeptide-GPCR interactions in *C. elegans*

**DOI:** 10.1101/2022.10.30.514428

**Authors:** Isabel Beets, Sven Zels, Elke Vandewyer, Jonas Demeulemeester, Jelle Caers, Esra Baytemur, William R. Schafer, Petra E. Vértes, Olivier Mirabeau, Liliane Schoofs

## Abstract

Neuropeptides are ancient, widespread signaling molecules that underpin almost all brain functions. They constitute a broad ligand-receptor network, mainly by binding to G protein-coupled receptors (GPCRs). However, the organization of the peptidergic network and roles of many neuropeptides remain elusive, as our insight into neuropeptide-receptor interactions is limited and many peptide GPCRs in animal models and humans are still orphan receptors. Here we report a genome-wide neuropeptide-GPCR interaction map in *C. elegans*. By reverse pharmacology screening of over 55,384 possible interactions, we identify 461 cognate peptide-GPCR couples that uncover a broad signaling network with specific and complex combinatorial interactions encoded across and within single peptidergic genes. These interactions provide insights into neuropeptide functions and evolution. Combining our dataset with phylogenetic analysis supports peptide-receptor co-evolution and conservation of at least 14 bilaterian peptidergic systems in *C. elegans*. This resource lays a foundation for system-wide analysis of the peptidergic network.

**Highlights:**

- System-wide reverse pharmacology deorphanizes 68 *C. elegans* peptide GPCRs
- Discovery of 461 peptide-GPCR pairs and additional ligands for characterized GPCRs
- Neuropeptides show specific and complex combinatorial receptor interactions
- Peptide-GPCR pairs support long-range conservation and expansion of peptide systems

## Introduction

Neuropeptides are one of the largest and most widespread classes of signaling molecules that regulate physiology and behavior in all animals. Although bioactive peptides are also secreted from non-neuronal tissues, most are released in the nervous system and signal via G protein-coupled receptors (GPCRs). Neuropeptides are among the most ancient neuronal messengers, likely predating the evolution of neurons^1, 2^, and many peptidergic systems have broadly-conserved functions across bilaterian animals^3–7^. Because of their crucial roles, neuropeptide-receptor systems are increasingly gaining traction as potential therapeutic targets for human diseases^8–11^. GPCRs represent over 30% of the targets for prescribed drugs, and abnormalities in neuropeptide transmission have been implicated in multiple diseases^8, 10, 12^. Indeed, these signaling systems are involved in many, if not all, biological processes, ranging from growth to feeding, reproductive behavior, nociception, learning and memory, as well as the regulation of behavioral states, such as sleep and arousal^2, 3, 7, 13–19^.

While several efforts have been made to comprehensively map synaptic connectivity in animal brains^20–23^, our insight into the system-wide organisation of the peptidergic network is limited, as huge gaps remain in our understanding of neuropeptide-receptor interactions. Single-cell transcriptomic studies have revealed broad expression of neuropeptides and receptors in animal brains^24–27^. However, understanding the actions of neuropeptides also requires a detailed knowledge of their biochemical receptor interactions. Neuropeptides mainly bind to GPCRs that activate secondary messenger pathways and, because the number of second messenger responses is limited, the diversity of neuropeptide-GPCR couples is thought to provide the complexity required for brain function^28^. Indeed, most animal genomes harbour around 100 to 150 peptide GPCRs and a similar number of neuropeptide genes^29–33^. These encode hundreds of bioactive peptides that are cleaved from prepropeptides containing one or multiple peptide sequences^34, 35^. Some receptor genes also code for multiple GPCR isoforms, which may have different ligand interactions^36, 37^. In addition, neuropeptides can interact with multiple receptors and *vice versa*^38–40^. These complex interactions further diversify the number of peptidergic pathways and complicate the characterization of the neuropeptide-receptor network.

Neuropeptide-GPCR couples are challenging to predict from prior knowledge, such as neuronal connectivity, expression or sequence information, as neuropeptides have only short sequences and activate GPCRs that are often distantly located from their release site^41–43^. In the last decades, reverse pharmacology has proven to be a successful approach for the deorphanization of GPCRs, by expressing receptors in heterologous cells and identifying their ligand(s) in a compound library^44^. This way, a broad range of peptide GPCRs have been deorphanized, including at least 138 receptors in humans, 36 in *Drosophila melanogaster*, and 29 in *Caenorhabditis elegans*^40, 45–61^. Nevertheless, many peptide GPCRs remain orphan receptors, hampering investigations into their functions^62^. Although some large-scale efforts have been undertaken, *e.g.* for *Platynereis* receptors and human peptide GPCRs ^40, 63, 64^, most deorphanization studies only focussed on single receptors or a subset of GPCR candidates, which may also overlook complex interactions of peptides with multiple receptors^40, 65^. To gain a deeper understanding of the peptidergic signaling network, system-wide deorphanization approaches are required that examine as many interactions as possible between neuropeptides and peptide GPCR variants encoded in animal genomes.

Due to its compact and well-defined nervous system, the nematode *C. elegans* provides an attractive model organism to map and functionally characterize the neuropeptide signaling network at an organismal level. The complete anatomy and wiring diagram of the *C. elegans* nervous system were defined by serial section electron microscopy, facilitating studies of peptidergic interactions with the wired circuitry^20, 66–68^. The *C. elegans* genome encodes over 300 bioactive peptides that are classified as FMRFamide-like (FLP), insulin-like (INS) and other neuropeptide-like (NLP) peptides^69, 70^. In addition, its genome encodes around 150 predicted peptide GPCRs^47, 71^. Many of these neuropeptide systems are shared between nematodes and other animals, including humans, and have conserved functions^6, 42, 72, 73^. Transcriptional profiles of all neuropeptide and peptide GPCR genes are also available for individual neuron classes among the 302 neurons of the *C. elegans* hermaphrodite nervous system^25, 74^. Like in mammals and other animals, distinct neuron types in *C. elegans* were found to express unique codes of neuropeptide genes and receptors, suggesting different roles for neuron classes in transmitting neuropeptide signals^24, 25, 27^. However, the relationship between peptidergic sender and receiver cells largely remains elusive, as over 80% of the predicted peptide GPCRs in *C. elegans* are orphan receptors^47, 71^. Similarly, target receptors remain unknown for the majority of *C. elegans* neuropeptides, hindering further analysis of the structure and functional organization of the peptidergic network.

Here, we set up a reverse pharmacology pipeline for the large-scale deorphanization of *C. elegans* peptide GPCRs and generated a comprehensive resource of 461 cognate peptide-GPCR couples. System-wide screening of individual neuropeptides and GPCR isoforms revealed novel ligands for known and orphan peptide receptors, complex signaling interactions, and distinct peptide-receptor pairs encoded within single genes. We combine our dataset with phylogenetic analysis to reconstruct the evolutionary history of *C. elegans* peptidergic systems and suggest an evolution-based nomenclature for peptide and receptor genes. This resource also provides a basis for constructing the neuropeptidergic connectome in *C. elegans*^75^.

## Results

### Large-scale deorphanization of *C. elegans* neuropeptide GPCRs

The *C. elegans* genome has at least 149 genes for putative neuropeptide GPCRs (Data S1), most of which encode orphan receptors^47, 71, 76, 77^. To map the *C. elegans* neuropeptide-GPCR signaling network, we set up a reverse pharmacology platform in which we systematically screened for neuropeptide ligands of these receptors in cultured cells (Figure 1A). We first cloned the cDNA sequences of peptide GPCR candidates in a pcDNA3.1 vector for expression in heterologous cells. Multiple receptor isoforms were included in the GPCR library, as receptor variants may be differentially activated or inhibited by neuropeptide ligands^36, 78^. In total, we cloned 161 receptor cDNAs, covering 87% of the neuropeptide GPCR genes that are known and predicted in *C. elegans* (Data S1 and S2).

**Figure 1.**
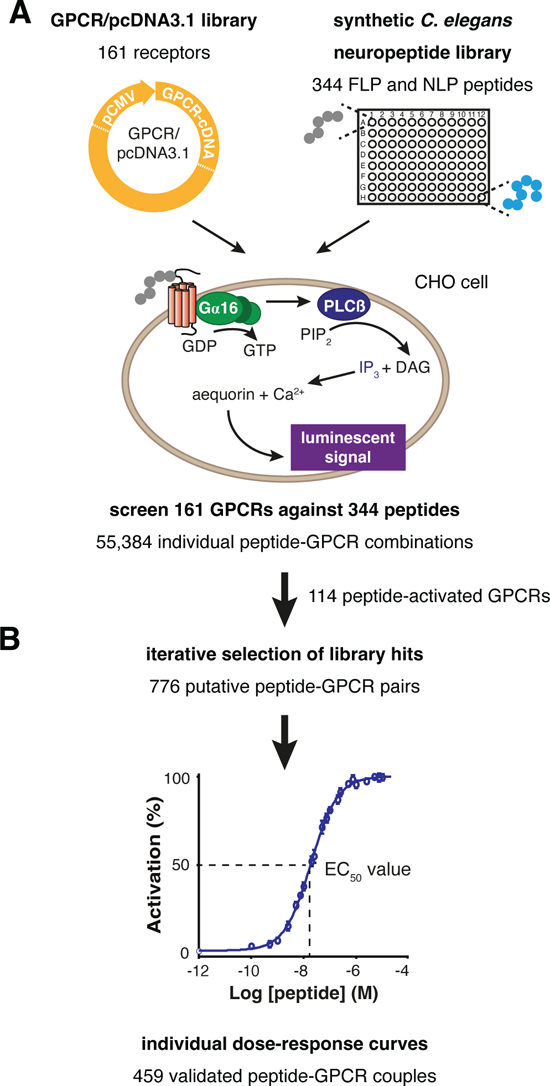
Large-scale deorphanization of *C. elegans* neuropeptide GPCRs. (A) Reverse pharmacology platform for deorphanization of *C. elegans* peptide GPCRs using a luminescence-based calcium mobilization assay. Peptide GPCR candidates predicted in the *C. elegans* genome are cloned in a pcDNA3.1 expression vector, under control of the human cytomegalovirus immediate early promoter (pCMV), for heterologous expression in mammalian cells. Each GPCR is expressed in CHO cells, stably co-expressing the calcium-activated photoprotein aequorin and the promiscuous human Gα_16_ subunit, the latter of which couples activation of most agonist-induced GPCRs to the calcium pathway. Cells are screened with a synthetic library of *C. elegans* peptides, including 344 FLP and NLP peptides. Upon GPCR activation, phospholipase Cβ (PLCβ) is activated and hydrolyzes phosphatidylinositol bisphosphate (PIP_2_) into diacylglycerol (DAG) and inositol trisphosphate (IP_3_), which activates IP_3_-dependent calcium channels, resulting in calcium release from intracellular storage sites. Binding of calcium to aequorin generates a luminescent signal by which GPCR activation is monitored. Screening of 55,384 individual peptide-GPCR combinations identified potential neuropeptide ligands for 114 out of 161 peptide GPCR candidates. See also Figure S1 and Data S1-S4 for an overview of the peptide library, neuropeptide receptor candidates, and peptide-activated GPCRs. (B) For each GPCR, peptide hits are iteratively selected from library screens, and each neuropeptide-GPCR interaction is validated by measuring receptor activation in response to a concentration range of purified peptide. A half maximal effective concentration (EC_50_ value) is calculated from the resulting dose-response curve to determine the potency of interactions. Out of 776 selected hits, 459 peptide-GPCR pairs show dose-dependent receptor activation. See Figures S2-S7 and Data S4-S6 for an overview of peptide-GPCR couples tested and validated in dose-response assays.

To identify neuropeptide-receptor couples, we screened each GPCR with a comprehensive library of 344 synthetic *C. elegans* peptides, belonging to the FMRFamide-like (FLP) and neuropeptide-like protein (NLP) families, that we compiled from biochemical peptide identifications and bioinformatic neuropeptide predictions (Data S3). Insulin-like peptides were not included in the library, as they mainly activate receptor tyrosine kinases and are structurally less well characterized in *C. elegans*^79, 80^. To monitor GPCR activation, we used an established calcium mobilization assay in Chinese hamster ovary (CHO) cells (Figure 1A), which provides a robust heterologous system for deorphanizing neuropeptide GPCRs^6, 81, 82^. Each receptor was transiently expressed in CHO cells that stably expressed the calcium-sensitive bioluminescent protein aequorin together with a promiscuous Gα_16_ protein. Gα_16_ allows coupling of nearly all GPCRs to stimulate the phospholipase Cβ (PLCβ) pathway, leading to an increase in intracellular calcium concentration upon receptor activation^83–85^. In total, we tested 55,384 individual peptide-receptor interactions by challenging each GPCR with the synthetic peptide library at high peptide concentrations (10 μM) and monitored calcium responses on a FLIPR^®^ high-throughput screening system (Figure 1A).

To identify hits in the peptide library, we manually inspected peptide-evoked calcium responses and determined an activation value for each peptide-GPCR pair based on Z-scores (see STAR Methods). We calculated Z-scores for each peptide-receptor pair, standardized the scores for each GPCR by dividing by its maximal activation, and log2-transformed this fraction to rank individual peptide-GPCR couples (Data S4). Known peptide-GPCR couples with nanomolar half-maximal effective concentrations (EC_50_ < 500 nM) nearly all showed Z-scores above 20 and log2-standardized values above −1.2 in our assays (Figure S1A and Data S4). Therefore, we considered all peptide-GPCR couples with a strong calcium response over background (Z-score > 20), reaching at least 40% of that of the receptor’s maximum Z-score (log2-standardized Z > −1.2), as potential hits in the peptide library. Based on these criteria, we identified 416 peptide hits for 114 GPCRs, derived from 81 neuropeptide- and 95 receptor-encoding genes (Figure S1 and Data S4).

### Validation and potency of neuropeptide-GPCR interactions

Peptide hits were identified using high peptide concentrations (10 μM); however, neuropeptide GPCRs typically show dose-dependent activation with EC_50_ values in the nanomolar range^43^. To determine whether the identified peptide hits are cognate GPCR ligands, we quantified GPCR activation for decreasing peptide concentrations (Figure 1B). To ensure accurate dose-response measurements, synthetic peptides were first purified by reversed phase high performance liquid chromatography (HPLC) and verified by mass spectrometry to obtain peptide stocks of high purity.

We then performed dose-response tests for all 416 hits identified in the library screens. Because false-positive ligand-receptor pairs could bias hit selection, we assigned hits by iterative ranking of peptide-GPCR couples. When the top peptide-GPCR interaction was not confirmed by dose-response tests with purified peptides, we restandardized the Z-scores for that receptor relative to the next top hit in the library screen. Dose-response assays were then repeated until all peptide-GPCR pairs matching the hit criteria were tested (Z-score > 20, log2-standardized Z > −1.2). For each hit, we included all peptides derived from the same neuropeptide precursor, which may be co-released from peptidergic cells, in dose-response tests (except for promiscuous receptors, see STAR Methods). In total, we tested 776 peptide-GPCR couples, of which 459 pairs showed dose-dependent interactions (Figure 1B, Figures S2-S7 and Data S4). None of the neuropeptide ligands evoked a calcium response in CHO cells transfected with an empty pcDNA3.1 plasmid (Data S5).

Among the 459 validated pairs, we identified ligands for 66 GPCRs (including isoforms) and receptors for 151 neuropeptides (Data S4). We characterized GPCRs for neuropeptides of all 31 RFamide-encoding (*flp*) genes, as well as peptides derived from 22 *nlp* encoding genes (Table 1 and Figures S2-S7). Among these, 39 receptors were activated only by RFamide neuropeptides, 24 GPCRs interacted with peptides of the NLP family, and 3 receptors were activated by neuropeptides from both families. For each GPCR, the potency of neuropeptide ligands was determined by calculating EC_50_ values from dose-response curves. These ranged between 0.1 pM and 22 μM but were in the nanomolar range for 74% of the interactions (Figure 2A and Data S6), suggesting that most of the identified interactions are likely cognate peptide-receptor pairs. To probe this, we set up a collaborative network through which *C. elegans* researchers could prioritize GPCRs for deorphanization and functionally validate peptide-receptor interactions *in vivo* (The Peptide-GPCR Project, https://worm.peptide-gpcr.org). Functional studies validated several neuropeptide-GPCR pairs discovered *in vitro* and, guided by receptor interactions, characterized novel roles of neuropeptides in the regulation of food searching, nociceptive responses, locomotory arousal, aversive learning and other behaviors^72, 73, 86–89^. Here, we present the complete resource of neuropeptide-GPCR interactions (Figures S2-S7); neuropeptide-receptor couples that were validated and reported in functional studies are listed in Table 1.

**Figure 2.**
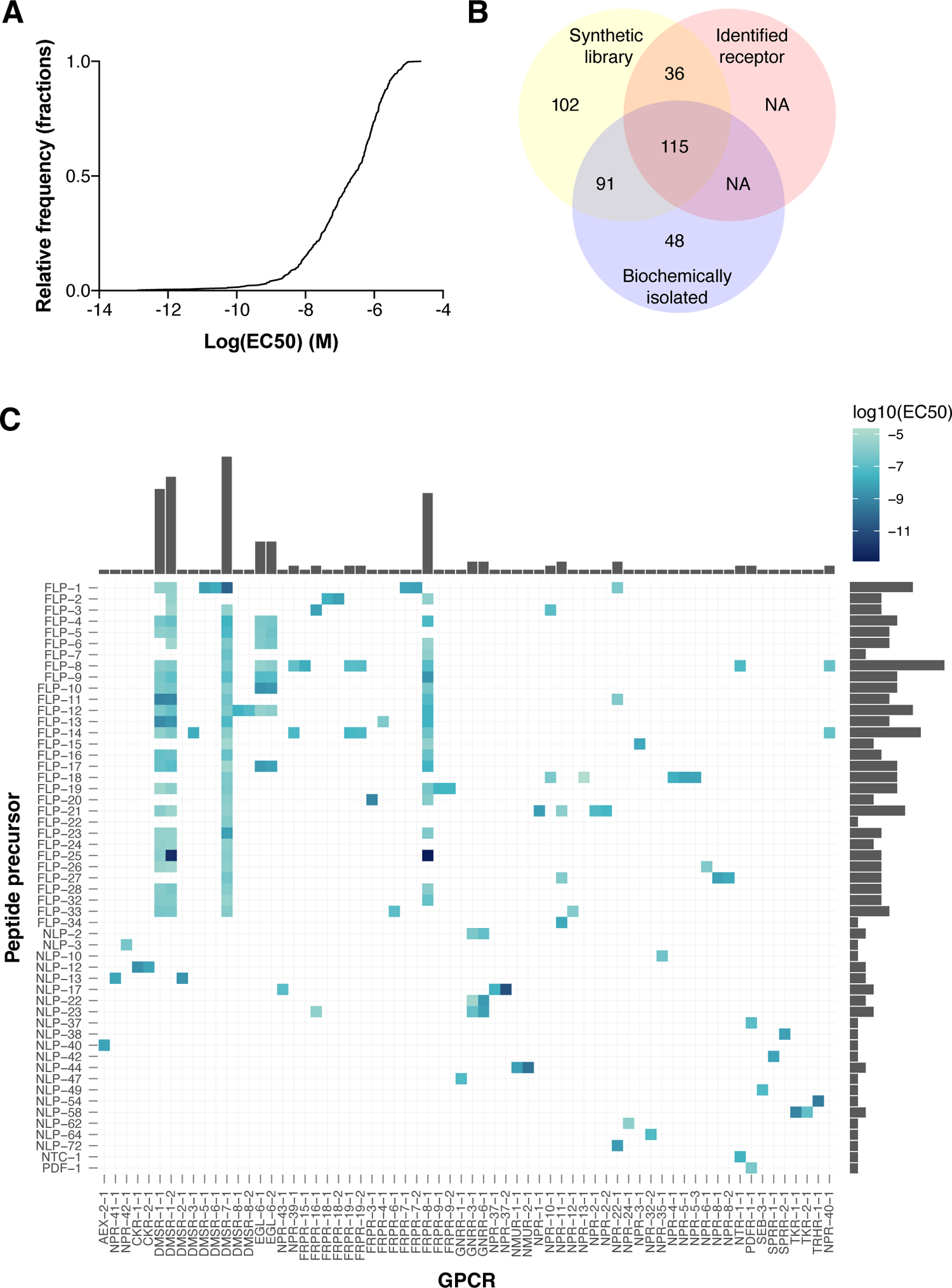
System-wide peptide-GPCR screening identifies a complex network of potent neuropeptide-receptor interactions. (A) Cumulative frequency plot for log(EC_50_) values of 459 identified neuropeptide-GPCR couples. 74% of peptide-receptor pairs are high potency interactions with EC_50_ values in the nanomolar range (log(EC_50_) < −6). See Data S6 for a list of all EC_50_ values. (B) Venn diagram of neuropeptide sequences in the synthetic peptide library that have been biochemically isolated using peptidomics^69^ and/or for which a receptor has been identified. For 36 predicted peptides (see Data S7), the identification of a receptor indicates a bioactive role. (C) Heatmap of log10(EC_50_) values for validated neuropeptide-GPCR pairs. For GPCRs interacting with multiple peptides from one precursor, only the most potent neuropeptide-GPCR interaction for each precursor is included. Histograms indicate the number of interactions for each GPCR (top) or neuropeptide precursor (right), respectively. See also Figures S2-S7 and Data S6 for dose-response curves and EC_50_ values.

**Table 1.** Deorphanized neuropeptide GPCRs in *C. elegans*

| Receptor | Known ligands (EC<sub>50</sub> range)<sup>a,c</sup> | Ligands in this resource<sup>b,c</sup> | In vivo interactions<sup>d</sup> |
| --- | --- | --- | --- |
| AEX-2 | **NLP-40-2** (3 nM) <sup>53</sup> | NLP-40-2 | *nlp-40* <sup>53</sup> |
| CKR-1 | NLP-18-2-NH<sub>2</sub> (13 nM) <sup>54</sup> | NLP-12-(1,2) <sup>87</sup> | *nlp-12* <sup>87</sup> |
| CKR-2 | **NLP-12-(1,2)** (15-28 nM) <sup>99</sup> | NLP-12-(1,2) | *nlp-12* <sup>146</sup> |
| DMSR-1-1 | **FLP-13-(1-7)** (2-65 nM) <sup>50</sup> | FLP-1-6; FLP-4-(1,2); FLP-5-(1-3); FLP-8-(1,2); FLP-9-(1,2); (pyro)FLP-10-1; FLP-11-(1-4); FLP-12-(1,2); FLP-13-(1-7); FLP-14-2; FLP-16-(1,2); FLP-17-(1-4); FLP-19-(1,2); FLP-21-1; FLP-23-3; FLP-24-1; FLP-25-(1,2); FLP-26-5; FLP-28-1; FLP-32-1; FLP-33-2 | *flp-13* <sup>50</sup> |
| DMSR-1-2 | - | FLP-1-6; FLP-2-2; FLP-3-8; FLP-4-(1,2); FLP-5-(1-3); FLP-6-2; FLP-8-(1,2); FLP-9-(1,2); (pyro)FLP-10-1; FLP-11-(1-3); FLP-12-(1,2); FLP-13-(1-7); FLP-14-2; FLP-16-(1,2); FLP-17-(1-4); FLP-19-(1,2); FLP-21-1; FLP-23-3; FLP-24-1; FLP-25-(1,2); FLP-26-5; FLP-28-1; FLP-32-1; FLP-33-2 | - |
| DMSR-2 | - | (pyro)NLP-13-(1-7) | - |
| DMSR-3 | - | FLP-14-(1,2) | - |
| DMSR-5 | - | FLP-1-(1-7,10) | - |
| DMSR-6 | - | FLP-1-(1-7,10) | - |
| DMSR-7 | - | FLP-1-(1-10) <sup>117</sup>; (pyro)FLP-3-(3, 6); FLP-4-(1,2); FLP-5-(1-3); FLP-6-2; FLP-7-(1-3,5); FLP-8-(1,2); FLP-9-(1,2); FLP-10-1; FLP-11-(1-4); FLP-12-(1,2); FLP-13-(1-7); FLP-14-2; FLP-15-3; FLP-16-2; FLP-17-4; FLP-18-7; FLP-19-(1,2); FLP-20-1; FLP-21-1; FLP-22-1; (pyro)FLP-23-(1-3); FLP-24-1; FLP-25-1; FLP-26-2; FLP-27-4; FLP-28-1; FLP-32-1; FLP-33-2 | - |
| DMSR-8-(1,2) | - | FLP-12-(1,2) | - |
| EGL-6-(1,2) | **FLP-10-1** (11 nM); **FLP-17-(1,2)** (1-28 nM) <sup>120</sup> | FLP-4-(1,2); FLP-5-(1-3); FLP-6-(1,2); FLP-8-(1,2); FLP-9-(1,2); (pyro)FLP-10-1; FLP-12-1; FLP-17-(1-4) | *flp-10*,<br>*flp-17* <sup>119,120</sup> |
| FRPR-3 | FLP-7-2 (1 μM); FLP-11-1 (1.3 μM); **FLP-20-(1-3)** (1-3 nM) <sup>55,147</sup> | FLP-20-(1-3) | *flp-11*,<br>*flp-20* <sup>55,148</sup> |
| FRPR-4 | **FLP-13-(1-7)** (67-541 nM) <sup>51</sup> | FLP-13-(1-7) | - |
| FRPR-6 | - | FLP-33-(1,2) | - |
| FRPR-7-(1,2) | **FLP-1-6** (10 nM) <sup>56</sup> | FLP-1-(1-7,10) | - |
| FRPR-8 | - | FLP-2-1; FLP-4-(1,2); FLP-6-2; FLP-7-(1-3); FLP-8-(1,2); FLP-9-(1,2); (pyro)FLP-10-1; FLP-11-(1-3); FLP-12-(1,2); FLP-13-(1-7); FLP-14-(1,2); FLP-15-1; FLP-16-(1,2); FLP-17-(1-4); FLP-19-(1,2); FLP-20-3; FLP-23-3; FLP-25-2; FLP-28-1; FLP-32-1 | - |
| FRPR-9-(1,2) | - | FLP-19-(1,2) | - |
| FRPR-15 | - | FLP-8-1 | - |
| FRPR-16 | - | (pyro)FLP-3-(1-5,7-9) <sup>149</sup>, NLP-23-1 | - |
| FRPR-18-(1,2) | **FLP-2-(1,2)** (4-54 nM) <sup>150,151</sup> | FLP-2-(1,2) | *flp-2* <sup>152</sup> |
| FRPR-19-(1,2) | - | FLP-8-1; FLP-14-1 <sup>88</sup> | *flp-14* <sup>88</sup> |
| GNRR-1 | **pyroNLP-47-1** (150 nM) <sup>98</sup> | pyroNLP-47-1 | - |
| GNRR-3 | - | NLP-2-(2,3); NLP-22-1; NLP-23-2 <sup>91</sup> | *nlp-2* <sup>91</sup> |
| GNRR-6 | - | NLP-2-(1-3); NLP-22-1; NLP-23-2 <sup>91</sup> | *nlp-2*,<br>*nlp-22* <sup>91</sup> |
| NMUR-1 | - | NLP-44-(1-3) <sup>73</sup> | *nlp-44* <sup>73</sup> |
| NMUR-2 | **NLP-44-3** (18 nM) <sup>101</sup> | NLP-44-(1-3) | - |
| NPR-1 | FLP-14-1 (24 nM); FLP-15-2 (22 nM); FLP-18-5 (1.8 μM); FLP-18-(2-4,6,7) (ND); **FLP-21-1** (1-207 nM); FLP-34-2 (578 nM) <sup>57,153-155</sup> | FLP-21-1 | *flp-18*, *flp-21*<br><sup>153,156,157</sup> |
| NPR-2-(1,2) | **FLP-21-1** (34-35 nM) <sup>49</sup> | FLP-21-1 | - |
| NPR-3 | **FLP-15-(1-3)** (56 – 599 nM); FLP-21-1 (50 nM) <sup>57,154,158</sup> | FLP-15-(1-3) | - |
| NPR-4 | FLP-1-6 (0.4-9.0 μM); FLP-4-2 (5-80 nM); FLP-14-1 (2.4 μM); FLP-15-2 (170 nM); **FLP-18-(2-7)** (3-1100 nM); FLP-21-1 (358 nM) <sup>57,154,159</sup> | (pyro)FLP-18-(1-8) | *flp-1*, *flp-18*<br><sup>159,160</sup> |
| NPR-5-(1,3) | FLP-1-2 (1.7-4.7 μM); FLP-3-(1,7,8) (0.9-2.9 μM); FLP-15-2 (6.6 μM); **FLP-18-(2-7)** (13-819 nM); FLP-21-1 (0.2-5.0 μM); FLP-34-2 (2.0 μM) <sup>57,154,159,161</sup> | (pyro)FLP-18-(1-8) | *flp-18*<br>156,159,162 |
| NPR-6 | FLP-1-10 (ND); FLP-14-1 (1.2 μM); FLP-15-(1,2) (1.5-3.0 μM); FLP-18-5 (425 nM); FLP-21-1 (409 nM) <sup>56,57,154</sup> | FLP-26-2 | *flp-1* <sup>56</sup> |
| NPR-8-(1,2) | - | (pyro)FLP-27-(1,3-4) | - |
| NPR-10 | **FLP-3-(1,3,5,7,8)** (60-300 nM); **FLP-18-(3-7)** (0.06-4.6 μM) <sup>154</sup> | (pyro)FLP-3-(1-5,7-9) <sup>149</sup>; (pyro)FLP-18-(1-8) | - |
| NPR-11 | NLP-1-1 (ND); FLP-1-6 (1-8 μM); FLP-5-2 (1-8 μM); FLP-14-1 (1-8 μM); FLP-15-2 (998 nM); FLP-18-(3,5) (80-800 nM); **FLP-21-1** (1-102 nM); FLP-33-1 (414 nM); **FLP-34-(1-2)** (0.7-19 nM) <sup>57,154,163</sup> | FLP-21-1; (pyro)FLP-27-(1-4); FLP-34-(1-4) <sup>72</sup> | *nlp-1, flp-34*<br><sup>72,163</sup> |
| NPR-12 | NLP-29 (ND) <sup>58</sup> | FLP-33-1 | *nlp-29* <sup>58,164</sup> |
| NPR-13 | - | FLP-18-7 | - |
| NPR-17 | NLP-24-6 (ND) <sup>48</sup> | - | *nlp-24* <sup>48,165</sup> |
| NPR-22 | **FLP-1-8 (ND)**; FLP-7-3 (1.1 μM); FLP-9-1 (ND); **FLP-11-(1-3)** (ND); FLP-13-(2,4-7) (ND); FLP-22-1 (ND); **NLP-72-(1,2)** (14-22 nM) <sup>52,166</sup> | FLP-1-(8,9); FLP-11-(1,2); NLP-72-(1,2) <sup>89</sup> | *flp-7, flp-11, nlp-72* <sup>52,89,148</sup> |
| NPR-24 | - | NLP-62-1 | - |
| NPR-28 | RGBA-1 (ND) <sup>59</sup> | - | *rgba-1* <sup>59</sup> |
| NPR-32 | - | NLP-64-2 | - |
| NPR-34 | - | SNET-1-1 | - |
| NPR-35 | - | NLP-10-(1-4) | - |
| NPR-37-(1,2) | - | (pyro)NLP-17-(1-3) | - |
| NPR-39 | - | FLP-8-1; FLP-14-1 | - |
| NPR-40 | - | FLP-8-1; FLP-14-1 | - |
| NPR-41 | - | (pyro)NLP-13-(1-7) | - |
| NPR-42 | - | NLP-3-3 | - |
| NPR-43 | - | pyroNLP-17-(2,3) | - |
| NTR-1 | **NTC-1-1** (19-83 nM) <sup>6,100</sup> | NTC-1-1; FLP-8-1 | - |
| NTR-2 | NTC-1-1 (ND) <sup>100</sup> | - | - |
| PDFR-1 | **NLP-37-1** (34 nM); **PDF-1-(1,2)** (127-361 nM) <sup>78</sup> | NLP-37-1; PDF-1-1 | - |
| SEB-2 | - | NLP-73-1 | - |
| SEB-3 | - | NLP-49-2 <sup>86</sup> | *nlp-49* <sup>86</sup> |
| SPRR-1 | - | NLP-42-(1,2) | - |
| SPRR-2 | **NLP-38-(1-4,6)** (8-486 nM) <sup>61</sup> | NLP-38-(1-4,6) | *nlp-38* <sup>61</sup> |
| TKR-1 | **NLP-58-(1-3)** (2-8 nM) <sup>60</sup> | NLP-58-(1-3) | - |
| TKR-2 | - | NLP-58-(1-3) | - |
| TRHR-1 | **NLP-54-(1,2)** (0.3-0.5 nM) <sup>5</sup> | NLP-54-(1,2) | *nlp-54* <sup>5</sup> |

<sup>a</sup> Known peptide ligands with EC<sub>50</sub> values below 10 μM. The range of reported EC<sub>50</sub> values is indicated between brackets (ND, not determined). Ligands in bold were also identified in this resource.
<sup>b</sup> Peptide ligands identified in this resource. EC<sub>50</sub> values of all peptide-receptor interactions are in Data S6. References cite reports of neuropeptide-receptor couples in functional *in vivo* studies.
<sup>c</sup> Peptide sequences of ligands are summarized in Data S3.
<sup>d</sup> Ligand(s) identified *in vitro* that were shown to act in the same genetic pathway as their receptor *in vivo*.

### System-wide deorphanization captures novel ligands for known neuropeptide GPCRs

Deorphanization efforts in the past two decades identified neuropeptide-receptor couples for 16 *nlp*, 19 *flp*, and 29 peptide GPCR encoding genes in *C. elegans* (Table 1). We compared these reported interactions to the 459 peptide-GPCR couples discovered in our screen. We confirmed known agonists for 23 (79%) of the characterized peptide GPCRs (Table 1). Our results recapitulated the majority of known, high-potency interactions with an EC_50_ value below 500 nM (Table 1). Most of the previously reported ligand-receptor couples that were not validated in our screen have EC_50_ values in the micromolar range (Table 1); it remains unclear if these interactions are relevant at physiological neuropeptide concentrations.

Besides known neuropeptide-receptor pairs, we also discovered novel neuropeptide ligands for six receptors that were deorphanized previously. By systematically screening GPCRs with the synthetic peptide library, we identified 63 additional ligands for DMSR-1, NPR-6, NPR-11, NTR-1 and two EGL-6 receptors (Table 1). This underscores the importance of system-wide screening approaches to map neuropeptide-GPCR signaling networks.

### Deorphanization of novel peptide GPCRs and discovery of receptors for predicted neuropeptides

Using a system-wide reverse pharmacology approach, we more than doubled the number of deorphanized peptide GPCRs in *C. elegans*. In total, we deorphanized 35 neuropeptide GPCRs for which no ligand was known previously (Table 1), increasing the number of characterized peptide GPCRs from 29 to 58 receptors (not considering isoforms) in *C. elegans*. These include receptors for neuropeptides derived from 58 neuropeptide-encoding genes (31 *flp* and 27 *nlp* genes).

We also asked whether our deorphanization screen might reveal receptors for predicted neuropeptides that have as yet not been biochemically isolated. The identification of their receptor(s) would suggest a biological function for these peptides. To probe this, we compared our dataset of peptide-GPCR couples to the *C. elegans* neuropeptidome, a resource of peptide sequences identified by mass spectrometry^69^. Of the 344 peptide sequences in our synthetic library, 206 peptides were detected in whole-mount peptide extracts using mass spectrometry. Among these, 115 neuropeptides dose-dependently activated one or multiple GPCRs in our deorphanization screen (Figure 2B). In addition, we identified receptors for 36 neuropeptides that so far have not been biochemically isolated (Figure 2B and Data S7). Of these peptides, 32 activated receptors with high potency and EC_50_ values comparable to those of peptide-GPCR couples validated *in vivo* (Table 1), which suggests bioactive roles.

### The RFamide neuropeptide-receptor network is characterized by peptidergic crosstalk

Neuropeptides of the FLP family share a C-terminal RFamide motif, whereas NLPs represent a structurally diverse class of peptides that lack a common consensus sequence. In agreement with these structural differences, we found that most *C. elegans* neuropeptide GPCRs are activated by either FLP or NLP ligands. Nevertheless, we identified three receptors (NTR-1, FRPR-16 and NPR-22) that have ligands from both neuropeptide families. In the case of NPR-22, these ligands show clear sequence similarity; NLP ligands of this receptor (NLP-72-1: PALLSRYamide; NLP-72-2: AVLPRYamide) have a C-terminal RYamide motif that resembles the RFamide sequence of FLP ligands (Figure S2 and Data S3). By contrast, the oxytocin/vasopressin receptor ortholog NTR-1 and the FMRFamide-like receptor FRPR-16 are activated by structurally diverse ligands. We found that NTR-1 is activated by the cyclic nematocin peptide (NTC-1-1: CFLNSCPYRRYamide), an oxytocin/vasopressin ortholog, as well as the RFamide neuropeptide FLP-8 (FLP-8-1: KNEFIRFamide), which both activate NTR-1 with similar EC_50_ values (Figure S3 and Data S6). The FRPR-16 receptor interacts with RFamide (FLP-3) neuropeptides as well as the non-amidated NLP-23 peptide (NLP-23-1: LYISRQGFRPA), albeit at different potencies (Figure S4 and Data S6). These receptors may have different interaction sites for diverse ligands.

To further probe the diversity of ligands for individual neuropeptide GPCRs, we generated a heatmap of EC_50_ values for the identified neuropeptide-receptor pairs (Figure 2C). The majority of receptors (48/66 deorphanized GPCRs) were activated by neuropeptides from a single peptide precursor, showing specific ligand-receptor interactions. However, we also identified six GPCRs with ligands from at least eight or more distinct neuropeptide-encoding genes (Figure 2C). These promiscuous receptors are encoded by *dmsr-1*, *dmsr-7*, *frpr-8* and *egl-6* genes, and are all activated by diverse RFamide neuropeptides. Most RFamide peptides also activate multiple GPCRs encoded by different receptor genes (Figure 2C). FLP-1, FLP-8 and FLP-14 are among the most versatile neuropeptides that interact with eight to twelve GPCRs. By contrast, neuropeptides of the NLP family activate only a few (one to three) receptors (Figure 2C). The RFamide signaling network thus shows a higher level of ligand-receptor crosstalk in comparison to non-RFamide (NLP) neuropeptide systems. RFamide neuropeptides also interact with a receptor network that is largely distinct from that of other neuropeptides.

We further investigated the structure of FLP and NLP ligand-receptor networks using a bipartite network representation and its projections. We focused our analysis on interactions with nanomolar EC_50_ values (< 1 μM), which comprised most peptide-GPCR couples in our dataset. Bipartite graphs of both networks showed different topologies and more crosstalk for ligands and receptors of the RFamide family than for other neuropeptide signaling systems (Figure 3A). To investigate the relationships among FLP and NLP peptides, we projected the bipartite networks into simple, monopartite ligand networks in which the nodes (peptides) are connected only if they have at least one common receptor. Nearly all clusters in the NLP network include peptides from a single precursor, which often have similar sequences and activate the same receptor(s) (Figure 3B). The only two exceptions are the *C. elegans* RPamide peptides (encoded by *nlp-2*, *nlp-22* and *nlp-23* genes) and PDF-like neuropeptides (derived from *pdf-1* and *nlp-37* genes). Both are families of homologous neuropeptide genes, which likely arose through gene duplication and share significant sequence similarity^90, 91^. By contrast, the monopartite projection of the FLP network shows a higher number of edges, consistent with a high level of crosstalk between RFamide peptides and their receptors (Figure 3C).

**Figure 3.**
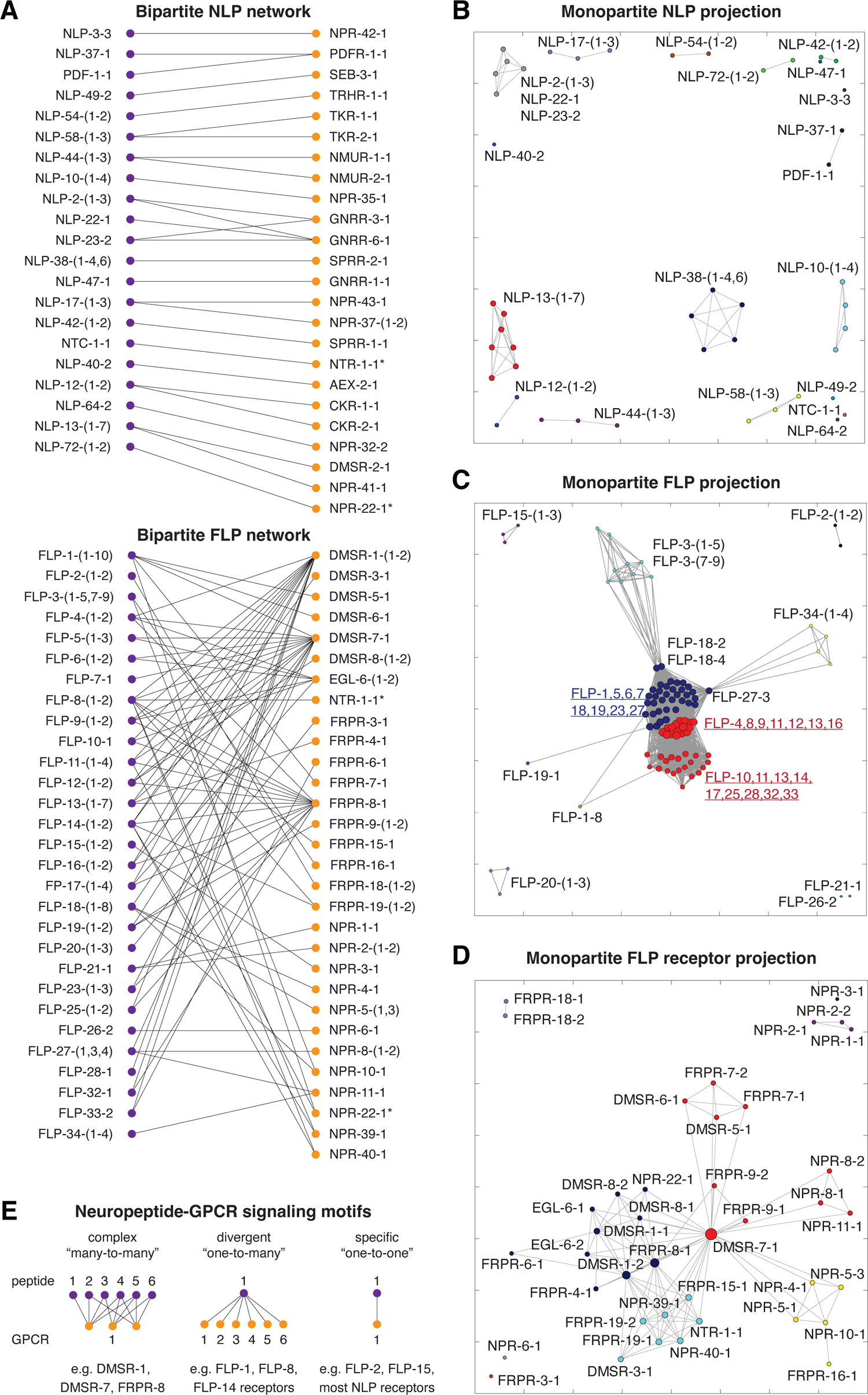
Specific and complex combinatorial interactions in the peptidergic signaling network. (A) Bipartite graphs of the NLP and FLP neuropeptide-GPCR networks. Nodes represent peptides (purple) and receptors (orange). Edges between two nodes depict neuropeptide-GPCR couples with nanomolar EC_50_ values (EC_50_ < 1 μM) identified in this resource. Asterisks indicate receptors included in both NLP and FLP signaling networks. (B-C) Monopartite peptide projections of the NLP (B) and FLP (C) bipartite neuropeptide-GPCR networks. Nodes are sized by the total number of connections (degree) and colored by module. (D) Monopartite receptor projection of the FLP bipartite neuropeptide-GPCR network. Nodes are sized by degree and colored by module. (E) Different types of neuropeptide-receptor interaction motifs can be distinguished in the peptidergic network: 1) complex “many-to-many” interactions mediated by promiscuous neuropeptide GPCRs that have a large number of different ligands, interacting with multiple receptors, 2) diverging “one-to-many” interactions involving a single neuropeptide that activates multiple specific GPCRs, and 3) specific “one-to-one” interactions between a single neuropeptide and its receptor.

We further analyzed the modularity of the FLP ligand network and identified several groups of highly interconnected nodes that form modules within the signaling network (Figure 3C). The majority of RFamide neuropeptides clustered in two modules based on their activation of two promiscuous GPCRs, DMSR-1 and DMSR-7. One module comprised all ligands that activate DMSR-7 but not DMSR-1 (dark blue in Figure 3C). A second module grouped all ligands that activate DMSR-1 (red in Figure 3C), of which some – closest to the DMSR-7 module – also activate DMSR-7. Most of the RFamide neuropeptides in *C. elegans* thus can be categorized based on their interactions with these promiscuous GPCRs. DMSR-1 and DMSR-7 share a set of common RFamide ligands, but each receptor also uniquely interacts with other RFamide peptides, which often activate other FLP receptors as well. In addition, we found eight smaller groups of RFamide neuropeptides that form separate modules in the ligand network. Several of these peptides, including FLP-15, FLP-21, FLP-26 and FLP-34, belong to the evolutionarily conserved families of short neuropeptide F (sNPF) and neuropeptide Y/F (NPY/F) peptides (see below). These neuropeptide systems arose in a common ancestor of bilaterian animals^42, 92^, and through ligand-receptor co-evolution may have evolved interactions with different receptor targets than those of other RFamide neuropeptides.

We discovered similar modules in a receptor-focused projection of the bipartite FLP network, in which nodes (receptors) are connected when they share at least one peptide ligand. Two modules are centred around the promiscuous receptors DMSR-1 and DMSR-7 (Figure 3D). Other receptors that share ligands with these promiscuous GPCRs cluster in the two modules as well. For example, DMSR-7 is potently activated by FLP-1 peptides and clusters together with other FLP-1 receptors (DMSR-5, DMSR-6 and FRPR-7). We also found seven modules distinct from the DMSR-1 and DMSR-7 clusters. Consistent with the ligand projection, receptors of the sNPF receptor family (NPR-1, NPR-2, NPR-3, NPR-4, NPR-5, NPR-6 and NPR-10) clustered into separate modules, as they share none or only a few ligands with other GPCRs. The FLP receptor projection also revealed a novel module in the FLP signaling network, consisting of seven GPCRs that are uniquely activated by FLP-8 and FLP-14 peptides (DMSR-3, FRPR-15, FRPR-19-1/2, NTR-1, NPR-39 and NPR-40) (Figure 3D).

Taken together, our results indicate three main types of ligand-receptor interactions in the neuropeptide signaling network, which may serve specialized functions (Figure 3E): 1) complex “many-to-many” interactions mediated by promiscuous receptors (e.g. DMSR-1 and DMSR-7), which have diverse RFamide neuropeptide ligands that often activate other GPCRs as well, 2) divergent “one-to-many” interactions by RFamide neuropeptides (e.g. FLP-1, FLP-8 and FLP-14) that activate multiple specific receptors, and 3) specific “one-to-one” interactions between peptides and receptors from single neuropeptide- and GPCR-encoding genes. Most non-RFamide systems belong to this category, whereas the majority of RFamide neuropeptides cross-interact with many GPCRs.

### GPCR isoforms diversify neuropeptide-receptor interactions

Signaling properties can differ between GPCR isoforms encoded by the same receptor gene^37, 93^. The extent to which GPCR isoforms contribute to functional diversity of ligand-receptor interactions, however, remains largely unexplored. To probe this, we compared neuropeptide-GPCR interactions for receptor isoforms of 11 GPCR genes that differ at N-terminal, C-terminal or intertransmembrane regions (Figure 4A). Most isoforms of a specific receptor displayed similar ligand profiles (Figure S8). However, two C-terminal variants of the *Drosophila* myosuppressin receptor ortholog DMSR-1 showed distinct interactions. We identified neuropeptides of the RFamide gene *flp-25* as potent ligands of DMSR-1B (DMSR-1-2), with picomolar EC_50_ values, whereas FLP-25 peptides activated the DMSR-1A (DMSR-1-1) isoform only at high micromolar concentrations (Figure 4B). DMSR-1 is a promiscuous RFamide receptor and most other ligands activated the two receptor isoforms with similar potencies. However, DMSR-1A was strongly activated by one of the FLP-11 peptides (FLP-11-4), which did not activate DMSR-1B. Besides DMSR-1, two N-terminal variants of the NLP-17 receptor NPR-37 also displayed a thousand-fold difference in EC_50_ value for their ligand (Figure 4C). These results suggest that GPCR isoforms can differ in their ligand interactions and further diversify the *C. elegans* neuropeptide signaling network.

**Figure 4.**
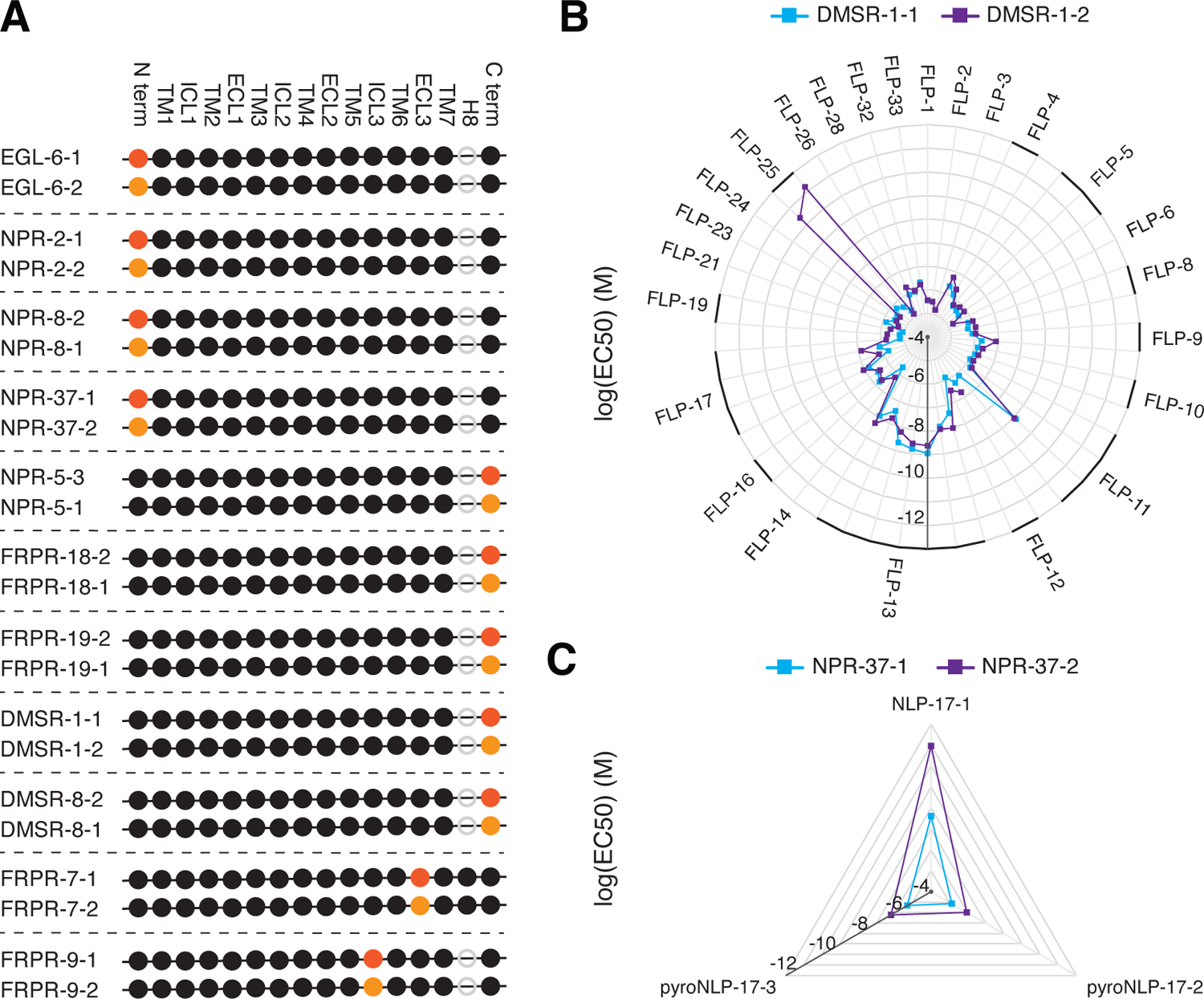
GPCR isoforms diversify neuropeptide-receptor interactions. (A) Structural fingerprints for deorphanized neuropeptide GPCRs with multiple isoforms. For each GPCR, the longest isoform is shown at the top. Black and orange dots indicate segments that are identical or different between two isoforms, respectively. Grey circles depict segments that are missing from the receptor sequence. N and C term, N and C terminus; TM, transmembrane region; ICL, intracellular loop; ECL, extracellular loop; H8, helix 8. (B and C) Protein variants of the neuropeptide GPCRs DMSR-1 and NPR-37 show different neuropeptide-receptor interactions. Radar plots depict log(EC_50_) values for each identified neuropeptide ligand. Neuropeptides of the RFamide precursor FLP-25 are potent ligands of DMSR-1-2 but not of DMSR-1-1, whereas DMSR-1-1 is specifically activated by FLP-11-4 (B). Two protein variants of the NLP-17 receptor NPR-37 display a thousand-fold difference in EC_50_ value for the NLP-17-1 peptide (C). See also Figures S8-S11 and Data S6.

### Neuropeptide genes encoding multiple peptides expand the peptidergic network

Besides GPCR isoforms, neuropeptide genes often encode multiple bioactive peptides that are generally thought to be co-synthesized and co-released^43, 94^. To probe how this multi-peptide signaling affects the neuropeptide-receptor network, we compared sequences and receptor interactions for peptides encoded by the same neuropeptide gene. Most neuropeptides derived from a single precursor show clear sequence similarity and activate the same receptor(s) (Figures S9-S11). However, we also found several neuropeptide precursors of which individual peptides differ in their potency to activate specific GPCRs. For example, peptides encoded by the RFamide neuropeptide gene *flp-1* show distinct GPCR interaction profiles that correlate with differences in neuropeptide sequence. FLP-1 peptides with the PNFLRF-NH_2_ motif (FLP-1-1 to 7 and FLP-1-10) all activate DMSR-5, DMSR-6 and FRPR-7. These receptors are not activated by the two PNFMRY-NH_2_ FLP-1 peptides (FLP-1-8 and FLP-1-9), which activate the NPR-22 receptor (Figure S10 and Data S3). FLP-11 peptides also differ in their receptor interactions (Figure S10), and similar differences were found for neuropeptides encoded by *nlp* genes (*e.g.* NLP-17, NLP-23 and NLP-44 peptides; Figure S11). These findings indicate that distinct neuropeptides in the same precursor often target similar receptors, which may amplify peptidergic signaling. Alternatively, they can further expand the peptidergic network by acting on different GPCRs in different target cells, so that multiple neuropeptide axes can signal across circuits when a peptidergic cell is activated.

### Characterization of ancestral bilaterian neuropeptide systems in *C. elegans*

Many neuropeptide systems show long-range evolutionary conservation and co-evolution of ligand-receptor pairs. Comparative genomic analyses of peptidergic systems across widely divergent animal phyla revealed at least 31 neuropeptide-receptor systems that are ancestral to bilaterian animals^42, 95–97^. These include well-studied peptidergic systems, such as oxytocin and vasopressin, cholecystokinin, neuropeptide Y (NPY), and gonadotropin-releasing hormone (GnRH), that already evolved in a common ancestor of protostomian and deuterostomian animals^5, 6, 42, 98, 99^. Several bilaterian systems are also thought to be conserved in *C. elegans*, but not all predicted neuropeptide-receptor couples have been experimentally demonstrated. To further clarify the conservation of neuropeptide systems in *C. elegans*, we performed a phylogenetic analysis of nematode GPCRs in our screen, identified receptor orthologs for bilaterian neuropeptide systems, and compared their neuropeptide ligands to peptides that activate known representatives of the same receptor family.

To reconstruct the phylogenetic relationships of nematode peptide GPCRs, we first used the sequences of all rhodopsin and secretin-type peptide GPCRs from species in 8 bilaterian phyla as bait to identify potential homologs in transcriptomic and EST datasets of 34 phylogenetically dispersed nematode species (Data S8). We then used these sequences to construct a maximum likelihood tree, which revealed 31 bilaterian receptor clusters, containing a diversified set of protostome and deuterostome GPCRs (Figures 5A-B, Data S10 and S11). Representative *C. elegans* receptors were present in 17 of the 31 ancestral receptor families (Figure 5C and Data S9). These GPCR families were well conserved in nematode species across different clades (Figure 5C). In addition, one receptor from *Trichinella spiralis* (clade 2) clustered in the bilaterian family of parathyroid hormone/glucagon/pituitary adenylate cyclase-activating polypeptide (PTH/GCG/PACAP) receptors, although this GPCR family was not represented in *C. elegans* or other nematode clades (Figure 5C).

**Figure 5.**
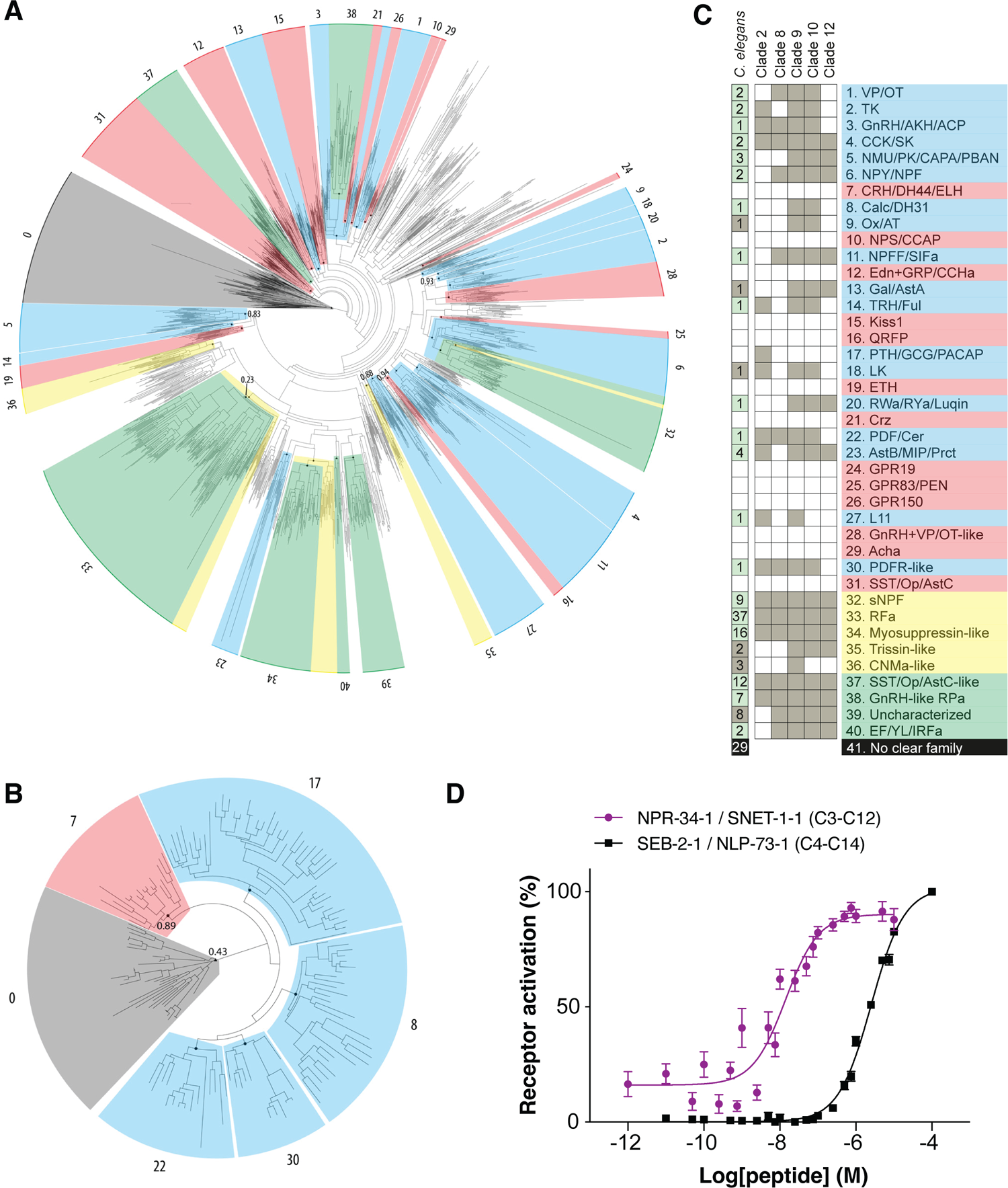
Biochemical peptide-GPCR interactions support conservation of bilaterian neuropeptide systems and expansions of peptide GPCR families in *C. elegans*. (A) Maximum likelihood tree of bilaterian rhodopsin peptide GPCRs. Subtrees are numbered according to the neuropeptide-receptor families in panel C. Subtrees comprising bilaterian families with nematode representatives are indicated in blue; those without nematode representatives in red. Subtrees containing only protostomian sequences are colored yellow, and nematode-specific subtrees are green. Node support values at the root of all delineated subtrees are above 0.95, unless depicted otherwise on the tree. Atypical peptide receptors (angiotensin, bradykinin, and chemokine receptors) were used as outgroup, depicted in grey. See also Data S10. (B) Maximum likelihood tree of bilaterian secretin receptors. Color-coding, numbering and node support values as in (A). Adhesion and cadherin receptors were used as an outgroup, depicted in grey. See also Data S11. (C) Inferred evolutionary relationships for the different nematode neuropeptide-receptor systems. Names of characterized neuropeptide systems are assigned according to the classification by Mirabeau and Joly (2013) and Elphick et al. (2018), and color-coded as in (A). The presence of a receptor ortholog is marked with a grey square. A receptor is considered present in a given nematode clade when it is positioned inside a well-supported subtree (branch support value > 0.95). The number of *C. elegans* receptors present in each family is depicted in the left column and squares are colored green if at least one receptor of the family is deorphanized. See also Data S8-S9 for nematode receptors identified in each family and full names of neuropeptide signaling systems. (D) The elevenin-like peptide SNET-1-1 (LDCRKFSFAPACRGIML; C3-C12) and calcitonin-like peptide NLP-73-1 (NRQCLLNAGLSQGCDFSDLLHAQTQARKFMSFAGPamide; C4-C14) dose-dependently activate the *C. elegans* elevenin and calcitonin receptor orthologs NPR-34 and SEB-2. Calcium responses of CHO cells expressing NPR-34 or SEB-2 are shown relative (%) to the highest value (100% activation) after normalization to the total calcium response. EC_50_ values (95% confidence interval) for SNET-1-1 and NLP-73-1 are 14.2 (9.8 – 20.6) nM and 2.5 (2.3 – 2.8) μM. Error bars represent SEM (n ≥ 6).

We deorphanized receptors in 12 of the 17 bilaterian families conserved in *C. elegans* (Figure 5C and Data S9). In most cases, the *C. elegans* receptor was activated by a peptide that was a previously recognized ortholog of the neuropeptide ligands of related GPCRs^42, 95, 96^. Consistent with previous reports, our biochemical data supports the conservation of ligand-receptor couples related to vasopressin/oxytocin (NTC-1/NTR-1), tachykinin (NLP-58/TKR-1), GnRH (NLP-47/GNRR-1), cholecystokinin (NLP-12/CKR-1/CKR-2), neuromedin U (CAPA-1/NMUR-1/NMUR-2), NPY (FLP-34/NPR-11), thyrotropin-releasing hormone (TRH-1/TRHR-1), luqin (LURY-1/NPR-22), pigment dispersing factor (PDF-1/PDF-2/PDFR-1), and myoinhibitory peptide (MIP-1/SPRR-2) signaling systems in *C. elegans*^5, 6, 52, 57, 60, 61, 72, 73, 87, 90, 98–101^. In addition, we deorphanized four novel *C. elegans* peptide GPCRs related to neuropeptide FF/SIFamide (NPR-35), tachykinin (TKR-2), NPY (NPR-12), and myoinhibitory peptide (SPRR-1) receptors (Data S9). The deorphanization of these receptors provides biochemical evidence for their predicted interactions with orthologs of neuropeptide FF/SIFamide (NLP-10), tachykinin (NLP-58), NPY (FLP-33) and myoinhibitory peptides (NLP-42)^42, 57, 60, 102^, which further supports evolutionary conservation of these neuropeptide systems in nematodes. Furthermore, we identified a ligand for SEB-3, a GPCR that belongs to the same bilaterian secretin superfamily of receptors as bilaterian CRH/DH44 and calcitonin/DH31 receptors, vertebrate VIP/PACAP receptors, and invertebrate PDF receptors^42, 86, 95, 96^. SEB-3 is most closely related to PDF receptors (Figure 5B) and orthologous to orphan GPCRs in many other species, including molluscs, annelids, echinoderms, hemichordates and cephalochordates^42, 96^. The identification of a neuropeptide ligand for SEB-3 may help to characterize the ligands and functions of other receptors within this conserved GPCR family. A functional study of this peptide-receptor pair validated their interaction *in vivo* and characterized a role of this signaling system in the regulation of locomotion, arousal and reproduction^86^.

Receptors in 5 bilaterian families conserved in *C. elegans* were not deorphanized in our reverse pharmacology screen. For the *C. elegans* galanin/allatostatin A (NPR-9) and leucokinin (TKR-3) receptor orthologs^42^, neuropeptides orthologous to allatostatin A (NLP-6) and leucokinin (NLP-43) were included in the synthetic library, but were not found to activate their predicted GPCR. Reasons for this lack of interaction include: (i) the functional expression of GPCRs in CHO cells may have failed or, (ii) in the case of NPR-9, the predicted receptor cDNA sequence could not be cloned (Data S1). For *C. elegans* orexin/allatotropin (NPR-14), elevenin (NPR-34), and calcitonin/diuretic hormone 31 (SEB-2) receptor orthologs, the predicted neuropeptide ligands were initially not included in our synthetic peptide library. *C. elegans* homologs of orexin/allatotropin (NLP-59) and elevenin (SNET-1) peptides were identified after we assembled the peptide library^96, 103^, whereas a calcitonin-like neuropeptide (NLP-73) was included but lacked a disulfide bridge that may be critical for receptor activation (Data S3). We therefore tested these receptors with their predicted neuropeptide ligands in the calcium mobilization assay and found dose-dependent activation for SNET-1/NPR-34 and NLP-73/SEB-2 ligand-receptor pairs (Figure 5D). SNET-1 and NLP-73 peptides specifically activated NPR-34 and SEB-2 receptors, respectively; these peptides did not evoke significant calcium responses in cells transfected with an empty control vector (Data S5). We did not observe activation of NPR-14 with NLP-59 peptides, which may be because this receptor did not express well in CHO cells or is activated by another, unidentified orexin/allatotropin-like neuropeptide. Taken together, our results biochemically support the conservation of at least 14 ancestral bilaterian neuropeptide systems in *C. elegans* (Figure 5C). Based on these findings, we suggest naming these neuropeptide and receptor encoding genes in accordance with their evolutionary relationship to bilaterian neuropeptide-systems (Data S9).

### Nematode expansions of evolutionarily conserved neuropeptide systems

Besides the conservation of ancestral bilaterian neuropeptide systems, phylogenetic analysis revealed large expansions of neuropeptide receptor families in nematodes, including GnRH, somatostatin, sNPF, myosuppressin, and FMRFamide receptor families (Figure 5A, Data S9 and S10). In each case, we found that receptors of these expanded families were activated by neuropeptides of the corresponding related peptide family, which supports co-evolution of neuropeptide-receptor couples and the conservation of these neuropeptide systems in nematodes.

A first expanded family of nematode receptors clustered near the bilaterian family of somatostatin (SST), allatostatin C (AstC) and opioid (Op) receptors (cluster 37, Figure 5A and Data S10). This group of 12 *C. elegans* GPCRs (Data S9) includes NPR-17 that was previously shown to interact with the opioid-like peptide NLP-24^48^. We found two other receptors in this family, NPR-24 and NPR-32, to interact with previously recognized orthologs of the SST/AstC neuropeptide family, NLP-62 and NLP-64 (Figure S2), supporting the conservation of SST/AstC-like receptors in *C. elegans*^42, 104^.

A second expansion occurred for GnRH-related receptors (cluster 38, Figure 5A and Data S10). The *C. elegans* receptor GNRR-1 clusters most closely to GnRH-like receptors in other animals and is specifically activated by NLP-47 (Figure S3 and Data S10), a neuropeptide ortholog of GnRH and the related insect adipokinetic hormone (AKH)^98^. In addition, the *C. elegans* genome encodes 7 GnRH-like receptors (GNRR-2 to GNRR-8) that form a paralogous group. Two of them are activated by RPamide peptides (Figure S3), a nematode family of neuropeptides that shows sequence similarity to GnRH and AKH peptides^91^, which is in agreement with an expansion of the GnRH receptor family.

Three protostomian families of neuropeptide receptors also largely expanded in nematodes. The largest expansion occurred in the FMRFamide-like receptor family, related to the *Drosophila* FMRFaR, including 37 GPCRs in *C. elegans* (cluster 33, Figure 5A, Data S9 and S10). As expected, all deorphanized receptors in this family are activated by RFamides (FLPs) or neuropeptides with a similar RIamide motif at the C-terminus (NLP-17) (Data S9). In addition, the protostomian sNPF and myosuppressin receptor families diversified in nematodes (clusters 32 and 34, Figure 5A and Data S10). The protostomian sNPF system was shown to be orthologous to deuterostomian prolactin releasing hormone signaling^92^. In *C. elegans*, this family includes nine receptors and ligands for eight of them are all sNPF-like neuropeptides with the canonical motif XLRFamide (FLP-15, FLP-18, FLP-21 and FLP-26; Figure S2, Data S9 and S10)^105^. The myosuppressin-like receptor family includes 16 *C. elegans* receptors (Data S9 and S10), of which three are activated by FLP-1 peptides that show sequence similarity to insect myosuppressins (Figure S6)^106^. Interestingly, these FLP-1 receptors – DMSR-5 to DMSR-7 – all cluster together in the phylogenetic tree along with several orphan GPCRs (DMSR-4, DMSR-9 and DMSR-11 to 16), whereas other myosuppressin-like receptors form separate sub-groups and are activated by different, more divergent neuropeptide ligands (Data S9 and S10). Promiscuous RFamide receptors activated by multiple RFamide neuropeptides also belong to these expanded neuropeptide receptor families (Data S9), which may explain ligand promiscuity. Taken together, the biochemical interactions identified for all these receptors support co-evolution of neuropeptide-receptor pairs and provide general insights into the evolutionary history and diversification of neuropeptide systems in nematodes.

## Discussion

Neuropeptides and their receptors form the largest and most diverse ligand-receptor signaling network in nervous systems. Deorphanization of neuropeptide receptors has allowed establishing the actions of several well-studied peptidergic systems, such as oxytocin and vasopressin, neuropeptide Y, orexin, and tachykinin, in the regulation of feeding, sleep, arousal, learning and many other physiological processes^3, 7, 13, 14, 17^. The complexity and organization of the neuropeptide signaling network at an organismal scale is, however, not well understood. By systematically identifying neuropeptide ligands for known and predicted peptide GPCRs, we have generated a comprehensive resource of neuropeptide-GPCR couples in *C. elegans*, uncovering an extensive signaling network of specific and complex multichannel interactions. This interaction network is crucial for spurring targeted research into the functions of neuropeptide systems, and lays a foundation for characterizing the brain-wide organization of the neuropeptide signaling network in the *C. elegans* nervous system.

### System-wide deorphanization reveals a rich complexity in peptide-GPCR interactions

In total, we identified 461 peptide-GPCR pairs for bioactive peptides of 60 neuropeptide-encoding genes (31 *flp* and 29 *nlp* genes; Table 1) and expanded the number of deorphanized peptide GPCRs in *C. elegans* from 29 to 60 receptors (not considering isoforms). The repertoire of peptide-GPCR couples is thus much more diverse than the number of neuropeptide and receptor genes. This diversity in part results from complex interactions between neuropeptides and their receptors. For example, many neuropeptides in our dataset activate multiple GPCRs. We also discovered promiscuous receptors and additional ligands for known peptide GPCRs, which interact with multiple peptides from different genes. Furthermore, neuropeptides or GPCR isoforms encoded by a single gene can show different interactions, further increasing the diversity of neuropeptide-receptor couples. These findings suggest that the peptidergic system forms a complex interaction network in which information is not only coded through a binary mechanism of specific ligand-receptor couples, but also through complex combinatorial peptide-receptor interactions. As most peptide-receptor couples that we identified are potent interactions with nanomolar EC_50_ values, and many ligand-receptor pairs in our dataset have been functionally validated (Table 1), such complex peptidergic interactions most likely exist *in vivo* as well.

The complexity of neuropeptide-receptor interactions in *C. elegans* may be explained by the large expansion of specific neuropeptide families, such as the RFamide peptides, during nematode evolution or may represent an adaptation to its compact nervous system^42, 96, 107^. Mounting evidence, however, indicates a similar molecular complexity for peptidergic signaling in other animals, including mammals^40, 65^. Animal genomes typically contain a large number of neuropeptide and receptor genes that, in some cases, have been shown to encode multiple peptides or GPCRs with different signaling properties. A well-studied example is the mammalian pro-opiomelanocortin (POMC) precursor which generates several peptides, such as β-endorphin and adrenocorticotropic hormone (ACTH), that activate different receptors^108^. Similarly, ligand-receptor interactions have been shown to vary between isoforms of mammalian peptide GPCRs^36, 37^. Differential expression of receptor variants may be important for fine-tuning peptidergic modulation^37^, although the functions of receptor isoforms in regulating behavior remain largely unexplored. Moreover, promiscuity of neuropeptide GPCRs has been observed for mammalian neuropeptide systems that did not undergo large expansions^40, 65^. Human neuropeptide FF receptors, for example, are activated by all five groups of mammalian RFamide neuropeptides^109–112^. Receptor promiscuity and peptidergic crosstalk thus appear to be general features of the peptidergic system, especially of RFamide neuropeptides and their receptors. Deorphanization efforts should therefore strive to remain relatively unbiased, including deorphanized receptors and different GPCR isoforms, when mapping neuropeptide-receptor interactions.

### Specific and combinatorial signaling motifs in the peptidergic network

Our dataset reveals three main types of peptidergic interactions that show different levels of complexity: specific, one-to-many and many-to-many interactions. Around 70% of deorphanized receptors interacts with neuropeptides from only a single peptide-encoding gene, generating a large diversity of specific interactions for information coding. Neuropeptide-GPCR couples from 14 peptide and receptor genes show unique interactions, *i.e.* they do not share ligands or receptors with any other identified pair (Figure 2C and 5D). Several of these are orthologous to bilaterian neuropeptide systems, including thyrotropin-releasing hormone, GnRH, calcitonin, elevenin, myoinhibitory peptide, and somatostatin related systems. These are broadly-conserved regulators of key biological processes, such as growth, reproduction, learning and memory across animals, including *C. elegans*^5, 61, 98^. Both neuropeptide and receptor genes thus likely co-evolved under strong selective pressure, which may account for their interaction specificity. By contrast, we found a few neuropeptides that interact with many specific GPCRs, which can diverge signaling from one peptide into multiple channels. It is noteworthy that these peptides (*e.g.* FLP-1, FLP-8 and FLP-14) are among the most abundant and highly conserved RFamide neuropeptides in nematodes^25, 107, 113^. They are essential to the regulation of diverse biological processes, including pathogen and carbon dioxide avoidance, nociception, locomotion, and reproduction, and during evolution may have acquired multiple functions mediated by different receptors^56, 88, 114–117^. Some RFamide neuropeptide systems, however, show a high level of crosstalk by interacting with promiscuous GPCRs. Among these, the promiscuous receptors DMSR-1 and EGL-6 have been implicated in regulating behavioral states, such as those related to sleep, foraging and reproductive behaviors^50, 118–120^. Although more functional studies of peptide-GPCR couples are needed, distinct types of interactions may be specialized for different biological functions. We speculate that neuropeptide-receptor couples with complex or promiscuous interactions may be important for orchestrating global states and cellular activity, for example, by integrating or broadcasting peptide signals. Unique neuropeptide-receptor couples, on the other hand, may have important roles in mediating specific local or long-range signaling.

Combinatorial signaling can also occur between bioactive peptides encoded by the same neuropeptide gene. We show that peptides from a single precursor protein in most cases activate the same receptors. As peptidergic communication is limited by diffusion, one way to amplify signaling is by increasing the concentration of secreted neuropeptide molecules that activate a given GPCR^121^. This is corroborated by our finding that peptides from the same precursor often activate a receptor with similar potency. Some neuropeptide genes, however, encode peptides with distinct receptor interactions that diversify rather than amplify signaling. The peptide-GPCR couples in this resource provide handles to further investigate the functional architecture and processing of neuropeptide precursors.

### Limitations of the study and prospects for future deorphanizations

Although we more than doubled the number of deorphanized peptide GPCRs and recapitulated most of the known peptide-receptor interactions in *C. elegans*, we could not identify ligands for 59% of receptors in our screen. One explanation could be that some peptide GPCRs are not functionally expressed in the heterologous system that we used. The promiscuous Gα_16_ subunit efficiently couples many GPCRs to the phospholipase C pathway but may be less effective in linking G_i_-coupled receptors to the release of intracellular calcium^83, 122^. In addition, GPCR signaling can be mediated by G protein-independent pathways and may require specific co-factors, such as receptor-activity modifying proteins (RAMPs), for some receptors^123–125^. To further complete the neuropeptide-receptor network, different heterologous systems and screening strategies can be combined in the future, for example, using chimeric G proteins and readouts of other key signaling events in the receptor activation cascade^57, 82, 122, 126, 127^. Such multifaceted screening approaches have improved ligand discovery for human peptide GPCRs and may also probe interactions with other receptors besides GPCRs^40, 128, 129^. Alternatively, the remaining orphan receptors may be activated by neuropeptides or other ligands not included in our synthetic library. Our phylogenetic analysis suggests that additional neuropeptides remain to be discovered in the *C. elegans* genome. For example, we identified a large family of orphan nematode GPCRs, including eight *C. elegans* receptors, that may be activated by potentially unknown peptides (cluster 39, Data S9). Likewise, we discovered two orphan receptor families related to the *Drosophila* trissin and CNMamide receptors (clusters 35 and 36, Data S9), suggesting that these neuropeptide systems are conserved in ecdysozoans, but trissin- and CNMamide-related peptides have not been identified in nematodes so far^130, 131^. Recently, peptidomics and comparative genomic studies also identified novel neuropeptides in *C. elegans*^69, 132^. Peptide discovery by these and other approaches, such as machine learning methods^40, 133^, will facilitate future deorphanizations. Finally, the G proteins by which peptide-GPCR pairs signal endogenously remain unknown; uncovering these pathways will be important to better understand the cellular effects of peptide-receptor couples, and may yield further insights into complex interactions, such as promiscuous receptor signaling. With the ligands identified in this resource, G protein coupling of deorphanized GPCRs can be investigated in the future.

### Evolution of neuropeptide systems and other resource applications

Neuropeptide-receptor interactions are one of the most ancient types of signaling pathways that link up cells into complex chemical networks^1, 121^. While synaptically-wired circuits can be anatomically mapped, the brain’s chemoconnectome constituted by numerous neuromodulators, such as neuropeptides, monoamines and their receptors, cannot be resolved solely from anatomical expression, as this extensive ‘wireless’ network relies on chemical messengers that often act remotely from their release site^41, 134, 135^. Therefore, knowledge of ligand-receptor interactions is crucial to delineate which nodes or cells in the network can communicate with each other. This dataset of peptide-GPCR couples lays a foundation for charting the neuropeptide signaling network and, together with single-cell transcriptome resources and the characterized monoamine network^25, 41, 74^, map the chemoconnectome in the *C. elegans* nervous system^75^. Determining the cellular circuits formed by neuropeptide transmission *in vivo* is the next step forward toward understanding the functional organization of this broad signaling network. The development of genetically-encoded sensors for peptide GPCR signaling provide a promising avenue to further map functional peptidergic circuits and validate peptide-receptor interactions *in vivo*^136–138^.

In addition, our dataset allows for more targeted research into neuropeptide functions, guided by receptor interactions, and further clarifies evolution of peptidergic systems. Because biologically-active neuropeptides typically have short peptide sequences and co-evolve with their receptors, identifying orthology between peptidergic systems from distant animals remains a challenge. Annotations of the first invertebrate genome sequences revealed a large number of GPCRs that resembled vertebrate neuropeptide receptors, but relatively few vertebrate-like peptides^139–141^. Initially, this led to the assumption that vertebrate neuropeptide systems are not well conserved in *C. elegans* and other invertebrates. The increasing number of available genome sequences, however, has enhanced our knowledge on the long-range evolution of peptidergic systems^1, 5, 42, 92, 95, 96^, revealing deep conservation of neuropeptide systems across bilaterian animals, which in some cases has been confirmed by biochemical interactions^42, 63, 92^. Our dataset provides further evidence for orthologs of at least 14 bilaterian neuropeptide systems in *C. elegans* (Data S9). In addition, we clarify the diversification of specific neuropeptide receptor families in nematodes. Indeed, many of the receptors we deorphanized belong to expanded GPCR families for which specific ligands could not have been predicted based on available data, as the corresponding neuropeptide families (*e.g.* RFamide, sNPF, and somatostatin-like peptides) have diversified in nematodes as well^42, 104^. The neuropeptide-receptor pairs that we discovered can facilitate future predictions of peptide-GPCR couples in *C. elegans* as well as other species, such as parasitic nematodes, which may speed up the discovery of drug targets. Conservation of neuropeptide-receptor interactions has been shown between *C. elegans* and nematode parasites^142, 143^. Furthermore, peptidergic signaling regulates key aspects of nematode biology, such as reproduction, development, and locomotion^47, 132, 144, 145^. The discovery of neuropeptide-GPCR pairs in parasitic species may thus provide promising leads for the development of novel anthelminthics. Comparisons of neuropeptide-receptor pairs between nematode species, in the future, could also further clarify the diversification and ligand promiscuity of GPCRs.

Taken together, our dataset lays out a genome-wide interaction map of neuropeptide-GPCR couples in *C. elegans* that, along with the synaptic connectome, neuronal expression atlas and other resources in this model system, provides a unique framework for dissecting the structure and functions of the peptidergic signaling network. This lays a foundation for further exploring the mechanisms by which neuropeptides govern physiology and behavior in animals, as well as the regulation of neuropeptide signaling and biosynthesis.

## Acknowledgments

This work was supported by the European Research Council (Grant 340318), the National Institutes of Health (Grant R01 NS110391-01), and the KU Leuven Research Council (Grant C19/19/003). J.D. was a postdoctoral fellow supported by the European Union’s Horizon 2020 research and innovation program (Marie Skłodowska-Curie Grant Agreement No. 703594-DECODE) and the Research Foundation – Flanders (12J6921N). P.E.V. is a fellow of MQ: Transforming Mental Health (Grant MQF17_24). All research from the Department of Psychiatry at the University of Cambridge is made possible by the NIHR Cambridge Biomedical Research Centre and the NIHR East of England Applied Research Centre. The views expressed are those of the author and not necessarily those of the NIHR or the Department of Health and Social Care. For the purpose of open access, the authors have applied a Creative Commons Attribution (CC BY) licence to any Author Accepted Manuscript version arising from this submission.

## Author contributions

I.B. and S.Z. designed experiments. E.V., J.C. and E.B. performed GPCR deorphanization screens. P.E.V. and W.R.S. conducted the bipartite network analysis and O.M. performed the phylogenetic analysis. I.B., S.Z. and J.D. analyzed and interpreted results. I.B. and L.S. wrote the paper with input from all authors.

## STAR METHODS

### LEAD CONTACT AND MATERIALS AVAILABILITY

Further information and requests for resources and reagents should be directed to and will be fulfilled by the Lead Contact, Isabel Beets. The CHO-K1 cell line stably expressing mitochondrial-targeted apo-aequorin and human Gα_16_ is under MTA (Perkin Elmer, ES-000-A24) and cannot be freely distributed.

### EXPERIMENTAL MODEL AND SUBJECT DETAILS

#### Animals

The wild-type *C. elegans* (N2 Bristol) strain was obtained from the *Caenorhabditis* Genetics Center (University of Minnesota). Worms were cultivated at 20°C on nematode growth medium (NGM) plates seeded with *E. coli* OP50 bacteria and used for mRNA extraction and subsequent cDNA synthesis. cDNA prepared from mixed-stage culture plates was used as template for PCR of peptide GPCR coding sequences.

#### Microbe strains

The *Escherichia coli* OP50 strain was used as a food source for *C. elegans*.

#### Cell lines

CHO-K1 cells stably expressing mitochondrial-targeted apo-aequorin and a promiscuous human Gα_16_ protein were used for GPCR deorphanization (Perkin Elmer, ES-000-A24). The Gα_16_ protein couples most agonist-induced GPCRs to a calcium second messenger pathway. This cellular expression system allows quantification of changes in intracellular calcium concentrations as a measure for GPCR activation^83–85^. Cells were cultured in Dulbecco’s Modified Eagle Medium F-12 HAM (DMEM/F-12, Gibco®, ThermoFisher Scientific), supplemented with 10% fetal bovine serum (FBS, inactivated at 65°C; Sigma-Aldrich), 1% penicillin/streptomycin mixture (10.000 units/ml penicillin and 10 mg/ml streptomycin in 0.9% NaCl; Invitrogen) and 250 µg/ml zeocin (Invitrogen). Cells were cultured in stable conditions of 37°C, 5% CO_2_ and high relative humidity in an incubator. Mycoplasma tests were performed using the MycoAlert™ Mycoplasma Detection Kit (Lonza) to verify that cells were free of mycoplasma contamination.

## METHOD DETAILS

### Selection and cloning of peptide GPCR candidates

A list of neuropeptide GPCR candidates (Data S1) was compiled from *in silico* searches for neuropeptide GPCR genes in the *C. elegans* genome^47, 71, 76, 77^. Oligonucleotide primers for amplifying receptor cDNAs (Data S1) were designed based on gene models in Wormbase (http://wormbase.org, version WS240). Only receptors with a seven transmembrane topology, as predicted from the translated cDNA sequence (http://www.cbs.dtu.dk/services/TMHMM), were considered as peptide GPCR candidates. Forward primers included a “CACC” sequence at the 5’ end that introduced a partial Kozak sequence for increased translation efficiency in mammalian cells. Receptor cDNAs were amplified by PCR with Q5 High-Fidelity DNA Polymerase (New England Biolabs) from cDNA of mixed-stage populations of wild-type *C. elegans*, and were cloned in a pcDNA3.1 vector (ThermoFisher Scientific). Plasmids for C16D6.2, T14B1.2^53^, F59D12.2, and C54A12.2^51^ were kind gifts from Dr. Derek Sieburth (University of Southern California, Los Angeles, USA), Dr. Rachel McMullan (The Open University, Milton Keynes, UK) and Dr. David Raizen (University of Pennsylvania, Philadelphia, USA). All plasmids were verified by sequencing of the full receptor cDNA and purified with an EndoFree Plasmid Maxi Kit (Qiagen). cDNA sequences of all GPCR variants cloned in pcDNA3.1 are summarized in Data S2.

### Peptide library composition and synthesis

A library of 344 synthetic peptides was generated for deorphanization of *C. elegans* peptide GPCRs. Prior to the deorphanization screens, we compiled a list of known *C. elegans* neuropeptide precursors of the FLP and NLP families that yield potential peptide GPCR ligands, by assembling peptide sequences from peptidomics studies and *in silico* searches for neuropeptides in the *C. elegans* genome^5, 6, 42, 70, 90, 98, 100, 102, 106, 141, 145, 167–180^. Insulin-related precursors were excluded, because insulin-like peptides typically activate receptor tyrosine kinases^79, 181^. We assembled 94 FLP and NLP precursors and searched each of them for peptide sequences identified by peptidomics or flanked by mono- or dibasic cleavage sites for proprotein convertases (RK, RR, KR, KK or [RK]-X_2/4/6/8_-[RK])^102^. This search yielded 344 mature peptide sequences. Replicates of the identified peptides were synthesized by GL Biochem Ltd. and ThermoFisher Scientific, and were chemically modified to match post-translational modifications (e.g. disulfide bridge formation, C-terminal amidation or N-terminal pyroglutamation) that are commonly found in mature neuropeptides^69^. Sequences of all synthetic peptides in the library are listed in Data S3.

### Transfection of CHO-K1 cells

GPCRs were heterologously expressed in CHO-K1 cells (Perkin Elmer, ES-000-A24), stably expressing mitochondrially tagged apo-aequorin and the promiscuous human Gα_16_ subunit, by transient transfection. Transfections with GPCR/pcDNA3.1 plasmids were performed when cells reached 70-80% confluency using Lipofectamine® LTX with Plus™ Reagent (Invitrogen) in serum-free medium. Transfected cells were supplemented with complete growth medium the day after transfection and were transferred to an incubator at 28°C and with 5% CO_2_ and high relative humidity conditions, 16 hours prior to cell harvest for the aequorin-based GPCR activation assay. The aequorin receptor activation assay was performed two days after transfection.

### Aequorin-based GPCR activation assay for library screening

We used a luminescence-based calcium mobilization to monitor GPCR activation and screen for peptide-GPCR couples in the synthetic *C. elegans* peptide library. Two days after transfection, CHO cells transfected with a receptor of interest were detached and collected in phosphate buffered saline (PBS) containing 0.2% EDTA. Viable cells were quantified using a NucleoCounter NC-100 (Chemometec, Allerod, Denmark). Cells were pelleted by 4 min centrifugation at 800 rpm at room temperature and resuspended in DMEM/BSA (DMEM/F-12 without phenol red, with L-glutamine and 10 mM HEPES, 0.1% bovine serum albumin; Gibco®, Thermo Fisher Scientific) at a concentration of 5×10^6^ cells/ml. Coelenterazine H (Invitrogen) was added at a final concentration of 5 µM, and cells were gently shaken at room temperature for 4 h in the dark, allowing the aequorin holoenzyme to be reconstituted. After a 10-fold dilution in DMEM/BSA, cells were incubated for 30 min before starting the assay. Library screening was performed on a FLIPR^®^ Tetra High-Throughput Cellular Screening System (Molecular Devices). Using this system, cells were transferred to 96-well plates (25,000 cells/well) containing 50 µl of synthetic peptide, dissolved in DMEM/BSA, and calcium responses were simultaneously monitored for 36 seconds. Peptides were individually tested in black 96-well plates with clear bottom (Greiner Bio One) at a final concentration of 10 µM. Wells containing 50 µl DMEM/BSA were used as a negative control. ATP (1 µM in DMEM/BSA), which activates an endogenous receptor in CHO cells, was used as a positive control. After measuring peptide-evoked responses, 50 µl of Triton X-100 (0.2% in DMEM/BSA) was added to lyse the cells and obtain a measure of the maximum calcium response.

### Dose-response assays

All peptide hits identified in library screens were further tested for validation in dose-response assays. Crude peptides, used for library screening, were first purified by reversed phase HPLC on a Symmetry-C18 column (4.6 x 250 mm HPLC cartridge with pore size of 5 µM) to obtain peptide stocks of high purity. Peptide masses of purified peptides were verified by MALDI-TOF/TOF mass spectrometry on a rapifleX® MALDI Tissuetyper® (Bruker Daltonic). Purified peptides were tested in concentration series using the aequorin-based GPCR activation assay on a MicroBeta2 LumiJET (Perkin-Elmer) luminometer. Positive and negative controls were similar to those used in library screens. Calcium responses were measured for 30 seconds after adding cells to the compound plate. After the addition of Triton X-100 (0.2% in DMEM/BSA), a maximum response per well was measured for 30s, which was used for normalization of peptide-evoked calcium responses. Dose-response measurements were performed in triplicate on at least two independent days.

### Bipartite network analysis

Bipartite graphs were constructed for all FLP and NLP neuropeptide-GPCR couples with nanomolar EC_50_ values (< 1 μM) identified in this resource (Data S6). Monopartite network projections were generated and analyzed using custom scripts in MATLAB; scripts are available upon request, MATLAB version 9.10.0.1602886 (R2021a).

Modularity: Many complex networks have a modular structure, whereby they contain subsets of nodes - called modules - that are more densely interconnected with each other than with the rest of the network. Various algorithms have been designed to find such modules in real-world networks. Here, modules were defined using a popular consensus clustering approach^182^, with the Louvain community detection algorithm as implemented in the Brain Connectivity Toolbox^183, 184^. The Fruchterman-Reingold force-directed layout algorithm was used to layout graphs in 2D space for vizualisation^185^.

### Peptide sequence alignments

Pairwise peptide sequence alignment scores in Figure S9 were generated using the *stringDist* method from the R/Bioconducor package Biostrings. The BLOSUM50 substitution matrix was used with gap opening and gap extension penalties set to 13 and 2, respectively.

### Phylogenetic analysis of nematode peptide GPCRs

We first drew a list of 150 *C. elegans* sequences predicted to be peptide GPCRs (Data S8B). We then added to this list the predicted peptide GPCRs from non-nematode bilaterian genomes of *Homo sapiens* (Vertebrata), *Takifugu rubripes* (Vertebrata), *Branchiostoma floridae* (Cephalochordata), *Saccoglossus kowalevskii* (Hemichordata), *Strongylocentrotus purpuratus* (Echinodermata), *Capitella teleta* (Annelida), *Lottia gigantea* (Mollusca), *Dapnia pulex* (Arthropoda) and *Drosophila melanogaster* (Arthropoda) and those from the nematode species *Strongyloides ratti*, *Trichinella spiralis*, *Brugia malayi*, *Pristionchus pacificus, Loa loa* and *Onchocercus volvulus* downloaded from the Ensembl *metazoan* database, using the strategy described in Mirabeau and Joly, 2013. We complemented these sequences with the predicted peptide GPCRs from the complete set of nematode ESTs available from NCBI (https://www.ncbi.nlm.nih.gov/nucest). All nematode species used in the construction of the phylogenetic tree are listed in Data S8A.

We retrieved 150 secretin-like and 1963 rhodopsin-like GPCRs for this analysis (Data S8C-D) and annotated each bilaterian sequence that was included in Mirabeau and Joly, 2013 for its membership in a conserved bilaterian group (for example, the *Capitella* sequence jgi|Capca1|219484|estExt_fgenesh1_pg.C_20057 was annotated as jgi|Capca1|219484|estExt_fgenesh1_pg.C_20057_Calc/DH31). We then constructed two separate phylogenetic trees were constructed with the SATé-II maximum-likelihood algorithm^186, 187^, with default settings (Aligner:MAFFT^188^, Merger:MUSCLE^189^, Tree Estimator:FASTTREE^187^, substitution model:GTR+G20, Maximum size subproblem:50%, Decomposition:Centroid, Stop after Last Improvement). Branch support values were generated using the implementation of the Shimodaira-Hasegawa test of FASTTREE, displayed as “label” in supplemental tree data.

### QUANTIFICATION AND STATISTICAL ANALYSIS

#### Selection of peptide hits for validation in dose-response assays

Peptide-GPCR interactions were initially tested at high peptide concentrations (10 μM) in luminescence-based GPCR activation screens with a synthetic peptide library. From this data, we selected putative receptor ligands by manually inspecting relative light unit (RLU) plots of calcium responses and by ranking peptides based on standardized Z scores. We integrated peptide-evoked calcium responses over time and normalized the signals per assay plate and subsequently per receptor to compute an activation value for each peptide-GPCR pair. Specifically, we performed Z-score normalization on the integrated RLU signal using the mean and variance of the on-plate negative controls to obtain an activation value for each peptide-GPCR pair. Next, we standardized the Z-scores for each GPCR by dividing by its maximal activation Z-score, and log2-transformed this fraction to rank individual peptide-GPCR couples. Based on the scores obtained for known peptide-GPCR interactions, we considered all peptide-GPCR couples with a strong calcium response over background (Z-score >20), reaching at least 40% of that receptor’s maximum activation (log2-standardized Z > −1/2), as potential hits in the peptide library. When the top hit (log2-standardized Z = 0) could not be validated in dose-response tests, Z-scores for that receptor were restandardized to the next top hit in the library screen and dose-response assays were performed for all peptides matching the selection criteria above. The NLP-2-1 synthetic peptide was excluded from the standardization procedure (log2-standardized Z was set to NA), as it produced a signal (Z-score ≥ 5) in 136 out of 176 assays and was therefore considered to be a likely false positive.

When a peptide dose-dependently activated its receptor, all peptides derived from the same neuropeptide precursor were systematically tested in dose-response assays. An exception was made for GPCRs with peptide hits originating from more than 10 different FLP precursor proteins, i.e. DMSR-1, DMSR-7 and FRPR-8. For these promiscuous receptors, at least one FLP peptide from each FLP precursor was tested for dose-dependent receptor activation.

#### Dose-response curves and EC_50_ values

Calcium responses for dose-response curves were measured in triplicate on at least two independent days. For each peptide concentration and replicate, calcium responses were first normalized to the maximum calcium response (sum of peptide- and Triton-evoked response) and subtracted with the negative control (BSA) value. Next, a relative calcium response (%) compared to the maximum peptide-evoked response (100% activation) was calculated. Dose-response data were fitted in function of log[peptide] using GraphPad Prism 5 or 7. EC_50_ values were calculated from dose-response curves by fitting a 3- or 4-parameter dose-response curve. In empty vector control experiments, calcium responses of peptide and control samples were compared by two-way ANOVA with Tukey post-hoc test.

## Supplemental information legends

**Figure S1, related to Figure 1.**
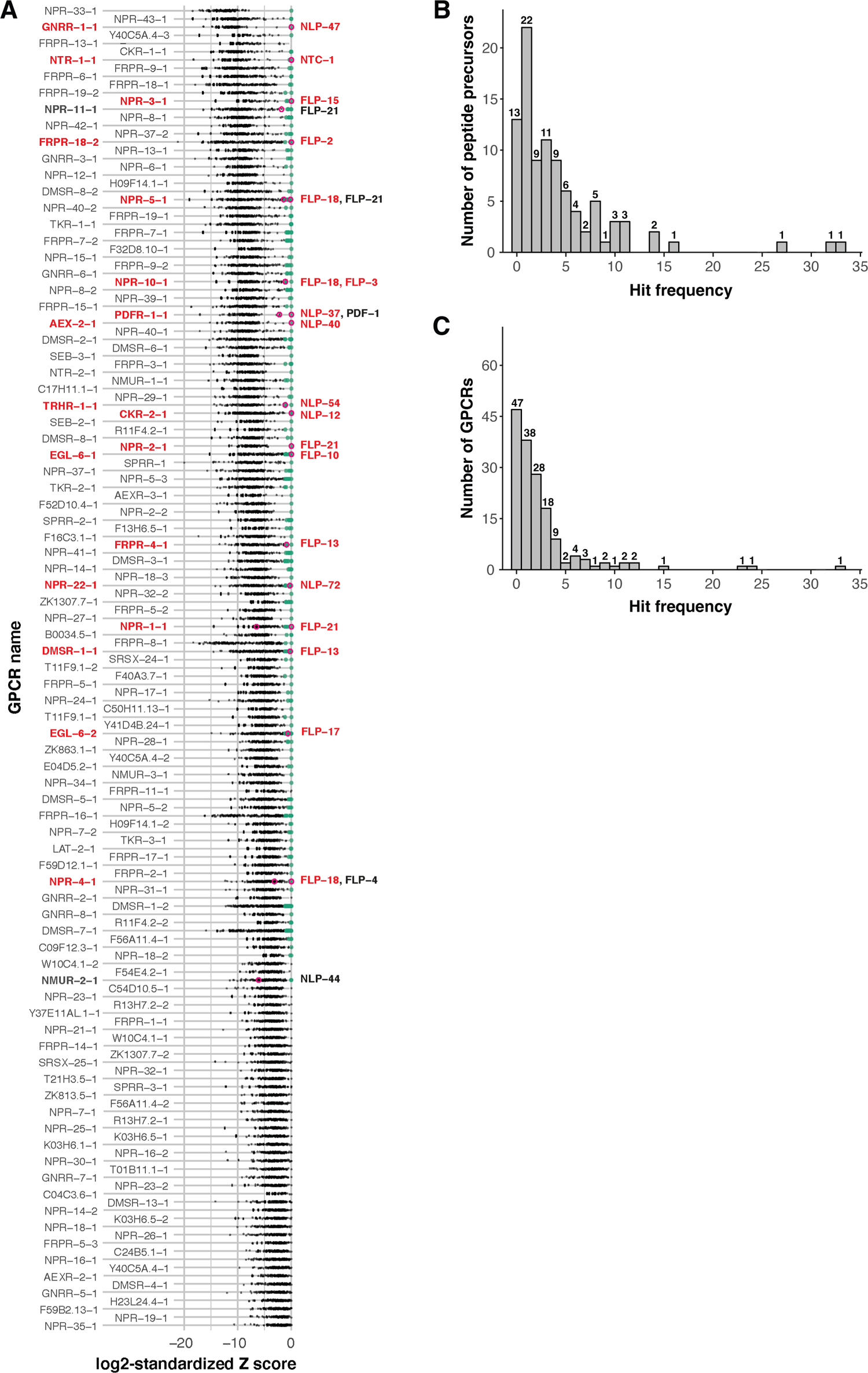
Peptide hits in reverse pharmacology screens of *C. elegans* peptide GPCR candidates. (A) Log2-standardized Z-scores for peptide-GPCR interactions and identified hits in library screens. Receptors are ranked, from low to high, according to their median log2-standardized Z-score. Peptide-GPCR couples with nanomolar EC_50_ values (< 1 μM) known prior to this study are circled in pink (see Table 1 for references; GPCR isoform with highest log2-standardized Z-score is labeled). Green dots indicate 416 peptide-GPCR couples that show a significant calcium response of at least 40% of the receptor’s maximal activation value (Z-score > 20, log2-standardized Z-score > −1.2), which are considered as peptide hits. In total, 79% (19/24) of the reported interactions are identified as peptide hits in the screens (labeled in red). See also Data S4 for a list of Z-scores. (B) Frequency distribution for neuropeptide precursors according to the number of peptide hits in reverse pharmacology screens. (C) Frequency distribution for peptide GPCRs according to the number of peptide hits in reverse pharmacology screens.

**Figure S2, related to Figures 1 and 2.**
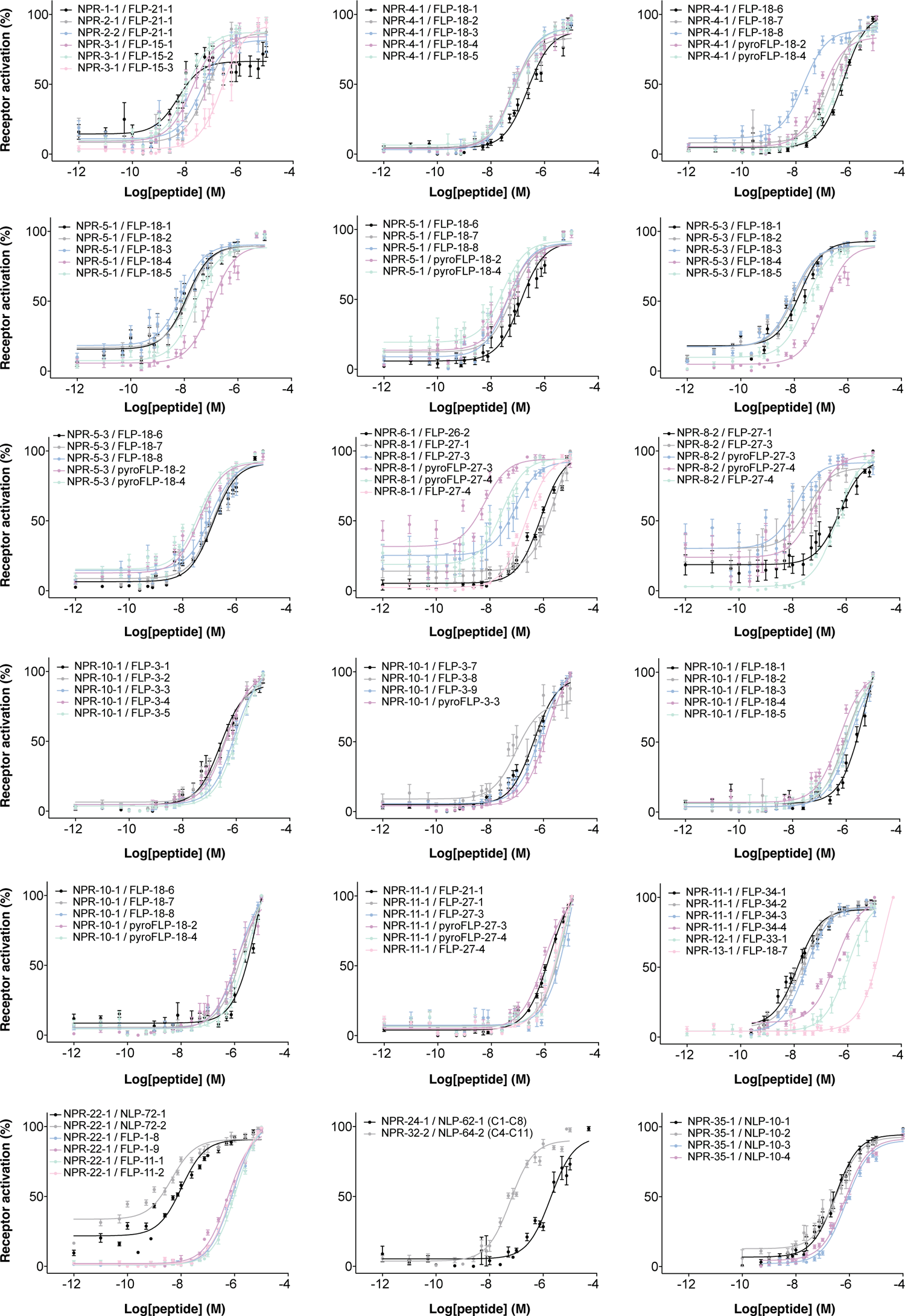
Dose-response curves for neuropeptide ligands activating *C. elegans* NPRs Dose-response curves for *C. elegans* NPRs co-expressed in CHO cells with the promiscuous Gα_16_ subunit and the luminescent calcium indicator aequorin. Calcium responses are shown relative (%) to the highest value (100% activation) after normalization to the total calcium response. Error bars represent SEM (n ≥ 6). Interactions of FLP-3/NPR-10, FLP-34/NPR-11 and NLP-72/NPR-22 were reported and validated in *in vivo* studies^72, 89, 149^. See also Data S6 for corresponding EC_50_ values.

**Figure S3, related to Figures 1 and 2.**
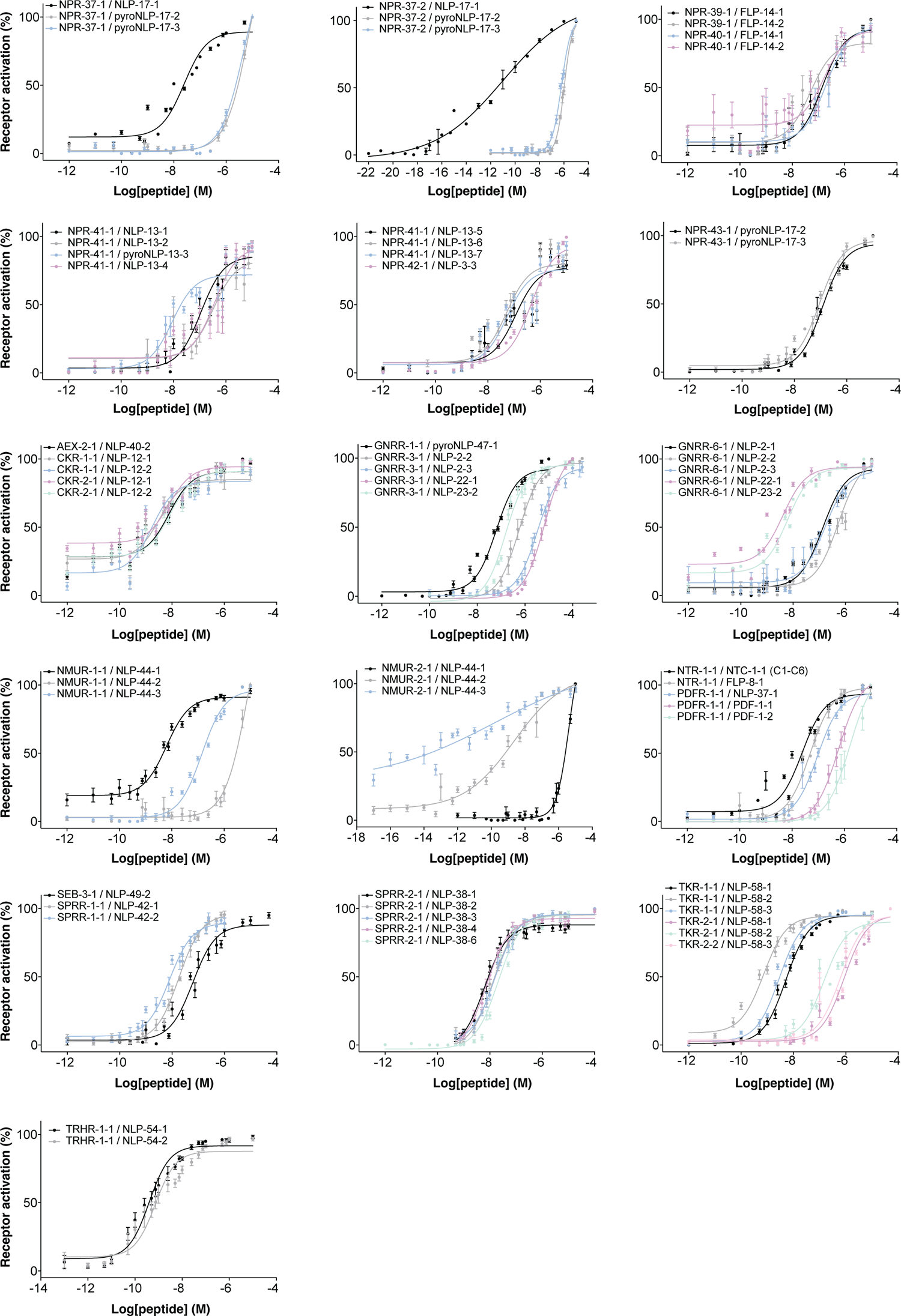
Dose-response curves for neuropeptide ligands of NPRs and orthologs of ancestral bilaterian peptide GPCRs Dose-response curves for *C. elegans* peptide GPCRs co-expressed in CHO cells with the promiscuous Gα_16_ subunit and the luminescent calcium indicator aequorin. Calcium responses are shown relative (%) to the highest value (100% activation) after normalization to the total calcium response. Error bars represent SEM (n ≥ 6). Interactions of NLP-49/SEB-3, NLP-12/CKR-1, NLP-44/NMUR-1, GNRR-3 and GNRR-6 were reported and validated in *in vivo* studies^73, 86, 87, 91^. See also Data S6 for corresponding EC_50_ values.

**Figure S4, related to Figures 1 and 2.**
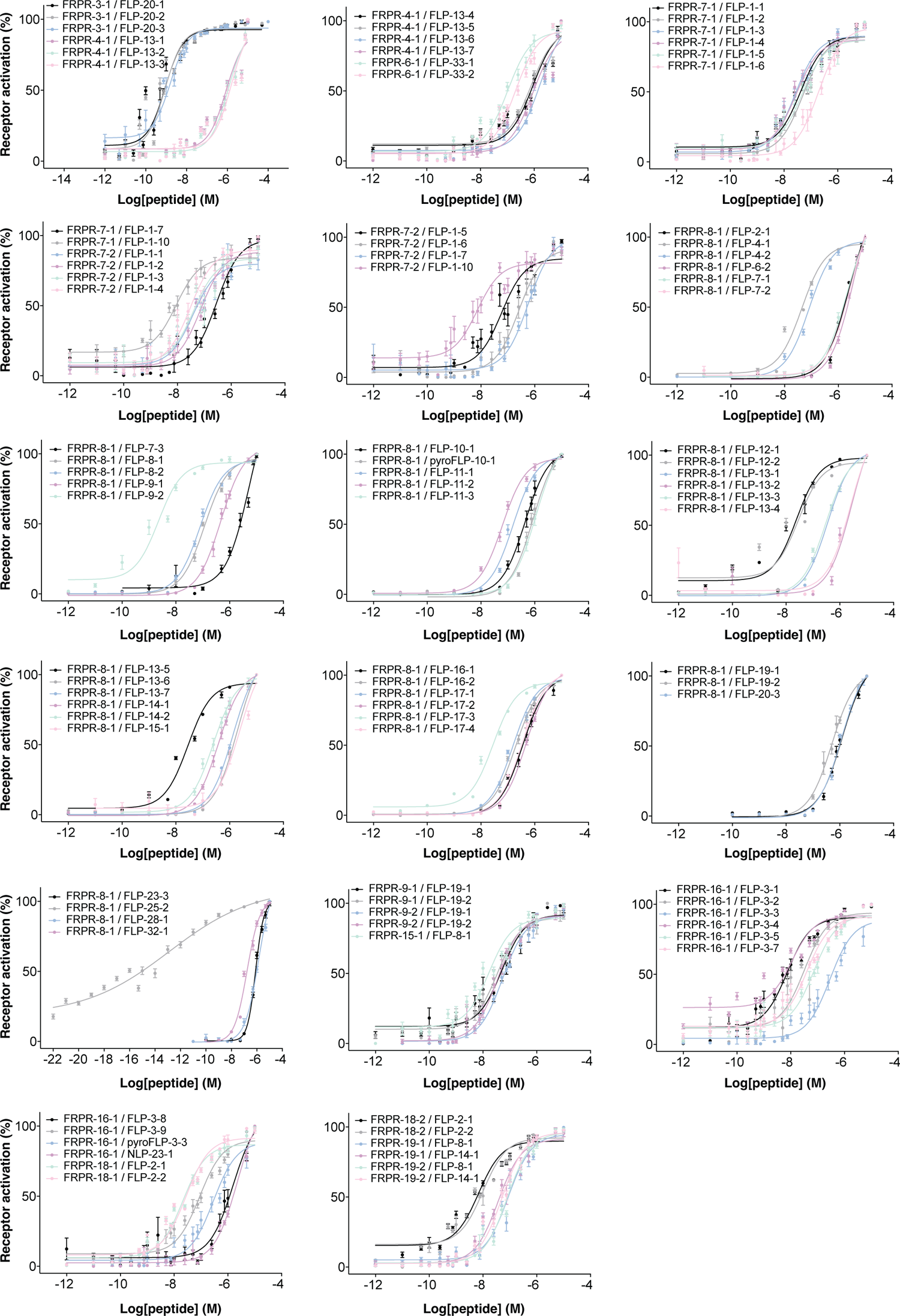
Dose-response curves for neuropeptide ligands of ***C.*** *elegans* FRPRs Dose-response curves for *C. elegans* FRPRs co-expressed in CHO cells with the promiscuous Gα_16_ subunit and the luminescent calcium indicator aequorin. Calcium responses are shown relative (%) to the highest value (100% activation) after normalization to the total calcium response. Error bars represent SEM (n ≥ 6). Interactions of FLP-3/FRPR-16 and FRPR-19 were reported and validated in *in vivo* studies^88, 149^. See also Data S6 for corresponding EC_50_ values.

**Figure S5, related to Figures 1 and 2.**
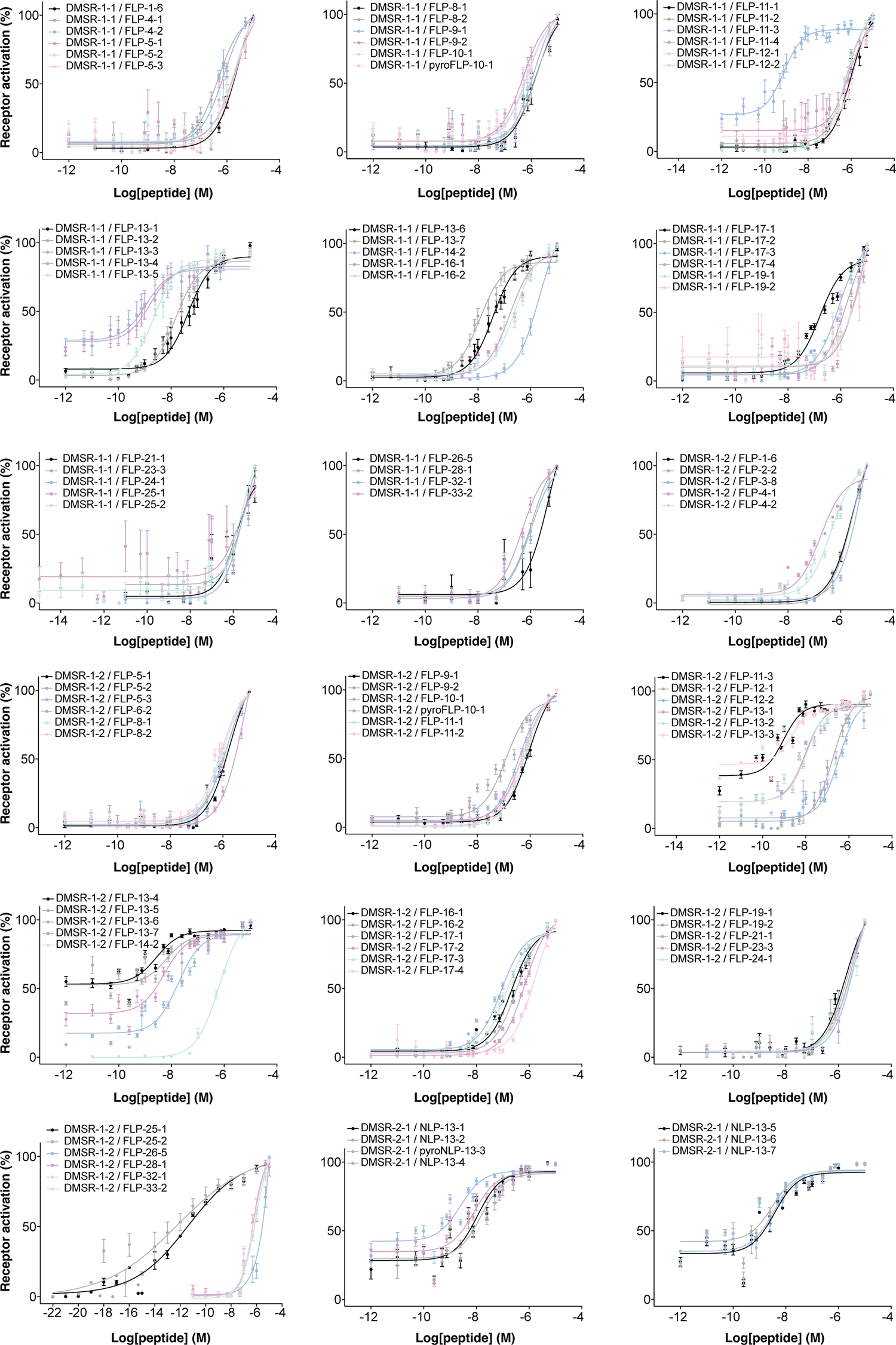
Dose-response curves for neuropeptide ligands of DMSR-1 and DMSR-2 Dose-response curves for DMSR-1 and DMSR-2 receptor isoforms co-expressed in CHO cells with the promiscuous Gα_16_ subunit and the luminescent calcium indicator aequorin. Calcium responses are shown relative (%) to the highest value (100% activation) after normalization to the total calcium response. Error bars represent SEM (n ≥ 6). See also Data S6 for corresponding EC_50_ values.

**Figure S6, related to Figures 1 and 2.**
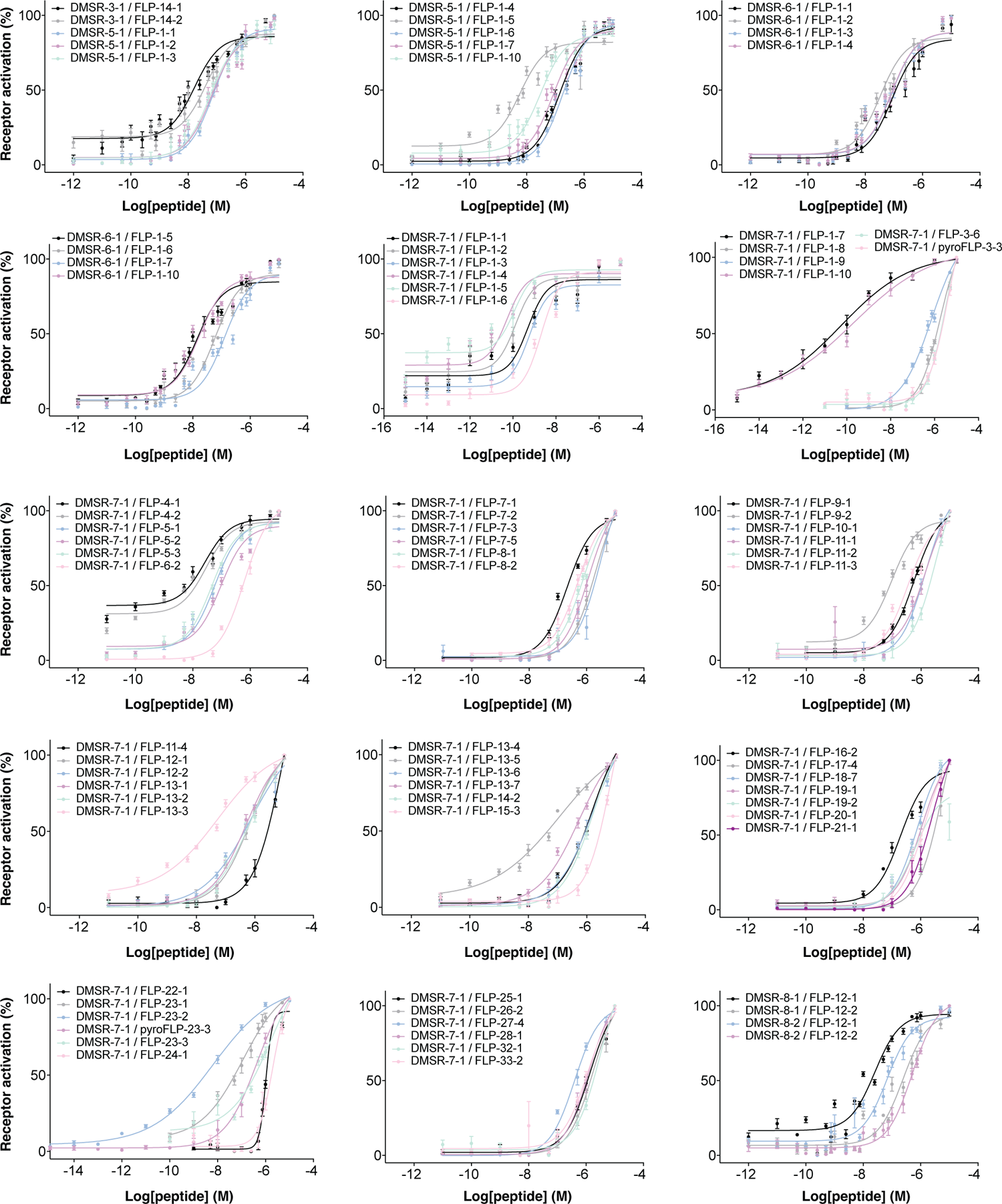
Dose-response curves for neuropeptide ligands of DMSRs Dose-response curves for *C. elegans* DMSRs co-expressed in CHO cells with the promiscuous Gα_16_ subunit and the luminescent calcium indicator aequorin. Calcium responses are shown relative (%) to the highest value (100% activation) after normalization to the total calcium response. Error bars represent SEM (n ≥ 6). Interactions of FLP-1/DMSR-7 were reported and validated in *in vivo* studies^117^. See also Data S6 for corresponding EC_50_ values.

**Figure S7, related to Figures 1 and 2.**
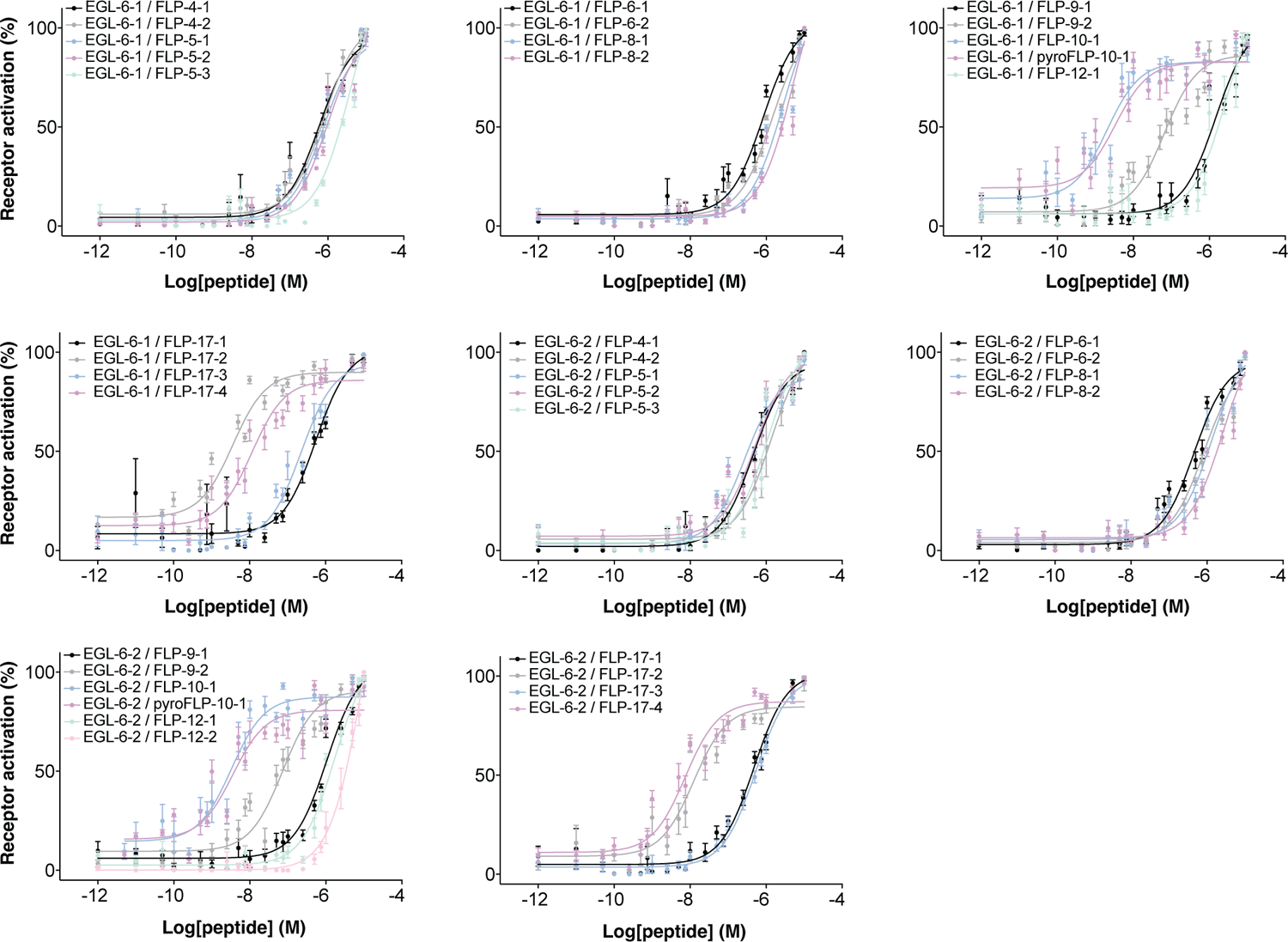
Dose response curves for neuropeptide ligands of EGL-6 Dose-response curves for EGL-6 receptor isoforms co-expressed in CHO cells with the promiscuous Gα_16_ subunit and the luminescent calcium indicator aequorin. Calcium responses are shown relative (%) to the highest value (100% activation) after normalization to the total calcium response. Error bars represent SEM (n ≥ 6). See also Data S6 for corresponding EC_50_ values.

**Figure S8, related to Figure 4.**
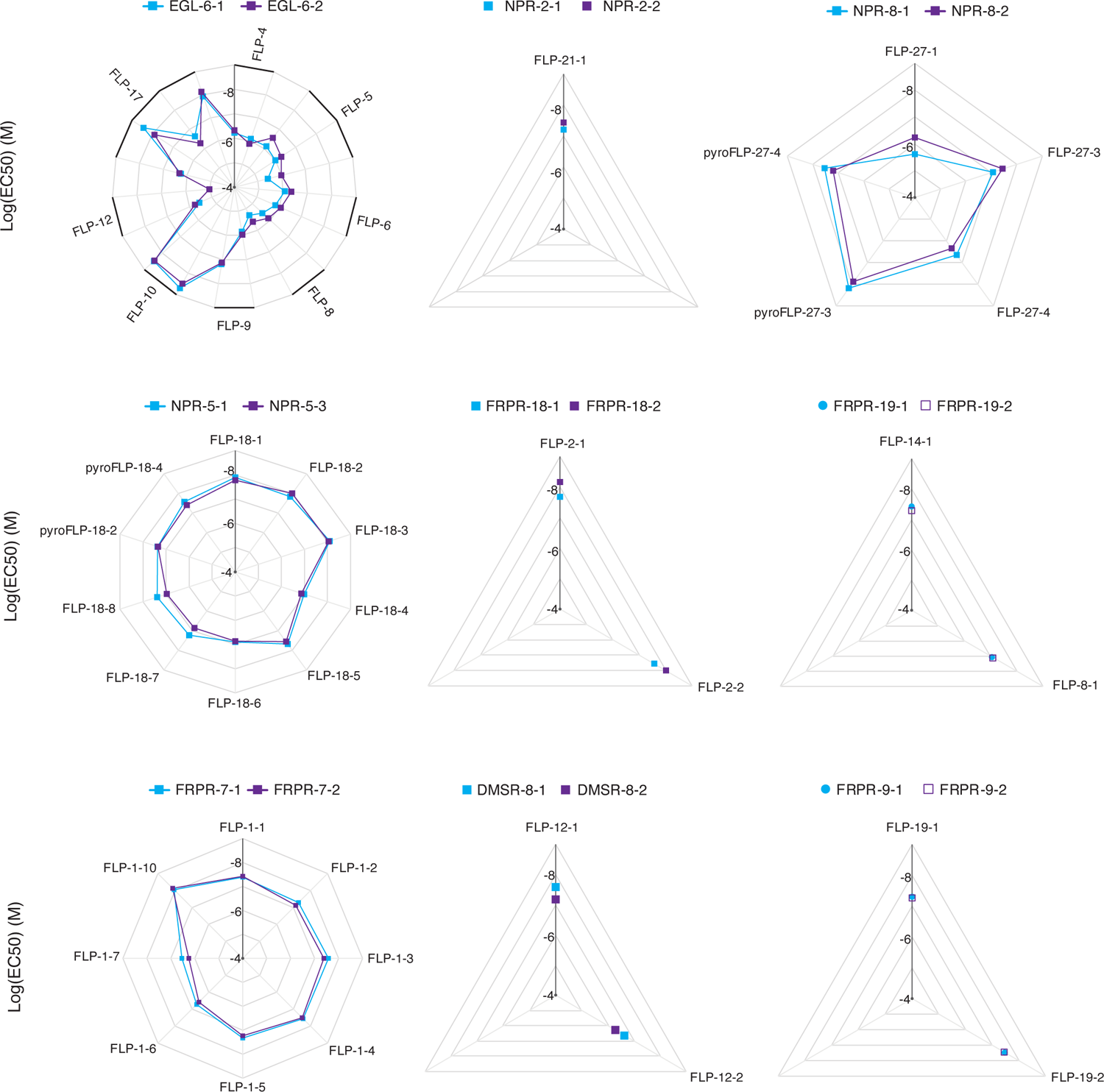
Neuropeptide-GPCR interactions for different receptor isoforms Radar plots depicting log(EC_50_) values for each identified neuropeptide ligand of GPCR isoforms (see Data S6 for a list of EC_50_ values). Different receptor isoforms show similar ligand-receptor interactions for most peptide GPCRs.

**Figure S9, related to Figure 4.**
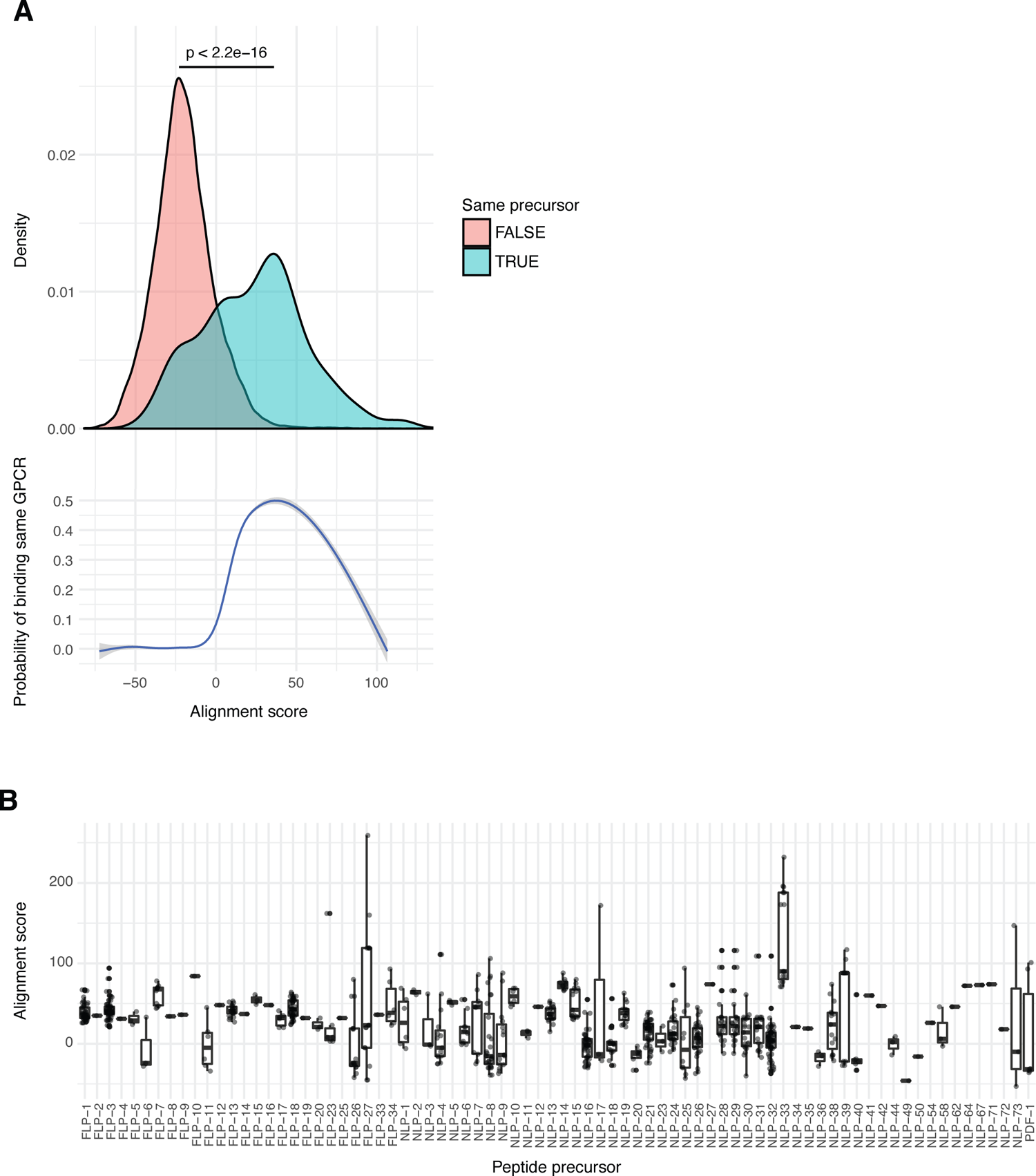
Neuropeptides derived from the same precursor protein share sequence similarity and activate similar receptors (A) Neuropeptide sequences encoded in the same precursor protein show significantly higher pairwise alignment scores. In addition, similar peptides are more likely to bind the same GPCR than dissimilar ones (Pearson’s Chi-squared test, χ^2^ = 1604, df = 1, p-value < 2.2e^-16^). (B) Boxplots of pairwise BLOSUM50 alignment scores for neuropeptide sequences encoded in the same precursor protein. Boxplot height indicates neuropeptide precursors encoding at least one more divergent neuropeptide.

**Figure S10, related to Figure 4.**
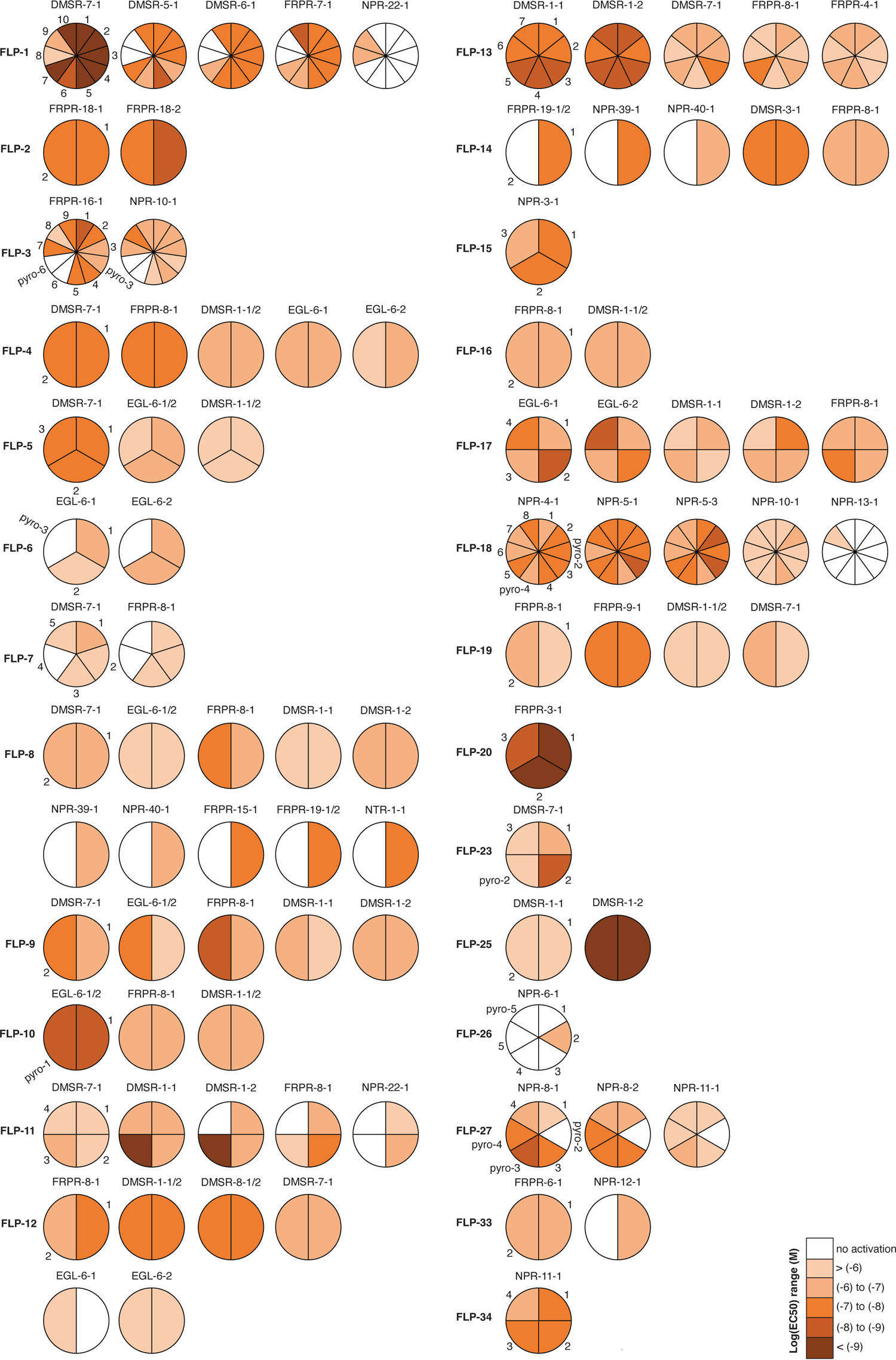
RFamide neuropeptides encoded by the same precursor protein can activate distinct peptide GPCRs Pie charts for neuropeptide-GPCR couples involving FLP neuropeptides from the same precursor protein, colour-coded according to the log(EC_50_) range of ligand-receptor interactions. In several cases, FLP peptides encoded by the same neuropeptide precursor differ in their potency to activate specific GPCRs. For example, the RYamides FLP-1-8 and FLP-1-9 activate DMSR-7 and NPR-22 receptors, but not DMSR-5, DMSR-6 and FRPR-7 that are strongly activated by RFamide FLP-1 peptides (FLP-1-1 to FLP-1-7, and FLP-1-10). Similarly, FLP-11-3 is the most potent ligand of DMSR-1 and DMSR-7, whereas FRPR-8 and NPR-22 are more strongly activated by FLP-11-2.

**Figure S11, related to Figure 4.**
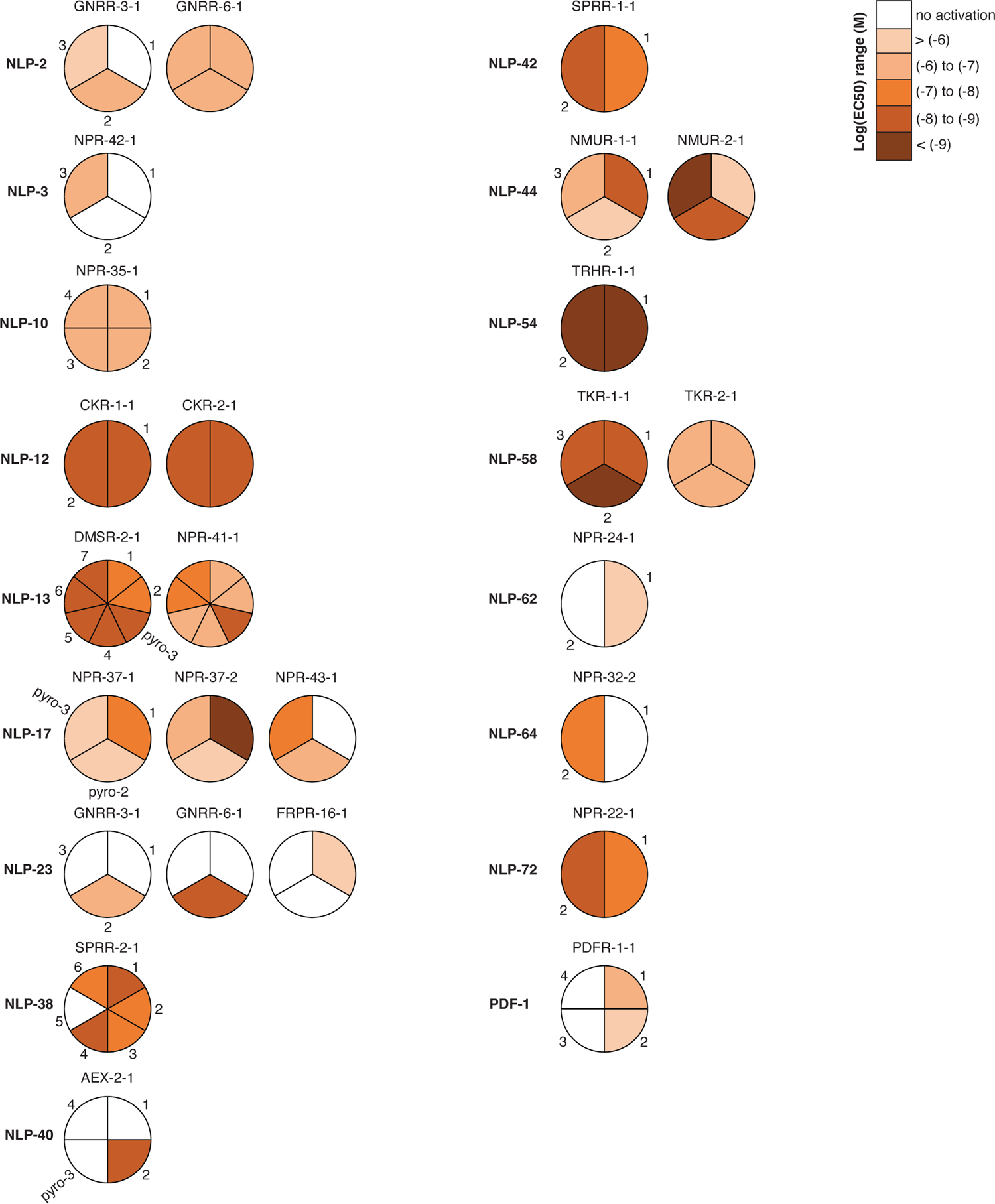
NLP peptides derived from the same neuropeptide precursor show different neuropeptide-GPCR interactions Pie charts for neuropeptide-GPCR couples involving NLP neuropeptides derived from the same precursor protein, colour-coded according to the log(EC_50_) range of ligand-receptor interactions. Peptides derived from the same NLP precursor do not always activate similar GPCRs. For example, NLP-17-1 is the most potent ligand of NPR-37, while NPR-43 is strongly activated by pyroglutamated NLP-17 peptides but not NLP-17-1. NLP-23-2 peptides activate GNRRs but not FRPR-16, which is activated by NLP-23-1. Similarly, NLP-44-1 is the most potent ligand of the NMUR-1 receptor, whereas NMUR-2 is more strongly activated by NLP-44-2 and NLP-44-3 peptides.

---

**Data S1, related to** **Figure 1. List of *C. elegans* peptide GPCR candidates and GPCR/pcDNA3.1 plasmids used in reverse pharmacology screens** List of peptide GPCR genes predicted in the *C. elegans* genome^47, 71, 76, 77^, with primers used for cDNA cloning. GPCR/pcDNA3.1 expression plasmids are indicated by a unique plasmid ID (pR) number. See also Data S2 for cloned receptor cDNA sequences. Gene names and isoforms according to Wormbase version WS240 (https://wormbase.org). For most receptors, translated cDNA sequences matched the predicted protein sequences on Wormbase. We also found 30 additional isoforms that showed a typical seven-transmembrane topology.

**Data S2, related to** **Figure 1. List of receptor cDNA sequences cloned in GPCR/pcDNA3.1 plasmids** Overview of cDNA sequences cloned in receptor/pcDNA3.1 (pR) plasmids for heterologous expression. Gene names of cloned receptors for each pR plasmid are listed in Data S1.

**Data S3, related to** **Figure 1. Composition of synthetic *C. elegans* neuropeptide library** Names and sequences, including modifications, of all FLP and NLP peptides in the synthetic peptide library used for GPCR deorphanization. Cte, C-terminal; Nte, N-terminal.

**Data S4, related to** **Figure 1. Peptide hits and Z-scores for neuropeptides in GPCR deorphanization screens** List of luminescence intensities, Z-scores, and log2-standardized Z-scores for individual neuropeptide-GPCR pairs in reverse pharmacology screens. In addition, all peptide hits tested in dose-response assays are indicated. Peptide hits were selected based on manual inspection of calcium response traces and by calculating Z-scores. Peptides were considered hits when their Z-score and log2-standardized Z-score were greater than or equal to 20 and −1.2, respectively. Note that these scores were restandardized when the top hit (log2-standardized Z-score = 0) could not be validated in dose-response assays.

**Data S5, related to** **Figure 1. Neuropeptide ligands of *C. elegans* peptide GPCRs do not elicit calcium responses in CHO cells transfected with an empty pcDNA3.1 plasmid** Calcium responses of CHO cells transfected with an empty pcDNA3.1 plasmid in response to ATP, BSA or synthetic peptides. ATP activates an endogenous receptor in CHO cells and is used as positive control. BSA medium without synthetic peptide is used as negative control. ATP elicits a significant calcium response compared to the negative BSA control, whereas synthetic peptides do not. Two-way-ANOVA with Tukey test.

**Data S6, related to Figures 1-4. List of EC_50_ values for identified neuropeptide-GPCR couples** Overview of EC_50_ values and 95% confidence intervals for peptide-GPCR couples identified in GPCR deorphanization screens. See also Data S2 and S3 for GPCR and peptide sequences.

**Data S7, related to** **Figure 2. List of predicted neuropeptides activating peptide GPCRs** Predicted neuropeptides, not yet biochemically isolated^69^, activate peptide GPCR candidates in reverse pharmacology screens, suggesting a biological role for these peptides.

**Data S8, related to** **Figure 5. Overview of nematode GPCR sequences used in phylogenetic analyses** (A) Nematode species included in phylogenetic analyses of rhodopsin- and secretin-type peptide GPCRs. (B) *C. elegans* peptide GPCR sequences used for phylogenetic tree construction. (C) Receptor sequences used in phylogenetic tree analysis of rhodopsin-type peptide GPCRs. (D) Receptor sequences used in phylogenetic tree analysis of secretin-type peptide GPCRs.

**Data S9, related to** **Figure 5. Evolutionary relationships of *C. elegans* neuropeptide-receptor systems** Names of characterized neuropeptide systems are assigned according to the classification by Mirabeau and Joly (2013) and Elphick et al. (2018), and an evolution-based nomenclature for *C. elegans* representatives of bilaterian neuropeptide systems is introduced. Bilaterian families with nematode representatives are indicated in blue; those without nematode representatives in red. Families containing only protostomian sequences are coloured yellow, and nematode-specific groups are coloured green. All *C. elegans* peptide GPCRs with their known and newly identified neuropeptide ligands are included (see Table 1 for all individual interactions). Asterisks mark receptors with additional ligands besides orthologs of bilaterian neuropeptide families (summarized in Table 1).

**Data S10, related to** **Figure 5. Maximum likelihood tree of bilaterian rhodopsin peptide GPCRs** Node labels indicate support values. Subtrees are colour-coded as in Figure 5A. Non-typical peptide receptors (angiotensin, bradykinin, and chemokine receptors) were used as an outgroup.

**Data S11, related to** **Figure 5. Maximum likelihood tree of bilaterian secretin peptide GPCRs** Node labels indicate support values. Subtrees are colour-coded as in Figure 5B. Adhesion and cadherin receptors were used as an outgroup.

