## Supplementary material for "System-wide mapping of neuropeptide-GPCR interactions in *C. elegans*": Data S2

>pR1

ATGAACTTTTCGGCCACCGATTCGATATTGGCATCAACGATAACAACGGTGATTGGTGGAGCTGGAGTTTTGGCAGAAGC

AGGCGAAGCTGAACTATCTGGTGATGATGATTTTTATGAGCTGACTCCTGTAGAATTGATAATATGGTGCATGCTGTATG

CAATTATAGCCTTCATGGCAGTTGTTGGAAATCTTCTGGTTCTCTACATAACACTGTTCAGATTAAGAGTCCGTTCCATC

ACAACCTACTTCATTCTGAACCTCGGATTTGCTGACCTCTTCACTGGTATTTTTGCGATTCCCTTCAAGTTTCAGGCTGC

TCTTTTTCAAGAATGGTTCCTGCCGCGATCACTCTGCCGGATAGTTCCATACGTGGAAACAGTTGCTCTGACAGTTTCAG

TCTTCACACTTGTGACGTCAGCAGTTCATGAATTCCGTACAATGTTCTTCTCGAAATGCTCACAAATGAGCCCAAGATCT

GCAAAACGATGTGTACTTTTGATATGGATAATGGCGGTTCTTGTGTCTCTACCACATGGATTGTTCCATAATACATACGA

ATTTCCAGATGACAATAATACTTCAATTGTACAGTGTCTCCCAGTATATCCTGATGCTGGTTGGTGGAAAACATACAATG

TCTACCTTGTCATAATCCAATATTTTGTTCCAATGATTATTCTTGACACTGCGTACACAATGATTGCTGTTAAAATATGG

TCATTGAGTCAGTCAAGAGTTGAACTTGATGAAACAAAAATGGCAACCCAGAAGATATCAGTGGTATCAATGGTTTCACC

AAACACTCAATTATCGCAGCTTATGCGTACTCTCATCATTGTCGTTGCCTGTTTCTCATTGTGTTGGTTTCCATTGGAGA

CGTATCTACTTTTGAATGAATTGAAACCGGAAATTAATGGATGGAAATACATCAATTTGGTGTTCTTCTTTTCACATTGG

CTGGCGATGAGCAATTCTTGTCTTAATCCAATTATTTATGGACTTTACAATACAAAATACAACGAGGAATATCGTCGTTT

GTTTCGCCAAATTGGATGCATTTGGCAACGGCAGAAAAGTTTGGACGATTCGATGAAACCGGAGCGTCGTTGGAATTCTT

CAAATGATTGTCAAGATCAACAGGAAATTGATCAAATTGTTGATATTCCACCAGTTATTTCTACAAATAATCTTTCTCCC

TGA

>pR2

ATGTGCGAAGAAGATGGTCCGAAATCCACAATCTCGTTTGCTATAGTGATAGCTGGATCTCTTATTTCAATTATTTCAAT

TTCGAACAATGTTCTTTTATTTATATCACTAATACGGAATAATAGATGCTTCAAATGCTATTTTCATTTCATTCTTGCAC

TTTGCTTTTTCGACATTATCATATCTGTATGCTATATGCCTGTTATACTGGTAGATTCTTTAAAAGATTGGACTAAATGG

ATTGAACTCGCCAGAGCCTGGTGGCCATTTTTTGTGTATGGTCTGGCAATGACGCATGTCTGCATGACGACTGCATGTTA

TATTCTGATAGCCGTAGCGTATGAGAGATACTTAATAACTGTCAGAAGTTACATGCTAAAACAATTTCAAAAACGACGGA

GTTGGTGGTGCTTTGCATGCCTTTCCCTTGGAGTACTAACAAAAGGTGGAATGCTCATAGAGCTTGATGTGTTTCCAAAT

GAAGATCCTTCATGCAAAAATACAGTAATGGAGTATTATGTAGATGTAACAGATATTACAAAATCTGTTTGGTATGGTAC

AATTTACAAGTTTTGGATTCGAAACATTGCCACTGTGTTTCTTCCATTTGCCCTTCTCTTATTAATAAATCTTGGAATTG

TCCTGGAGCTTCGATCTCAAATGCAACACGCTTTTGGGAATAGATCAAGAAGAAGATTTTCATTGAGAATGCAGAGTCGG

ACAAACGTACGTCAGGCGACAGCCACAATGCTATTCATTTGTGTCATTTATTTGATATCAAACGTAGTGAATGTGTTCAT

CACCGCTTGGGAATTCATTGATATTGAGTCATTGCAAACACGGTTCTTGGAAGAGTACATGTTATCCGCTGATCTTTCTT

CAGTTCTGGTAGTGACTGCTTGTGCACTTCGACTCCCAATATATATGCTGTGCAATCCGGAGCTTCGAAGAGCTGTAAAA

AAGTCATTCACTCATAAAAGCGAACATCAACAAAAGATGCATCAAACCTTTTCTCTGATTGCAAAGATTTTAGTTTAA

>pR3

ATGGACATATCCGGTCAAGTTGGACGACTCTCCGGTGGACTTGCATCGGCTCTTGACGTCGTTTCAACAATTGGATGTGC

AATTTCAATAGTGTGTCTTGCATTGAGTGTATGTGTATTCACATTCTTCCGAAATCTTCAAAATGTTCGAAATTCGATTC

ATCGAAATCTTTGCCTGTGTCTTCTGATTGCCGAGTTGGTTTTTGTGATAGGCATGGACAGAACTGGAAATCGTACCGGC

TGCGGAGTTGTTGCAATTCTCCTGCACTATTTCTTCCTCTCCTCGTTCTGTTGGATGCTTCTAGAAGGATATCAATTGTA

CATGATGCTTATTCAAGTATTCGAACCAAATCGTACAAGAATCTTCCTTTATTATCTATTTTGTTATGGAACTCCGGCTG

TTGTCGTGGCAATATCTGCTGGAATTAAATGGGAAGATTATGGAACTGATTCTTACTGCTGGATCGACACCTCCACCCCG

ACGATCTGGGCATTTGTGGCTCCAATTATTGTCATCATCGCTGCCAATATCATATTCCTACTCATCGCCCTAAAAGTCGT

TTTATCAGTTCAAAGTAGAGATCGTACAAAGTGGGGAAGAATCATCGGATGGTTGAAAGGGTCTGCCACGTTGCTGTGTC

TTCTTGGAATCACATGGATCTTCGGATTCCTCACAGCAGTAAAAGGTGGTACTGGAACCGCTTTTGCCTGGATCTTCACG

ATTTTGAATTGCACCCAAGGAATTTTCATTTTCGTGCTCCACGTTGTGCTCAACGAGAAGGTTCGAGCATCCATTGTAAG

ATGGCTGAGAACTGGAATTTGCTGCCTCCCGGAGACAAGCAGTGCTGCATACAACAGCAGAAGCTTCTTGAGCAGTCGGC

AGAGAATTCTGAATATGATAAAAGTAAACGGACATAGTTATCCAAGTACAGCAAGTACAGATGACAAAGAGAAACAGTTG

ACTCCGATTACAAAGACAACGGATTGGTTGAGCAGGTTGCCAAATCAGGACTCGGTTTCGATTCCGGAGAGCAATTTTAA

CAATTTGAATGGGACCCTGGAGAACTCAAATCTCAATAGCGCAGAAATCAAGGAAGAAGACGAAATTCCAGAACTCCGTC

GACGAGTCACAGTTGACCTGAATCCAATGATAGTGAGCAACAACGAAATCGAGCGAATGAGCCATGCCTCTTCGGATCCT

AGAGGATCCCAGATCATCGAAGTAACGGCGGTGGAGAAGAAAGCGCCAGTAAAGAGAATCAAGTTCCCACTGGGCGCGAA

ACAAAGTGAACGGGGATCTCAGCACAGGACAAAGGCGAAAGTAATAGCACCTCCAGAGTCACCGGTTTCGGAATCTGGAT

CCAAGGATTATCGATTTTAA

>pR4

ATGGACGAACTAATCACCCTGGAAGGTGCTTCTCAATCTGAACAAATCATTGGATCCCATGATTTTTGCAACTTTTCCAA

TATCACCCATCATGAGCACGATGAGCAATCGATTTCGATTGTCTGGTGGAGCAACGTGGCAGTTTTACCCGTGATTGCTC

TGATAGGATTGGCTTGTAATCTACTTAATATGGCGGTGCTGACCTCGAATAAAACAGCACGCCGAATCCCTTCGTGGAAC

CTTTTAATAGCTCTCGCAGTATGTGATAGCTTATTTTTGATATTTGCTACTCTGGATGTAACTCCATTATCTATTCCATC

ACTGGCATTTTCCACTTCTTTCAATCATTTTTATTCAAGAATTGTGCTTTATATTCGGACATTGGCATCAACTTTCTACA

AAAGCAGTGTTTTAATTGTTGTGGCCTTCAACATTGAACGTTACCTGTGCGTCGTTTGTCCGCTAAACTCGCATCGATGG

TGTACTTCAAGAAACTCGAAAAATGCGATTGCCACTGCAATTGTGCTCTCATTCCTGTGCTCCATTCAATGGCCATTGGC

CTATGACACAATTCGTTGTTTCGAAAGCAATTCAAATCAATACTACTACGTGATTCTAATGTCGACAAACCGTGCTCTTC

AGATTTACTACCGAACAATGGACTATGTGTCTTTATTCGCTTTCAATGTCCTCCCAATTATTGGTCTTCTCTACATGAAT

TCTCGAATAATCTTCACTCTTCGCAGAGTTGTTGATGAAGATTCAAGAAAATACGAAGAAACCAAACTTTCCGATGGTTT

GATTCAGCATGATGCTCATAACAACCGAACTATGAGAGCAAATGCAATGCTTTTCGCAGTCGTCTTCATGCTTTTCTTCT

GTGTTGGACCTCAGGCTCCTGCAAGAATTCTGTTCGATATGTATGGACAGTATCATCCAAAAGCCATACTTTATGTATGT

TTGAGCCAACAACTGGTATTCCTGAATGCGTCACTCAATTTCTGTTTGTATTGTGTTGTGTCAAAGAGATATCGAACGTT

GATGAAACAGACTTTGAAGAAGTTCTTGCACAAACTTGAAGGAGTCGACCATCCATTCCAAATCAATCTGAAACAAACAA

AGAGCAGCTCGGCACATGTGACATCTCTTGAGGATCATCATGCTCACCTCATTCAAAATGTCTAG

>pR5

ATGGACTCAAACGTGAAATATTTCATGTATGAAATTTTTATTCCATCAATTATCATACTATGTTGTGTCGCCGCATTTCT

CAATTTCATGGTTGTCATCTCCCGGTTGTACTGTAAAATGCGATCAGCCTCACTGGAATTGACCTACTCGTTGGCGCTAT

CTGACACGTGGACAAGCATCGTCATTGGATTTTCCCTTTTTTGGAATAGTTACAAGCCGGTCGTCCTGAATATTCCACAT

TCCAGTTATTGCTTTCCGTTAACTCTTGAGGCATTCCGAACCGGAGGTCTGCTTACCGGAATTTTCCATCTCGTCGCACT

GGCATTCACCCATTACATGACCATTAAACGACCATTCGACCATCATAAAGTACTCCCGATTCGAACAATCTACATTATGA

TCTTCTTTATGTGGGCAACACCGCCAATGGCTCTTATGATCTATTTCGCCAGCAATTCGGGACAGGGATATCAAAGCGAG

AAATGCATGGGCATCAAATTTTACGAGAACTTCTATTTCCGCGCACTTGTCTCTCTTATCATTGTCTTTCTCATCATTCT

TACCACCATCTTCTACATCAAAATGTTACAAAAAATTACCGAGGTCCGTAGCAAAACCGCTTCCAACTCCCAAACCCTCG

GAGCATCTGCCCGTGGACGGCGTACAGTTGTGACTGCAGTTCTGATCTTCGGAACTTTCCTGATCGGGTGGATGCCAGCA

AGTATCCTGTACATCTTGACTGCAGAAAGTATGCCCCTGTATAATAAGCACAGTGTCTCGATAACTATCATGTCGATCGC

CGTGCTGGTCAGTATTATGGCGAAGACGTTGTGTAATCCAATTATCTATGCGACAAGAATCCCTGAAATCAATCAATTCG

TGTTCCAAAAGCTGCTCTACCGCGTCCTACCCGGCCGGAATCCGACTATTCGAAGACAAAGTGAGCTTGAGCCACTGAAA

ACGAGATGCTCACAACCAAATGCACATTCTGTGATGCTTTAA

>pR6

ATGGAAGGTCTCAACATGAGCTTAGAATGTACACCGGCAAGTCGTTCAAACGGGACCTGTTACGGTTTGGCAGTGTGTGG

CTATTGTTACGACTTTGTAGAAACAAGGCAGGTTGTCAAGGAATACGAACAGTTCAATCTGGTCGTCATTGGACTAATGC

TTCCTCTCATCGGATGCCTTGGGCTGATTGGGAATGCTCTTTCGGCATTCACATACTCACGAAGGGAGATGATTTCTAGC

TTGAATGTATACCTATTTGCGCTTGCCTGCTCCGACATAGTTATCATCCTAACCGCATTTTTCCTTTTCTTTCTGGAAAA

CATGAGAAAACGGTCAGAATGGGCAACATACTATTTTGCCGTGCTATCACCTGTCATGTTCCCATTGGGACTTACTGCTC

AAACAATGTCTGTCTTCATCACGGTTGCATCAGCGTTTGACTGTCTAGTTTTGGTGGCTGCCAGTGAAAAATTTAAGAGC

AAATTCTGTTCTGTGAACACTTCAATTCTGATAATTGTGAAAATATTCCTGCTGGGAATATTCTATAATTCACCTCACAT

GTACGAGATTTACGTCATTGATTGTTGGAGTACCATGTACAACACTGCTTCTAAGGACGTTTGCCCAACAGCTCTACGAT

CAAATGTGGATTATGTCCGAATCTACTACGTTTACATGTACACTATTGTGATGGCAGTCGGCCCAGTGCTTCTCCTCATT

GTCATTAATACTGCCATTGTAATTAGTATGCGACGGTCGTCTTCTCCGAACAGCGAATCCGACATTATCACCTTAGTGCT

GGTAGTTTGCCTGTTTATTTCGTGTAATGTTCTGCCCTTGACAGTCAACTTTCTGGAATTGCTCTTCGGAATTATTAACA

GCTACTTGATCGATTTATCCAACTTAATGGTCGTTGTCAACTCTTCTTGCAACTTTTTGATCTACTACACATTTGGCTCT

AACTTCCGTCGAACTCTCCGTTATTATGTGCGTGCTGCTCTCAATCGACGCGCGCCGGCTCAAAATGCTGCCAACAACCG

AACACCACCAAGAGTGAAATTGTGCCTTCCCCCAACTGAAGTTCTTATTTGA

>pR7

ATGCAGACTGACGATTCTTGGAAACATAATGTGTCGTTTTATACATTGCAAGCTTTATTCACCTCAGCAAACCGTCGAGA

TGACTTCATAGCTGTATCAATATGGACCATAATGCTACTTTATGCCTTAATTTCAAACATGCTCATTTTAGCTGGAATTG

CTCGAAGCTCAACTATGAGATCTGCCACAAGTTACTGGTTTATCATATCAATTGCAATCTGTGACATATTAATGACATTT

ATCTCACTTGGTCATTTAGTACCAGCTACTGCATTTCATGAAGAATATGTACAATTCAAATCGATTCGAAATATTGTTAT

GATATTTTTCTACGACCTGTTTTGGTACACCGGAGTTGTTCAGCTGGGGTTGATGGCTGGAAATAGATTTGTTTCAATTG

TGTACCCTATGGAGTACAAACACATTTTTTCGCGAACAAGGTCGTTATATCTTATTTTATTTGGATATTTCCTGGGTTTT

CTCGTTTCTTTACCAACATTATTTGATTGCTGTCATACACTTTGGGACTCGAATTATTACATCACAGTGTACGAAAAGCC

AGACACGTTATACAAGTACGTTGACATGGCTGTAAATAGCATATCGCTGTGCATGATGATTATTTCCTATGCGCAACAAA

ATGCACTTGTGAATGGAGTATCGTTGTCCTGTCAAATGAGTGAATGCGGGCGAACTTCCTCAGTTAGGCCGCCAAGAAGT

CAAGTTAGCAAAAAGGAAATGCGGTTATTTATTCAGTTTTTCGTGGTCTCCCTGGTATTCCTACTAACCTGGACAACTTG

GCAGTGGCTGCCCTACATGTCCGAATCCAAATGGGCGTACTTTGTAATGACATCCTTATTCTTCATCAACAACTCGGTTA

ACCCAACAGTCTATATCATTTTCAACACTCAACTTCGACGGGAACTTCATTATCTCATTTGCCGACATCATGTTATAACA

ACTGCTCAAAATAAGAGGAAACAGACGTTGTTTGGGCGCGGAATCGCGGCTGCCAATAAAATTGAAACGGATTTTCAAAA

TAATACTCGAGACGACGCCACCAAATCATTGGTTGATCAAGCCGGCTCTCTTTCCTCTCAATCGCACGGAACAATTGACG

AGCATGTAAGACACCAGCTTCTGATAAAGAATTTGGACTACACCGATAAGGATACCAAAATCAGTGCAGTTTAA

>pR8

ATGTATAAACTGGAATTTCCTCTTTATGTAGTTATTAGTTACATAGCTCTAGCTATTATCGGTATTCTTGGAAACTTTAC

AATAATTCTAATAACTATTACAAATAAATGTTTACATTCCCGTTGTTATATTCTTATTGCACTCATGGCTTGTTTCAATT

TTGTATATTGTATTTATGCGGCTCAATTGAGAGTGATGATTTTGATCAATGAATCAGTAATGAGCTCATCCACGTGTTTT

CTCATCTCAATTTATGGAATTTTCGCTATGAACATGCAAAGTCTATTGGCACTTTTTATTGGAATCGATAGAGTATATGT

GATCGTGTATCCGGGAAAATATGCAAAGCTTTCAATCAATTACTATTACTTAATGGTTACCTTGTGTGCTTTGATTTCCT

TACTAGTCACGTTTTGCAAGTTTGTTTTTTATTCAGATGAAGTGGTAACTAATTGTCTCCTGAGCACTGCACTTCACGGT

ATCCCATTTAAAATTTGGCAGTTCATGAATTTTTTCATAACTGTGGCTGTAATTTTTGTTTACAGCTTGGCTCATATAAA

TTTCCAAACTTTAAAAAAAAGTACCGTTCATATGAAAACTGCAAAATCAGTCGATCGAATTATGAAGTCAATGCTGATAG

TCATCGCATTCTACGTTTGTACTTGGACTCTTGGGCTTTGTGGGATGTTTTTGTCTCAGTTTTTACAAGCTGACGATATT

CATGCATTTATTATTCAGAAGTTATCTGGATGGCTCGTCCTTTCTAATTCTTCTCTCAATGTTTTTATTTACTTTTGGAG

AACTCCCGATTACAGAAAAGCAATTTTGAAACTTTTTGGTATATCCAAGTCCGTCGTGGATTCAAGGATTCATGTGTCTT

ATATGCCGAATAATCCTGTTGCTATTTTTTGA

>pR9

ATGACGTCCTCATCATCAGAACCCCGGGACCTTGACGCAGAACGTGACGCCTTCATAAAAGAATGGACTGCCACGTCACA

ACTACTGTACTATACAGTAATCTGCGTGTCGGTGGTCACTTTACCTGCGTTGATATTCATGTTCATCGTGCTTCGAAAGT

ACAAAAAATCCAACTATTATCACTTTTTGTGCGCAATTATGGCCGGCAATTTCGTGCTGCTTGCCACAATTTTTTCCAAT

GTGATTTCGGATAGGAATGTAATGATTTTTGCTGGTATCATCCCGGGAGTAGTTGTCTGCAAGCTCAGCGCATTTCTGGT

GAACACCTCCTCGTTTTTTGTACATTGGGCGTGGGTGGCCATGTATGTGCAGCGATTCCTTCACGTTTTCTTCCCATTGC

GGTCCCATAGAGCTGGCGACAAGAGCAAGGAGATTATTGGCGTATTATTCGCATTCTCCGTCATATCCCAGCTATGGACC

CCAATCCTTATCACGGAGTTATCAGTAGGTGCAGACGCCTCATCAGGCACCTACTGCGCCGAGGACCCGAGAATCTTCGG

CGAGATCACGCTCCGATATCTGATATTTTTCGAGTGCTTCATGACATTTTTCCTGCCGTTAGTGCTCACAGTTATCACGG

ATTTTTCGGTGTTGATTATGCGAAATCCGTGTACTTCGAAAAATGCGTTTACTCTGATTTCGGCGGATGAGATTGTGCAC

CATGAGAGTACAGATGAGAAGAGTCCGCTGAAAATTGTGCACAAATCGAAGATCATGCTAGATATTCGTCGGAGAAACGA

CGCGATCCGCCGATGTTTACTTTTGGCTACCGTAACCCTTTTACTCAATCTTCCAAATTATTCTCTACAACTTATTGATG

AATTCTATCATTTCCGGGAGAGCGAGGATTTGGCAGATCGGCGGAGGTTTGTGAGGGCTGACGCAATTGTCTACATAATA

TACCTACTTCAATTCCCAACCGTTCCATTGTATATGTACTGCCTGAAAACTGATTTCGATAGGACCAAATTGTAA

>pR10

ATGCTCAACCATACGGTAGTTATGCGTGAGTGTGAATGCCTTCATGAGCCGATAGAGGGGTACGCGGGAGTTGCCAATCT

GCTGCTTATTGTCATCTCGCTCCCGTTAATATCATTCGTCGGCGTCGTCCTCAACTTCTTTAACATCTTCATTTTTTGCG

ACCAGAAGAACACCGCCGCCAAGTATTTGACGGCACTCTCGTGCAGTGACGTAGGAGTATGTATGGCAGGGATCTTCGTC

ATTTGCTCCGACTCTCTCCGAGCACACTCATTTGTCATTGACCAAGTCTTCGTGTTCTTGTTGCCCAAAATCATCCCGTT

GGGCCTCTTTTTTCAGATGCTCAGTGTATATATCACAGTTTTGGCTGCATTTGACTGTTTCTATTCTGTATATTGTGGAA

CAAAATGCGAGCCAAAACGCAGCACATGGGCTCCACGGGTTCTCGCGCTTGTCGTCATTTCGGTGGCCGGCTATAATATT

GTTCAATTTGGAGATTTGCAGGCTATTGAATGTCTTCACCCGGACAATTACACACTTTTTGAGTTATGCCCTACTGAAAT

GAGGGTCAGTGAAACATATGTCATTGTGTATAAAGGATATCTTTATGCACTTTCAATGGCATTTCTCCCATTTGTACTTC

TCACGTTTTTAACTGTTTCGATCATTGTAATGCTTCGTCGGAAAAACGAAACAATGCACATGGACAAGGCAGAGAAAGAA

GAATGCGAAGATGACGGGGGCAATAATCCGGTTGTACTTCTTCTAGTGGTTCTCCTATTTCTCTGTTGCAATTTGACATC

ACTTCTAGTCAATGTTTTTGAAATGCTCAAGTTTAATATGGCATTTGAAACAGAAGCAGTGCTCATTGACATTGGAAATT

TTTTGGTGGTTATCAATGCGACGGCCAACTTTTTTGTATACATGGGGTCGTCGGAAGAGTTTCGCTCGTCATTTTATCAG

CGGATTCGGCAATTGTGGAGGCCAAAATCGACGAGATTACCCTTGCTTCATGATCATAGAATACGACTTTTCAATAGGAA

TCCGGTTGCGAGTAGCATATGA

>pR11

ATGTCTACAAATTTGGTGGACTATGTCGATGATTCGTATTTGAATCAATCAATGAATTCAGAAAATGGATTGGATTCAGT

CACACAGATTATGTATGATATGAAAAAGTACAATATAGTGAATGATGTTCTACCTCCTCCCAATCACGAAGATCTACATG

TTGTAATAATGGCAGTTTCATACCTTCTTTTATTTTTATTAGGTACTTGTGGAAATGTGGCAGTGTTAACTACAATATAC

CATGTTATTCGATCGTCTCGAGCCACGTTGGATAACACATTAATATATGTCATTGTGCTTTCTTGTGTCGACTTCGGAGT

TTGTTTGTCACTTCCAATTACGGTTATTGATCAGATTCTCGGTTTCTGGATGTTTGGCAAAATACCATGTAAACTTCATG

CAGTATTCGAGAATTTCGGTAAAATCCTGAGTGCTCTCATTCTGACGGCAATGAGTTTTGATCGGTACGCCGGAGTTTGC

CATCCACAGCGAAAACGGTTGAGATCAAGGAATTTTGCAATTACTATACTTTTAGTTCTTGCCGTATATGCATTCATCAC

ACTCTGCCCGTTATTATGGTCTTTTACTGCACGGGAAATTATACTTTATGCGAAGGAAACAGCACCCGGAATGCTGACAA

GAATGAAAATTGAGAAATGCACAGTGGATATCGACTCACAAATGTTCACAGCTTTCACGATTTATCAGTTCATTCTCTGT

TATTGTACTCCATTGGTTCTCATCGCCTTTTTCTATACGAAACTTCTCAGCAAACTCCGTGAACACACAAGGACGTTTAA

GAGCTCTCAAATCCCATTTTTGCACATTTCGTTGTACACTCTAGCAGTTGCATGTTTCTATTTTTTATGCTGGACCCCAT

TCTGGATGGCTACATTATTCGCAGTTTATCTCGAAAACTCAGCAAATTCGAGTAGTGTTCCACCAGTTTTTGTATATATT

ATGTATTTTATTCATGCTCTACCGTTCACCAACTCTGCCATTAATTGGATTTTATACGGTGCACTAAATGGCCAACTTCA

ACAAAGATATCGATCAAATCGTTCAAATTCTACAAAAAAGACGACAACAACAACAGCTTCAACAGCTTTATTGGAAAAGA

AAATCACAAATTTGAATACTAACTCTAATTATCAGGTAAATGGCTCAATGAACTCAATAGCCACTGCAGCTCCAACAAAA

ACGATTGGAAATAATGAAGTACTTGTTGCCACGTCAACAATTGATGATGATGTTGCAACTGATGTTGTAGATGTTCGACT

TTTGAGTAATCATAATCCAACTTTTCTTTGA

>pR12

ATGGATGATAATTATGGTTTATTTTTCCCAAGTCATTCGATGGTGTGCTACCATGAACCTCGAATACACGACCATGAGCT

GTATTTTCAACTTCGGAATCTCAATCTTCTTTTATTGCTTCCAATTGCAATTGGTGGAATTCTTTGCAACGCGTGTGCTA

TGGTCTCATTATATAGACCACCAAAAATTACATCTGGAGTATTTGTTTATCTGAAAGCACTTCTTCTTTTGGATCACGTG

ATGCTCACCACCACCTTAGCAGCCGAATTTTTCCCTCAGATCTGTGATCAACACCATATGAAAAATCATACATTATATGG

TCCATGCATGTTTGAGAGAAGATTTTTAAAATACACAATGCCTAGAGTCGAAGTTACAATACATACACTGCACGTCTGGA

CAATAGCCACACTTTCAGCTCATCGTTATTGGAAGATATCACGTCCAATGATTGCAAGACTTCAAGACACGGTTAGCAGA

GCTCGGACAATGTTGATAGTAATGTTTTGTCTGGTAATGTTGTTCCGGCTGCCTATTTTTGTATTGGAACTTGAATTCCG

GTCACATCCTCAATTGAGAATTACCAGAAGAATATCCACAACAGAGATCCTTTCTACGTATCGATTTGTTTACCATTCCA

TCTTGGATCCTCTCTTTTTCAACATTGTCCCATTTATGTGGATGTGCATTTTTTCGCTGCTCACTCTTTATGAAATCTAT

AAAAGTCGCCACAACACGTATCAACATTTGGCTTTTGATCATCCCAAGCAAACCAATGTGACACATCCATTAGCAGGTTG

CTTTCCAAAAAAAGCTGAACTAATCCGGCAGAAACAAGAACTTCGAGCCACTTTCTCAATTGTGGCTATAATCCTTTTGT

ACTTGTGCTTTCACTCGCTTAAGTTATTTGCCGTTGTTCGAAAATGGCAACTTTTGGTAAAAAGGGAATGCCCAACGAGA

AAAGACTACTACCATTCACACCTCTCTGACGTTTTGAGTATGATTTCCGCATCGGTGAACGCATTCGTGTTCATTGCGTT

CACGAACCGACTAAAAAAGTATATACGACTGCTTTTACGGAAAACATCACGGACTCTATCCAACTCCAGTGATCCGCCCA

TGAGCCCGAAAACTGTGACGTCATATGAAAGTGGATGCAACAACAATTTTAATATTAACATTTGA

>pR13

ATGGAGGGTGGTCGAAACTGTGTAATGACAGTACAACAGTGGCAACCTGAATACAATGATATGAACCAGATAAGAGCAAT

ATTCTCGTTACTGTACCTTCTTGTTTGGGTTGGAGCTATTGTTGGCAACACTCTAGTACTTTATGTACTCACTTTTAATC

AGGTATCACTGTCAGTTAGGACGGTATTTGTTGGATGTCTAGCTGGTTCAGACCTTCTAATGTGCCTTTTCTCGTTGCCA

ATTACCGCGATTTCTATATTTTCAAGAGTTTGGGTATTTCCTGCTATATTTTGCAAGTTGATCGGAGTTTTCCAGGGCGG

TACGATTTTTGTCTCATCATTCACATTAACAGTTATTGCTCTGGACAGATGTGTACTAATTCTACGTCCAAATCAGGAGA

TAGTAAATTTCCCGAGAGCTGTCTTCATTGTTTTCTGCATTTGGCTTCTCGGATACTCTCTAGCACTTCCTGTAGGCATC

TACAGCGACATTGCAGTATACGACGAAATTTGCGGCACATTCTGTGAAGAGAATTGGCCTGATTTCAATCCGGATACTGG

AAGATCGGGAATTCGAAGAGCTTATGGACTTTCTGTGTTGGTACTTCAATTTGGTATTCCTGCATTGATAAGTTCAATTT

GTTACTGGATGATTAGTCGAGTGATGTCAGATCAATTAGCAAGAAGAAGAGGGCACAATATTCGACCGGAATCTGAAACA

AAGCTGGTGAATCGAAAGACAAGAGCTAATCGAATGATGATCGTAATGGTTGTTGGATTCGTTCTCGCGTGGATGCCATT

CAATGCAGTCAATCTCTACCGTGACCTATTTGGAATTTCTAAATGGTATTCTACAGTCTTTGCGCTTTGCCACGTATGCG

CAATGTGCTCCGCTGTGCTCAACCCAATCATCTATTCCTGGTTCAATCCTCAATTCCGACAAAGTATCACCACTTTGTTC

AAGGGTACTGATGAAGCTAGATTGATCAAGAAGAAACCACAATCTACCAGTAAAATGGTTTCGTATCCGACTAATTTCTC

CGAGATCCGAAAAGAAACAGAAATTGCATCGACAAAGACAAAAATCACAATTGCTGAAAACGACTATCGAGCTGGAGATC

AACTTTTATAA

>pR14

ATGGCGGATGCCACGTCACCGTTCAACGTTTCGATTCTCGATAATTCGACGAAGCTGAGCGAAATGGTTGAATCTGGTTG

GAACGTATTAGCGTCGACCAGTGTTCAGGCGTTCAACGAAGCAATGGATGTTCTTGAAGAGTCTTATCCGTTGTGCAAAA

AGATGCTTGATCACAACAATTTATTCCCTGAACGAGATCCAAATGATACACGAATTTGGTGTAATGCCACGTATGATACA

GTACTATGTTGGCCGCCGACACCTGCAAATTCATCAGTCACATTACAATGTCCGCATATGAAAGGCCTAGACCCAAACAA

ATACATTGTAAAACGGTGCGACGAGACTGGACGATGGGCAGGAAAGAAGCCGGGTCACTACGAGAATCCCTGGGGATGGA

CCAATTTCACCGTCTGCTTCAAGATCGATTATGAAGATGCAAAAATAGCTCAAGAAGTAGCACGAAATGCAAGGAAACTG

GAATTCGTGGGACTCGGACTGTCATTAGTATCTCTGATATTGGCGATTTCAATTTTTTCGTATTTTCGGCGGCTTCGAGT

ATTCCGAAATCTGCTGCACTTACATTTAATGATTGCAATGCTTATGGTTGTAATACTTCGGCTGGTACTCTATATTGACT

TAATATTCACTGGTGAAAATGGTCCTCACACGAACTCAGCAGAAGGAAAAACGATAAATACTATGCCAATCGTTTGTGAA

GGAATGTTCTTCTTTTTGGAATATTTCAAAACCGTTACGTTTTGTTGGATGTTCTTAGAAGGAATTTATTTGAATAATCA

AATTGTTTTTGGATTTTTCAACTCGGAGCCAAAATTGTTACCCTATTTCATTGCTGGATACGGTATCCCATTGGTTCACA

CAATGCTCTGGCTACTTGTGGTGCTCATCAAAAAAGACTTCAAAGTGGAACGATGCTTAGGATCCTATTATTTAGAACCA

GAATTTTGGATTCTAGATGGTCCTAGGATGGCAGAACTTGTGATAAATCTATTCTTCATCTGTAATGTCATTCGAGTGCT

CTACAGTAAAGTTCGAGAATCTAACAATACGTCTGAAGCAGGTCTGAAAAAATCGGTCAAAGCTGCAATGATGTTATTGC

CGCTGTTAGGTGTTCCGAATATAATGCAAACAATTCCGTTTGCACCAACTAGGGATAATATTATGGTATTTGCTGTTTGG

ACGTACACCGCCTCTTTCACCTATATGTATCAAGGACTTATGGTTGCCAGTATTTATTGCTTTACGAACAAGGAGGTTAA

TCACGTCCTCAAGACCTTCTATGCTCGGTATCGACTCCTCCATAAATCACAAAACGAACTTCGACGAGGATCACGAAGCG

TCGCATCCCATTATGCTGCAAAAAATGGAACAGCTAATGCAAGTGCACCACAGACAAATAACGCTGACGAGTTTGGAAAA

CTTTCTCCGTTCCCGTCAAGGAGTAAAAAAGGTAGCGATGATTCCACGACGAAGCTGATGAAAGATGCAGTGATGGAAGA

AGAAAAGAATGCAAACAATAATGGATACGGATCGGCTGGTGAAATGACACCTCTTCGTGAAGGATCCAATCGCTCCACAA

AATCTCCATAA

>pR15

ATGTCACTAATCTTACCCAACTCAACACCACCGTTCATCAATCCATTAGATGAAATTTCAAATCAAGAGACACTTGGACA

TGTACCATCGGACTATGCTCAGATGCTCCTTTATGGAATTTTCGCGCTAATCGGACTTCCAGTGAATATCTCAACTTTGA

TATACATGCTAAAACGGTATCGACATGCAAAGTCATTCCTACTTCTTCTCCATATCAACCTGAACATTTCCGATATTTTG

GTTCTTGGTCTATACGTGCCTGGATTAATTGGGTGGCTGGTGACTTTGGAATGGCGTGGAGGCGCATATCTATGTAAATT

CATGAGATTCGTTGACGCCTTCGTTTTTGCGATCAGTTCAAACATAATGGTATGTATCGCCTTGTATCGTCTCTCGGCTC

TCCGTTATCCTCTTTGGGTGAATGCAGTCGGACACTCAAGAGTCCCGAGAATGCTCATTTTAGCGTGGGCTCTTGCGGTG

GTCACAATGCTCCCACAGATGCTTGTTTGGAATGAAGTTCAATTCTCGAACATAACTCAATGTGTGACAGTTTGGACAGA

AATAATTAATAAACGGGAAAAATTGAGTGAAAGTGAGCTTCGTAACATGAAGCTCTATTCAATTCAAAATGCGACCATCA

TTTTCTATATTCCGTTGATGATTCTCGTTGCTTGTTATGTTCTAATTCTGAAAGATATTTACAAGACTCTGAATACGGAC

ACCGAATGCTCTTCAGCTGCATATCTTTCTGAAATGAGCTCATCAAAGACTGGAGGAAAAGCTGCTCTTCACAAGAAGGA

TCAAGAGAGTTTTGTCACCCTAACTACTCGAACAGTTCGTGGACAGGAAAAGTTTCGTCGAGCAAAAGTTCGGAGCCTCC

GAATCACACTTTTGCTCATTTTGACGTACGCTGTCACTTGGCTTCCATACAATTTGCTCTCCTGGTGGATGGTTCTTCAT

TGGGATTCGTATCGTGAGAATCTGGATTCCAACTATATTCTGAATTCATTGGTGGTTCTAAACAGTGTCATTAATCCATT

CATTTATGGAAGATGCCAGGGGATTCGATTCTTGTTCAAGTGTAGAGAGCGATTGGCTGCTCCCAAAACAAAGAAGATTT

GCAAGTTATTGCAAAACTAA

>pR16

ATGAATGGCTCCGATTGTCTGAATCTCAACTCAGAATTATGGTTGTATCGAGAAGATTTGTCATCAAGGTGGTACATAAT

GTTAGTGTTTGCATTTCTCTACCTGATAATCATTGCCGCCGGAATAATTGGAAACTCATGTGTGATTTTGGCAATCACAA

GGAACAAATCACTTCAAACTGTTCCGAATCTGTTTATTCTTTCTTTATCATGTTCTGATATTGTGGTATGCTGCACATCT

GCAACAATCACTCCGATTACTGCATTCAAGAAAGAATGGATCTTTGGAGAGGCTTTATGCCGAATTGCACCATTCATTGC

TGGTATCAGCCTTTGTTTCTCAACTTTCACATTGACTGCAATCTCCATCGACAGATACATCCTGATTCGATTTCCGATGA

GGAAACCTATTACGCATTATCAAGCGGTTGGAGTGATTGCTATTATTTGCGCTTTTGCTGCAACCATCACATCCCCAATA

ATGTTCAAGCAAAAGCTGGGAGAGTTTGAGAATTTTTGTGGGCAGTACTGCACGGAAAACTGGGGAGCCAATGAAAGCCA

GAGAAAAATTTATGGTGCAGCTCTGATGTTTCTTCAGCTCGTCATTCCGCTTACCATCATCATCATATCCTACACTGCGA

TTTCTTTGAAGATCGGACAAAGCATGATTCTCAAAGGGGCGAAAAAGCAAAAAACAGACAATTGGGAAATGGAATTAAGT

GATCAACAAAGAATCGCTGTGAAGAGAAGACAAAGAACTAATAGAATGCTTATTGGTATGGTAGTCGCATTCGCTTGCAG

CTGGATTTGGTCAGTGACGTTCAACATTCTGAGGGACTATGAATATTTGCCAGAGCTCATCAAAACTCAAGAATATATCT

TTGGAATTGCTACACATTGCATTGCAATGACCTCAACGGTATGGAACCCGTTACTCTACGCAGTGCTCAACCTCCAACTG

CGTGCAGCATTCATTGACCTGATGCCTCACTGGCTTCGTCGTCATTTAAACCTGGAAGGAGACAACAGCTCTCCATTGCT

CAACCATCCGACGATGACAATTACAAACAAGTACGGATCAACTGCAACAAAGACCGTCAAAGCAACATACATTAATACCA

GCAATGGACAACCATACGTGTCAACAAGTTTAGTGGGTAAAGTGCAACCAGAAGCACCTAGTTTTAAATTTAACGGATCC

GGCCGCAAAAAGTCGGCAATGATGCGAATTCTAGTGCAAAAAAGAAATGCAGAAGAAGAAGAGCAACTAATCACAAAAGA

ATCACCGTCACCACCAGAAATCCAAATGGATACTCTATGTGCGGCATCCATAATCCCTCGAAGGAAGAGTGCCCAGCCAA

GGTCCACAAATGAAAAAGTTGTGCTACCAAGGAAGGCTTCTTTCTAA

>pR17

ATGCTTCAAAATACTACAATGAGTTCTTTGTCTTGTGGATTCTCCACTTGGATTATACCATGGCTCCAAATAATACTCAA

CATGGATTCATTTTGTATGGAAACCAATGGCACACGTGCAGAGTTAAGTCCAGAGCTGATTAAAGTCAAAGATGCTTTCA

CAATACTACAGGTAACATGCTGTATACTAGGAGTATTTGGTAATATTTTGAATCTCCAAACGCTAAGAAGTCCTTCTCTG

CAAACTGTTCCATTCATGTACATAAAATCCCTTGCCATTTTCGATCTGATTGCCCTAACTATGATCTTAATTCATTATTG

TCTTGTGAGCACAACGAAAATGGCGTTTGTGATGTATTATACAACCTATATCGAAGCCCCATTCATCAATGGGTTTTTAA

TATCCGGACTATATTGCTCATTTTTGCTCACGCTAGAACGTTATTTCTTAATATCTAGGCCACACCAGAAACCAACTCCA

TTTCCACGAGAACAAGCTCGCAGGAGAATTTATACCATGCTGGCCTTGTCGTTCCTAATTCACTTACCGATGGTGCTTCA

AAGAAGACTATCAGTGAATGAAAATGGAGAGTATGTGATAAAAAACAACGTGGAACTGTTATGCCAAGAGCCTAACTGGT

CGTTGTACAACTATTACAAGTTAATAAGAGAATGTGTGAGGTTTTTGATTGTGATCAGTATGACCATAATTAATATCATC

ATTGCAAGGAAATTACAGATCACAAAAAAACGCCGAAGACGTCTTGTCCGACGATCTTCCCCACAAAACGAAATCCCAAA

TGGAATCTCTGCCGATTCAGAATCCAATGTAAATTCATTCACTATGAGACGCGAAGATTCGAATTTAATACGAAGTTTCA

CGGAAAAGAAACTTACTGTATTGATGATATCGATTTGTGTGATTTTCTTGCTAGGAAATGTTCCTCAAATTGTCGTGATG

ATACTGCAAAACGAGGCCATGGAGACTAATTTCAACTTTCAACTTTATCGTCATTGCTCAAACACTTTAGAAGTGCTGAA

TCATTGTCTCAATTTTTATGTCTTCTGCATTGCTAGCTCGGAATACACAAGGGCATTTCTTCTGAATTGCAACTGTCTGC

AGAATATGATGTACAGATTTCCTAGAATTGCAAGATTTGTGAGTAGGAGAAGAAGCAGCAGTGTGATGGTGAGTTCCGGT

GGATTTGGAGTTGCAAACCGAGAATACTTGAGCATGGAATCAGTCCAGGAGGAACAAAAGAAATGGGTTGTTGATCCAAG

CACAAGCCCATACTCTCAAACGGATTCCAATTTGAGAAGTATTTTGGTAACAGGTGCTCGTGAACCTCGACAGAAAAAAT

CACTCACAATTGTAAACCATTGCGCCGTAGAAGAAGACGTCAACTCTCATGAAGTTACACAAATTTAG

>pR18

ATGACAAGTATTATACGAGCCGAGTATGCAGCCTGCCAAGAATTGAAAAAACTGGAAAACAGTTCGTATAATCCGGGAGG

TTGTTCAGTTGACTTTGACAAATCATTATGTTGGGCAAGTGCACATATTGGACAGCAGATGACCCGAGACTGCCCATTCA

CATTTTGCACTGCAATTCCTGGGTGCGAAGAAATTAAAGACAGATACATGGTTTCCCGGAATTGCAACAGTATGGGCGTA

TGGCAGGACTCGAATTACACAATGTGCATCAAAGTTGTTGAGGAATATGCTCAATGTCTACAAGGATTTTGCAGAGTTTG

TCCTGATGTTCTTCGAGACTTGGTGATATCGGTATCTTTAACGTTATCCTTTGTATCTGTCATACTTCTAGTGGCGGCAA

TCGTTTTGTTTTCGATATTTGACTCGATTCAGTGCCGGAGGTTATCAATTCACAAGAACTTAGCAACTGCGTTCGTTTTT

CGATTTGCTGTTTTGGCGATTTGGACGATTGTGCAAACAACGAACGTGTTTCAAGATTGTACAAGATTAACTCCGCTGCC

TCTTTGGGACTACGAATGGATTTGCAAAGCTATTCTCTGGTTCGTCATATATTTCAACGTTGCGTCCGTCATGTGGATGT

TGATCGAAGGCGCCTTCCTCTACAGTCGATTCACTGTATTCGCTATGCGTCACAGTGATGCACCATGGTCTCTTTACCTG

GCGTGTGGGTGGGGCGTCCCGTTCGTGGTTGTAACGGCATGGGCATTGGTTCATCAATACATATCCAGCCAACAAACAAA

TTCATTTTGTTGGCTGCCTTATGCCCAGGGACTCCATTTGTGGATTCTTGCTGGAACTATGGGATCCGCATTAATTATGA

ATCTTATATTCTTGCTTATGATTGTGGTGATATTGGTGCAAAAGTTACGAACGGAGAATTCTGCGGAATCCAAGAAAATT

TGGAGAACAATAAAAGCAACTCTTTTGTTAGTGCCACTCCTCGGCATTTCAAACATCCCTCTTTTTTACGAGCCGGAGCA

CCCGAGCTCTGTCTACATGCTCGGCTCGGCTATTTTACAACATAGTCAGGGTATTTTTATTGCGGTCTTGTATTGCTTCT

TGAATAGTGAAATCCAGGGAGCGCTGAAGCGGCAATTGTCAAAAGTGCCATTTGAGTTCTTCAAAACTAGGAATCGATTC

GAAACTGAAAGAACTTACGTGCCTGAAGCAAGAAATGCCACGAAAAATGGAGTTCCGATGGAGGAAATGAACAAAACTAA

AAATATTGAGAGTGGTGAAAATACGGAATCTCAAGATCAGGTTAGCACTGGAAAGCAAATCTACTCGGTGTCTACGAAAT

CTTAA

>pR19

ATGTCATCGGCGGCTCAACTGCTACTGGATTCGGCGGTGCGAGACGACGACGATCCGATCTTTCTTTATCATCATCTGAA

TGCTAGTCTAGAAGCAGCAAAAGATGCAGTTAATGCTACTTCTGTAAAACAAATTCTCGAAGCTCAAACTCATTATCTAT

TCAACGATACTATCTTTGGACCAAGTCCAGCAATGGAGTCTACACCGGGAAGATCACTCATTCAAAATCTCTATTACTTC

TACATGCCATTCTGCGTTTTTGTTGGGCTCACTGGCAATACTATGGTTTGGATTTTGATCAGAAGCAACCGAATGTTGAA

TAAACTTCCAACGAACGTCTATCTCCTGTGCCTGGCAGCAATGTCATCAATCTTCTTGCTTTCATTACTAGTGTTTTGGA

TAGAAGAGGTCGCATATATCTACTTCTATGACATATTCCAAGATTCATTGCTCCGAAATTCCTACTACAGTTGCATTTTC

AACACGTTCTTGGCGCATGTCTGCGATTTTGCATCAGTATGGTTGATTGTGCTAGTTGGATTGGAAAGACTATTGCTACT

TTACAGGAAAACAAGAGGATTAACTTGTGAAAAAGCCCGTGCCCAAGTGTTCATCCTTCTCGGCTTCGCAATGGTCTTCA

ATGGATGGATCCTTTTTGTTGCTGATATGGATGAAGAAGGCCAGTGTGACATTAAAATTGAATTTGTTGGAATGTATCAA

ACTATGACTATTGTGGAAACGATCATTTGTATGCTGGTCCCATCGGTTCTAATAATTACTTGCAACTGTCTAGTTGTGAG

CAAATTGAATAGCCACATCAAAAAAAATCCAGGATCTCCGGCAGTGTCCTTCAATACTGCGGATGTGGTGCTGACCACTC

AGACATCAGCACCAAGTGCTACATTGAAAAGCTACACACGAATATCAAGCAGATTCTCAATCGATGTTGAAAAGGCACCA

AAGAAGAAAAAGAAGGGAATACGATATACGGATATTCAATTGACTCGGTCACTTCTTGTTGTCACATGGGCATTTATTTT

GTTAAATATTCCAAATTATGGATACAGAATTGCCAGCATTTTATTTGGAATCTCAGAGCAGTCCTCCTTGATGACTGCAA

TTTCTCTTGGAGCTCACGTATTTCTTTACACTCATCATGCATTCCTATTCTACCTTTACATTTTCTACTCACCACAGATG

AAGCGTCGTCTGAAACCAACTGCAATGAAATTGCTCGAATGTTACTGTTTTAAGCCACCCGGTGATTATACTGATCATAC

TTAA

>pR20

ATGGCATTTTCATTGAGCCAACGAGTCGTGGAAACGATGCCATCTGGAACATGGAACGATTTCACTGATCTTCCGGCAGA

ATTCATTTCTCAATTTGGATCTTCTTCCGAGGCAATGCAGTATTCATGTTATTACTCCGATTCCTTTATTCGCTTTGCAA

CCGCTATGGACGACGTATTAATTGGAGCATGTCTCATATCTACTGTGATTAATTTTATCGTGATTGCATGTTCCGCCAAG

TTATATAAAAAGAAAGGAGACACACTTCACTTGTTTATTTTGAACATGACTATTGGAGACACAATATTAACACTGTTTTG

CCATCCGTATGAGCTAGTCACGAGAAGATATTCTGGAGCACATGTTCATTTCATAACTGTTTTTTTGAATTTTGCAAACT

GGGTGGGTCTTGCAGTATCCGGGCTTTCACTCACTTTGCTGAATATTGATAAGCTAATATTTTTTTGCTGGCCATTCAAA

TATGACATATGGATGTCTTATTTTAGAGCCAAGCTATTTTGTTATCTCTCCTGGATTATATCAATAGGTTTTGCGACATA

CTACTGGATGTATAGTTATATGTACTTTGTAAATGCAACAGTTGATATTCAACAACAGTTTGAAATACAATTTGTGTTTC

TTACAGGATTAATGAACAGTGGCCAGTTTTCTCCGGTGAACAAAATTTTCTACGAAGTATTCACAGTGGTGTTCTGTGTT

ATTCCGATAGTGTCATCGCTGTTAGTATCCTGTTACTTATACGATTTAACTAAAAGGAAACGAAAAACCGTGATTAAGAC

TAGCAGTGCAAGCAAAATTGAGAATAAAGCAACATCTTTTGCATTCATCTTCGCGACAACACTTTGGACATCTTGCAGTT

TGCTTCCTTATAGAATAGCGAATCTAGCGAGAATTCATATTATAGCTTCGTTTAGGTGGCCCAACCTAGACTGTGAATCT

CGTCAAAACTTGAGCTGGCTGACCTGGAGTATGCTTTACTTACTTATTTTGAATCCAATAATTAATCCCCTCATAACTGC

ATTTGCCTACGCGCCATACCGTCAGATGATATACTCAAGAGTTCGGAAATCATCTAGAAAAAACTATCAAGATCAGAATT

CCTACGAAACAAACAAAAGCAACATAACATCAACAAATTCTGTATCCTCGAAAGTGTTTTATGTTGATCTCACGATGGCT

AAACAGAACCATACCAAAACTTTGAGGAAAGCTCCACTTGCCAAGTTCCCAAGCATTTGCTCCTGCGGCTCTTTTGTGAC

CAACGAAAATCTACAGTGCACACGGTTTTAG

>pR21

ATGGGATCGGTGAATGAATCATGTGACAATTATGTAGAAATTTTCAACAAAATCAACTACTTTTTTCGAGATGATCAGGT

TATCAATGGGACTGAGTATTCACCAAAAGAGTTCGGATATTTCATCACATTCGCATACATGCTGATCATTTTGTTTGGAG

CAATAGGCAACTTTTTGACAATCATCGTAGTCATACTCAACCCAGCAATGCGGACAACAAGGAACTTTTTCATTTTAAAC

TTGGCGCTGTCGGACTTTTTTGTTTGTATTGTGACAGCGCCGACCACATTATACACGGTTCTCTACATGTTCTGGCCATT

TAGCAGGACATTATGCAAAATTGCGGGTTCGCTGCAAGGCTTTAACATATTTTTATCCACATTCTCGATAGCCTCAATTG

CTGTTGATAGATACGTGCTCATTATCTTCCCAACGAAGCGAGAACGACAACAAAATCTGTCGTTCTGCTTTTTTATCATG

ATCTGGGTGATTTCCCTAATCCTTGCGGTTCCACTTCTGCAGGCTTCTGATTTGACACCGGTTTTCGTTGAGCCATCGTG

CGATTTGGCTCTTTACATTTGCCATGAGCAAAATGAGATATGGGAAAAGATGATCATATCAAAAGGCACCTACACGTTGG

CAGTTCTTATCACCCAATACGCATTTCCCCTGTTTTCACTAGTCTTCGCCTACTCCCGGATAGCACATCGGATGAAGCTG

AGATTTGCAAACCGAAATCAGAATGTGACAACAAATACCAATACGAGTCAGAGGAGACGATCCGTCGTGGAACGTCAACG

ACGCACCCACCTCCTTCTTGTATGCGTTGTAGCTGTATTCGCCGTCGCCTGGCTGCCACTCAACGTTTTTCATATCTTCA

ACACATTCGAGCTGGTCAACAGTTTTTCCGTTACAACGTTCAGCATCTGTCACTGCTTGGCAATGTGCTCGGCGTGCTTG

AATCCGCTGATCTATGCATTTTTCAATCATAACTTCCGGATCGAATTTATGCATCTTTTCGATAGAGTTGGGTTGAGATC

CCTTCGAGTCGTGATATTTGGAGAGCAAGAGTCGCTCAAGAAAAGTATGCGCACTGAATTCAGAAGCCGTGGAGGCTGTA

AGACGGTAACTACTGCAGAGCCAGCAACCTTCCAGAGAATGAATGAAAGTATGATTCTGAGTGCGATGGAGCAAGATGAG

CAGCTGTGA

>pR22

ATGTTGCTACTACTCTCCCGGGTTCTGGTGAACGTGATGGTGCTGATGTGCTTTGGTGGAATTCTCACGAACTTCATCTC

ATTCTACATCTACACTCGTAAAACATTCCGGAAAAAGAGCATAAACGTCTTGCTCGCAGCCTTGTCAATGTCTGACCTTT

GCGTCTGCGTTCTTGCAATTCCAGTTTTTGCAAGTACTCAACTTCAACAAGTCATTCCGCCAACAATTACTGCAATGATC

ATGGTGTACTTGTATCCAGTGACCATAATGTTCCAGTCAGTGTCAGTATGGTTGCTGGTATCCATAACAATTGACAGATA

CTTAGCCGTATGTCATCCATTCATGGTCAATACGTATTGTACAAGGAATCGCGCTCTCATAACTGTTGGAGTTGTCGTGA

TATTCAGTGTTGCTTACAATTTGATTCGAATTTGGGAGTACACGATAAATTTCGATGTCGCTCCTGAGAATCGTACGATT

GAAGATCTTGTTGTACCGAAGCTTCGAGCAAATCCGCACTTCTTATTGTGGTACCAGAATGTTGCCACGTTGGTGTCGCA

ATTCGCATTTCCACTTACAGTGTTATGTGTTCTCAACATTCAAGTAGCGCGAACAATTATCGAGGCATCCGAGCAACGAA

GAGAACTAGTGGCAAGTGTGAAACGAGAACATTCGACTGCAAAAATGATGATAATGGTTGTTTTAGTATTCCTCGTTTGC

TACATCTTCTCATTTATTCTAAATATATGGGAAATTCTTGACAAGGAGACTTTTGGAGGAGATATTGGTTGGTTTATGAA

TGACATCAACAATGTACTTATTGTTGTCAACTCGACGAGTGCAATTGTTTTCTACTATAAATATTCAACAAGGTTCCGAA

ATCAAGCACGGACACTTCCCGGGATACGGTGGTATGCTAGTATGTCAAAGTTCAATGTATATGATACAGAGGCAACGTCA

AATCGAACAATGGTCACTAGATACAAAGAATCCATGATTTCAATAAGAGGCACGTCAACTCGCCTGAGCAGCTCTCACAA

TTTGCTCTACAAACCGAGCTACTCAAAGCCATGTGATATCTAG

>pR23

ATGACGTCAGCCTCATTGGCAGGCATATCAATATCGCCAATTGTGCCGGAAGAACTCGACGGCACAATAGCACCAATTGG

AACACAACCAATGGACATCACATTAGCATTTGTGTTTTCTGCGATTTCAGTCATTGGAGTACTTGGAAACCTTTTAGTAA

TTACAGTTGTACTTAAAGTTAGAGGAATGAAAACACCAACTAATTGCTATTTAGTATCATTGGCCGCCAGTGACACATTA

TTCTTTTTTGCTTCAATGCCTCATGAAATGATGTATTTATTGGGTCCAAATGATCATTATTTATTTGGAAGTTTAGGTTG

TGTACTTCTAACCTATCTTCCATATCTTGCAATGAACACTTCAAGTCTGTCTATTTTAGCGTTCACAATTGAACGGTATT

ATGGTATTTGTAATCCATATAAAGCCAGAACCATGTGCACTGTTAAACGCGCAACGTGTATTATCTGTGGTATCTGGATC

TTCTCAATGCTGTATCATTCCTATTGGCTTTTCTTGGCTACGTTAATCAAAGATGATATTGGAACATCTTGTAGTTTTCG

GTTGGAAAGAAATAGTCATGCTTACAAGATAGTTTTCCTTCTTGATTTCGTCCTGTGGTACGTGCTTCCAATTATGTGCG

ACATCATTATCTACGCAAAAATCGGTATCACTTTGAGTCAATGTGGAGATAAGATTAAGAAATCGGTGAAGCCGAAAATA

CCAAATGAAGTGATCGTCGAAAAATCAAAGACATCGTCGACTTCAATGGGACATTATTCCGGAAGAGATTCTCATATTTC

GGGGAAACGGAACTCGGCAAAAGGAAGAAATCAGGTTGTAAAAATGCTTGCAATCGTTGTGGCCGTCTTCGCAGTGTGTT

GGCTTCCTTATCGTGGAATGGTTGTCTATAATTCATTTGTATCGGATCCAAAGTATTCATGGAGTCCTGATTGGTATATC

AATTTGTCAAAGACACTTGTATTCATCAACTGTGCTATTAATCCAATACTATACAACTTGATGTCTGCTAGATTCCGTGC

TGCATTCAGATCGTTGTTATCGAAAAGAAAGAATTCCGGATTTAAATCAACAGCTTTGAACCAACGTCACCGATTAAACA

CAATGGATATAATGTCATCAGCTGAACCAAAATCTCCATTAATAACATCAATGGCTGGAACTCCGTGA

>pR24

ATGGAATGTCCGCACGATGCTCAGTTGTTTGATCCGAACAGTAACAGTACGCAGGCATTTCTTCGGAAACTTGCACATTT

TCAAAGATGGTACCAGCCAATTCACGGATATGTATGTGTGCTCATCTGCGTTTTTGGTATCTTCACCAATTTTGTGCATG

TGGCAGTTTTATCGAGACCCAACATGCGCAATTCGGCCGTCAACTGCATACTGACAGCAGTGGCTGTCTGTGATATCGGA

ACAATGGCATCCTATCTCATTTATATTATACATTTTGTCTTACGACGAAATAATTCCTGCACTCCAACGTTCACTCATTC

TTGGTTACAATTCCTATTGTGGCATGTTGTTTTGAGCATCACACTACACACAACTTCTTTATGGCTCGCTGTGGCGATGG

CGTTCATTCGGAGGATGACATTAAGGGTTACCGCTTTGAATAGTCAATGGCAACGACCAAAGTTCGCATGGAAGTTGTGC

CTGGGAATCTATATGGTCGTATTTGTACTTTGTATTCCAAATATGCTGGTCCACGAAATCGCCCGTGTAGAAGGAAATGA

CTGGCGACCTGGCAAAACGTGTCAAAAATTTGCATCGAACTACTCTGAACCGGTGTACACTTTTATGGTCAGCCACGCGG

CGACTATCAACAATTGCCGGTTGTTTAAATGGAATATTTGGATGATTGGAATTCTGTTCAAGATCATTCCATGTATTCTG

CTAATTTTCTTGTCTTTCGGCCTTGTATCAAAGATTAGAGACGCAGAAAAGCATCGGAGGAAACTAACGAGTGTTCCGTC

AAATGCATCAACAGATTCAAAACCATTGAAAAAAAAGAATGGAACATCAGATAGAACCACATTGATGCTTGTTGTTATTT

TATTGGTATTTTTAATAACGGAGTTTCCACAAGGAATCATCTCGATTCTTTGCGCAATTTTTACTACAGACGTTCATAGA

TACCTCTATTTCTACATTGGTGATGTTCTTGATCTCCTATCGCTTGTCAATTCCTCTGTAAATTTTGTTTTATATTGTGT

GATGTCTTCAAGATACCGGCAGACGTTTTGGGAAGTTATCATTCCTTCGTGGGCTTGTGGATTGTGGACGACCCGTCGTG

GTTCTCCGACGGAACTTTCACAGCTTCAGTTGAACAACAGCTTACGACTAGGAAGAAAACAAAGCTATACGCCACTGGCA

ACTGAACCGGAACCAGCACGCAATGAGTATTGGCGGTATGAAGCTTCCGTGTCGGACAATCAGAACAGCAGTAGAGAACA

TCAACTTTAA

>pR25

ATGGGTTCGGTGAATTTCACGGGTGCCGAGTGGCTTGAGACAGCAGTGACAAATATAAATACAGCGTATGCTCATATTCA

TCCGTACGTCTCTGTGATTCTCTGTCTTGCAGGTACTGCAATGAACATAGTAACAGTGATTGTTCTAACTCGTCCATCAA

TGAGATCAGCCGTCAATTCACTTCTTTGTGCAATTGCTCTTTGCGACATTCTTGTTATGACCAGTGTCCTGGTTTTTGTT

ACTCATTTTCTACTATTTGCGGGGTACAGATGCGATCCAACAGACTACAACATTTATTGGGCATATTTTTTATACTATCA

TTCCCAAGCCACGGTTATTTTCCATGCAACAAGTATTTGGTTAACAGTTCTACTAGCCCAAATTCGAGTGTTTTCTATCC

GGCGAGCAACATCTGTAGCTGGAGAATCTGTAACTAATCAAATGACTTGTATTATTGCTGTTACAACATTTATTGTAGTT

TGTTTGTTAAATGTTCCAAATATGCTGACATTTGAGATAATTGAAACGCCAGCATCTCTTTGGCTTCAATGCAAAGCAAA

CGAGACAGCTGAAGACGATATGCTCGTTTATTTGGTTGCTCCATCTGATCATTGCGGACTACTTAATATTGCATTCTGGA

CAAATGGAGTACTTTTCAAAGTTGTTCCTTGTCTTCTTCTAACATTTTCTATAGTTGCACTTGTCAGTATTATTCGTGAC

GTTGGAAAACGAAGGAAACAATTAGCTCAAGTGATGAATAAGAAAAGAATGCCACGAGATCATACTACACCAATGCTTGT

CGCCGTTCTTTCAATATTTTTATTTGCTGAACTTCCACAAGGAGTACTTCACGTCTTCAATGCAATTTTCACAAAAGAAA

CATTCTACGATAAAATATACATTCATCTCGGAGATGTGATGGATGTCCTCTCGTTGCTCAATTCTGCCGTTAACTTCATC

ATTTATTGTGCAATGAGTCGGAAGTTCCGAGCCGTATTTATCCAGATATTCCTTACATGCTTGCCACAAAAAATTATACG

AAAATACGCAATGGAAGCATTCCTGGACGTTGAAGTATCTCGCATGAGACCTACTGCTGATATGACAAAAAGTGAACAAT

TGGCATTAACGTCACACCGAGTTTCAGCGTCCTCCGTTCTTTTACCTATTTCTAGACCTGATCCACCCAGAATGTATTCT

GGAAATCTGCTTACAGCTGACATCAACGGCGGATACTCTACATCTCCTCGTGTTTCCTTTGATATCGTTCGTAAAGAATC

GCAGATCAGTAGCGTCGTCGATTTCCAACTTGTGGTACCAAATATGCAGCAAGACTCCCTAAGCATTGTTCAAAAAATTC

TCAGATTTTTCCGTTCTACAGATGAGCTTTCCAATAGTCCACACAGAAGAATTAAATTAATGCAATGTGAAACAAGTACT

GCTCTTTATTAA

>pR26

ATGGGTTCGGTGAATTTCACGGGTGCCGAGTGGCTTGAGACAGCAGTGACAAATATAAATACAGCGTATGCTCATATTCA

TCCGTACGTCTCTGTGATTCTCTGTCTTGCAGGTACTGCAATGAACATAGTAACAGTGATTGTTCTAACTCGTCCATCAA

TGAGATCAGCCGTCAATTCACTTCTTTGTGCAATTGCTCTTTGCGACATTCTTGTTATGACCAGTGTCCTGGTTTTTGTT

ACTCATTTTCTACTATTTGCGGGGTACAGATGCGATCCAACAGACTACAACATTTATTGGGCATATTTTTTATACTATCA

TTCCCAAGCCACGGTTATTTTCCATGCAACAAGTATTTGGTTAACAGTTCTACTAGCCCAAATTCGAGTGTTTTCTATCC

GGCGAGCAACATCTGTAGCTGGAGAATCTGTAACTAATCAAATGACTTGTATTATTGCTGTTACAACATTTATTGTAGTT

TGTTTGTTAAATGTTCCAAATATGCTGACATTTGAGATAATTGAAACGCCAGCATCTCTTTGGCTTCAATGCAAAGCAAA

CGAGACAGCTGAAGACGATATGCTCGTTTATTTGGTTGCTCCATCTGATCATTGCGGACTACTTAATATTGCATTCTGGA

CAAATGGAGTACTTTTCAAAGTTGTTCCTTGTCTTCTTCTAACATTTTCTATAGTTGCACTTGTCAGTATTATTCGTGAC

GTTGGAAAACGAAGGAAACAATTAGCTCAAGTGATGAATAAGAAAAGAATGCCACGAGATCATACTACACCAATGCTTGT

CGCCGTTCTTTCAATATTTTTATTTGCTGAACTTCCACAAGGAGTACTTCACGTCTTCAATGCAATTTTCACAAAAGAAA

CATTCTACGATAAAATATACATTCATCTCGGAGATGTGATGGATGTCCTCTCGTTGCTCAATTCTGCCGTTAACTTCATC

ATTTATTGTGCAATGAGTCGGAAGTTCCGAGCCGTATTTATCCAGATATTCCTTACATGCTTGCCACAAAAAATTATACG

AAAATACGCAATGGAAGCATTCCTGGACGTTGAAGTATCTCGCATGAGACCTACTGCTGATATGACAAAAAGTGAACAAT

TGGCATTAACGTCACACCGAGTTTCAGCGTCCTCCGTTCTTTTACCTATTTCTAGACCTGATCCACCCAGAATGTATTCT

GGAAATCTGCTTACAGCTGAGGATTCTGCCTAAGTTAGTTGAAAAGACATCAACGGCGGATACTCTACATCTCCTCGTGT

TTCCTTTGATATCGTTCGTAAAGAATCGCAGATCAGTAGCGTCGTCGATTTCCAACTTGTGGTACCAAATATGCAGCAAG

ACTCCCTAAGCATTGTTCAAAAAATTCTCAGATTTTTCCGTTCTACAGATGAGCTTTCCAATAGTCCACACAGAAGAATT

AAATTAATGCAATGTGAAACAAGTACTGCTCTTTATTAA

>pR27

ATGAATCAAGAATTCTTAATTCAACTTGGCGAGCGTGCGTGTAAAAATGCTGAAAATCTTACTCTCCCTGCTGAACTCGA

GGGAATATTCTTCTGTGCTCCAAGTTCCAGAGAATCATTAGCAACTCAAGTATTTGTGGCAATTGCATTTGTCCTTCTGA

TGGCAACTGCAATAATTGGAAACTCTGTCGTGATGTGGATAATTTATCAACACAAAGTAATGCACTACGGCTTCAACTAT

TTTCTTTTCAATATGGCATTCGCTGATCTTTTGATTGCTCTCTTCAATGTTGGTACTTCATGGACCTATAATTTATACTA

TGACTGGTGGTATGGTGATCTATGTACACTTACTTCCTTCTTCGGTATTGCACCAACTACCGTATCCGTGTGCTCAATGA

TGGCTCTCAGCTGGGATAGATGTCAAGCAGTCGTGAATCCTCTACAAAAACGTCCACTATCTCGAAAAAGATCCGTCATT

GCTATTCTCATCATTTGGGTTGTTTCAACGGTTACTGCACTTCCGTTTGCAATTGCTGCATCTGTCAACTCTCTCTACAC

ATATGACGTGGTTACATCAACTGTTTCAAAAGCTCACGTTTGTTCAGCACCAGTTAATACTTTCTTTGAAAAAGTGCTCT

TTGGGATTCAATATGCTCTACCTATAATCATTTTGGGATCAACGTTCACACGGATTGCCGTTGCATTTCGAGCAACAAAT

GAAGCCACTGACAGTAGTCTGAAGAACAATCACACACGCGCAAAGAGCAAGGCAGTAAAAATGCTCTTCCTAATGGTCGT

TGCATTCGTTGTCTGCTGGCTTCCATACCACATTTATCACGCATTCGCTCTTGAAGAATTCTTCGATGCAGCTCGTGGAA

AATATGCGTATTTGTTGATTTACTGGATTGCAATGTCTAGTTGTGCATACAATCCAATTATTTATTGTTTTGCCAACGAA

AGATTCCGCATTGGCTTCCGGTACGTCTTCCGATGGATTCCAGTGATTGACTGCAAAAAGGAACAATATGAATATTCACA

ATTATTCCCGGATAAAATGAGAAGTATGGCTATTTCTTTGCAAAAAGGAAGAGTGAACTCTTCCTGTTTGGATAAAAAAG

TCAAAGAGAATTCTTCTCAAGATTTGGTTTGTGTGATGCATTCCGAGAAAAATACGAAGAAATATTCAAAAGTACATCTT

CTGAGTTGCCATGAACGGTGA

>pR28

ATGGAAGTTGAAAATTTTACCGACTGTCAAGTATATTGGAAAGTGTATCCAGATCCTTCTCAAAGTATATATGCGATAGT

GCCATTTCTGACCGTGTACCTTTTTCTCTTTTTTCTTGGACTCTTTGGAAATGTGACCTTGATTTACGTAACTTGCAGCC

ATAAAGCTTTACTGAGCGTTCAAAACATATTCATTCTGAACCTGGCGGCGAGCGATTGCATGATGTGCATATTATCGCTT

CCAATCACTCCAATCACAAATGTGTACAAAAACTGGTACTTTGGAAATCTACTCTGCCATTTGATACCATGCATTCAAGG

TATCAGCATTTTCGTATGCACATTCAGTCTCGGTGCGATTGCTTTGGATCGGTATATCCTTGTAGTAAGACCACATTCTA

CACCACTATCCCAAAGAGGAGCATTTCTTACTACTGTTCTATTGTGGATCCTCTCTTTTGTTGTAACTCTACCCTATGCG

TTCAATATGCAAATGATTGAATACACAGAAGAGAGAATATGCGGCTACTTTTGCACTGAAAAGTGGGAATCTGCCAAGTC

TAGAAGAGCCTACACAATGATCGTGATGCTCGCCCAATTCGTGGTCCCATTCGCTGTCATGGCCTTCTGCTATGCAAATA

TTGTTTCGGTGCTCAGCAAGCGTGCTCAGACAAAGATACGCAAAATGGTGGAGAGAACAAGCGCGTTGGAGAGCTCTTGC

GCATTCCCGTCGCATGGTCTTGAACAGTATGAAAATGAGTTGAACGAATTTCTAGACAAACAGGAAAAGGAGAAACAACG

AGTTGTACTTCAGAACAGAAGAACAACGTCAATCCTAGTTACCATGGTTGTCTGGTTTGGGATAACTTGGCTGCCACATA

ACGTCATTTCTTTGATTATTGAATATGATGACACACAATCGTTTTTCCGACTTTATGGCAGAGATGATTACGATATCAGT

TATTTACTGAACCTTTTCACTCACAGTATTGCCATGTCGAACAATGTTTTAAACCCGGTACTCTATGCGTGGCTGAACCC

AAGTTTCCGTCAACTGGTCATAAAGACATATTTTGGAGACCGGCGTAAAAGTGACAGAATAATCAATCAAACATCAGTTT

ACAAAACAAAGATCGTGCATGATACGAAGCACTTGAATGGAAGAGCCAAAATTGGCGGTGGTGGTAGCCACGAGGCGTTG

AAGGAGAGGGAGCTGAACTCGTGTTCGGAAAATTTAAGCTATCATGTGAACGGTCATACGAGGACTCCTACACCGGAAGT

TCAGTTGAATGAAGTTTCAAGTCCTGAAATAAGTAAACTTGTTGCTGAGCCGGAAGAATTGATCGAGTTCAGCGTCAACG

ACACGCTAGTCTGA

>pR29

ATGTCAAGCTTTTATAACGAAGCAAAGTTTTTAATTTCGGATGGAGAGCAAATGAGGATCGATACATATGCAGTTGGTCG

GAAAACCATGCAGCAATCCGACGCATTCAAGGGTACATTTTATGAAAACTTTACATTGGTGTATGCTCTTCCACTTTCAA

ATCATGACAACTCTTCGTTGATGCTGATTGCGGGTTTCTACGCTTTGCTGTTCATGTTTGGAACATGTGGAAATGCTGCT

ATTCTTGCAGTAGTTCATCATGTTAAAGGACAAGATCCGCGATCGCGACACAATACCACTCTGACATACATTTGCATATT

GAGCATTGTGGATTTTTTATCCATGTTACCAATTCCAATGACCATCATCGACCAGATCCTTGGATTTTGGATGTTTGACA

CATTTGCCTGCAAATTGTTTCGCCTCTTGGAGCACATAGGAAAGATTTTCAGCACGTTCATTCTTGTGGCATTTTCAATT

GATAGATACTGTGCCGTTTGTCATCCGCTACAGGTTCGAGTGCGCAATCAACGAACAGTTTTTGTGTTTCTTGGAATTAT

GTTCTTTGTGACCTGTGTCATGTTATCGCCAATCTTGTTATATGCTCATTCAAAGGAACTTGTCATGCATGAAAAAGTCG

ACTTGGATCAAGAAGTCATCACTCGAATGCATCTGTATAAGTGTGTCGATGATCTTGGTCGCGAGTTGTTTGTTGTATTT

ACTTTGTACTCTTTTGTTTTGGCATACCTGATGCCACTTCTTTTCATGATCTATTTTTACTACGAAATGTTGATCAGGCT

CTTCAAACAAGCAAATGTCATCAAACAGACTCTTGTTGGAAGAAGAAGTGGTGGAGAAGAAAAGAAATTGACGATTCCAG

TAGGCCACATTGCAATTTATACACTTGCCATTTGCTCTTTCCACTTCATTTGCTGGACACCATACTGGATCTCAATTCTC

TACTCTTTGTACGAGGAGCTGTACCAAGATACAAAGAGCACTGCTTCTCCACCAACGTACGCGTTCATATACTTTATGTA

CGGCGTACACGCACTACCATATATCAATTCTGCATCCAACTTTATTCTCTATGGACTTTTAAATCGTCAGCTGCACAATG

CTCCAGAGCGAAAATACACTAGAAATGGAGTGGGAGGCCGCCAAATGTCACATGCATTGACGACGAACACCCGACCTGAA

TATAGCGAGCTAATCGCAATACCATCCAGCAGTTGTCGCCCGGATTCTCGAGTGTCGGCAATGATCCATAACAACAATAA

CAACACGGAATCTTTACCTCTGGCACAGAACATCTCGAATATGACTAACAAAGATTCTGCAACTATTGTTCCAATGCCAA

TGTCTGCCAACGTCGACGGGAATGAGATCTACAATTGGATCACTCCAGATACGGAGTCTGTAATTCTGTAA

>pR30

ATGCAGCAATCCGACGCATTCAAGGGTACATTTTATGAAAACTTTACATTGGTGTATGCTCTTCCACTTTCAAATCATGA

CAACTCTTCGTTGATGCTGATTGCGGGTTTCTACGCTTTGCTGTTCATGTTTGGAACATGTGGAAATGCTGCTATTCTTG

CAGTAGTTCATCATGTTAAAGGACAAGATCCGCGATCGCGACACAATACCACTCTGACATACATTTGCATATTGAGCATT

GTGGATTTTTTATCCATGTTACCAATTCCAATGACCATCATCGACCAGATCCTTGGATTTTGGATGTTTGACACATTTGC

CTGCAAATTGTTTCGCCTCTTGGAGCACATAGGAAAGATTTTCAGCACGTTCATTCTTGTGGCATTTTCAATTGATAGAT

ACTGTGCCGTTTGTCATCCGCTACAGGTTCGAGTGCGCAATCAACGAACAGTTTTTGTGTTTCTTGGAATTATGTTCTTT

GTGACCTGTGTCATGTTATCGCCAATCTTGTTATATGCTCATTCAAAGGAACTTGTCATGCATGAAAAAGTCGACTTGGA

TCAAGAAGTCATCACTCGAATGCATCTGTATAAGTGTGTCGATGATCTTGGTCGCGAGTTGTTTGTTGTATTTACTTTGT

ACTCTTTTGTTTTGGCATACCTGATGCCACTTCTTTTCATGATCTATTTTTACTACGAAATGTTGATCAGGCTCTTCAAA

CAAGCAAATGTCATCAAACAGACTCTTGTTGGAAGAAGAAGTGGTGGAGAAGAAAAGAAATTGACGATTCCAGTAGGCCA

CATTGCAATTTATACACTTGCCATTTGCTCTTTCCACTTCATTTGCTGGACACCATACTGGATCTCAATTCTCTACTCTT

TGTACGAGGAGCTGTACCAAGATACAAAGAGCACTGCTTCTCCACCAACGTACGCGTTCATATACTTTATGTACGGCGTA

CACGCACTACCATATATCAATTCTGCATCCAACTTTATTCTCTATGGACTTTTAAATCGTCAGCTGCACAATGCTCCAGA

GCGAAAATACACTAGAAATGGAGTGGGAGGCCGCCAAATGTCACATGCATTGACGACGAACACCCGACCTGAATATAGCG

AGCTAATCGCAATACCATCCAGCAGTTGTCGCCCGGATTCTCGAGTGTCGGCAATGATCCATAACAACAATAACAACACG

GAATCTTTACCTCTGGCACAGAACATCTCGAATATGACTAACAAAGTTTGA

>pR31

ATGGAGATTAACTTAAGGATCGATACATATGCAGTTGGTCGGAAAACCATGCAGCAATCCGACGCATTCAAGGGTACATT

TTATGAAAACTTTACATTGGTGTATGCTCTTCCACTTTCAAATCATGACAACTCTTCGTTGATGCTGATTGCGGGTTTCT

ACGCTTTGCTGTTCATGTTTGGAACATGTGGAAATGCTGCTATTCTTGCAGTAGTTCATCATGTTAAAGGACAAGATCCG

CGATCGCGACACAATACCACTCTGACATACATTTGCATATTGAGCATTGTGGATTTTTTATCCATGTTACCAATTCCAAT

GACCATCATCGACCAGATCCTTGGATTTTGGATGTTTGACACATTTGCCTGCAAATTGTTTCGCCTCTTGGAGCACATAG

GAAAGATTTTCAGCACGTTCATTCTTGTGGCATTTTCAATTGATAGATACTGTGCCGTTTGTCATCCGCTACAGGTTCGA

GTGCGCAATCAACGAACAGTTTTTGTGTTTCTTGGAATTATGTTCTTTGTGACCTGTGTCATGTTATCGCCAATCTTGTT

ATATGCTCATTCAAAGGAACTTGTCATGCATGAAAAAGTCGACTTGGATCAAGAAGTCATCACTCGAATGCATCTGTATA

AGTGTGTCGATGATCTTGGTCGCGAGTTGTTTGTTGTATTTACTTTGTACTCTTTTGTTTTGGCATACCTGATGCCACTT

CTTTTCATGATCTATTTTTACTACGAAATGTTGATCAGGCTCTTCAAACAAGCAAATGTCATCAAACAGACTCTTGTTGG

AAGAAGAAGTGGTGGAGAAGAAAAGAAATTGACGATTCCAGTAGGCCACATTGCAATTTATACACTTGCCATTTGCTCTT

TCCACTTCATTTGCTGGACACCATACTGGATCTCAATTCTCTACTCTTTGTACGAGGAGCTGTACCAAGATACAAAGAGC

ACTGCTTCTCCACCAACGTACGCGTTCATATACTTTATGTACGGCGTACACGCACTACCATATATCAATTCTGCATCCAA

CTTTATTCTCTATGGACTTTTAAATCGTCAGCTGCACAATGCTCCAGAGCGAAAATACACTAGAAATGGAGTGGGAGGCC

GCCAAATGTCACATGCATTGACGACGAACACCCGACCTGAATATAGCGAGCTAATCGCAATACCATCCAGCAGTTGTCGC

CCGGATTCTCGAGTGTCGGCAATGATCCATAACAACAATAACAACACGGAATCTTTACCTCTGGCACAGAACATCTCGAA

TATGACTAACAAAGATTCTGCAACTATTGTTCCAATGCCAATGTCTGCCAACGTCGACGGGAATGAGATCTACAATTGGA

TCACTCCAGATACGGAGTCTGTAATTCTGTAA

>pR32

ATGGGGAAATGGACGGAACTCGGATTGTCGGGTCGTGAAAATGCAACAAATGTCTCGTCGCGAGCCATAACTGACAAGTC

CTTTTTGCAGTACTACGATGAAATTCATATTCCTCTTTCAATTTCTATCTGTATATTCGGAGCCGCTTCAAACGTCTTCA

ATATTATTGTGCTCACAAGAAAACGGATGCGAACCCCAATAAACATCCTGCTCACAGGTCTCTCCATTGCACAATGGCTC

CTAGCGACCAATTACTTCCTTTACTTATTGCTTGAGTATTATAGATATCAATGTGTTCAATTATTATGGTCCGAAGCATT

TACAAGATATAGATTCTTCAATGTCAACCTGAATACAGTATTTCATACAATTGCATTCACAACGACGATCGTTGTCGCAG

TGTTTCGGTATTGTGCACTGAAGTTTCCAATTCAAGCGAATAGATTCATCTATAAATGCCAGCCAGCAATCGCAGCAAAC

GTTATCATTTGGATCATCATTCCTATCATCAGTCTTCCATTGTTTTTCATATCCGAGGTTAAAATTGTCGCTCGTGACCA

CGTTGCTTATGATTTGCAATGCGAAATGGAAGGTCCATTGTACGACCTCAGTTATCAAGAGAGTCCACTTCTCGTGTCCG

CAGTTTTCTGGGCATTTGGTATTGTGTTCAAACTACTTCCGTCGTTAATTCTCTCAATACTACTCATTGCTCTTATAAGA

TCTTTGAAAAGCGTAGAACGAAGGCGTAAAAACTGGAAGCGTACACAAGGTGCCAACATTTGCACCAACTCCGAGAGAAA

AGCAAAAAGAAAGTTGACGACCAGACCGCGCACAACACGAATGCTCGTCATCATTCTACTCCTATGTGTAATGGTAGAAC

TACCCATGGGTATTCTTAACTTGTGCGTGGCTATCTACGGTGAAGAGTTCGGAAATCGGTATTACGATCCAGTGGGAAAT

CTGATGGAAATGCTGACGCTGTTGTACAGCTCTGTTAGCTTCGTTCTCTATTGCACCATGTCCAATGAGTATCTCTCCAC

GTTCCGAGCACTATTCTTCCCATGGACCAGAAAGAATAGTTTAAGAGGAACTCGAAGAAGTTGGAATCATCGACATGACG

ACGAAACAAAGTCCCCACGAACTTTTCTGATAAATAGAACTGCGCCAAGCTCATATGTCGGGTCTTAA

>pR33

ATGAATGACACACTGATCTGTACATACTTCGAATGCTATATCCACATGTTTGCCATGTACTACGATGAAATTCATATTCC

TCTTTCAATTTCTATCTGTATATTCGGAGCCGCTTCAAACGTCTTCAATATTATTGTGCTCACAAGAAAACGGATGCGAA

CCCCAATAAACATCCTGCTCACAGGTCTCTCCATTGCACAATGGCTCCTAGCGACCAATTACTTCCTTTACTTATTGCTT

GAGTATTATAGATATCAATGTGTTCAATTATTATGGTCCGAAGCATTTACAAGATATAGATTCTTCAATGTCAACCTGAA

TACAGTATTTCATACAATTGCATTCACAACGACGATCGTTGTCGCAGTGTTTCGGTATTGTGCACTGAAGTTTCCAATTC

AAGCGAATAGATTCATCTATAAATGCCAGCCAGCAATCGCAGCAAACGTTATCATTTGGATCATCATTCCTATCATCAGT

CTTCCATTGTTTTTCATATCCGAGGTTAAAATTGTCGCTCGTGACCACGTTGCTTATGATTTGCAATGCGAAATGGAAGG

TCCATTGTACGACCTCAGTTATCAAGAGAGTCCACTTCTCGTGTCCGCAGTTTTCTGGGCATTTGGTATTGTGTTCAAAC

TACTTCCGTCGTTAATTCTCTCAATACTACTCATTGCTCTTATAAGATCTTTGAAAAGCGTAGAACGAAGGCGTAAAAAC

TGGAAGCGTACACAAGGTGCCAACATTTGCACCAACTCCGAGAGAAAAGCAAAAAGAAAGTTGACGACCAGACCGCGCAC

AACACGAATGCTCGTCATCATTCTACTCCTATGTGTAATGGTAGAACTACCCATGGGTATTCTTAACTTGTGCGTGGCTA

TCTACGGTGAAGAGTTCGGAAATCGGTATTACGATCCAGTGGGAAATCTGATGGAAATGCTGACGCTGTTGTACAGCTCT

GTTAGCTTCGTTCTCTATTGCACCATGTCCAATGAGTATCTCTCCACGTTCCGAGCACTATTCTTCCCATGGACCAGAAA

GAATAGTTTAAGAGGAACTCGAAGAAGTTGGAATCATCGACATGACGACGAAACAAAGTCCCCACGAACTTTTTTGATAA

ATAGAACTGCGCCAAGCTCATATGTCGGGTCTTAA

>pR34

ATGTTGCAAGCTTGCCTAAACACCACCGAAGACCAATGTGATTGCCTTGCATTCAATTGTCCAATTGTTTATAGTCATTC

GGAAAGCGAAAAAGAAGCGTGTTACATGGAGCACTGCTTTATTTCAAAACGAGCACTGGATGACGTCACGTTGTATAAGG

TGACTGCTCTTTACATTTTCATTTTCTTAGTTGGTGTAATTGGAAACACTACAACCTGCTTAGTCATGAAAAAACATCCC

ATGATGAAAACCCATGCAAGCATGTATCTCATGAATCTGGCGGTTTCGGACTTGGTCACGTTATGCGTGGGTTTACCGTT

TGAAGTAATGATGAACTGGAATCAGTACCCATGGCCATTTCCGGATTACATATGCAACTTGAAAGCGCTCATTGCGGAAA

CAACGAGTTCCGTTTCTATTCTGACAATTCTGATTTTTGCAATTGAACGTTATGTAGCGGTGTGTCATCCACTTTTTCTA

ATGAAGGTTCAACCATTCAAGAGAAATATTGGAACTATAATTGGCTTTACTTGGATTTTCTCTATCCTTTGTGCTATGCC

CTTTGCGATCCATCACCGAGCCGATTACATTATGAAAAGCTGGCCAGGGACAGACAACAGAATACCGGTTAAATCTTCAA

AAATGTGCATGATAGCAGTGATGTTTGAACCAAAGCTAGCGTCAACTTTTAAGATTCTATTTCACTTCTCTGCCATAGCA

TTCTTTGCACTCCCACTGTTTACAATTGTAATTCTCTATGCAAGAATTGCATGTAAGGTATCCAGCAACAGAACAATTCA

ACCAGGCGAACTTGATATCACTGAGGAACTGCAAATGAGAATCAATGCAATTTTATGTGCAATCGTTTCGGCTTTCTTCA

TCTGCTACCTTCCGTTTCAATTGCAACGTCTCTTGTTTTTCTATTTTGATAATGAAGTTATTTTGACATGGGTCAATCAG

TATATGTATTTTATCTCAGGATTCCTTTTCTATCTTGCCACTATCATCAATCCTATTGCCTATAACCTTGCATCCAGCCG

TTTTCGAAGAGCATTCAAAGACATTCTTATTGATTACTGTTGGAGAGGAGGATCTGAGCGTTATCCAAGAAGCTCATTCA

GCAAATATAGCTTAGCTCATACTCCTCTCAGGCAAGCCATGTCCAACCGGGTTCCAATTCTCGACTCCAAAACAAACGCA

TAA

>pR35

ATGGGGGAACAAGAGGAGGAAGGACCTATATGTATTGAATCTACATTTCTTACAATGCGAGTGATGGAAAATTATTTGAA

ATCTTTTGAATATTTTTTGCTGGTTTCTTGCGTCTGTTCTACTTTTTTACAAATATACGTATTGCTGAAAGCGGTGAAGT

ATATACGCAGGGCGACTGGCGATGAATGCCTCCACGTTTTCCTGCTCAGCATGACCATGGGCGATCTGTTGCTAACTAGT

TTCTGTTATCCAATCGAGCAGCTTCGCGAACAGGAAATTATCAAACCGCCACAGTGGATTAACGTCGCTCAGCATTTTCT

AACATGGGTCGGATTGTCTGCATCATCGGCCAGTTTGATTCTGCTCAATGCGGATAAATTGCTATATTTCAAGTTTCCAT

TGAGATATGCCAACTGGGTCACTTCATTCAAAGGTGTCTCCCTTGCTGTTTTCGTTTGGTTCGGATGCTTTGTCTTTGTG

TTCATTTGCTGGTATCTGGAATGTTTCACATGTGAAGATGACTGCCGGCAACTAATGATTCTTCCCAACAAAGTTGTAAT

GTATATCGTGTTCACAGTTTCTGCGTGCATTGCTCCAAGCTTAACATCACTTGGAGTAGCAATTTATATTTTAAACGTAG

TGACGTCGCATAGAACAAAATTATCAGACGAGAATGGGAATTCTTCTCACGCTCTTGCCACCCGCCTTCGAACTTTTTAT

TTCATATTTATGACCACAATCTTTACTGTTGGCACATTACTTCCTTACCGAATTTATAATATTCAACGTCAGCTGACCCC

GAGGGAAACTGATGAGATAAGCTGCGGAAGTATTATCTTTTCATGGACTTGTCTCTACTTTGTCTCGCTCAATGCGATCC

TGAATCCAATAATAACGGTCACGGTTCTCCCACAATACCGTTTTCAATGGCTTTGTAAACGGCTAGGAAGTAGCTCTTCT

CCAGCCGCCGTCTATGTTTAA

>pR36

ATGACAACGTGTCCCCTACCACCCAGTTTAGACGAAATGGATCTGCGATTAGCTGCCGATAAAGTTCTAAACGGTTCACT

GATCAACTGTACATTCCAATCGTTTTATGATCAAATGTATCAAACACACGGAGTATACTTCATATTTGAACCAACTCCGT

TCGTTCATCCAATTGTTTCACAGATTTTCTATGGAATCCTGTTCACACTAACAATATTTCTGGCGTTAATGGGCAATTTT

ACAGTGATGTGGATAATCCTGTACCACCGTCAAATGCGAAGCGTCACAAATTACTATCTGTTCAACTTGGCAGTGGCGGA

TGCCTCGATTTCAGTGTTCAACACGGGATTCTCATGGTCTTATAATTATTATTATGTTTGGAAATTCGGAAGCTTTTACT

GTCGAATAAACAATTTGATGGGAATAACTCCGATTTGTGCAAGTGTATTCACAATGATTGTCATGAGCATTGAAAGATAT

TATGCCATAATCCATCCATTGAAAAAGCGTCCAGGACGACGATCAACTGTTACCATCATCATAATGATTTGGTTCATGGC

ATTTTTGTTCGGGGTTCCAGCATTTCTTGCGTCAAAGGTTGATGTCTATTACTTCTACGATGGTTACACATTATACGAGA

ATCCACTGTGCCTTGCAGACAATTATCCCGGTGGAAATGAATCACTACTTGGACAGGTATACAACAACGGACTGATAACT

GTTCAATACATTCTTCCACTATGCATTTTATCAGCTGCTTATTATCGAGTTGGTGTTGAGCTGAGAAAGGATAAAACCGT

CGGTGACGTGAGACATGCAAAATCAGTGGCCGCAAAGAAGAAGGCGTCAATCATGTTGGCAGTGGTTGTGTTCATTTTCA

TGATTGTCTGGTTCCCATACAATGCCTACTATCTCACATTGCATTTAGTTGAGCCAATTGGGAATAAAATGTTGAGTCTG

TACATTTATATCAACATCTATTGGTTGGGAATGTCGTCAACTGTCTTCAATCCTGTCATCTATTATTTTATGAACAAACG

ATTCCGTGTCGGATTCCATCACGCATTCCGTTGGCTTCCATTCGTTCGTTCCGATAAAGATGAATATCAAACAATTCTTT

CACAGACACGTCCATCTCTGATGCCACCCACAACGATGGCGCATACGGATTTCTGA

>pR37

ATGAATACGTCATTTGTAGAGCCATTATATGCAGATGTTGAACAACTTGAACCAGTCCCTCTACTTAGACATTCATATCA

ACTAACTGTTCTTTATACAGTTGCATATGGAGCTGTATTCTTCACTGGTGTACTTGGAAACACGTTTGTTGTCTTAGCGG

TTTGGGCTCATAAGAATTTAAATATAACCACGGATTATCTAATTTTGTCTCTTGCACTTGCTGATCTGTTTATTTTATGG

ATTTGTCTGCCAACTACGTTGATTAATAGCATTTTCACAGAATGGCTTTGGGGTCAATTTTTCTGCCGATTGTCCACATG

GGCTAACGCATCTACGTCATTTGCATCAGTTTACACCTTGGTTGCAGTGACGGCTGATCGTTATCTAGCTATTTGCCATA

CGTTGAAATACAACACTAGCTGGGATCGAGAATATACAAAATATGTTATATTTGCTGTTTGGCTAGTAGCTGCAATATTC

GGAATACCTAATTGGTATAACTATGATTTGATAGTATGGCAAGAAGGCAGTTATGGTTACCGATTATGTACGTCACAAAC

GGATCAAAAATTATATTTTTTATTTGTTAACTTATTGCTGGCTTTCATAGTTCCATTTGGTTTGATTTCGGGTCTATACA

CGAGAATATTTATCACTGTATCAACACATAGAAGTCTGGCAGTTGATGCAAGAGCCCGAGAAGATCGAGTAAAACTACGA

GTTGCCACAATGATGTTAACAGTAATAATTGTATTTGCTTGTTGCTGGTTACCTCTCTATTGCATCTTCACATATTTTTT

CTTCTTTGCTGATCAGCGATCGGATCTTTTTCAAATTACTTCAATGTTGATTCGGCCAATATTCCAATGGATGTCATTAC

TATCGAGTTCTTTAAATCCAATAATTTATATTGCATACAGTCACAAGTACAGAAGAGCTTTCAAAAGTATATTACTGATG

CCATGTAAGACAAGATACGAAAGAGTCCGAAGCACAATACTCCGTCGTCACTCTCGTGGCTTCAAATCAACTGCAACAAT

TTCAATGTCTAATTTTGGAACTGAACCAACAAATTTAGGTGGAGCATCTAGTTTACTTATTGAACTAGATGGAAAACAGG

TGGAAAGATCTACTTCTGATTGTTAA

>pR38

ATGTCTGTCACAATGGCTCTACAATCCCTTATAAAAAGTCCAACTCTAGACGACGAGCAAGAATATCTCGAAGAATGGAC

GAACAGCTCTCGACTTCTCTTCTACTCTGTCATTTCCATTTCTCTACTAACTCTCCCAATTCTTCTCTTGACCTTCTGCT

ACATTCTGTGCAGAAGCAAACGAAATGCGCACTTTTTGCCGTACTTGTGCTCAATATTGGCCGCTAATTTCGTTCTGCTG

TCTACAATTTTTCTGAGTGTGCTTGCAAAGAATACGGATTTGGTCTATGACACCATTCCCGGCTTCCTTGTCTGCAAAAT

CTCGACATTTCTCGTCAACTCCTCCTCCTGCTTCATCTACTGGACATGGGTCGCCATGTTTGCAGAAAGGTGCTGTAACA

TATTCTTCCCACTTCGGTTTCGAACTTCGAGCTCATTCAAAACCGCCGGAGTTTTGTGCTCAATTTTGATATTTTCAATG

AGTATTCAGTTGTGGACTCCTATTTTTATAACGGAAAAGCGACTTGATAATCATATGGATTATATTTATTGTGGGGAAGA

TCCAAGATATTCAAGCCAAACCCGAATAATAATTGTGCTGGAATGCCTAACTACCTTCTTCCTACCTCTAATCCTTACGA

TTTTCGCCGATATTTCGGTTTTAACATGGAAAAGTTCGTTTGGAATTGACATTGATTTGGTGTCCCGTGAAAAGATTAGC

GGAAAAAATAGTGAGACAATGAAAATTGTGTCAACTAATAGTTTGAAGAATTCAAAAAAGCGCCGGTCAAATGCAATCCG

TCAGTGCTTGATTTCATCGACAATCACGCTTTTTCTAAATCTTCCGAATTATAGCTTACAGCTCCTCGACGAGTTTTTAA

ACTTTCGCGACAGCAATTCCATTCAAGCGCGAAGAATATTTTTAAGGATAGACGCATTTGTTTACGTGCTGTATTTGATG

CAATTTCCCATAACTCCACTTCGAATGTTCACTTTGTCGAAATCTCATACCAGACGGGGATCTCGTCATCGAAGGAATAC

GCTTATTGCTTAA

>pR39

ATGAGTTCTTCGAATCACTGCATCGACATCCGTGCATACTTGTGGCAGACAAAGCATGACCTGACGCTCCACCCGATTCC

CATCGCGATTCTTGCAACCATCTACACTATAATTGTCGTAGTTGGCGTAACCGGCAATTTGTTAGTAGTGATGTCGGTGA

TGAGGTTCAAAGTTCTTCAATCAGTCAGGAACATGTTCATCGTATCTTTGTCAGTTTCTGACATTTTTGTGGCGATTGTT

AGTGGTTCAGTAACGCCGATAACCGCATTCTCTAAAGTTTGGTTATTTGGTGGACCATTGTGTCATTTACTACCTTTGTT

ACAGGGTACCGCGTTGAGTTTTTCCACGTTAACGCTCACCGCAATTGCAATTGACAGATATATTCTCATCTGTCATCCGA

CGAAAGAACCGATACGCAAAGATCAAGCATTGAAAATGATAAGTTTCAACAGCGCCATCTCAGTTGGGCTTTCGGTACCA

TTATTCATGAAACAGGAACTTATGCAATTCCGAAACTATTGCGGAGAATATTGCTCAGAAAACTGGGGACCAGATGCTTA

TTTGAGAAGCGTTTATGGAACAGTGGTGTTCATTATTCAATTCGTGTTTCCATTGATCACCATCACATTTTGCTATGCAT

CTATTTCTATCAAACTACGACGTGGTGTCTTTGTGAGAGGAAGCCAAAAAGAGCTGATGTCTGAGGCACGTCGTCAATTG

ACGCAACGTCGACTTCGCACAAATCGGATGCTTATTATCATGACAGTCACATTCGCTCTCTCATGGCTGCCATCTGTTGG

CTTCAACTTTCTCCGGGATTACTCAGCGCTTCCCGGCATTATTGATTCACAAGATTACCTATTCGGAATTATTTTCCATT

GCATTTCAATGACATCGGTGATTGTGAATCCCTTCCTTTACGGTTACTGTAATGAACACTTTCGTGCTGCATTTGCGGCT

CTTCTTGACACGGTGAAGGCAGCTTGTGGAATGAGACGAGGCAATCCAGCGTGCTCCCAGCTACTCAGTACTCACTTTGA

AAGCACCACAAGACGATCCGTGACTACCACGATTCCAAGTTCAATTTAA

>pR40

ATGATGGATTCGATGGACTTTCACAGAATAATGGCTTTCGTGTATTTACCGACTATTCTGATTGGGCTTCTTGGTAATTT

ACTATCACTTTACGCATACAGTCGTAAAAATAGAAAATCCATGGTCGGTTTCCTGCTGTACTCACTGTCCGTCTCTGACA

TCTTCCTGCTAGTCTTCGCTCTTCCACTTTTCAGCATCATGTATCTACCCATTTGGACTGACGAGCAGAGAAGTTTCGTC

GTCGCGTACACTGCTAAATATGTATATCCGTTGTGTATGATGGCAAAAACATGCAGTCTTTATATTATGGTATTGATTAC

TATTGAGCGATGGATTGCTGTTTGTCGACCACTTGAGGTCCAAATCTGGTGTCGATACACCACATCATCATACTCCATCG

CCGCCATCATCACTTTCGCAGTGGTTCTGAATTTTGCTCGATTCTTTGAATTTGAAATTGAATACATCGATGGTTTGGCA

TTCTTCAGACGGGATCTTCTGGATTCTGAGAAACATTGGTGGTACTTCATGTTTTATTTCATCATCATTTCTATTATATT

CGATTATCTCGTTCCATTTGTGATCATGTTTGTTGCGAATATGTTGATTATCAGCGAGCTGAGAAGAACGAAAAAGGAAA

GAAGTTTGATGACAATTCAACAACAAAAAGAGCAAAATACGACAGTAATGCTTCTGGTTATCACAATTTTCTTTGGATTC

TGCCACTTCTTCTCAATGGCCCTAAAGCTTGCGGAGAGTTTTGTTGGAAAGTTTCTAACAATTCAGAATATTTATCTAGA

AATGTTTGGAGAAATTTTTAACTACCTCATTATCATTCACACCGCCTCAACATTTTTCATCTATTACATGTTTTCGGAGA

AGTTCCGGCAAATAATCAAAGGAATTTGGAGGCCCGATCAATACCGACACGGTAGCCTTCCTGATGGAACTCTGAACATT

ACTGACAGGTATCAAAAAATTTTGTGA

>pR41

ATGAACCTTACTGCACTCGACGAGCTTGTTGAGCTTCGTGCCATGGCTGAAGCTGAACTTCTGTGTGCTCATTATGAAAC

ATACACACCCGTTCGTTTTGTGCTCATCATGGTTGCAACGGTAATCGCGTCGCTTGGAACAATTGGAAATATTCTTTTGC

TGATTGTTTTCTCAGCTAAACAGATATCAAATACTCCAGCAACGCTTTACCCGTCCGTTTTAGCTTGTTTGGATTTTGCA

ATATGCTTCGAATATATCCTTCTGTTCGGCGTTGATGCTCTTGTGAGCTACTTGAAAATTGAGAGCTTGTTCTCTCTATA

TTATGTGTACATTGTTCCCGCCTACGTGATGGCTCGAATCACTCAGCTGGCTATCCCTTACATGCTCATTTTCGCCACAC

TTGAACGCCTCTTCTGGACGTCTAAGAACAAAAGCAACTTATTGAAGGCATTTCATTCAACTACAGGAAGACATATTACT

GTCATTGTATCACTTATTATGTGCATTTTGTTAAGATCTCCGAGTGCTTTTGCTATTAGAGTTGATGAGTATCCGAAATG

TCCTGACTTTTTTAGAACAAAAACTACAAATCCAAGAGAATGGGCACTTGAAAGCCAGCTCTACCATTTCTTTGATTTTC

AAGTAATGACAATTGCTCAAACACTTGTTCCATTTGTTTTGCTAGTTGGATTAAACTTGATTATTGTTCGAAGAATGTGC

TCAGATAGTGTTCAAAAGGAAAAACTACCAGAACCTACGGAAGTTTGTTTCAAGGAAGATCGGCTACTTACTCAAATCAC

TCCAGTTCATAAGTCATCTACAACAATTGGTCTATCGTTCCCGCCTCTCAAAAGCATGTCACCAGCGGTTAGAAGTGCTG

TATTTACCATGGCAGCTATCGTAACATCATATCTAATCTCTAACATTCTTCATCTCAGCTTGACAGTTCTTGAAAGATCT

GGTCATCAAATACTAAAGTCAGAGGAGGATCCACAAATGTCTTCCACATTTCACACATTCTTCTCCGATTTGGTTTCATT

CGTTTACATGTTTACTTCTGCTATCCGAATTGTCATTTACTATCTGTGTAATCCAAAGATCCGTGGAGATCTTATCGATT

TCTTTGCCAATCGTAAAAATGCCGTTTATCTTTAA

>pR42

ATGATTCTGGAGTGGGAAGGCAGCGAGAACTCGACAGAATGTTTATGCTCCGATATTCAGAGAGACGATTATTCACCGTT

TTTCGAATGGGCCAACTACCTGACGATCATCCTTGCCCTCCCCGTCCTTTCAGTTTTCGGAGTGCTCACAAACATAATCA

ATGTATTTTTATACACAAGAAAACGATTACAAAACTCCGCAAACACGTACCTATTATTTCTGGCGTGCAGTGACTTTATG

GTCATTGTGACCGGTCTCTTCATATTCTGGATCGACTCTGCCCGATCGTACATTCCAGAGCTCACTCAGGCTCCCTACAC

CACTGTTTACACACTTCCATTTGGATATATGGCACAAACTTGCAGTATCTATTTCACAGTGGCTGCCGCTGTCGATTGTT

TTGTGAATGTATGCTGGGCAAAACAAGCAAAACATTATTGCACCGTTCGTCGAGCCAAACAAATATGCATTTCAATAGTT

GTTGTTTCAATTATGTACAACTCATTACGCTTTCCGCAGTTCAATTTACGAAAGTGTTTTAATGATATTACAAAAGAACA

AGTCATCGAGATTTGCCCAACATCACTTTTTGTCACAATTAATTCAGTTTACAATATATATATGTACATGGTCCTAATGA

CACTTTTACCATTTTTCTTCTTACTGTGTATCAATGCAATTATTGTGAAAAGGCAGTCTAAGGCAAAAACAGATGAATCG

GCACCAAAAAATGAAGGAACATCTGATGATACAATTACAATGATCATGGTTGTTATTTTATTTTTGGCTTGTAATACTTT

GGCTCTTGTAGTAAACTTTATCGAAAACTTCACTGAGCCATCGCCGGTTCTGCTAAACTTTTTGTCAGACACCAGTAACT

TCTTAGTTGTGTTCAACTCTTCAGTCAATTGCATTATCTATTTTATTTTTAACACAGACTACAGAGATGTTTTCCTAATA

TACTGGAAGAAGCTGAAAAGAGTTTTAATCGAGGAGTATTGCTGTTGTTGTGTACCAGCCGGCTCACGTAACTACTCAGC

TTATCAGCCAGTTTCCACAAGACTCGTTATCGAAAAGAGTCAAAATGGAAGCAGTAGTACAAAATTGATTCTACCATCAC

TTGATCTGAATAGAAGTGATCAAGCCAGTTTTGCAGAATCTGAAAGTGCAGCATCCCCAATCTGGCAACCCTTAACCACG

AGAAACCTTCCAAATGTTGATTGGCCAAACATCGATGATGACATAGATTCCGGATGGGATGATGGAAGTCAGATGAAAAT

GCAGATCTCCTCGCGTACACCAAAACGTTGGCTTGCCGAGGTGAACATTGTAGATGTTGAATCATCGAATGGAAGACATC

CAAGAATTTATGTTCGTCCTTTGAATCAATTGCCAAATAATGAAACAATCTCTATCACTGCTCTTTAG

>pR43

ATGATTCTGGAGTGGGAAGGCAGCGAGAACTCGACAGAATGTTTATGCTCCGATATTCAGAGAGACGATTATTCACCGTT

TTTCGAATGGGCCAACTACCTGACGATCATCCTTGCCCTCCCCGTCCTTTCAGTTTTCGGAGTGCTCACAAACATAATCA

ATGTATTTTTATACACAAGAAAACGATTACAAAACTCCGCAAACACGTACCTATTATTTCTGGCGTGCAGTGACTTTATG

GTCATTGTGACCGGTCTCTTCATATTCTGGATCGACTCTGCCCGATCGTACATTCCAGAGCTCACTCAGGCTCCCTACAC

CACTGTTTACACACTTCCATTTGGATATATGGCACAAACTTGCAGTATCTATTTCACAGTGGCTGCCGCTGTCGATTGTT

TTGTGAATGTATGCTGGGCAAAACAAGCAAAACATTATTGCACCGTTCGTCGAGCCAAACAAATATGCATTTCAATAGTT

GTTGTTTCAATTATGTACAACTCATTACGCTTTCCGCAGTTCAATTTACGAAAGTGTTTTAATGATATTACAAAAGAACA

AGTCATCGAGATTTGCCCAACATCACTTTTTGTCACAATTAATTCAGTTTACAATATATATATGTACATGGTCCTAATGA

CACTTTTACCATTTTTCTTCTTACTGTGTATCAATGCAATTATTGTGAAAAGGCAGTCTAAGGCAAAAACAGATGAATCG

GCACCAAAAAATGAAGGAACATCTGATGATACAATTACAATGATCATGGTTGTTATTTTATTTTTGGCTTGTAATACTTT

GGCTCTTGTAGTAAACTTTATCGAAAACTTCACTGAGCCATCGCCGGTTCTGCTAAACTTTTTGTCAGACACCAGTAACT

TCTTAGTTGTGTTCAACTCTTCAGTCAATTGCATTATCTATTTTATTTTTAACACAGACTACAGAGATGTTTTCCTGTAA

GACTTTTCATTTATTTTTTCTCTTTCAAATCAAAATATTTTCAGAATATACTGGAAGAAGCTGAAAAGAGTTTTAATCGA

GGAGTATTGCTGTTGTTGTGTACCAGCCGGCTCACGTAACTACTCAGCTTATCAGCCAGTTTCCACAAGACTCGTTATCG

AAAAGAGTCAAAATGGAAGCAGTAGTACAAAATTGATTCTACCATCACTTGATCTGAATAGAAGTGATCAAGCCAGTTTT

GCAGAATCTGAAAGTGCAGCATCCCCAATCTGGCAACCCTTAACCACGAGAAACCTTCCAAATGTTGATTGGCCAAACAT

CGATGATGACATAGATTCCGGATGGGATGATGGAAGTCAGATGAAAATGCAGATCTCCTCGCGTACACCAAAACGTTGGC

TTGCCGAGGTGAACATTGTAGATGTTGAATCATCGAATGGAAGACATCCAAGAATTTATGTTCGTCCTTTGAATCAATTG

CCAAATAATGAAACAATCTCTATCACTGCTCTTTAG

>pR44

ATGAACCGGACGGCTACTATACTCGAAGGAATGATACCATTGATACCAGACATCATGATTCTGGAGTGGGAAGGCAGCGA

GAACTCGACAGAATGTTTATGCTCCGATATTCAGAGAGACGATTATTCACCGTTTTTCGAATGGGCCAACTACCTGACGA

TCATCCTTGCCCTCCCCGTCCTTTCAGTTTTCGGAGTGCTCACAAACATAATCAATGTATTTTTATACACAAGAAAACGA

TTACAAAACTCCGCAAACACGTACCTATTATTTCTGGCGTGCAGTGACTTTATGGTCATTGTGACCGGTCTCTTCATATT

CTGGATCGACTCTGCCCGATCGTACATTCCAGAGCTCACTCAGGCTCCCTACACCACTGTTTACACACTTCCATTTGGAT

ATATGGCACAAACTTGCAGTATCTATTTCACAGTGGCTGCCGCTGTCGATTGTTTTGTGAATGTATGCTGGGCAAAACAA

GCAAAACATTATTGCACCGTTCGTCGAGCCAAACAAATATGCATTTCAATAGTTGTTGTTTCAATTATGTACAACTCATT

ACGCTTTCCGCAGTTCAATTTACGAAAGTGTTTTAATGATATTACAAAAGAACAAGTCATCGAGATTTGCCCAACATCAC

TTTTTGTCACAATTAATTCAGTTTACAATATATATATGTACATGGTCCTAATGACACTTTTACCATTTTTCTTCTTACTG

TGTATCAATGCAATTATTGTGAAAAGGCAGTCTAAGGCAAAAACAGATGAATCGGCACCAAAAAATGAAGGAACATCTGA

TGATACAATTACAATGATCATGGTTGTTATTTTATTTTTGGCTTGTAATACTTTGGCTCTTGTAGTAAACTTTATCGAAA

ACTTCACTGAGCCATCGCCGGTTCTGCTAAACTTTTTGTCAGACACCAGTAACTTCTTAGTTGTGTTCAACTCTTCAGTC

AATTGCATTATCTATTTTATTTTTAACACAGACTACAGAGATGTTTTCCTAATATACTGGAAGAAGCTGAAAAGAGTTTT

AATCGAGGAGTATTGCTGTTGTTGTGTACCAGCCGGCTCACGTAACTACTCAGCTTATCAGCCAGTTTCCACAAGACTCG

TTATCGAAAAGAGTCAAAATGGAAGCAGTAGTACAAAATTGATTCTACCATCACTTGATCTGAATAGAAGTGATCAAGCC

AGTTTTGCAGAATCTGAAAGTGCAGCATCCCCAATCTGGCAACCCTTAACCACGAGAAACCTTCCAAATGTTGATTGGCC

AAACATCGATGATGACATAGATTCCGGATGGGATGATGGAAGTCAGATCTCCTCGCGTACACCAAAACGTTGGCTTGCCG

AGGTGAACATTGTAGATGTTGAATCATCGAATGGAAGACATCCAAGAATTTATGTTCGTCCTTTGAATCAATTGCCAAAT

AATGAAACAATCTCTATCACTGCTCTTTAG

>pR45

ATGGAAGTGAAAGATATAGATAACTACTGTGATCGTGGAATCAGTCCGAATGCATCCAATTATCTCACGTACCCATTTGA

CGGGCTCTGTCTACAGAAATTTTTTTATCAACTCCAAACTTCTTTGCGAAGGTTCACTCCTTACGAAGAAATCATTTACA

CAACAGTTTACATCATTATCTCTGTAGCAGCTGTTATTGGAAATGGATTGGTGATAATGGCTGTAGTACGGAAAAAGACA

ATGAGAACAAACAGAAATGTTTTGATTTTAAATCTCGCGCTTTCAAACTTGATACTCGCCATCACCAACATCCCATTTCT

ATGGCTTCCGTCAATTGATTTCGAATTTCCGTACTCTCGATTTTTCTGCAAATTTGCCAATGTGCTTCCGGGTAGTAATA

TCTACTGCTCAACTTTAACCATCTCGGTGATGGCAATTGATAGATATTATTCGGTGAAGAAATTGAAAATTGCATCAAAT

CGTAAACAATGTTTCCATGCTGTTTTGGTTTCATTGGCTATTTGGATTGTGTCATTCATCCTCTCGTTACCTCTGCTCTT

GTACTATGAAACCTCAATGCTCTACGTCATGAGAGAAATTCGAGTTGTTGATCAAAGCGGTCAGGAAGTCATCCGAAGCT

ACGGATGGAGACAGTGCCGTCTAGTTTCTGCCGGACGATTACCGGACATCACCCAAAGCATCCAGTTGCTCATGTCTATT

CTTCAAGTCGCTTTCCTATACATCGTTCCACTCTTTGTTCTTTCAATCTTCAACGTGAAACTCACCCGGTTTTTAAAAAC

AAATGCCAACAAAATGAGCAAAACTCGTGCTCCGCCAAAACGATTTGACAGATCCGATAGCCACCATAATTCGTTGAAAA

ATAACAACAACCACACGTCTTCTCTACGTTCGCCATCAATGCCCTCAATCAGAAGTTCGATAACGGAGAGAAACAAGACG

AATCAGAGAACAAACAGAACTACTTCGTTACTGATTGCAATGGCCGGAAGCTATGCGGCTCTTTGGTTTCCATTCACTCT

CATCACTTTTTTGATAGACTTCGAGTTGATCATCAATCAAGACTACGTGAACTTGGTAGAACGAATTGACCAAACCTGCA

AAATGGTATCCATGCTGTCCATTTGTGTGAACCCATTTCTCTACGGATTCCTAAATACCAATTTTCGACACGAATTTTCC

GACATCTACTACCGATACATTCGCTGTGAAACAAAGAGTCAGCCAGCCGGCCGATTCCATCATGATGTATCATCAATCGC

TCACCATAGACAAGACTCTGTTTACAATGATGAGGCAACACTTTTGACTACAGGGCGTCAGAGTAATGGTAAAGATGGGA

GCTCGTCTCCAATAGGATTCCGCTCTAGCGTCCGTGTTTGCTCGGGTCAAACAAAAATGATTGGAGATCGCATTGTATTG

GACGATGATATCGAGAAAGATAGTTTTGTCTAA

>pR46

ATGGACTTCATCAATGAGCTGTTTTTCACCGATACCGATATAGAGGATCTGGGAATGATTATGGAAGTGAAAGATATAGA

TAACTACTGTGATCGTGGAATCAGTCCGAATGCATCCAATTATCTCACGTACCCATTTGACGGGCTCTGTCTACAGAAAT

TTTTTTATCAACTCCAAACTTCTTTGCGAAGGTTCACTCCTTACGAAGAAATCATTTACACAACAGTTTACATCATTATC

TCTGTAGCAGCTGTTATTGGAAATGGATTGGTGATAATGGCTGTAGTACGGAAAAAGACAATGAGAACAAACAGAAATGT

TTTGATTTTAAATCTCGCGCTTTCAAACTTGATACTCGCCATCACCAACATCCCATTTCTATGGCTTCCGTCAATTGATT

TCGAATTTCCGTACTCTCGATTTTTCTGCAAATTTGCCAATGTGCTTCCGGGTAGTAATATCTACTGCTCAACTTTAACC

ATCTCGGTGATGGCAATTGATAGATATTATTCGGTGAAGAAATTGAAAATTGCATCAAATCGTAAACAATGTTTCCATGC

TGTTTTGGTTTCATTGGCTATTTGGATTGTGTCATTCATCCTCTCGTTACCTCTGCTCTTGTACTATGAAACCTCAATGC

TCTACGTCATGAGAGAAATTCGAGTTGTTGATCAAAGCGGTCAGGAAGTCATCCGAAGCTACGGATGGAGACAGTGCCGT

CTAGTTTCTGCCGGACGATTACCGGACATCACCCAAAGCATCCAGTTGCTCATGTCTATTCTTCAAGTCGCTTTCCTATA

CATCGTTCCACTCTTTGTTCTTTCAATCTTCAACGTGAAACTCACCCGGTTTTTAAAAACAAATGCCAACAAAATGAGCA

AAACTCGTGCTCCGCCAAAACGATTTGACAGATCCGATAGCCACCATAATTCGTTGAAAAATAACAACAACCACACGTCT

TCTCTACGTTCGCCATCAATGCCCTCAATCAGAAGTTCGATAACGGAGAGAAACAAGACGAATCAGAGAACAAACAGAAC

TACTTCGTTACTGATTGCAATGGCCGGAAGCTATGCGGCTCTTTGGTTTCCATTCACTCTCATCACTTTTTTGATAGACT

TCGAGTTGATCATCAATCAAGACTACGTGAACTTGGTAGAACGAATTGACCAAACCTGCAAAATGGTATCCATGCTGTCC

ATTTGTGTGAACCCATTTCTCTACGGATTCCTAAATACCAATTTTCGACACGAATTTTCCGACATCTACTACCGATACAT

TCGCTGTGAAACAAAGAGTCAGCCAGCCGGCCGATTCCATCATGATGTATCATCAATCGCTCACCATAGACAAGACTCTG

TTTACAATGATGAGGCAACACTTTTGACTACAGGGCGTCAGAGTAATGGTAAAGATGGGAGCTCGTCTCCAATAGGATTC

CGCTCTAGCGTCCGTGTTTGCTCGGGTCAAACAAAAATGATTGGAGATCGCATTGTATTGGACGATGATATCGAGAAAGA

TAGTTTTGTCTAA

>pR47

ATGTGTCAAGATGAAGGAGGAACACTGTTCGACCCGCAGGATCCCGCAGTTTCTGGCCTCATCGATGCTCTCGAGCAATT

TCAGAGCTTTTACACATATTTCCATAGATACGCCTGTCTTTTTATATGTATTGTTGGCGTTTTCAGCAATGCGATTCATA

TTGCGGTACTATCTCGTCCCCGAATGCGAAGATGTGCTGTGAACAGTGTGCTCACCGCCGTAGCCTTTTGTGACGTCATC

ACCATGACGTCATACAGTATTTATTTGATGCGATTTCGATTTTATGAGACCGACCATGGCTACAGCTACATTTGGCTGGT

ATTTCTGAAATTTCACGTGTGGTCCTCGATGACATTACATGCTATTACGCTTTATATGGGTGGTGCATTGGCATTTATAC

GGTGGCAAGCACTAGGAAATATTCACAGCAAATGGCTGCAGCCAAGGAATTCTTGGCAACTATTCGGCGTTGTTAGCGTT

GTGTTATCAATTGTCTGCCTCCCTACACTTGTTCTACACAAAATATATGAAATAGAGAGTCCGGAAATCGAGACCTCAAC

CGTTTCCGAAGTTTTGAGGCTTTCCGGTCAGAGTATACCAAAAGAAGTTCGATATAGTTTGAATTTTTCTACGTATTCGT

GCGCGTTTTTCAAGTTTAACTTGTGGATGTTGGCGATTGTGTTGAAGGCAATCCCCTGCGCACTTCTCCTCTGGTTTACC

ATTGCTCTGGTGGTAAAACTCCGTCAAACAGATGAGAAACGAAATTATTTATACTCGAAAAGCTTCCGAAAACACGTCAA

GAAAACTACAGTACCCGACCGTACCACCTATATGCTCATCATTATGCTTGTGGTTTTTTTGGTGACAGAGCTTCCGCAGG

GGTTTTTGGCACTTCTGAATGGCCTGTACACAGGAGATGTTAATATTTATATTTATAAGAATTTATCCGAATTGTTGGAT

TTTCTATCGCTGATTAATTGCAGCGTGGATTTCCTGCTCTATTGTGTGATGAGCTCCCGGTACCGTCAAACATTCGGTCA

TATGCTGATACGTGTCGAATCGTGGCTTCGCAATCACGGAGAACGTCGTCGACTTGCGCGGGAAATCAAAAAGAAATTGC

CGGCGCCCGTTGGGGTCTAA

>pR48

ATGGATCAAATAACAAGCACAATGATGTTAGATTTAATGGAAGAAGAGGAAGAAGAAATTTGTGGATTATATGAAGGATA

CAGTACTCAAAGATTTGTTATTATATCCGGATGTACTACGGTTGCACTTTTTGGCGTTTTAGCTAATATGTTACTGATGG

CAGTCTTCCGTCGATCCTTACCTTCTTCCATATTTCTAGCCACACTTGCCACGTGTGACATGCTGATATGTCTGACGTAC

ACATTATTATTTGGTGTAGATGCTGGAATTTGGTATCGAAAAAATACGACATTATTCTTCCTCTATCACCGTTACATTGT

TCCCGTATTCTTTATTGCAAAAGTTGTTCAATTCGCCATTCCATTTATTCTAATTTTAATTACTTTCGAACGATATCTCT

GGACATGCACAGAACGAAAAAGAAAAGCCTTCTCTGCGATTTTCAACGAACGTGGACGAATTGTTACTGTGATATTTGTG

TGTTTCTTCTCGGTGGCAATTCGGATTCCTGTGCTTTATGCGATGAAGGTCAAAACGTTTCCTCTGTGTGACGACTATTT

CAGAAGTGAATCATTAGACGGAACACCTTTCGCCGCCACAGAAGCATACGAAATATACGATTTCCATGTTATCACTGCAA

TTCAGATGATGTTTCCGTTTGTGGTACTATTGCTACTGAATCTAACAATTATCAAGAGATTGGTAGCTGAGAAAAGGGAA

AATATGTACCCTATTCTCCGAGGTGCTGGAACTACAACAGAAGTGAAAAAAGCGTCATTTGTTCAGGGAAATCTTCCAGA

AAACTATGTGCTTCTGCAAGTTGCCGCTGACGTCATCAAGGAATCATTGATCCACAGATCATCCCGAAGTAAGCGATCAC

AGCTTCGAAATGCAATTTACACGATGTTGGCTATTGTCACTTCATATTTGGTTTGTAATGGCGTTCATCTTTTTCTGACG

ATCCTTGAGAGGTTCGATCCATCTTATCTTTACGAATCCACGGATCGAATGCAGTCGAGCACATTTTATATCGTTTTGTC

TGATACTGTCTCCATCTGCTATATGGCATCCTCGGCAATTCGTATTTTCATCTACGCCAAGTGTAATCCAAAGTTGCGAC

AGGAGATCACCGACTACATCAAACGAGAGAAGAGCATCGAGACGAACAGCTCGTAA

>pR49

ATGACACATGACAATGAATATCAATCTGCACTTGTATTCGATAGAATTAGACGATTAATCTTAAACAACTCGAATACCCC

AATAACGACAAAAGAGCTTATAAAGCATTGCTTTCATCCAGATGTTCAGTTTCTTGCTCACATGACCCTGAAATCGTTGG

ATCATCCCGGTGAACAGTTTAATAATTCATCAAATCAACTTTTATTAGATCGCGCTTTTCTAAACCCAATACTCACACTC

ATTTTTGTTTTTGTCGGGCTTGTGGGTCTCATTGGCAATTTGCTCACAGTTATTGTGATTTTCAAGACAAATTCCCTGCA

CTCGCACACAAACTATTTCTTGGCAAATCTTGCTACCAGCGACTTCTGTCTCATCGTTGTTGGAGTTTCGTTTGACTTGG

TGAATATCTGGAACGATGAGGAACCGCTGGACATTTTTGGATATTGCTCTCTTACAAGCACTTTTATATCTCTGTTCACA

TTTGCTTCAATTCTCACAATTGTTCTTTTGACGGCAGAAAGATTTACAGCGATTTGTTATCCTTTTTCCCATAGAACAAT

TTTCGACGAGAAGCGTGTTAAAAGGTTTATACTGCTTATTTGGTTCGTAGCACTTCTTCCATCAATTTTCATTGGCTCTA

TGTTCAAACGAGTGTCTCAAGACTTTTGTGGCTTCAATCGCCAAATGACTTATATCGGTCGATGTGATTTGGTGACATCT

CCAGATAGTTTCTTCCGGTACCCATTTGAATCAGCAATAACCATCACTTTTGTACTTCCCTTGTTCTTCATTATTTATTG

TTACTTCCGGATTCTAGTCACATTGAACGAGATGTCCAATTCGACTCATGTGCACACTCCAGTCGGAACTGCTCGTAGTG

ATAGTGGAGCTTTTCCTTTTCCCCATACATCTAATAACTCTAACACTCAAAGTTTCCCGCTAACAGTCCATACTAAAAAT

GTCCAACCGCCGCGAAGTCAACAAGCCCAGAAAATGGTCATTAAGATGCTTGTTACTGTAACCGCTGTATTTTTTGTCTG

CTATCTTCCATATCACGCTCAACGTCTCATCGTCAAATACAACAGCAAGGATTGTTCCAACTCAGACTTCTGCAAGCTTC

TCTACCCAATAGCAGGAATCCTCCAATACATCTCCGCTTCTTTGAATCCAATTTTCTACAACCTCATGTCTGTACGATTT

CGCAACGGATTCAAGAAGCTCATCAAAGACGTCTGGGCACATCGAGCCCGTAGTTACAGCAATCTAGCTCGTGTCTGA

>pR50

ATGTTTGACAGGCAGATGGAGGTGGATAATTCAACAATTGTTGAAACAGTGGCACCCACCTATCTGATAAGTGATTATAT

AGAAATTGCATATTTGGGACTTGTTCTGCTATTTGGAGTTCCAGCAAACGCCGTCATACTTCAAAAGTTGATTAAAGAGA

TGAAGATGTCAAATCGGGATATGGTTAAGAGTGGTTTCGTTATGCTGAAGATAAATTTGAACATAACAGATCTTCTGATC

TTGACCTACTCCCTCGGTAAACTCATCTGGCTTATCACTTATAAGTGGATCGGTGGAGATTACGCGTGCAGATTTTATCA

GATGTTCTCCATGTTCTCATTATATAGCTCTTCAAATATCGTAATGTGCATCGCACTTGATAGGTTACGGAATGTGATCT

ATGCAAACCAAATTCACACAAAAACTGATAAGATAAGTACAGTATCAATTTTGGCGTACAGTTCTTGGCTTGCTGCACTA

GTCTGTAGTTTACCACAATTCTTCCTTTTCCAAACAATCGAGGTGTATCCAAATTTTGTTCAATGCTCAGACATTTGGCA

AATTCGGAGACATAGTCAAGATGATATTGCATTTTTTGGGAAAGACTCGTTTGTACTAACTCAGACATTCGAGAACTCTT

ACAACATTGCTCATTTGCTTCTTGTATTCTGGGGCCCACTGATTGTTCTAATCGTAACGTATGCGGTGATAGCAACCAAG

CTCACGAAGTACTCGTTAAAAGCACCTGGAACCCAGAGCATGAGAAGACCTATGCCAACTGCAGCCGATGATTTACCGAA

AGAAGTTGTTGTCCACGTTAACGTCGAAGACGGTCTACTCAAAAAAGAGGGCAGTATAAAAAAGGTGGTAAGATGCTGTG

CTGAGGAGCTCATTACAAAGCAACATCGAAAGAAGTCGAATCGTAGAATGAGAGCCAAAGAATCAACAACGACAACATCA

TCAACATCTACACGAATGCCAACGTGGCGGAAACAAATGCGCAGTCGTGTGTTTCGAACCACAATGCTTGTCATTCTGAC

CCACTTCCTCTTTTGGTTCCCTTACAATGCACTCGGCTTAATGAAATACATCAACCAATCTATGTTTGAAGTTCTTAGCG

CAAATGCCAATATTTTCAAAGACTTGCAAATTCTCATCACACTCATCAATCCGTTCCTATACGGTTTCTCAACTGGAAAC

TAG

>pR51

ATGATTGAAACATCTACAAGAATATCCCAACTAGTACCAGAAAGGCTGTGGTCAGAAGTGACAGAAGCAAGCATACTTTT

TATATATGATATTGCATATATTATAGTGGGATTCATTATTGCTATCAAACTATATAATCAGAGAAAAGTCAAACGACCAA

TTCAACACAATGGGCAGTCAGGAAACCACAACTCATTTCTACTTTTCAAAACTAGCTTGTTCGCAAGTGATTGCATGATT

ATGTTTATCTATGCAGCTGTGAAAGCTCTCTGGCTATTACAATTCGAATGGAAATATGGTTCATTATGGTGCAAAATGTA

CAGATACTGGTCATCTGTGGCATTCTTTTCGAACTCGAATATAGTTTGCGGTATAGCATTGGATAGGTATCTTTCGGTGT

ACAGTAATCATATTATTGGTGTTCGTCAGTACCAAAGAACTAAACGGATGCTATATGCTGTGTGGATTATTGCATTAATC

GCCGCGCTTCCGCAGCTCATCGTCTGGGAAACGTATCGTCCCAGTTCAGAAGACTGGGAGCAATGTGTCACCGTATTTGC

CATTGATCTACACAAATTACCACTCAATTCACCAAAAAAAGATGAGATCAACTTTTGCTCCATGCTCTATGAAGGGTACC

ATCAAGCAATGGCATTCTGGTTGCCCTTAAGCATAACAATTGCTTCCTACGTTCGCATGATGAGCAGGCTTATTCCATTT

TGGCCGTTCACAGTGCTCAACCACTATGAAGACGAGCGACAACAAACACTATGTACGATTTTCTGGAGCAAAATTAGTGA

GAAAATACGAAGCTTCTTCTGCGTAACCATTTTTCGACGGAAAAGCTATGCATCTCAGCAATCAGTGAGAGTTCCATTGG

CTGAAACAATACGTCATCCACCAACTACAGCTCTTCGACGACAACTAGGCACAACAGTATTTAAAAATGCTTGTGCAATA

ATTGTGACCCATATTATTCTTTGGCTCCCGTATAATATTATCAGTTTGTCAAGATTCGTAAACGAAGGTTTTTACGAGAC

CATTTCTCAAAATGGCGGAAATCTTTTCGAGCTCCTTATTCTGCTCAGCTCGTTTTTGAATCCAATCCTATACTCTGGCG

GGACCAACACAGCTCATCGAGTCTAG

>pR52

ATGGACCCTAAACCAGCCGATATAGTCAAAATGTGTGTTGATCATTTCGACAAACTTCTTGAACCAATAAATTGCATTCA

AGTTATTGCCACAGTACAAAGTCTCGAAGAACAGGGAGTGGCAAAATGTTCGATATCACCAACTCTTCCAACATATCGAC

ACATTATGAATTTCACATCAGGATGCGATTACGTGGAACTCACATATCTCCAATACGCCGTACCACCTCTTATGATGCTC

TGCTTTATCGGAAATATGCTCAATGTGCTCATCTACGGACTTCCGTATTTCGAAGGCTCGAGCTCGGTGCATTTTCTTCG

TGCGAAAGCAATCGCCAACATGGTGTTCATGTTTTCTCGAATTTTGGAGGTAATGCATGCCTCGTCTCCTCACCCAATTT

ATTGGCTGGAGCCCTTGTTTTGGAAATCTCGACCATATATGATGATGGTTTCGAACATTAGCGGAACAATGTCCACGTGG

CTAACTTTGATGGTTACAATGGAGACCGTCATGTGCATCATGACCCCATTCGTTTTCCGGAAATATTGCACAAAAAAAAT

GACATGGATTGTCCTAATCCTATCAACAATTGCTGCAACTCTTCTTCATGTTGCTATTGTTATAGTGACCGATGTGCAAG

AAATCGTTCAAGTGCGGCAATATATTCAAAATTTCAAGAAAGAAAATGCTCCTTGTTGGTTCCTGCAATCAGCGTTTCGT

GCCAGAAATAATCCGTACTATGAGATTTATCGACGGTTCTATGCAACCACTACTATGGCTGTTTCCATAGTCATTCCAAC

AATTGCCATGCTTGTTTGCACAGTGCTTATTATCAAGAAATTCACGTTCAAGAACCTCGGAGAAACGTTCTCTCAACGTC

GGAAGTGTGTGATCCGAATGACAGTTGCCACAACGCTCACTCATTTGTTTTTTGAAGGACCAGCTACGTTAACTCACTCA

GCAAGTGCTATTCAGGGAGACAACTACAGTTTTCTGATGTGCGTGCTTAATCATGGAAACAATTTGGGATCTCTGGTCAA

TGCCACCATTCCATTCTTTGTCTTCCTCTTCTGCAACCAACAATTCCGGCATATGACTGTTATGTATATCAAGGCTGTGA

TACAAACGGATCAAGTAAAACGAAAATCATATTTCTCTCAAGCAGGAATGCGTTGTGGTCGAATGTCTCGAATTGAAACG

GATCGGTCGATGGTTGAAACTCGCCTCGTTTCGAGACCCTCTAATGTTTAA

>pR53

ATGAACAACAACACGTTGAACATAACCAACCAGCGCACTGCTGCTGCGATGAGTCAAATCTACTTCCTGGTGGTCTACCA

AACGGCGGTGATGATAGTTTCCCTATTGGGAAACCTATTCCTGCTCTTCGTCATTTTTCGTGCCAACCAAGTAATGAAAC

GACGGGTGTCACCAGTTCAACTTCTCATAATTCATACGTGCGTTGCTGACCTTCTCTTTGCTCTTTTGTCTCTGGGCACG

GAAATTTTGACGTTGCGAACATATCCTCAATACTATGGCTCAAACTTTGTATGCAAATTGATGAGATATGTACAAATGTT

CCCAATGTATGCAAGCCCTTTTCTGCTGGTTGCAATCAGCGCGGACCGGTATCAGGCAATATGCCGTCCACTGGCTCACT

TCCGCTCATCACGATATCGTCGTCCAAATTGGATGGCTGCAATTGCCTGGGGCCTCGCCTTGGTGCTCTCCATTCCGCAA

TTTTTTGTCTGGACGAAGCACTCGAAAACTGGACGCTGCTCGACTATTTATGGACAGAACAAGAACACCGTGAAAATTAC

ATATGTGATCATGTTCAACACACTGGCCTGGCTTCTTCCCTCAATTCTTGCAGCAGTTTTTTATTATTGCGTTTGCAAAG

CCGTCCGCCTATCTTCGACAAAATCAGTTAGAGCAATGGATAGTCAAAAAAGGAATGGAAAATACTCTTCTGGAGCAACC

GAGGACTACATCGAAGAGCTCCGCAAGAAATCTAAAGGCTTCCGTCAGCAAATGTCAGAGTTTGATAGAAAACGTGTTCA

GACAGTCCGTTTAACAATCACAATTGTTGCATGCAATTTCTTCCTTTGGATGCCGTTCTGCCTTATTAATGTTATTCAAG

CATTGTGGCCTGAGATCTCACATATCATGTTTATCAATTATGTCGCTATACTTGGAAATCTAAATTCTTGCCTTAACCCG

TGGATCTACATTTTGTTCAACCGATCACACGTTCGCAAGGCACTTTGTCGTTCAAGACGCTCGTTCACAGAAGTTACTAA

AAAACGAAGTTTTGAAAACTTCGAGTGTTCCTCAACAGCAACAATGAACAATAATTATAACAACTGTCATGCTTACACCG

CTTTCAGTAACCGGTCTCAACTGAAGTTTGATTCATATGCAACAGACTCTACTTCCTTGAAGACCAACTCTAATTAA

>pR54

ATGTCAATTGCAGTAATTCTTGAATCAATTCAAGGTTCAAGTTCATTTGAATGGATTCGAACTAATTTGAGGCATTTATT

GCCTATATTTTGTTTTATTGGAATTCTTGGAAATTCTATGGCATTGATTCTGATCAGGACAAATTTCTGGTTGAGAAGGC

TAACATCAAACATCTACCTCTGCACTTTATCAGTATCATCATGTTTTTTTCTACTTACGGTTATCATTTCATGGGCAGAT

ACATATATGGGATTACCACTTTATTCAGAATCTGAACTTGGTTGCAAGTTCTTCTCATTTCTTGCTCATTTTAGTGATTT

TATCTGTGTTTGGATGATTTCATTGATCTCTTGTGATCGAATGATCGTACTTTATAGACCTAGAATCAGAAAATGGGTTT

GCACGAAGAAATTCTCTCGAAACATGACAATTGGCTTTGTGTTATCATCAGTTGTCCTCTACAGCTGGCTATTTCTACTT

GCTGGTCTTCAAATTTATAGATTAAAAGATGGAACTGAAAACACGTTTTGTGGATTGAGTCAGGATGTCAATTTATTTGG

ATTCCAAGTGGACCAACATTACTTCATGTTCACCCTCATGGATACAGCTCTTTGCACACTTCTTCCAGCAATTCTCATCA

TCATTGTCAACTCATTCTCCACATATCGGTATCGTCAATGCATGAAAATCTACTCGTCTGGCGTGCTCCGTGTTCGATTT

GTGCGTGCTCCAACAACTCAACAACAGCAGCAACAACTCAACAATGATACCATTCAATTTGAAGAGACAACAGTGAAAAA

ATATCTTCTGAGCACAGACAATACACATTCAAATCTTCCCACGAGTTCAACTCAATCCAGAAATTGCGGAAAACTTCGAA

GTTCTGATTTACAATTGAGCAGAACACTTATCATTGTCACTAGCACATTTGTTCTTCTGAATGTACCAAGTTATGCAATG

CGTATTCTTCAATCAATAATCAGTTCAGCTGGACCATTATTCAATTTTGTATATTATGTTACTTTGCTCATCTATTATTT

ACATCATGCTGTTTTATTCTATATGTACATTTTCTGGAGTCCTCAAATGAAAAAACAACTGAAACCGACGGCGATGCGAC

TTTTGGAATGCTATTGCCTGAAGACTGTTCCAGATTTTGGACATCGATCTACTTCAACACAAGGTAAAGAACTTGGATAA

>pR55

ATGCAATTGACTGGATGGCCCGAATATGTTTCAATGATTTATCTTCCAATAATACTAGTTGGATTAGTTGGAAATGGATT

GTCCCTATACGTCTACACAACTCCAAACATGCGAAAGTCAACTGTCGCCTTCTTACTCTACTCACTTTCAATTTGTGACA

TTTTTGTGCTTCTTTTTGCGCTTCCACTATATAGCATCTCATATTTGCCTATTTGGGACAATGTCTATGGTGCTTGGTCA

ATGCGCCGTATGTTTATTGCATTTTCAACAAAATTCTTCTATCCGTTATGCATGACTGCAAAAACTGCAAGTCTTTATAT

AATGGTGGTGATTACTGTAGAACGATGGATTGCAGTCTGCCGACCGTTACAGGTCCACATTTGGTGCACATTCAAAAACT

CAGTTAGAATTGTTATAGCCATCATAACATTTTCAATTATTCTGAACTTCCCCAAATTCTTTGAATACCAAATTGGCTAT

TCGGATGCATTGGGATACTGGCCAAAACGTGGAATTTTGGATGCAGAAGAGCACTGGTGGTACTACATTTCATATTTTAT

CATCATTTCTGTCATTTTTGACTACCTTCTGCCATTTGTTATAATGTTCATTGCAAACATGAAAGTCATCAATGAACTGA

GAAAATCAAGGAAGGAGAGAGCACTTTTGACGACTTCGCTTCAAAAAGAGCAAAATACAACTGTGATGTTGTTGGTTGTG

ACTATTCTCTTTGGATTCTGTCACTTTTTCAGTATGGCTCTCAAATTGTTCGAGAGCATTTTTAAAGATTTTCTCAATCG

ACACAATGAATATTTCGAAGTGATGATTGAAGTATCTAATATTCTCATCGTCATTCATATTGGCACGACATTCTTCATAT

ATTACTTTTTCTCGGCTAGATTCAGGAATATTTTGTGCTATTTGTTCAAAAAACGAAGCAACCTTCCGGATAATAGCCTC

ACAGACTTGAACAAACGGAAGCTTCTTCAAAAATCCGATTCAATATGTACTTCACTGGCAAAAACATCACCAAAAACATC

AATGGCATAA

>pR56

ATGATAGACCCGAACAATATGACTACAGTGATCGAAAAAACGACTGAAATCGTGAACAGTGTTCTAACGAGCAACGAATC

CGATGGTGTTTTGGAACAAACGAGCACATTGATAAGTGTACTGGCCACCCAATCAGCTCCGCTTCAATGCTTTTCCTGTC

ATCCGAAGCTGTACTTCATCGTGTTTGGTCTGGTTTTTATGATCATCATCTGTGCGGGCGTCATCGGAAATATTTTCATA

GTTTTTGTGATTTTAATGGACCGAAAGCTGATGAGCTCCTCAGTTAACCAGTTTCTCCTGAATCTTGCCATTGCGGACCT

TGGAAACCTGATATTCTGCTCACCAGATGCAATACTGGTATTGATTGATCGGGGTTGGCTCCTGCCGAATTTCGCGTGCC

ACTTGCTTCGATTCCTTCAAGAATACTTTCTATACGCTAGTGTCTTGCTTCAGATGGCGATCGGTGTCGAGAGATTTCTG

GCCATCTGCTCCCCAATGAGAATGCAGAGATTCTCCACAAAAACTACGATTTCGGTGCTCGCAGGTGTTTGGTGCGTGGC

TGCGTGTTTTGCCTCCCCGTACTTTTTGTATCAAGGCATTGTGTTCCACAAGTTATACTTCTGCTTCTGGAAAGGAATCT

CCCATAAAACGCGAACCTACTTCAAGTACTGTGAGCTGATTGTTCTTTATGCGATTCCGTTGGTTTTCCTCACCACACTC

TATTCCATCATGTGTCGTGTTCTCTGGGGAAACGAGGATAATCACAACATTGCAAATCATAGTCAACAAGAGGCCATTCT

CAAATTGCGACGTTCTGTGGTCAAAATGCTCATTATCTCAATGCTCCTATATTTTTTGTGCTACACACCAATTCAGGTAC

TATTTATGCTCGAGAAAATACTGGATCACAGCGTTCAACTTCCTCAATGGCTGCGTCTGCTGCTCAATGTGCTTTCAGTA

ATGAGTTCCTCCACAAATCCCATCGTCTACATCATTTGTTGCCGTCACTTCCGCCTACGCCTCCGCGACGTGGCCGCCGG

AATTTCATCGTTCTGTTGCTGGCTGCTCCCTTCGTTCTCGAAATCGGAATATGAGTGTGTGGATGAGTCAATGACGGCAA

AGTTATCACGTTCTCCGTATGTCTCCTTTCGCAGCTCTCGTCGACAGAATTCACGTGCGAATCTTTCAACGCTTCTATGA

pR57

ATGGACATCGGCAATCAGACAGCGTTAGAATCGCAAGCAGAAAATATTTCCAGTTATGACAGCACGATCATGCAACCTGA

TTACACAAATCCGTGCTCGGCGGCGGATGACTATCTGATCACGGAACGATTCTGGTTATGTGTCGTTGCTGGAGTGACTG

TGTCCATTATATCAATTGTGTTCAATACATTTATATTCTTTGTTTTTGTCACAAATAAGCAACACCGACGTTCACCAAAC

CTGTACCTTTTGCTACTTTCCCTTTTTGACGTCTTTATCGCATTTGCTTATATAGCCGTGATGTCAGTTCGGATTCTTGT

GAACTTCACGTCCAGCGTATTTCTAAAATCTATTTGGGTCCATTATATGATTCCAATGCTAACTGTTTCTCACATCGGAA

TCACTTCTTCAACTTTTCTCATTTGCTTTGCATCAATTGAGCGATATTGTATTACAGTCAATCATTTCTTTGTACCGTAC

CTTCAAAGATTTCGGCCTCTTCTTGCATTTACTGCAATAATGTGTGGAGTTGTGAGCAAGGGAACAATTGTAAAGGAAGT

CGATATTTCTTACAATCCAGAATGTTATGGAGAACTGAACTATTGGACGGTTGTTCCCTCACAACTCCTATTTGATTTCC

CAAAGATCAATGAATATTGGCGTTTTTACTTCCGTAACATTTTCACAATCCTTGCACCATTCTTCATACTTCTTCTTGTA

AACTGCCTTCTTCTTTTCCAACTCCGCGAACATGTCTTGAAAAGTAAATGTGCTGATCATGATAAGCAGAATGTGAAGGA

GAAGAAAGCCCGAATTCGTGCAACTACAAAATCTGTCGTTATCATCGTTTGCACATATCTGATGTCCAACCTGTTGTCAG

TCATCATTACTATCTGGGAATATATAGATAAAGAGTCCATTTTTAGCGAAGATTGGATAGCATTCTACGTTCTTTCTGTT

GATGTTATCTCCCTTTTGACAATTGTGGCTTCGTCTGCTCGTTTACCAATTTATGCATATTTCCAACCGCTACTGAGAAA

GGAAATGGGACAGTGTCTAGGTGACTGGTGGTGTTGCTGTTTTTCTGAAAATGACAAGAAAATGTCTCTTCTGGACGATC

TTCAACTTCCAAAAACCCAATTAATTTCAACACCAGATGGTGAAACTCCAAGCATCAGCTCCAAAATTGAATTCGTCTAA

>pR58

ATGAATATCGACTCGAATTTAACAGATTTTCAATGCGAGTACCCGGCGATTGACATAGGAATTCATATGAAAACCTTGTT

AGCAGTCGGGTATGGTTTGGTGGGAGCCCTGAGCTTAGTTGGAAATCTTGCTGTATTGCTCATTGTGATATGCAGGAGAG

AAATGCAAACAGTCACCAACATTTTTATAAGTAGCGTCTCCGCGGCTGACTTGGTAATAACAAGTTTCTCCTTATGGGCG

ACCCCATTAGCCTATTATCAACGGGTATGGCATTTCGGAAAGTATATGTGCTATATGGTTTCCATCATACAAGGCTTATC

CCTCATGTGGGTACCACTGACCCTAGCAGCAGTTGCATTAGATCGCTACTCTCTAGTTGCTTCTCCGTTCCGTCAGCCAA

TGTCTAAAAAGACTTGCTTGCTGATCATCGCTGGAATCTGGATGGGAGGGTTTGCAGTTTTGTCGCCAATGATTCGGATG

GTAGATTTTGTTGACAGCTATGGACCATGCCATTTTTGCCTGGAATCCTGGGACCACGACAAACAACACTACCGACTTTT

CTACGGACTCTCGGTGCTCGTGATCCGCTCCGCAATTCCACTTGTTCTCATTTCCCTGTGCCACTGGAGAATCGCAGTTA

TTTTGAACACGCAGACGAAGAAATTCCAAACATTACGTAGTGCCAGCACAGTCACCCAATCGACTGACATCCGCCGTAAA

CAACGTCTTCAGACTCTCTTATTGGCAATGGTGGTCATATTTGCCGTGTCCAGTCTCCCACTGGACCTTTCAAATGTTCT

TCAAGATTTGATCGTAGTGTACCAGGTTCGACCTGTCCCCGACAACGTTCGCCATTTTATCTTCTTCTTTTGCCATTGGA

CCGCAATGGCAGGAACACTTCTCAACCCGTTAGTCTACGCGTACTACAACGAAAACTTCCGGCGTCAGATTCAAACATGC

TTTGGAGAAATGCGTGGACAGGGAGAGTTCAAGCGGGGGCTATACTCTATTGTCTCCGGCAGATACTCCTACCGAGAAAC

TTTCAGAGCAGACGACGAGGAAAATCATCACAGACAAAACACACGTATCGAACTGGCTAACAATCAAACTGGTGATATCG

AAGTTCTCAGAACGGATCTTTAA

>pR59

ATGAATATCGACTCGAATTTAACAGATTTTCAATGCGAGTACCCGGCGATTGACATAGGAATTCATATGAAAACCTTGTT

AGCAGTCGGGTATGGTTTGGTGGGAGCCCTGAGCTTAGTTGGAAATCTTGCTGTATTGCTCATTGTGATATGCAGGAGAG

AAATGCAAACAGTCACCAACATTTTTATAAGTAGCGTCTCCGCGGCTGACTTGGTAATAACAAGTTTCTCCTTATGGGCG

ACCCCATTAGCCTATTATCAACGGGTATGGCATTTCGGAAAGTATATGTGCTATATGGTTTCCATCATACAAGGCTTATC

CCTCATGTGGGTACCACTGACCCTAGCAGCAGTTGCATTAGATCGCTACTCTCTAGTTGCTTCTCCGTTCCGTCAGCCAA

TGTCTAAAAAGACTTGCTTGCTGATCATCGCTGGAATCTGGATGGGAGGGTTTGCAGTTTTGTCGCCAATGATTCGGATG

GTAGATTTTGTTGACAGCTATGGACCATGCCATTTTTGCCTGGAATCCTGGGACCACGACAAACAACACTACCGACTTTT

CTACGGACTCTCGGTGCTCGTGATCCGCTCCGCAATTCCACTTGTTCTCATTTCCCTGTGCCACTGGAGAATCGCAGTTA

TTTTGAACACGCAGACGAAGAAATTCCAAACATTACGTAGTGCCAGCACAGTCACCCAATCGACTGACATCCGCCGTAAA

CAACGTCTTCAGACTCTCTTATTGGCAATGGTGGTCATATTTGCCGTGTCCAGTCTCCCACTGGACCTTTCAAATGTTCT

TCAAGATTTGATCGTAGTGTACCAGGTTCGACCTGTCCCCGACAACGTTCGCCATTTTATCTTCTTCTTTTGCCATTGGA

CCGCAATGGCAGGAACACTTCTCAACCCGTTAGTCTACGCGTACTACAACGAAAACTTCCGGCGTCAGATTCAAACATGC

TTTGGAGAAATGCGTGGACAGGGAGAGTTCAAGCGGGGGCTATACTCTATTGTCTCCGGCAGATACTCCTACCGAGCAGA

CGACGAGGAAAATCATCACAGACAAAACACACGTATCGAACTGGCTAACAATCAAACTGGTGATATCGAAGTTCTCAGAA

CGGATCTTTAA

>pR60

ATGGCGGATGCCACCACCGCCTCGCCGTCGACCACCTTCATTGCCTTCATCTTTGACTGGGCCCTATACTTCATCCAAAT

GTTGTCGCATTTGTTGAACGGATATTTCACGGCTTTCTTTGTATTTACTGGAATTATTCTCAATTTGCTATCGGTGTCCA

TATTTCTGCGAAAAGAGCGAGCCGGAACACCTGCAATACAATATTATTTGGTGACACTAACATTATGGCAGACGGCACTA

CTTGCCAATGCTTTTCTACTATATTCATTCCCCAATCTCTGGTGGGGTCATCTTGTTTCACAAGGCACTTACGTGTACCT

ATACCCCTATGTGTACACTTTCGCCAACACCACTCACACCGGATCCGTCTGGATTGTTTTGACACTGACGATCGACCGAT

ACTTGGCTCTGTGCCAGCCACTCAAACATCGGGCGATTGGCAAGAAACGAAGAGTGCGAAGGCTGATGATAGTCGTGTCA

GCGATGGCAGTGATGTTCTCAATTCCAAGGTTTTTCGAAGTTCACGTGATCTTAATATGTGACGAAGATCAATTATCATG

TGTCGCCACAATTGATAGGACTGAACTTTTTGACAACCGTCTCTACTGGACAATCTATCATGTTATCCTTGCAATGGTCT

TTGTGACCCTTCTCCCTTGCCTTATTTTGTTTGCACTGACTCTAAGAATCACAATTGCCCTGCGTTCCGCTATCGCAAAA

CGGAAATCTTTATGCGCACCGAACTCGGATATCGATACACGGTGTAAATCAATAAAATCATCGCGATATAATTCTTCAAG

GAAGGACCATAAATCGAATATAATGTTGGTTTTGGTGATCGCAAAATTTTTGGTCTCCGACATTTTACCCACGGTCATTG

ATGTTTTGGAGCATGTTGTCGGCCAGTCGGCGTTCATGAGATCACCGCTGGCTTCCTTGTTTGTCGACATCTCCAATTTC

CTAATTGTGCTCAATTGCTCATCGAACTTTTGGGTGTTTTTCGTGTGGGGCAAGCGGTTTCGAAGGTCGTGCCGAAAGTG

CATAAAGTCGACGCTAATTGGAGCATGTATATATGATATGGCTCAATGGAATAATGACTCGGAAGTATCCCTCTGTGCAC

CATCATCATTCACAACTGGCAATGCCACAAAGGTCTATGGTAGCACGGAAAGGTCGTTTCGTTATAAACGGAACATGGAA

TTAATCGTATCTAGAACCGATGAAACGTCTGTAAGTAGAACTTGA

>pR61

ATGGCGGATGCCACCACCGCCTCGCCGTCGACCACCTTCATTGCCTTCATCTTTGACTGGGCCCTATACTTCATCCAAAT

GTTGTCGCATTTGTTGAACGGATATTTCACGGCTTTCTTTGTATTTACTGGAATTATTCTCAATTTGCTATCGGTGTCCA

TATTTCTGCGAAAAGAGCGAGCCGGAACACCTGCAATACAATATTATTTGGTGACACTAACATTATGGCAGACGGCACTA

CTTGCCAATGCTTTTCTACTATATTCATTCCCCAATCTCTGGTGGGGTCATCTTGTTTCACAAGGCACTTACGTGTACCT

ATACCCCTATGTGTACACTTTCGCCAACACCACTCACACCGGATCCGTCTGGATTGTTTTGACACTGACGATCGACCGAT

ACTTGGCTCTGTGCCAGCCACTCAAACATCGGGCGATTGGCAAGAAACGAAGAGTGCGAAGGCTGATGATAGTCGTGTCA

GCGATGGCAGTGATGTTCTCAATTCCAAGGTTTTTCGAAGTTCACGTGATCTTAATATGTGACGAAGATCAATTATCATG

TGTCGCCACAATTGATAGGACTGAACTTTTTGACAACCGTCTCTACTGGACAATCTATCATGTTATCCTTGCAATGGTCT

TTGTGACCCTTCTCCCTTGCCTTATTTTGTTTGCACTGACTCTAAGAATCACAATTGCCCTGCGTTCCGCTATCGCAAAA

CGGAAATCTTTATGCGCACCGAACTCGGATATCGATACACGGTGTAAATCAATAAAATCATCGCGATATAATTCTTCAAG

ATATTCCAGGAAGGACCATAAATCGAATATAATGTTGGTTTTGGTGATCGCAAAATTTTTGGTCTCCGACATTTTACCCA

CGGTCATTGATGTTTTGGAGCATGTTGTCGGCCAGTCGGCGTTCATGAGATCACCGCTGGCTTCCTTGTTTGTCGACATC

TCCAATTTCCTAATTGTGCTCAATTGCTCATCGAACTTTTGGGTGTTTTTCGTGTGGGGCAAGCGGTTTCGAAGGTCGTG

CCGAAAGTGCATAAAGTCGACGCTAATTGGAGCATGTATATATGATATGGCTCAATGGAATAATGACTCGGAAGTATCCC

TCTGTGCACCATCATCATTCACAACTGGCAATGCCACAAAGGTCTATGGTAGCACGGAAAGGTCGTTTCGTTATAAACGG

AACATGGAATTAATCGTATCTAGAACCGATGAAACGTCTGTAAGTAGAACTTGA

>pR62

ATGTCTGCGAATTTCTCGCATCTATGCCTAACCGATGAGCAGCAGAGAATGTCGGGTACCAATTTGGAATACAATTTACA

AAGGTACTTCTACCCAGCGTTAGCAATATTCGGAATTCTAGGAAACGTACTCAACCTGACAGTTCTGCTAAATCGAAGCA

TGAGGAGTCGAGCCAATTCGTTTCTAGCAGTTTTGGCATTTGCTGACATTATTTTTCTATTTCTCCTGTTTCCAAATATT

CTCGCGAACTACTCATTTTTCACATTTAATTGGTATTTTCGGTGGTTCTACTTGCACTCAAAGGTTCACCTGATCTCTCT

AGCAAACTGGTGTTCGTCTGTGGCGCATTGGTGCGTAATAGCAGTGTGTGGTGATCGGCTTCTTGGAATTCAGAACCCAC

TTTATGCGCGAGCGACTTGGAGATGGTGGAAACTCCCACTTGTTACCACAATCATCGTCTTCACATGTGGTCTTCTCACA

TGCTACCAGCATTTCGAATATTTCTGCTTAGTCAGATCTTACTGCCGCGAGACTCAACTCTATTCGAGGTGCCTTCCTGT

GAATGCAGAAAAATGGTTTGGCCACCGCCCTAACCCGTTCTCCCAGCGATACCAGAGTTTTATTGCAATTTGCAAGTTGG

CTCATATCTTCCTTATGATTATTCTCCCCATTATTCTTTTATTATTCCTTAACTTGACACTTCTTTGGGCGTTGAGAAAA

CGTCAGAAGCATTTGTCAATTGGAAAGGATTTTAATGCGGATAGACGACAAAACGACGTTCACATGCAAAAAACTGAGCA

CAGAGTCACGTTAACCGTAACGTTTATTGTGACAATGTTCACATTAACTAATGGGCCATCAGCTCTCGTGCATCTAGTAA

TGTATGCAACTCATGAGGAGCTCTATGATTTGACTATGATTTCCAGTACTCTTGTAATTTGTGGAAAGGCTTCTAACTTC

ATCCTCTTTTGCCTCGGCTCCAAACACTTCAGACTCCGGCTGCTCAAGCTGACTCGAAAGAAGATAAACCGAAAAATAGA

TTCTATCACTGGTTCACTTGTTCATACGACAAAACTTAGCACGGCTTCATCCAGACGTACTTCAATGCCAATTCAAGAGA

AAAAACGATCATCATCGTGTGCTCCACTGGTAGCCAAGAACAGGGCTCTAAGCACTGTCGAAACAATTTATCAACCACTT

TTGAATGGGAATCGAAAAGGAGATTTTTGA

>pR63

ATGTCGAATGATCTCGTGCCTTCAGTGTCTTCTATACTAAATGAAACAACACCTTCATATCAAAGTACATGTAAAATCAA

AAACAACCCCATGGAAATGGAATATTTCCGCCCATTCTTTATTTCTATGTATTGTGCTGTTTTTTTGGTTGCTAGTTCCG

GTAATTTTTTGGTGGTGTACGTTGTGATGACGAACAAGCGAATGCAGACGATCACCAACATTTTTATTACAAATCTCGCA

GTTTCTGATATAATGGTTAACTTTACATCGTTGTGGCTTACACCAACATACACCTCAATAGGACATTGGATATTCGGAGG

TGGATTGTGTCATGGTTTACCCCTGTTCCAAGGTACAAGTATCTTCATCAGTACGTGGACACTTACGGCTATAGCCATAG

ATCGATACATAGTGATCGTGCACAACTCATCAAATATCAATATAAATGATAGAATGTCCATGAGATCTTGCCTTTCGTTC

ATTGTCCTCATTTGGCTATGCTCATTGCTTCTGGTCACTCCGTATGCCATCAACATGAAGCTCAACTACATTCATGAACC

ATGTGATTTTCTGATATGTAGTGAGGACTGGAGCAATGCCGAATTTCGATCTATTTTTGGAATTGTGGTGATGATTCTTC

AATTCATTTTGCCATTTGTACTCATTGCCATTAGTTACATAAAAATATGGTTGTTCCTAAATAGCCGTCAAAGTATGACC

GAGAGAAAATCCGATATCAAGCGCAAAAAGCGTTTACTACGGATGTTGATTGTCATGGTGGTCATCTTCGCAATTTGCTG

GTTCCCATTCAACCTCTTAAACTGTCTCCGAGATCTCAAGTTGGATAATTTCATGCGTGGCTACTTCAGTTTTGTTTTCC

TTTCCGTGCATTTGATGAGTATGACAGCTACCGCCTGGAATCCAATCCTCTACGCATTCATGAATGAGACCTTCCGTGAG

GAGTTCGCAAAAGTTGTTCCCTGCTTGTTTGCGCGTCGTCCTGGAACTGGTCCAATTCGCGTCATCACTGAACGCACCGC

TATGATAACTAACCCGTTTCGACGTGCAAACCGAAAAAAAAAAGTGGAGGAACAGCCTGTTACGGTGATTTCTGAAACAG

ATTCAGAGAGTCCACTTCAAACTGCAGTGGAGCCGCAGCGCAGTATCGTGTATCTCGATGAGCCTGAGAACGGATCAAGT

TGTCAGACGCTTCTGCTTTAA

>pR64

ATGGAGGATTTTTCCTCGAATTTCACGACAACTTCAATTCAGAATGATAGTTATGAAGAAGATCAACAACCACATATCGA

TATGTCATTTGCAAAAGGTTTTATGACAGTTGTCTACATAGTCGTTTGTCTCGTTGGAACACCTGGAAATCTATGGATCA

TCTATAAACTTTTCCGTGCAAAATTATGGAGCGGTGCGTCAGTTCAATTAACGGTATCTCAAAGATCTCGAATCTACATA

TTTGCTCTCGCATGTTCGGACATGTTCCTTCTATTGACACTCCCTGCCACGGCAAGCTATAATTACCACGGGACTTGGAT

ATTTGGCTCTGCAGCTTGCTATATAATTCGATCCATCGAGATCTTCGCAAAACTTTTCTCGGTAGTATTGCTGACTGTGA

TGAGTTTGGAGAGATACATCATTGTGTGCACGCGGCTTCGTCACATCTACCGTGCATGGATGTCACTTGTACCTCTGGCA

GTCGGAACAATATTTGGAGTTCTCGTACCAACAATCATCCATTATTTCTACCTTCAACATTTCTCAGTACCATTTGACCC

AGATGTCACCTGGGTTTGTCTACCACTAATGTCAAATGAAGTCTTCAATCTGTTCGCCCAATACACATTTGTCGTTGGTT

TCATCATTCCGTTCGCTATAATGACAGCCTGCTACATCATGCTCGTTCGTCATGTTCGAACAAAATACAAGATGCGAAGA

GCTCAGACGACAACGACGTTGGCGAAACCAGGAAAAGAGCCAAGATATATGAGTGAAGTGAAGAAGAGTATTTGGAGAAT

TGCAGTGTTCCATTTTGTATGTTGGGCTCCGTTTTGGGGATTCACTATGCTTCCCAATTATATTTATCAAATCGATCAAT

TTTTGCATGGGGACAATGAGGAAGAAAGTGGTGGAGAAAGTATTTTCCTCGTCTATTGTCGTCTTGTCTCAAATTGTCTT

CCATATATAAATGCTGCAGGCAACTGGGTTCTATATGCTCTTCTGAATGTAGATGTCCGCAAGCATATCTACAATCAGCC

AAAGAAGAAGAGGAAGTTCACATTAAAATTCAATCCGGTCAATACATCATCGAATTGCTAA

>pR65

ATGAACTACGAAGTTTATTGCGGCAATGCGCATGCAGAGCCAAATAGCTCAGCAGTTCAACAACTTGCAGAAGCATGTGT

TGAACAAGACGCGTCTTATTTTGATGGTTGCTCGAGAACGTGTGTGAAAGACAAGTATCTACTGCCTCTGACACAATTCG

ACAATTTAGAGATTGTAGTTTATGGGCAGATTTTCCCAATTTTGGTGCTTTTCGCAGTTTTTGCGAATGCGGCTGTTGCG

CTAGTTCTATCGAAGAAGCACATGATCACTCCAACTAACGTCGTTTTGAAGTACATGGCTATTGCGGAACTTTTAGTGGG

ACTCGTCCCACTGCCCTGGACTCTTTTCTTTTTTTCAATGGGGAATATCAAAGAAACACATCGGTTAGAGTTATGGTGGT

GCTATCTACAAAAGTACAGTATGGATGCGTTTCCTCCAGTTTTCCATATGATTGCCATGTGGTTGACGGTACTTCTGGCG

GCACAAAGATACGTCTCCATAAGTCATCCATTGCATTCTCGATCCGCGTGTAATGTAAAAAATGTTCGATTAGCAACAAT

GATCATAACTGTCACATCTTTCCTTTGTGGACTTCCAAAGTCGTTTGATTATGAGTATGAAACAGTTCACGGATGGATTT

ATTCGCATGGTAACTGGACGTATGCAAGCAGTTGCGTCATGATGCCTACAGCGATATTAACCAATATGGGGCAAACTGTG

TACTTTAATATTTACTTCTGGACACGTGCGCTCGGCTTCATTATTTTGCCAAGTTTCCTACTTGTCCTGCTCAATGGCCT

TCTCATTAAGGGTATTAGAAGAGCGCAGCGGCGGAAGCTGCGATTGTTGAGAGAAAAACGTTCCGAAGAAGCTGCTCGGC

AACGAGATAGCAATTCGACAAGCCTTATGCTAGTTGCTATTGTTTCAATTTTTTTGATTGTAAATTTGCCGCAGGCTATA

TTTATGGGCCTGCTTTGTGTCTGTGAGACGTTTACAATTAAGATTCCAATATTAGAAGGAACTTTCCCTGCCGTGTTTCT

TATCGCTAGTAATATGATCGTTATTGCTACTTATCCCATCAATTTTGGAATCTACTGTTTTATGTCAAGCAGTTTTCGAC

AAACTTTCAAATTGTTGTTTTGTCCTGGCGCTAGTCAACTCCAATGCGAACGGCGAATCGAAGCTGCAAGTGCCGTACAT

TCCAGTCGTCGACGTTCCGACATTTGCTCGCATTTGGTGAATGTTTGCACAAACTCTGAGGGATTCATGCAAGTATCCCA

TCATTGTCTGCACGTCGATTATCTTGTATCGGATAGGCAGTCGACTCAGTTTACAACGATGGATAGGAGTGATTAA

>pR66

ATGGCAAATGAGTGCCCAAGTCCTCCTTCTTCGTGGGCTTTAGCATTTGATGGTCCATATTCATTTGTGGTTATTATCGG

AGGAATTATTGGAAATGTGTATTCATTGAAACAACTATTTTCTCGTTCTATCAACACATCAATGTTGGTGTCTCTCACCG

GTCTGGCTATATGGGATATTGTACTTTTGATCGCTGCTCTATGGCATCATTCACTGTGGGCAACTATGCACTACTTCTCA

CTACGCGACGAGCCGTGGGATGCTGAAATGGTGGCTACAAATGCGCTCGTTGAATGTGGCCACATTACATCCACTTGGAT

GTTAATTGAAGTAACTGCAGAGAGATTCATTGCGGTCACAAGACCTTTTCAATTTGCTCCAGTTCATCGAAAGCAGAGAA

GGAAAAGCTATGCAAGAGTTGTCGGAGGCTTGATTCGTATCCCGCTAATTATGACAATTGCTGCATGTGTCATATGTTTA

CCGTGTACTGTGGAATACACTCTAGAACCGTGTACATACAAGGGAATTGATTCACAACAGATGCTCGAAACGCCTTTAAT

GCAAAACATTTTCTACAGAGTGTTGTATAGGACGGCTTTCCTTTCTATTGGGAAAACGTTCGGTCCTTTTGTGATAATAT

CGTTTTTGACTATCTCTACTCTGAAAAGTATGAGAAAAAGCATGGATAGCAGGGCGTCAATTCTTATTGCGCAAGGCCAA

AATCACCTTTTCCAAGCCGACAAGGACAAAACAAAAAGCTTGCAGGCAATCTCCATCATGCTCTTAGGAAAATTTCTTTT

TCTTAGATGCCTACCTACTGCAATGGCAATTATTCAGATGTTTCCGGGTGCTGATTCAACAACTTTCCTGCCCGTTTATT

TATCACAGTTCTTCCTTCTTTTCAATTCGGCAACCAACTCTTTTGTTTTCGTGGTTGTAAAATCGGCGTTTGAAACCCGA

AGGCTCAAAAGAATTCGCCAGCGTCATCGTGAACTGGTGGCTCAACATGCAGAACAAGTGTTGAGCATAGGAAAAGCTCT

TGCTGGGGATAAGTTGTTCTTACTTCCCGAAGTAGAATTTGACAATCAATCATCTGAAGAAGAAACGCTCGAAATGCAAC

CAATGATGCCTTCTGCAACAACCTCTAATCCGGTTTAG

>pR67

ATGAACTCGACAGTTATTTCGCTTCTTAATAATTCTACACTGGTCACTACGGATCAGCCAGGTTTTTCTGGAAGTACAAA

AGGAACATGCAGCTATCCAACTAATTTTGCGGAGGTCGATTACATTAGCACATATATTTACCTGCTTGCCATTCCAACAA

TTTGCGTGCTGGGAGCCGGCGCTGCGGTAATGTGCATTGTGGTGTTCACTAGGAAACAAATGAGATCATCACTCAACGTA

TATCTTGCCGGATTATCGGTTTTCGATCTTATCCTGCTTTCATTCAGTGCTCTGATCTTTTCACCTCTCCAAGGTTGCGT

TTTGCAGGGTCACGGGGACACCGCGGTTTGTCATTTCTTCTGGAGATCAACTCCTTGGACGTTACCCATATCAAATATCG

CACAATGTGGTAGTGTCTGGACATGTGTTGCAGTGACGGTGGATCGATTTCTTGCTGTTAACTATCCGCTTCATAGCAAA

ATTTGGTGCACACCGCGCCGAGCAACAACAATACTTATTGCGATTACGGTTTTTAGCATATTGTTCAAAGCACCAATGTT

CTTCGAGTTGACCAACGATGACTGTGGTCGACTGAGAACCTCATTTCTGCGGGATAATAAGTACTACAAAGAGTACTACG

TAACATTTGGGTACCTCATCGCGCTGTTATTGATTCCATGGACAGTAATGATTATACTGAATGTGTTTGTGGTAAAGGCT

GTACACAAGGCGTATAAGATTCGGAGGTCTATGCAAGGAGGAAAAAATAATCAAGAAGAAAAAGATCGGAGAAACAGTCT

CAGATGCACTCTTATGGCCATCGCTATGGTGCTCACTTTCTTGATCTTCAACGTAGTAGCAGCAGTGAACAACTTGGCTG

AGACGGTGTTCGAAGTGAGCTTAGGATTTTGGAGTCCCATTGGAAATTTGCTAATTTGTTTAAACAGCGCTTCCAACATC

GTGATCTACTCGTTATTCGGAGCCCGATTTCGTCAAATGTGTGTCCTCATGCTTTGCGGGCGACAAAGTCGATGGATGGA

GTGGCTCAAAGTTGGAAGGAGCATGGTCAGTGATTGGGATGGTAGAGATGGTACGACGAGGAAAACTAGTTTGGACGCCA

GTACTGCTCTGTTAAGTGCGAGGAGTTCTCGCCGATGGAGTAAACGGAGTCAAAGTGGTGCACCACATCAAGCAGCTCCT

GCTCGAAAAGATACAGTAATCTACACAAGTGCTCAAAACAGACTTTCGCTTCAAAGCGCACCTTCATAA

>pR68

ATGACATCTACTGTCATCTACGTTCCAGACATTCAAGAATTACCAGAATGTGAGCCATCAGTATTCAATGATATTCAGGC

AACCATTCGATTATTCGGAGGCATGCCAATTGCTATATTTGGTTTTATCACAAACGTGATCAATGTGATAGTATTTTGTG

ATCCAGAGATGCGATGTTCTCTTGTAAATCATTTTTTATTGGTGCTCTCAATTTCCGATCTAGTACTTTTGGTGTGTAAT

TTCTTTATGCTCATATTCCCAGTTATTGCATCAATGAGCAATAGCTATTTGTTACATGATTATTATCCTGTATTTCTGTG

GTTTGCGTATCCAGTTGGATTGAGTACACAAACTTGTGGTGTTTATCTCACAGTTCTGGTTTCTGTGCACAGATATTTAG

GAGTTTGTCATCCTTTTCGAGCAAAACGATGGGTTTCTGGAAAACCTGTTAAATGGGCAATTATTGGCTCAATAATCTTC

AGTATAGTAATAAATCTTCATACGTGGCTTGAATTGGACATTCGTCCTTGCTATTCTATCAACTTCAATGCACCTATCAG

TTCAATTATATTAACAAGTTTGAGACAAAAATCATCATACAATTTGATAACAAAATGTATAATGTATACATTGATAATGT

TCATCATTCCATTCATTACACTTATAATAGTTAATTGTCGAATTGTGGTTGCATTGAAAGAATCAACGAGAATGAGAAAT

GGTCAATCAATGAAGAAAAGCACACAATCTAGATTCACCTGGAAATCTCTCGACAAACGAATTATGAATAATTTTCGGAT

GCTCAAAGGTGCCAAATACAGTGAGTTATTTGGAAGGTTCGGGCGACTCAATTTCAACCCGTTGAAAACGCCATCACTTC

TAAAAACAAATGGAAATTCTCTTCGTGATCGTTCTGTAACTCTTATGCTTCTTGCAATTGTTGCCATATTTCTCTGCTGT

AATTGTCTTGCATTCTGCAACAACATCTACGAAAACGTGCAACATGTAAAAAAACATGCATCGGATTCTCAATCCCTGAA

TCAAACTTCATTTGTTCCAGAGATCGAAGAACAATCACAAGATTATGATGGTGAATGGAGCATTTTTAATGAATGGAACT

TTGACCTTTCTGTTGAAATATCAAATCTTCTAATCAGTTTGAATTCATCCAGTTCTATGTTTGTATATCTGATTTTTTCA

TCAAAGTACAGATCAATTATCAAACACTGGCTGGGATTAGAGAAGAGAAAAAGAACAAATGGAGTTGCCTTAACAACAGT

GATGGCCGCTCAGAAAGCATTGGAGCTTTCGATTCTTCCAGATGAAGTTGAAGCCCGTCGTCATAGAAAAGAAAAATCAC

ATTTTGTAAAAAATAAGAAACAAATGAACAAATCAGCTCAATTATTTCTCACAACATCAGAAATTGATCTTCGAAAAGCA

AGAACGATTCGAAAAGAACAAGAAGACGAAGAAGAAGAAGAGGAAGAGATTCGAGAAATTCAAAGCGATGAGCCAAGTGC

AGCCAGTAATCTTTCTAGAGATGATCAAAACCGTTCAAGCTCGAAACGAAAACTTCTTCGATTTGCCACTCTCGCCTAA

>pR69

ATGACATCTACTGTCATCTACGTTCCAGACATTCAAGAATTACCAGAATGTGAGCCATCAGTATTCAATGATATTCAGGC

AACCATTCGATTATTCGGAGGCATGCCAATTGCTATATTTGGTTTTATCACAAACGTGATCAATGTGATAGTATTTTGTG

ATCCAGAGATGCGATGTTCTCTTGTAAATCATTTTTTATTGGTGCTCTCAATTTCCGATCTAGTACTTTTGGTGTGTAAT

TTCTTTATGCTCATATTCCCAGTTATTGCATCAATGAGCAATAGCTATTTGTTACATGATTATTATCCTGTATTTCTGTG

GTTTGCGTATCCAGTTGGATTGAGTACACAAACTTGTGGTGTTTATCTCACAGTTCTGGTTTCTGTGCACAGATATTTAG

GAGTTTGTCATCCTTTTCGAGCAAAACGATGGGTTTCTGGAAAACCTGTTAAATGGGCAATTATTGGCTCAATAATCTTC

AGTATAGTAATAAATCTTCATACGTGGCTTGAATTGGACATTCGTCCTTGCTATTCTATCAACTTCAATGCACCTATCAG

TTCAATTATATTAACAAGTTTGAGACAAAAATCATCATACAATTTGATAACAAAATGTATAATGTATACATTGATAATGT

TCATCATTCCATTCATTACACTTATAATAGTTAATTGTCGAATTGTGGTTGCATTGAAAGAATCAACGAGAATGAGAAAT

GGTCAATCAATGAAGAAAAGCACACAATCACGAATTATGAATAATTTTCGGATGCTCAAAGGTGCCAAATACAGTGAGTT

ATTTGGAAGGTTCGGGCGACTCAATTTCAACCCGTTGAAAACGCCATCACTTCTAAAAACAAATGGAAATTCTCTTCGTG

ATCGTTCTGTAACTCTTATGCTTCTTGCAATTGTTGCCATATTTCTCTGCTGTAATTGTCTTGCATTCTGCAACAACATC

TACGAAAACGTGCAACATGTAAAAAAACATGCATCGGATTCTCAATCCCTGAATCAAACTTCATTTGTTCCAGAGATCGA

AGAACAATCACAAGATTATGATGGTGAATGGAGCATTTTTAATGAATGGAACTTTGACCTTTCTGTTGAAATATCAAATC

TTCTAATCAGTTTGAATTCATCCAGTTCTATGTTTGTATATCTGATTTTTTCATCAAAGTACAGATCAATTATCAAACAC

TGGCTGGGATTAGAGAAGAGAAAAAGAACAAATGGAGTTGCCTTAACAACAGTGATGGCCGCTCAGAAAGCATTGGAGCT

TTCGATTCTTCCAGATGAAGTTGAAGCCCGTCGTCATAGAAAAGAAAAATCACATTTTGTAAAAAATAAGAAACAAATGA

ACAAATCAGCTCAATTATTTCTCACAACATCAGAAATTGATCTTCGAAAAGCAAGAACGATTCGAAAAGAACAAGAAGAC

GAAGAAGAAGAAGAGGAAGAGATTCGAGAAATTCAAAGCGATGAGCCAAGTGCAGCCAGTAATCTTTCTAGAGATGATCA

AAACCGTTCAAGCTCGAAACGAAAACTTCTTCGATTTGCCACTCTCGCCTAA

>pR70

ATGACAACGATCAACTGCTCTCGATCTGTTCCACCTGATGTTACAGTCAATGATTCTGTTCTATCAATTGTCTTCACATA

TTTGGCTTTATTTATTCTGGCATTTGTTGGAAATGTCACAATGTTCCTGATTTTATGTCGAAATCAACTGGTAAAAGTTC

GACGGGTGCATTCGTTGCTTCTACACATGAATATTGCTCATTTACTTGTCACTCTTGTTGTCATGCCAAAGGAGATTCTG

CATAACTACATGGTTGCTTGGTTTGCAGGAGATGTTATGTGCCGAATTTGCAAGTTTTTCGACGTCTTCGCAATTAGTTT

ATCAATGAATGTGCTCATTTGCATTACCCTAGACAGATTCTACTCAATATTTTTTCCGTTATATGCGATGAGAGCCCGTA

AAAGTGTCCAACGAATGGTATCATTTGCATGGACAATCTCATTTGTCACTTCTGCTCCACAACTGTATCTCTTTAAAACT

GCAACTCATCCGTGCTTCGATTGGTACACACAGTGTGTTTCCAAAAATTTCATTGGAGAATTGTCGAATGATGTAGTCTT

TTTCTTCTCAATAGTAAATATAATTCAAGTCTACATTGCACCTTTATTTGTTACCGTCGTCTGTTATTCCCTAATTCTAT

GGAGAATCTCAAGAAAGTCGAAGCTGGTCGGGGAGAAGGAATCTGAAAAATCCTCAGAACTGCTTCTCCGTCGAAATGGT

CAGAACAACTTGGAGAAGGCGAAAAGTCGAACTCTAAAAATGACATTTGTAATAGTTTTGGCATTTATTTTCTGCTGGAC

TCCCTACTCAATTCTCATGTTTCTTCACTTTCTTCGTCATACTGATTGGATTCCGAAGGATATTCGAAAGTTTATTTATG

CATTTGCAGTTTTGAATTCTGCAATTTCTCCATATCTTTATGGTTATTTCTCATTTGATATTCGAAAGGAGCTCCAATTA

TTATTTGCATGTTCTAAAGCAACTGCAGCTGATCGTCATTTATCTTGTTCAGCGAATGTTAGCAGAAATCAAGTCACTGA

GAGAATGCGCAAACGGAGTGCATCGGCCTGTAATTTTGACGGCGGCAAAACAAATAACACATTATCGCCACGACCTCCCC

GTGGACACAGTTTAAGGCATAAACCAAGTTCGTCAGGAATTGATAAACGAAATCATAATGTTCAATTAGAAATTATTGAT

TTTTGA

>pR71

ATGAAACATTTCATATTAATCTTTCCCATTGGGGATGACGTAATGGGTACTATTATTCAAACTATATATTCTGTTATTGT

GTTCGTCGGCATCATCCTAAACATTTACGTGGTCAACAAAATGTCTAGATTATACAAATCAGACAAAGATCAGTTCATCA

ATGGAACAGGAATCTACTTGATGTCAATGGCAGTTTGCGATGCCGTCAATCTTGTGTTATCCTGCTTTGAAATGCTCACT

TACTTACTATCTATCACATCAAGCGAGCAGGCAGCAAGTATTTTGTGCAAGATCGCCGAGTTCACACTTCGATGCTCCTA

CACCTATAGTATGTACTGCTGGTTATTCATGTCTGGACTACGGTATCTGGCAGTGTTCTCTCCGATCCACTATACAACGC

TTTGGAGAAGCCCGTGGCACATGGTGATACCATGCTTCATTGTGGCCATTATTTCGAACATGTATTTGCTGGTTGCAGTG

AAGCAGGAAAACTTCCAATGTATTTTGATTATTGATGAGTATTCAACATTGTACAACAGTGTTGATGTGTTCATCTCCAC

TTTGTTACCAGTCTCCTTTATTCTCGCACTGGATCTCTTGGTACTATGTTTTCGTCCATCTAGACAGCAAAACGATCCAT

TGCTACAGATTGTTTTCCATAGACTAGATGAAGATACGGAGAAGAGGAAAATGATAACAACTCGGAAATTCATGCTGGTC

ACGCTGCTCGCCGTGAGCTTATCCACTCCAGATGGTATCCTTCGAGCTATCAGGCCGTATATCGACGGGAACGTTGTGTT

CATTTTGTTTCAAGTCTTCAAAGGATCTTATTTAGCAAGGTTCACATTCAACGCGTTTTACCTAACACTGTTCGTTTTCG

ACCGGAACATTTTATCCAAAGTTTCATCTTCTCGTCACTTATCGGTCTCGATGCGTCGTCTTGAAGAAGAGCCGGCAATT

GTTCCCCGTGAAAGAAGCCGAACACTTTCCTGTCAAAGACCACCACTAGTGCACAATTTGATGCGAAATTCTTCCTGTAT

AATTTACAAGGATGACAAAGATGACAAGGAGAACAACGAGACGTCGTCAACTCTGTGGGTGTGA

>pR72

ATGTCGACAATGACTATCAGTTCCACATCAATTTTAACAACGGTATCCACAACAACATCTAGTACAATAGTTCCATCACT

CAATGATCCAATGGAAATTCAATTTGAAATCATGTTTGTTATGGTTCTATATTCTTGCTTAGCAGTTACTGGAGTCGGTG

GAAACTGTTGGGTACTTATAAAAGTAATCAAACAGTTATCCGGATGCTCTTCAAATAGGAGTAGGCCTTACAAGACAGTG

GTTCAATCATCAGCCTACGTCTACTTGATCATTTTGTCTGTGGTGGACTTGTTTTCACTCATCCCTGTACCGATGATTGT

CACAGACGTATATGTGAATCATTTTCCATTTGGTCTATGGGCCTGCAAGCTCAGCTATTTTTGTGAGGCAATCAACAAGT

CCTTGAGTCCCATGGTGCTGACTGCACTGTCTGTGGATAGATACATTGCTGTCTGCCATCCCACGTTGTACTGGTTACGA

ACAACAAAATTTTCGTTGGGAGTACTTGCAGTATGCTTTTCGGTTTCCCTGATTTTCATTATCCCAGTCACAAAAAAGGC

CACAATGCAAGTGATGAAAGATAACACAGACGAGGAGATCATTAAATGTGCATTTGACCATGGAGACTCTGCAAGCTACG

TGTTTGATACGGCTCAAGCAATTGTCTGCTATTTGGTTCCACTCTTCATCATCTGCGCAGTTTACCTTGCTATTCTTCAT

AAGCTCTTCATTCACACGAGATTCTCAACCGTTGGAAAGAAGACAAGCATTTCGCTGGGCAGAGTAGTCAAATGCAGTGT

TATGGTTGTTGCTTTCTACTTTATTTGCTGGACACCGTACTGGACAATGAGAATTCATTACTTGGTGACAAGATTTATCG

AGACCAATAGTGATGGAAACTCCACGGTTATTGATGAGGTTGTTGCTGCAGTTGCAGAAGAGGTGCAGGATGGAATTGAA

ACGTCATACTTTTTTGGGCTGATATCAAAGAAACGGATGCACAATATTGAAATCGCATTGATCTACATTTTACATTCTCT

CACATATGCCCAATCTGCATTCAACTGGCTCTTCTATTCCTTTCTCAACCGCAACCTACGCTCGAACAACGGCCGAGGAA

ACGGAACACGATCAGCCGCTAACACATCCGTATTTGATAACGGAAACGCAACAAGCACAGTCAACACTTCTTTGACACCG

ATTTGGAAGAATATTCAGCAAATGGGAAGTCATATCAAAACTGCTGGACTTGACACTCGTTCTGCTTTGATGAAGAAGTC

GCCGTTTAAGGGAAAATCCAAAATTCAGTCTAGAAGTGCCGCCTACCTCGATTGCTCATCTGCTCACATGCTCAGACCGT

CATCTGAAAACACGTTGAGTTCCTCGCTTCTTGACATTCCAAGAAGGCCATCATTAATTCCAAGACAAGAACCGTCGCTT

GTCGGAAAAGCTCTTTCTTTCACAAATATTCCGCAGCCGCAATACCAACGAATAGAGCCGAAGGAGGAATCTTCTCACGG

AGCAGTTTCAGAAAGCTCGTCTGTCGAATGGTTGTAG

>pR73

ATGATCGAAATAAACGACACATGCGCCGAAGATCCTCCACAAGCTGATGCCTTTCGAATGATCCTCATAGTAATAATTGG

TACAGTAGTATGCTCACTCGGAATTGTGCTCAACACCTTTCTGCTCCTTTCACTTCGCCGGCTAGACGTCTTCCGATCAA

ATATCCTTTACCTTTTTCTTCTGGCATGCCTTGACATTCTCGTAGAACTTTGCTTTATGTTAATATTCCCGGCATCCCTA

GTCTGGGACTATTTCCGAGTGGAGCTTCTCTACACATGTTGGCACTTCTACATCAAATACGTGTCAACTGTTGGGCAAGT

TCTAATCGCTGCATCGACACTTCTCATCGTGGCTGCATCTTTTGAACGCTACATTTGCTCCTTGAAATCTTCGATTCAAT

TTTCCCCGCAGAGACGATTTCTGTTCATTTCCATTGTTGGAGCATGCGCCCTTTTTATGAAAGGATCAGTATTTTTTGAG

TTGGAATTGCAGTCACTTCCACATTGTCCACCATTTCAGAATTTGCGGTTGGATCTGTCAGAGATTACTAGAAGCGAAAG

CTACAAAACGATATGGATGTTCTGGTGTCGATCAATATTCAACGTTTTTCTTCCATTTTCACTTCTTCTGATTCTCAACT

CCCTGACAATAACCAACCTGAACAAGCTCCATTCAAACGGATTTCAGTCGGTTTTGGTTGAGCAAATGCCTAGCCTGCTC

CGTCGCAATTCAGAGGCATGTGCAGCCCGTCGTCGGAAGCGCGACGCGACACGCACGTTGGCAGCACTCATCACAATCTA

CCTGTTGACAAACACATTAAACCTACTGATAACAATTATGGAGTTTATCAATCCAGATGTGTTGGGCAGTTTAGGTGAAG

GATGGACGTACAAATACCTGGCCGACTTGTCTTCTGTGCTCACTATCTCATCAACTGCCTTCCGACTACCTGTCTATTTC

CATTGTAATGGAGACATTCGCGCACAGATACGACACTTTGCAAAGGCTTGTTTTATTGAGCAGAATGAGAAAAAGAAATT

GAAAAAACTCACAATGCACCATCCAATTGTGTCCTCCTCCACGGAATCTGTCCTATGA

>pR74

ATGATCGAAATAAACGACACATGCGCCGAAGATCCTCCACAAGCTGATGCCTTTCGAATGATCCTCATAGTAATAATTGG

TACAGTAGTATGCTCACTCGGAATTGTGCTCAACACCTTTCTGCTCCTTTCACTTCGCCGGCTAGACGTCTTCCGATCAA

ATATCCTTTACCTTTTTCTTCTGGCATGCCTTGACATTCTCGTAGAACTTTGCTTTATGTTAATATTCCCGGCATCCCTA

GTCTGGGACTATTTCCGAGTGGAGCTTCTCTACACATGTTGGCACTTCTACATCAAATACGTGTCAACTGTTGGGCAAGT

TCTAATCGCTGCATCGACACTTCTCATCGTGGCTGCATCTTTTGAACGCTACATTTGCTCCTTGAAATCTTCGATTCAAT

TTTCCCCGCAGAGACGATTTCTGTTCATTTCCATTGTTGGAGCATGCGCCCTTTTTATGAAAGGATCAGTATTTTTTGAG

TTGGAATTGCAGTCACTTCCACATTGTCCACCATTTCAGAATTTGCGGTTGGATCTGTCAGAGATTACTAGAAGCGAAAG

CTACAAAACGATATGGATGTTCTGGTGTCGATCAATATTCAACGTTTTTCTTCCATTTTCACTTCTTCTGATTCTCAACT

CCCTGACAATAACCAACCTGAACAAGCTCCATTCAAACGGATTTCAGTCGGTTTTGGTTGAGAATCGGTGTCAAAGTATT

GCAACAACTTCTGAGCTTCTTCCTGACAACCTGTCCTTCCAACCACTTCTCGGGACCTCACTCACCTCGTCAACTATCAA

CAGTTTTTCTTCGTTTAATGATCAAATGCCTAGCCTGCTCCGTCGCAATTCAGAGGCATGTGCAGCCCGTCGTCGGAAGC

GCGACGCGACACGCACGTTGGCAGCACTCATCACAATCTACCTGTTGACAAACACATTAAACCTACTGATAACAATTATG

GAGTTTATCAATCCAGATGTGTTGGGCAGTTTAGGTGAAGGATGGACGTACAAATACCTGGCCGACTTGTCTTCTGTGCT

CACTATCTCATCAACTGCCTTCCGACTACCTGTCTATTTCCATTGTAATGGAGACATTCGCGCACAGATACGACACTTTG

CAAAGGCTTGTTTTATTGAGCAGAATGAGAAAAAGAAATTGAAAAAACTCACAATGCACCATCCAATTGTGTCCTCCTCC

ACGGAATCTGTCCTATGA

>pR75

ATGAATGAAACAGATGATAATTGGACGGTGTTTACAAAACTTTATTCATGTCAATACAGAGCATCAGATGATTCTCCACA

GTTTACATTATGGCTCGATGGCCCCGTCACAATATCCGCTGCAATTTTCTCTGCCATTGGCACTGTCTATGCTATTGGAT

TCCTTCGAAATGGGCACTTGAATAGAAGAATGTCAGCAGCTTTATATACTTTGTGTCTCATGGATTTTATGCTTACGATG

ACTACCGTTCTTTTCCTAAGCATTGAACCTCTTAGTATTCTCCTATTTCGTACCAATATCTTCTACCAACATCAGGATAT

GATTCTTATTCTGTATGGAATACGGAATAGCTTTGCCATGTCATCGCCGATGCTAGTTTGCTACATCACATACATCCGAT

ACAGAGTTGTCAACAATCCACTCAAATTTGCATCACATTATGGAAGATCGAGAAAATCAATATCTGCAAGCAAAATGAGC

ACAGCTGCTCAGCCATCTTCCACAGAGTCGGCAAAGTTCACAATTTCATTTCCAGCCGAAATGTTCTATGAATTTCACAC

CAGAAGCAATGCTGGAAAAAGAGGAGCAAATTTCAGAAGATTCTTCAGACCATTTCTAGTTCCAATCCTTCTGGTAATCC

TCTGCTTCGTCATTCATAGCACTTCCTACTTTGAATTCAACCTTATTACATGCTTTGATGAAGTTCATCAGACTGAATCA

AAAATGCTCCAGATGACACAACTCAGAGGAGATTCTTATTGGTATTTTCAATTCAAAGTTGCCCTTACCATGACAACAGA

GACATTGGGACCAATGCTTTTCATCTCCACGCTGTCATTGTTCACTGAATACAAAATGCATCAGAATGTCAAGGAACGGA

GGAGGCTTTTTGAGAGTCAGAAGAGAAGTAGAAACACACTGGTGACGGAGGAATTGAAGGATAAAGCATCGAAGGCGTTA

GCTGTGTTCATTGTTGTCAAATTTCTGATTTTGAGATCCCTGCCAACATTGATTGATCTCTACGAGGTTCTCATTGAAAA

CAGTATAAACTTCGGCCCATTCATGACAAAAGTCACGCGAATCTCAGACTTCCTTGTCATTCTCAACTCGGCAACCAACA

CATTGGCCTATTTCGGCAAAGTTCGATTCGAAAAGTGTTTCAGATGGCTGGAACGAAGGATTCGGTGTCGAATTGTGAAG

AAGGAGGCGAAGGAGATTTTGAGCACTTCACTTACTGGATAG

>pR76

ATGAGCGATTCCTCGCAACACCTACGGCTTGCGCTGACGGTCACACATCTGCTGCTCGTCGCTTTGGGATCCGTCAACCT

CATCATTATTCTGTTAATTATCACAAGACCATACCTCCGATCTATAACAAATGTGTACATGATAGGATTATGTCTTGCTG

ATTTCATATATCTAGCAGATTTAATACTCGTTGCCGCCACATCACTAAATGGCAAAAGCTGGCCGTTTGGTCCAACAATT

TGTCATCTTTTCCATGGCACGGAGGCGACGGGGAAATACGCATCTGTACTTTTCGTCGTACTTCTAGCCGCTGATCGATA

CATCGCGATGTGTAAAAGTGATTTGTGCGGCAGATATAGAACTTATCGCACAGCCATCTTCCTGAGCGGTCTTGCGTGGA

TCGCCGCGTTTATCTGCAGTCTTCCTCTCTACGTGTACGCCGATGAGATAAAAGTACGTATGAGACCGAAGAATGGTACA

GATTCGATAGAGAACAATCATACATTGTGTATTGCTCATTGGCCAAGTCCACCACACGCACAGTGGTATATCTCTGTGTG

CTCGGTGATGATCTTCATACTTCCGGGACTTGTGATTTTCTACTGCTATTATCATGTATTTTGTAAATTACGAGAAGCTG

CAAAGGGTTCTAGGAGATTGCATCGTAATAAGAGATCAAGGTCTTCATATCAAAGAGTCACTCGAAGTGTACAACGGGTT

GTATTGTTTCATTTACTTTGTTGGTCCCCATTCTGGTTGTTCAATTTGTTTTCTGCCATCTTCCGGGTTCGAATAACAAC

ACAATTAATGAGAATCATAGTCAACATCATTCACCTCTTCCCTTATGTGAACTGTGCACTTAATCCAGTATTATATGCAT

ACCGAGCAGAAAACTTCCGGACCGCATTCAAATCACTGCTCTTCTGGAGCCGACGGTCAGTTTCTAGCAAATTAAACCGA

CCACTTCCCTTACCGGAAGATGTGCGGAGCACGTCGGCTAGTACGTACTATAGGAACAGCAGCAACTTGGACTGTTTGAA

ATCCTCGTTGCCCATAAAGTCAACAATGGAGCCAGCATCCGTGTACCAACCAGAAGAAGAAAGTCCAGTGGACTCTACTG

AAAACTTGGAAATGGAGAGAAAAAAAGCGGAACTTCAAGCCTGGCGGCCTTATGATGGAACTAACTGTGAAGTATTAGAA

GTTCAATACGATCATCATAAATGCTCCGCCAGAAGTATCCTTCTCCCAGCTGTCACAGTGGACGAAGGAACAAAATTATG

A

>pR77

ATGAGCGATTCCTCGCAACACCTACGGCTTGCGCTGACGGTCACACATCTGCTGCTCGTCGCTTTGGGATCCGTCAACCT

CATCATTATTCTGTTAATTATCACAAGACCATACCTCCGATCTATAACAAATGTGTACATGATAGGATTATGTCTTGCTG

ATTTCATATATCTAGCAGATTTAATACTCGTTGCCGCCACATCACTAAATGGCAAAAGCTGGCCGTTTGGTCCAACAATT

TGTCATCTTTTCCATGGCACGGAGGCGACGGGGAAATACGCATCTGTACTTTTCGTCGTACTTCTAGCCGCTGATCGATA

CATCGCGATGTGTAAAAGTGATTTGTGCGGCAGATATAGAACTTATCGCACAGCCATCTTCCTGAGCGGTCTTGCGTGGA

TCGCCGCGTTTATCTGCAGTCTTCCTCTCTACGTGTACGCCGATGAGATAAAAGTACGTATGAGACCGAAGAATGGTACA

GATTCGATAGAGAACAATCATACATTGTGTATTGCTCATTGGCCAAGTCCACCACACGCACAGTGGTATATCTCTGTGTG

CTCGGTGATGATCTTCATACTTCCGGGACTTGTGATTTTCTACTGCTATTATCATGTATTTTGTAAATTACGAGAAGCTG

CAAAGGGTTCTAGGAGATTGCATCGTAATAAGAGATCAAGGTCTTCATATCAAAGAGTCACTCGAAGTGTACAACGGGTT

GTATTGTTTCATTTACTTTGTTGGTCCCCATTCTGGTTGTTCAATTTGTTTTCTGCCATCTTCCGGGTTCGAATAACAAC

ACAATTAATGAGAATCATAGTCAACATCATTCACCTCTTCCCTTATGTGAACTGTGCACTTAATCCAGTATTATATGCAT

ACCGAGCAGAAAACTTCCGGACCGCATTCAAATCACTGCTCTTCTGGAGCCGACGAAACCGACCACTTCCCTTACCGGAA

GATGTGCGGAGCACGTCGGCTAGTACGTACTATAGGAACAGCAGCAACTTGGACTGTTTGAAATCCTCGTTGCCCATAAA

GTCAACAATGGAGCCAGCATCCGTGTACCAACCAGAAGAAGAAAGTCCAGTGGACTCTACTGAAAACTTGGAAATGGAGA

GAAAAAAAGCGGAACTTCAAGCCTGGCGGCCTTATGATGGAACTAACTGTGAAGTATTAGAAGTTCAATACGATCATCAT

AAATGCTCCGCCAGAAGTATCCTTCTCCCAGCTGTCACAGTGGACGAAGGAACAAAATTATGA

>pR78

ATGAGTACGATTTCACCAGATATTGTGACGAACTCTACAGAAGAAGTAGCAGAATGTTCACGAAATGAAACTTATGACTT

TAATCGATACGCACTTATTGTATACGTTGGAACTCCGATTGCAGTTGCTGGAGTTATTTGCAATTGGATACTATTTAGAC

TATTCACCAAGTCAAAGTCGACGAAATCACCAGCCTTATACTTACTTCTTTTAGCAGTGCTAGATCTTTTAATGGATCTC

CTCTACATTCCCTTTTTCACAGTTGACGCATTAGCAATCTACCACAAAAATGAGTTTCTCTATCATATTTGGCATGATTA

TGCAATGTTTGTATTTGGACTGAGTAGGCTTGTCCAATTTGCATCAACTTACATAATTCTGTGTGCTACTATTGAGCGAT

TTATTGTGGTGGCCGAGATAAACTCACTCGATTTTTTAATCTCGTCAACAGGCCGATTTGTCACTATTGGAATCACCTTC

CTAGGTGTTGCAATTCTGAGGCTCCCAGCATTTTTTGAGTATTTCATCACATACCGTCCCGATTGCCCTCTTTATGAAAA

CTACGACTATACACCCATTCTTGCCAGTTGGGAACATTATCAAATGTTCAATTTTTATGTGATGACGGTTCTTCACATTT

TCGTACCATTTGCTCTTCTACTGATGCTCAATATTTCCATTGTCGTTGTTACGAAAAGGAAATTGACTGGAATTGGATGG

GCGGTCACAACATTTATTGATATGCCAAAAGTTAGTGAGATGATTCGAAAAGAGTCTATCAATTCAAACAAGAGGCGAAG

AGATGAGCTACGGTATGCTACATGGACAATGGTTTCAATTGCAACAACTTACCTTTGTTGTTCGTCTCTTTCCTTGTTTA

TAGGAATATTGGAAAATGTTTGGCCGGAAAACACACTTCTTTTCCAGGAAGATGGATCGAGTACGAAATTTTACACTTTC

GCATCAGACGCTGTCAGTATTTTGGTTGCAGTCAACTCACTTCTTCGAATTGTTGTGTACTCACTTTGCAGCCCCAATTT

TAGAAAACAATTAGTGAAAGAATATCCTTGTTTGAAATGTCTCACCTGTGGAAGTGACGGAGATGAAAAAGCGAAAAAAA

TCAAGGAAGAGAAGAAACTATTTATTGGATATCAGAATTTGATAATGGGTGCCAGGGTGGATATGCTGTGA

>pR79

ATGGAGTTTACCGAATGCAAAACTACATTTATTCATCTGCCCGATAAAAGTTTTTTATACGATGTTTTTGTAAGTGTATA

TAATTTCTACCATCCTATACATGCCTACTTATCAATATTTCTATGCGTGTTGGGTACAATCGCTAATTTCTGTAACATCG

TCGTACTAACGAGACGAACAATGCGAACGCCAGTTAATATGATTTTGACAGCAATGGCATCTTGTGATACAGTTGTGTTA

TTTTCAAATTTAATATACACAACACATTACTCATTTGTCGCTTTCAAGTTTTGTCATCCGAAACATTGGTCCTACTCTTG

GGCGTTATTTTTAATTGCTCATGCTCATCTTTCATTAGTTGCACATTCTTCAAGTGTTTGGTTGTCAGTTATGCTTGCAC

TTGTTCGATATGTAACACTTCGAAGTAGGGGAAATATGGGTGGTATGCAAGTGACGTTAAGGCATTCTTATTATGCCGTT

GCTGTTACTGTATCCCTTGTAGCAGTGCTTAATGCACCGAATTTTCTGAATTACAAAATCAATGAACAGCCATTGAATGA

AACGTGTACCGATTTGGATCCAATGTTCTGGAATTCGCCCGCGTATCTTCCTGGAATTGCAGACATTGCAAAAGCAAACA

GCTGTCTGGTCTTCCGGTTATCTTATTGGATATCCGGTATGGTATTCAAAGTATTACCATGTGCGCTTCTATCGTTGTTT

GTTTGGCTCCTTTTACGAATTCTTCGTGAAGTGCGTGAGAATCGTCAACGTCTTCTCAAGAACTCGCAACATCGACCACC

GAATCAAACGACTACTCGAAACGGACAAAGACTGAGCATTTCAGTTGCAGGCAACGAGAAATTAGGCAGAAATGGAAGCT

TACGGGGGAGAGGAGAACGTGTCGATCGGACAACTCATATGTTATTGGCAATTGTAGCAGTTATGCTAGTGACTGAATTA

CCTCAAGGAATTATGGCTGTCTTGTCTGGAATGTGTTCTGAAGAATTCCGAATTTACATTTATAACAATCTGGGAGACAT

TCTCGATTTGTTCTCACTTTGCGGTTCATGTTGTTCATTCATCATTTACTGCTCAATGAGTGGACAGTTCAGAAATGAAT

TCCACCGTGTCTTTGTACCTGCAAAGGTGAGATGTCTCCGAATGTCATCGCCGTCGATTCGTCGTCCATCCGACGCCTAC

AGTACCACAAAAATGACTTTTCTAAAACCAAACGAGAAAAACGGAAATGGAATGAATGGAAATGGCACTTATTCGGAAGA

TACAAGGTCAGCAAGTGTTAAAATGGTCGGAATCCAGGTACGAAGAAACAGTACGGAAATAACGAGAATGACTGGATGTG

ATTCAATTACTCCATGTTCTCCAATGCCAACATCATTTCCATCATCCCCACTTCCACCGATTCGAAGTGGAGAAGATGAA

TCCACTGATGAGACATCACATCTACTTAACAGCTCAGGACCCAACTCAACAGCCAGTGCTGATGGAATTCGTGGACACTT

TCAAAACATTTGA

>pR80

ATGGAGTTTACCGAATGCAAAACTACATTTATTCATCTGCCCGATAAAAGTTTTTTATACGATGTTTTTGTAAGTGTATA

TAATTTCTACCATCCTATACATGCCTACTTATCAATATTTCTATGCGTGTTGGGTACAATCGCTAATTTCTGTAACATCG

TCGTACTAACGAGACGAACAATGCGAACGCCAGTTAATATGATTTTGACAGCAATGGCATCTTGTGATACAGTTGTGTTA

TTTTCAAATTTAATATACACAACACATTACTCATTTGTCGCTTTCAAGTTTTGTCATCCGAAACATTGGTCCTACTCTTG

GGCGTTATTTTTAATTGCTCATGCTCATCTTTCATTAGTTGCACATTCTTCAAGTGTTTGGTTGTCAGTTATGCTTGCAC

TTGTTCGATATGTAACACTTCGAAGTAGGGGAAATATGGGTGGTATGCAAGTGACGTTAAGGCATTCTTATTATGCCGTT

GCTGTTACTGTATCCCTTGTAGCAGTGCTTAATGCACCGAATTTTCTGAATTACAAAATCAATGAACAGCCATTGAATGA

AACGTGTACCGATTTGGATCCAATGTTCTGGAATTCGCCCGCGTATCTTCCTGGAATTGCAGACATTGCAAAAGCAAACA

GCTGTCTGGTCTTCCGGTTATCTTATTGGATATCCGGTATGGTATTCAAAGTATTACCATGTGCGCTTCTATCGTTGTTT

GTTTGGCTCCTTTTACGAATTCTTCGTGAAGTGCGTGAGAATCGTCAACGTCTTCTCAAGAACTCGCAACATCGACCACC

GAATCAAACGACTACTCGAAACGGACAAAGACTGAGCATTTCAGTTGCAGGCAACGAGAAATTAGGCAGAAATGGAAGCT

TACGGGGGAGAGGAGAACGTGTCGATCGGACAACTCATATGTTATTGGCAATTGTAGCAGTTATGCTAGTGACTGAATTA

CCTCAAGGAATTATGGCTGTCTTGTCTGGAATGTGTTCTGAAGAATTCCGAATTTACATTTATAACAATCTGGGAGACAT

TCTCGATTTGTTCTCACTTTGCGGTTCATGTTGTTCATTCATCATTTACTGCTCAATGAGTGGACAGTTCAGAAATGAAT

TCCACCGTGTCTTTGTACCTGCAAAGGTGAGATGTCTCCGAATGTCATCGCCGTCGATTCGTCGTCCATCCGACGCCTAC

AGTACCACAAAAATGACTTTTCTAAAACCAAACGAGAAAAACGGAAATGGAATGAATGGAAATGGCACTTATTCGGAAGA

TACAAGATGGTCGGAATCCAGGTACGAAGAAACAGTACGGAAATAA

>pR81

ATGTCAAATAACACAACAATACCATCAAAAACAGCGACGGACATCTGTCTGACAGACAGACAGATGTCCTTATCAGTCAG

TTCAACAGAAGGAGTACTTATTGGAACGATTATTCCAATTCTTGTGCTCTTCGGAATTTCCGGAAATATTCTTAATCTCA

CAGTACTTTTAGCTCCAAATTTAAGAACGCGATCAAATCAACTTTTGGCATGCCTTGCAGTTGCTGACATTGTCAGTTTA

GTCGTAATTCTTCCCCATTCAATGGCTCATTATGAAACATTTGAGACGGCGTTATGGTTTCGGAAATTTTATGGAAAATA

TAAATTTCAAATTATTGCAATGACAAATTGGTCAATCGCTACTGCTACATGGCTGGTTTTTGTGATATGTTTAGAAAGAT

TGATCATCATAAAATATCCGTTATCAGTGAGAAAACAGGCAAAGTTTTTTACTCCTCGGAACGTCGTCACAATCATTGTT

GTCACCACTTTCATCCTCACTTCCTACAATCACGTGAGCCATGCTTGTGCTGAAAAGCTCTTTTGTAATGGCACACAATA

CCACGTGGCTTGTTTGGGAATTGATTCCGAACGTTGGTTCCGGAATGAGCCAAATCCCAACTCTGAATTTATGAAGTCTG

TTGTCCGTGTAGCTCCTCAAGTTAATGCAATTTTCGTTGTGCTCATTCCAGTGGTTCTCGTGATTATATTCAATGTCATG

CTTATTTTGACGTTGAGGCAAAGAACAAAGTTATTTGAACCATCAAAAACTATTCGAGGAGATTCACAGTTTACCCAGCT

TCAATCAAAAACAGAACATAAAGTGACAATCACAGTTACAGCAATTGTGACGTGTTTCACAATAACTCAGAGTCCATCAG

CATTTGTCACATTTCTCAGTTCCTACGTCCACAGGGATTGGGTTACACTGAGTGCAATCTGCACAATTCTGGTTGTACTT

GGCAAAGCCTTAAACTTTGTTCTCTTCTGTCTTTCATCTGCATCATTCCGTCAACGTCTTCTCATGCAGACAAAACAAGG

AATTCTTCGGAAATCGACGAGAACCGTCTCAATGGTGACATCATCAACAGTGGTGGTTGACAGTTTGCCAGAATCACGTA

AAAAAAGTCGTGTTGCAATGGAAATGATTGAGAGAAGAACATCAGCTTCTTCGTGTCTTGTCGGCGGAAATCGAGCCGAC

AGAACTCATCGAGCCAACAGCAGTCAGTCAATGATCGGAGAGAGAATGCCTTTGAAGGAGTTTCGTAGAGGAACGAGCTT

CGTGTGA

>pR82

ATGTGGCCGCCTGTCGATGTTCGCAGTCTGACCGCGGAGTCGTTGGGACGTGCGTCTCTCATCCTCGCCGTCGACTTGCT

TCTTTTTGTCACATCGATTCTCAACATCACTGTCATCGCGTTCACCCCGGACTTGTCTGACGTTATCGGATGTTATTTGA

TATCCCTATCAGTGGCGGATTTGCTAACCGCAATCTTCGTCATCCCCCTGAGCATCTATTCAACTTTGGAAGGCAACTGG

CGAATCGGTGGCGACAATTCGATCATTTGCAAAAGTGCGGCCTATCTCCAAATTGCACTTTTCTGCTCCACCGTTTACAC

TTTCGCATGGATATGTATCGATAGATACTCTGCAATGATGAAGCCATCACGATATAGCGAGCAGTCACTAACTAGGTGCA

AGTGTTGGATAGTGTTTAGTTGGTTAACTTCAATGCTTCTATGCTGCCCAATAATTGTTGCAAGAATGCAGGTTGTGTTC

TATCCCGACGCGCAGTTATGTGTCCTGGACTGGTCCGCAACATCTGCATACAGTGTGACCTTACTCCTACTTGTCTTTGT

ACCAACTTTAGTAACCGTCTTTAACACAGGCTGGAAAATCTGGTCAGCTATGAGAAATCCTGCAGCGCTTGATGATAGTC

AAAGAATGCTCGTGGAAACAGACCCAAACTTTGTATTAACCGGTTTTCTGTTTGTCACATTTTTCTTGTCATGGCTTCCA

TTGATTACATTGAAGGTGTGCGAATTAATATGGGGCCCATTTGATATGGATTTATCAATGCTCACATTCTTCTTCGTGTG

GTTAGCGGTGTCTGGTCCGTGTTGCAAGTTTTTGATCTACATGTTCACAAATCATCAGTTCCGAAGGTCCTTCCTTTCCT

ACATTTCATGCTCAATGTGCTGTTCGAGCCGGAGTCGCTATGAATATCATGACATAGGCAACAATTCTTTCCTTTGA

>pR83

ATGGAGGAAACGGGGTTTCAATTTGACGTCTACATGTATCTTCTGCCTCTGATCGTCATTTTTGGATTGTCTGGGAACGT

CATTTCTCTCGTTACCATATTTCATTCAAGACTAAGGAGGGTGAATGCTAATATCTATTTGATAGTTCTAACATTAGCTG

ATAGCATATTCTTGACTGGAATCCTTCTGATTTGTTTCAAAGTTGATTGGATTGCGTACGAATATTGTGTTGGTCTAGAA

TACGTTCTCATGACGGCTTCATATATCTCGAGTTGGTCAACAGCAGCACTAACTATTGAAAGATATTTGGCTATTGCTCA

TCCATTGTCTCATATGAAGTATGGTCACGTGGATCGAGCAAAAGTGATGCTTTACTGGGTTCCAATTCCTTTTGTTCTAC

AACTGTTTCAATTTTTCTCACTAGAGCCAGCAACCGAAGAAAGGAAATGTGCATTAAAAGAAGCAAACTATCAAATCATC

GCTCAAGCTGTAGATACAGTACTTTGCTACGTGGTACCATGTGGAATAATTGTTGTTCTGAACATTCTTGTTGCACTACA

AGTACAAAAATCACAAGAACATTTCATGGCAGAAACTAAAAAGAGCAATTCAAGAAGAACTGGAGGATCATCATCATCAT

CTGGTACATGGACCAGAATTCTTTGGGTTATGCCATTAGTGTTCGTAGTATTGAACACCCCATTCTATGTAATTATGATG

ATTGAAATTGTTTTTCAAATTATCTACCAGTCACCCCCATCAGGAGATACAAGATCAGAACTTTTTGTGACTGTTTATAA

TACTGCTCATTACATGTACTACATGAACACGGCAATTGATGTATTGGTCTACGCATTTTCCAGTGCAAATTTCCGCAAAA

CTGCTGTCATTGCTTGGAAGCGAATTCTTTGTCCAGGATACGCAGAACGGCAAAAGGGAAAGGTACTCGTAACGGATCAA

ACTTCGAGAATCTCCTATCGTATGACGTCTGAAAGAAGTACAATCCGTTTCCCAAACACTCCTTCTAGAGCCAATTCACC

AAAGTTAACTGCTTCAAATGCGGTTTCTGTTCCATTACTTGGAACCGAAATTCAAGGATCATCAAAAGAATCAACTGCAA

TTTAA

>pR84

ATGGAGGAAACGGGGTTTCAATTTGACGTCTACATGTATCTTCTGCCTCTGATCGTCATTTTTGGATTGTCTGGGAACGT

CATTTCTCTCGTTACCATATTTCATTCAAGACTAAGGAGGGTGAATGCTAATATCTATTTGATAGTTCTAACATTAGCTG

ATAGCATATTCTTGACTGGAATCCTTCTGATTTGTTTCAAAGTTGATTGGATTGCGTACGAATATTGTGTTGGTCTAGAA

TACGTTCTCATGACGGCTTCATATATCTCGAGTTGGTCAACAGCAGCACTAACTATTGAAAGATATTTGGCTATTGCTCA

TCCATTGTCTCATATGAAGTATGGTCACGTGGATCGAGCAAAAGTGATGCTTTACTGGGTTCCAATTCCTTTTGTTCTAC

AACTGTTTCAATTTTTCTCACTAGAGCCAGCAACCGAAGAAAGGAAATGTGCATTAAAAGAAGCAAACTATCAAATCATC

GCTCAAGCTGTAGATACAGTACTTTGCTACGTGGTACCATGTGGAATAATTGTTGTTCTGAACATTCTTGTTGCACTACA

AGTACAAAAATCACAAGAACATTTCATGGCAGAAACTAAAAAGAGCAATTCAAGAAGAACTGGAGGATCATCATCATCAT

CTGGTACATGGACCAGAATTCTTTGGGTTATGCCATTAGTGTTCGTAGTATTGAACACCCCATTCTATGTAATTATGATG

ATTGAAATTGTTTTTCAAATTATCTACCAGTCACCCCCATCAGGAGATACAAGATCAGAACTTTTTGTGACTGTTTATAA

TACTGCTCATTACATGTACTACATGAACACGGCAATTGATGTATTGGTCTACGCATTTTCCAGTGCAAATTTCCGCAAAA

CTGCTGTCATTGCTTGGAAGCGAATTCTTTGTCCAGGATACGCAGAACGGCAAAAGGGAAAGGTAATAACTTCTCACTAG

>pR85

ATGGCACAACGGAAAAGGTCACAAATGACTACTGATATTATATCAGTAGCTCAACATGTTATCACAACGTCGTACATAAC

TGTTATCTTATTTGGAAATATTGGAAATGTGTGGGTCTCCTGGAAGGTCGGAATCGTATTTGTATTTGATAAATCATCTC

TGGTACCCCGAAACATTGTGATGCTCATTTTGTTCATTTGCGTCGCCGATCTCCTGGTACTTCTACACCTCACACTTTTC

GTGCACTTCCAGTTCGAGAAGCAGTGGATTTTTGGGAACATCGTCTGTAAGTCATTCTACTGCATCGAGGTGGTCAACAA

ACTCATCATCCCATTGGCTCTTCTCCGTATCAGCAGGGAGTCTTACGAATCTGTTCGAATGACGTCACACAAATGCTCCA

AAAAGAGGCGGAGCATATGGAATATTGTATTTCAGCTGTACTCTGTTATGGCGTTGATTGCAGTCGGAGTTTTGACGATA

TCCGTACTAATTTTTGCGGAAACTCGAACTTACGAAATTCCACGACACGGAGAGCTCACTGCAGTCACCATGTGCATTTT

CCACCCTCCTCATCCATATGCAATGGTGTTCAATATCTTAGCGTTTGTGTTTGGCTACGGATTCACTAGCGCTGCCTACA

TGTACTTTTATTTGAGGGTCCCCATGATCTTGAAGAGGAGATATTCGTCTATTAAAACTACTAGTAGTAATTCTACACGA

TTTAATCACCAATCAATACTAAAAATCCGCCATACCGTAAACGCTTTCGTGGTGGTCTATATGGCCTGTTGGACCCCCTA

CTGGGCTCTTTTCTGGCTTTTCTTCACATTTCCCCCGTCGAATAATTGGATGGTGATCATATCTCAAATGGCTCACCTGC

TACCGTACATCTCGTGTGCCGCCTATCCGATTATCCTAACTTTTATCAATAAGGGAATCCGCAACGCCCACTCGACTATT

ATGACGAGCCAGAGGAAGAAGCTGATGACTATAAGAGCGGGGGCTTATCAGATGATAACTGCACAGCTCGGGTTTGTGCA

AACTTGGCTCCGCGACGACAATACTCTTGCTCCGCGGTGGAGCTCCTCCGGCAGTGTAGTGCCTACCGAAATCCACGTGG

AAAGTGTGGTGGAGGACGAGGAGACGGAGCCGGCGGCTCCGCAAATTTACTTGTAA

>pR86

ATGAGCTATTCCAATGAAAATCTGACGCTAGAAGAGCTTCAAGAGGCACATGCACTCAAGGAAGCCGAGGAGATTTCAGA

GATTTTAAGAATCAAGATCGAAGTATGCATCTACGTAATCGTATTCTTCATTGGTGGCCCATTGAATCTCATGGCTCTCT

CCAGATCTCTCCGTGCCTTCTCAAGAACTCACAAAGCGAAAAGCCAAATCCTCCTTCTCCGAATAACGCTTAATCTAGCC

GATTTGATGACTCTCTTCCTTTATGTCCCAAAACAAATCATCTGGCTTATCACATATCAATGGTATGGCGGAGAGTTTTT

GTGCAGAGCCTGTGCCTTCTTCTCCACGTTCTCCTTTTATCTCAACTCTTTTGTGATCGCGTGTATTGCAATTGATCGGG

TGTTTGGAGCCTATAACATAAGCTCTTTGAATGCTCATCGGAAAGCTTACATCAGATGTCGGAACCTTCTTGGATGCGGA

TGGGTTTTGGCGTTCTTGTTGTCACTTCCACAGGCGGTGGTGTTCACAACGACTAGCCCTTATGAGAATATTGATTTCAA

ACAATGCGCCACAATCTTCATGATCTTCCTTCACGAGAAGCGTATCGAGTACCACGATCCTGCGACCACAGACATCCGAC

GAGAACAAATCGACGAAGAAGCCGCTTCGATGGAACGATGGGAGAAGATGTACGCAATCAATCATTTTCTATTCGTCTTC

TGGATCCCGTGTATCGTCATCATCGGCAGTTATATAGTTGTGTTACTCATCCTTCAAGGTCATCTCAAGCAAGACAAGGC

ATCAACATCTTGCTTCAATTTCCGAGTCAAATCATCTGACATTATGACTATCGATGAGACTCAGTCGACTACGACGAGAT

CCCCGACAAGGTCGACTTATCTACTTCAACAGGAAGAATCATTTGCTCTGAGATCCTCGTCATCTTACGACTTCGAATCT

CATAACTCCCACCCAACGCGTCAAGGATCCATAAGATCACGAGCTTCCACTGGTGGAGGATCAAAGTTCGGAGCAATGGC

AGTGAGCACTATCCACAGAGCCAAGCAACACGCGAAACGGCAAGCTGCTTTGATAATCGTGGCTTATCTGTGCATCTGGA

GCCCATATAATCTGTTCACAGTGCTTAATTTGTTCGGGATGCCGATTATGGAAGAGCTCCGACATTTTCTCAGTTTTCTG

AATGCGGCAATTTGTGTGAACACTGTAGTGAATCCAATTATTTATGGTGTATTCTCAATGCCACATAAATAA

>pR87

ATGTGTGTTTCCGAAGAATTCGGGCCCCGGTTTTTCGGTTGTTTACTGAATTTATGGCTTTTACCTCTTGTCTGTGGGAT

CGGACTTATTCTAAATATTTGCTGCCTAATAGTATTTTTCTCTACAAGAGGTCATCCACTTGTACCCGCACTTATATTTT

TGTCATTTTGCGACTGTCTTCAATTATTCTTCTCGTTTTTAGTATTATTTCTGCCAGCTCTGCACGATTTCTCACAGTCG

ACACCTTATTCAGCTCTAGGACAACTTGCATATTTATCCACCGGTCTCTTATCACCACTACTGCTTACATTTAATTGTGC

AAGTATATGGACAATTTGTTTTATTTCAATTCAGAGGCATCGGGCTATTCTTCGTCCTCTTTCAAGTATATCTTCTCCAT

CAAAACCATTCAAGCCATTATTATTTATTTCATTAATGGCTTTAATGTTTAATGGTTGCAAATGGGCAGAATTTCGGTGG

TTTTGGTCACATGATCCAACGAATTCTAGTGACTATCAGTATATTCTGGCACACGAGCCTTCGGATTTAGCAAGAAATGA

AAATTATCATAGGGTGTTGGATAATCTTCTCTACCCATTATTAGTCTACCTGGTTCCTCTTCTCCTTTTATCTGTTTTGA

ATTTTCGAATTTTATCAAATATTTCGAATCGAAGAGTATCATTCGATCATAAGTCACGATTTGCACAGGAAAGACGTTCT

GTAACACTTTTGATATCAATTGTGACAATGTTCTTCCTGTGTCATACGGGTGGTCTTGCATATCGATTCGTGGATCAAGA

AAAATATAATAATTCGGAATTATTTGTTCTTTTGAAGGATGTCATCAATTTATTATTCAATGTGTACTCATTCACAAATC

CATTGCTATATTTTGTTTTTACCCGCCAATTTCGTGATCTTCGAACCATGTGGAATTCACACTTTGCATCGCCTGCTTCT

CGATCCGAGAGCTATGCAATAACGAACTCGAAGCCAAATTCAAAACATCGGCATCTCTCAATGCCATCAATTTATCGTAC

ACCTTGCGTTAGCGGAAAATTGATTTCATGA

>pR88

ATGGCATGCTTGAAAAAAGCACAACTTTTATTTTTTAGAGACCCCCTGAGTCGATTCATTTGGCAATACGTATTCCCACT

CCAATTCGCGTTAGGACTCCTTGGAAATTCTCTGAATTTGTGGGTGCTCGCCAGTGATGAAGTACCCAATATTGCTAGTG

ATATGCTAGCAGCAGTTTCATTTTGTGATTTGGCATTTTTACTGGTTATGTTACCCAATTGTCTGGCCGCATTTGACACA

ATTGCATACAATGTCTCGTTCAGGTATTTCTACCTGAACACCAAACAACACATGTCAGCAGTGGCGAACTGGATGAGTGC

CGCTGCGATTTGGCTAATCCTAGCAGTATCCATCGAACGCTACCTGATCGTCCGATCGCCGTTCCGCGCCAAACTCTATT

GGCAACGTGGAAAAATGGTGGTGGTCCTGTCGTCGATCTTCGTGACAACCGGGTTGCTCACAGTATACCACCATTTTGAA

TGGGACTGTGTGATCGAGGAGTTCTGTAACGGTACACAGCTTCTCGACTTTTGCTACTACTCTGGGATGCGGCACCAGAA

AACGTATCGGAACAAAGACTGGGTGACGCCAAGCAATGTGAAGAAGGTCTACCTGAAGTTTTCCACGTTTTTGAACGCGA

TTTTGGTTGTATTCGTCCCAATAGTTTTGGTCATAGTTCTGAATATGCTTATGATTCGGCAGTTGAAGATAAACTGGACA

TCACCAAGATTGGAGTCGATGAGAGGAGCCATGACTAGGAGCCACGTGCGTCAACGTCAACGAGTAGCCGTTACAGTCAT

CGCCATCGGGCTCTGTTTTTCTCTAACCCAAGGACCATCTGCATTCATGGCTCTTTACGAACTTTTTTCCGAATACGAAC

TCGGGCATACTTTCTACGCGATTTTTAGCATAACGAATTCGCTGGTGGTCACTGGTAAAACGATCAACTTTATTTTGTTC

TGCCTGTCATCGGAACATTTCCGGCAGAAATGTTTCAAGGCGATCTATCGGAAATTTCCAAAGCTCTCCCAAAGCAGTAT

CGGCAAGCGATTTCTCGACCATCAACGCTCCAACTCGTACGATCGATCCTCCTCGCTGCGCACCGTCCGCTCACGGCGTG

GCTCTGCAACTACACTTCCAGGTGTCAACCGAGACACCTTGCTGCTCCCGCCATCCGAAGAATGCGCGACGTGCTCACTT

TCAGTGTCGAGACAGAACATCTCATATCGGACAAGGTTGCACTCGGAGCCTTCGGTCTCATCCGAACGTTAG

>pR89

ATGCTAGAAGAACAGAATCTGACATCGACATTGGCTCCTCCGGCGCTGTCGCCACTGTTACAAAAGTGCTTCGACGAGTC

GTACTTGTTGTTGATGCCCGACATCGAGAAAAGAATTAATAATTACATCTTCCCGATCCAGTTCATGATCACAGTGGTGG

GAAACCTGCTGACAATGACAGTACTTCTCAGTGGACATATTAAGAATAGAGCCAATCACCTGCTGACAAGCCTTGCTCTC

TGCGACATGCTCGTCTTCTTCATGATGATCCCGCACTACCTCGCCTCACTGGACACTTTCTCTCGAAATCCCACATTCCG

TCTATTCCAATTCCACTCGAAACCTCATTTCGGCGCGTTGAGTAACTGGTTTTCAGCTGCAGCTATTTGGTTTGTTTTGG

CAGTTTCACTCGAGCGACTCTTGATCATCAAATTCCCATTTCGATCATTGGACGTGCACAATGGAAAACAAATAGTCGTC

GTATCGGTAGGGATCATGGTGTCCACCCTATTCCTCACCAGCTATCATCACTTCTCCCATACCTGCATCACCTATATCGC

GTGCCATGGCACTCAGATCCTTGGCAAGTGCTATCCGAACACTGAGGACATGCACGGCAGGAAGATCAACCCGACGAGCC

TATCCGTGAAGCAGTACCTTCATACTTCTGTGTACGCAAATGCGCTGCTCGCCGTGCTTCTGCCGATTTTCGCTGTGGCA

GTGCTCAATATTTCACTGATTCGTTTGGTAAAGCGGAGGCATTCTGAGGAACTTCTCGTCCGAAACGCTGCCGGACCATC

GAGTATGGCCGAGCAGGAGAAGAAGATGACGCACACCGTGCTGGCGATCGTCTCATGCTTCACACTGACACAGGGACCGT

CGGCCATAGTTTTCATTTCACAAAAGATGTTCCCACCAAATCAATATTCCATCTATATATCCGTGGTGGCGAATCAACTA

GTGCTCACCGGCAAAATGCTCAACGTAGTCCTCTTCTGCCTCACCTCAGAGACCTTCCGCCGCCGCCTCTGGCTAACCTG

CAGGTTTTGGTGCCAGCTCATCTTCTACGCGGGTCGCAACAAAACGTCGAGGTCGAACTTCACGCGTTCAAAGTCGGTCA

TCACTCAAAAAACATCGATTGTCTACTCTCCATGCTCCCGAAACTTTTCTACATCCCGAAAGGTCAGCAGTTCCAGTGTA

GTCGCGCCGAAAGGGGCTAGTCAGCTGATCTCCAATCCTTCTACGGATAGCTTCCCGACCCCGCCGACGAGCACGGCAAG

GACCACGCAGAGCACACAAGTTTAA

pR90

ATGCTAGAAGAACAGAATCTGACATCGACATTGGCTCCTCCGGCGCTGTCGCCACTGTTACAAAAGTGCTTCGACGAGTC

GTACTTGTTGTTGATGCCCGACATCGAGAAAAGAATTAATAATTACATCTTCCCGATCCAGTTCATGATCACAGTGGTGG

GAAACCTGCTGACAATGACAGTACTTCTCAGTGGACATATTAAGAATAGAGCCAATCACCTGCTGACAAGCCTTGCTCTC

TGCGACATGCTCGTCTTCTTCATGATGATCCCGCACTACCTCGCCTCACTGGACACTTTCTCTCGAAATCCCACATTCCG

TCTATTCCAATTCCACTCGAAACCTCATTTCGGCGCGTTGAGTAACTGGTTTTCAGCTGCAGCTATTTGGTTTGTTTTGG

CAGTTTCACTCGAGCGACTCTTGATCATCAAATTCCCATTTCGATCATTGGACGTGCACAATGGAAAACAAATAGTCGTC

GTATCGGTAGGGATCATGGTGTCCACCCTATTCCTCACCAGCTATCATCACTTCTCCCATACCTGCATCACCTATATCGC

GTGCCATGGCACTCAGATCCTTGGCAAGTGCTATCCGAACACTGAGGACATGCACGGCAGGAAGATCAACCCGACGAGCC

TATCCGTGAAGCAGTACCTTCATACTTCTGTGTACGCAAATGCGCTGCTCGCCGTGCTTCTGCCGATTTTCGCTGTGGCA

GTGCTCAATATTTCACTGATTCGTTTGGTAAAGCGGAGGCATTCTGAGGAACTTCTCGTCCGAAACGCTGCCGGACCATC

GAGTATGGCCGAGCAGGAGAAGAAGATGACGCACACCGTGCTGGCGATCGTCTCATGCTTCACACTGACACAGGGACCGT

CGGCCATAGTTTTCATTTCACAAAAGATGTTCCCACCAAATCAATATTCCATCTATATATCCGTGGTGGCGAATCAACTA

GTGCTCACCGGCAAAATGCTCAACGTAGTCCTCTTCTGCCTCACCTCAGAGACCTTCCGCCGCCGCCTCTGGCTAACCTG

CAGGTTTTGGTGCCAGCTCATCTTCTACGCGGGTCGCAACAAAACGTCGAGGTTTTCAGGTCGAACTTCACGCGTTCAAA

GTCGGTCATCACTCAAAAAACATCGATTGTCTACTCTCCATGCTCCCGAAACTTTTCTACATCCCGAAAGGTCAGCAGTT

CCAGTGTAG

>pR91

ATGACTTCAGAGCCTGATTTGCATTTCCAGGACCTCTACGATAAAATTGATCCCCCTTTTCAAAACGCGACACCCTTCGC

AAGAACAGGACAATTCTACGAAAATCGCTCGATTACCGATTTTAATAATACAACTCAAATGATAGATCAATGTCCCTTCA

AATTCTCACAGCACTTGGCGCTCGCTACAATTCCTCTTTTCATCGTTGGCCTAATTTTGACAGGAATATCAATTTATGTT

TATACCAGGCCATATCAGTTGAAAACGACTGTTGGTTTTCTGCTGGCTTGGTTATCCGTATTTGAATGGTTGTTTCTTTT

GGCATCTTTTCAAGTCTTTTCAATTTCCGAAACCGCATTTTTTGCATGCCCAACAACTGAAAATGAGTATCCAAAAATAA

AAGCATACAGTGAAATATACATCTTTCCATTAACTCATGCAACAAAGTACATGTGTGTTTGTATATCAATTATCATCTCT

ATTCAACAATGGATTGCTGTCTTCTTACCAATGGAGGCCAAAGATCTGTGCAACATGGAAAGAACAAAACGGTTTTTAGT

GGGTTCTCTAGTTATCTCATCAATTCTTTCATTACCAAAGTTTTTCGAAAAAGGAATACACGAGATAAATGGATGGCCCT

GGCAAACAAATGCCGTTGAGAAGTATACATTCTTGAAAGTGGTACAATACGTGGAAGAAAACGTGTTCAGTGTATTCAAA

ATATTCTTTTTGCTCTTTATTATTAATGTGTCGATTATATTGCTACTGAAATATTCTCACAACGAAAGAATTCGAATGAC

ATCACAGTCAATTAGAGATCCCCGTACATCTATTATGCTTCTGGCTGTACTTTTTGCATACGTTGTCACTCATCTGCCAA

GATTTTTGCTTCAGTACGATGGGCATTGGAAACATTATTTACCGAGCGATATGTATATTGCCCGGAATGATCTATCCAAT

TATTTCATGTGCTGCCATCCTTTGTTCTCATTTGCCGATTATTTAATCTTCTCTGAGAAGTATAGAAAAAGTGTTAAATC

AATTTTAAAGCGACAGCCAAGTGGAAACTCTTCAAGGAAAGCGCCTCAATCTGTTTGA

>pR92

ATGTCGTTAAATACAACTGCTTCCTCATCTTTAAACTCACACAGTTATGATATGGTTGAAGTGTTTGCTATGATTATGTT

GCCATTAATACTAATTGGGATATTCGGAAACATTATATCCATCTATGTGTACAGCAGACATCATATGAACAAAAACACGA

TTGGTTTCCTTCTTCTCAGTCTATCAACAATTGATCTCATTGTTCTTATCACCGCAATGCCATCGTTTGGAACCTATAAA

TTTCCATTTTTCCCCGGTTACCATGAAATTGGTTCCGTTCACACAATTTTTAGTGCATTCTGTTTAATCTATTTTTATCC

ACTTATGTGCATGGGAAAAATGATGGGTCAATACATCATTGTGCTAATTTCTGTGGAAAGATGGTTTGCAGTATGTAGGC

CCCTACAAGTACAAATTTGGTGCACACAGAAGAATACTATACGGGCTATGACCTGCATAATCGCAATTTCAATTTTGTTC

AATGCTCCTAGGTTCTTCGAGTTCAAGGCCAATCTAATCACAGGTGTCATCGGTTTTGGTCTAGCACATATTTCAAAAAA

TGAATGGTATTTCTACTTATATTATGGAATTCGTGCAATAATTTTTGACGCATTGATACCGTTTTCTATCATTGCTGTAA

CAAATATTCAAGTTATCCAGCAGCTTCGAAAATCAAACGAAGAAAGAAAATTGATGACAACGCAACAACAAAAAGATAAC

AAAACTACCACAATGCTTTTGGTTATGATATTTATGTATGCGATTTGTCATTTTTTATGTACAAGTGTCAAGTTTATCAA

TTTGTTCTCCCATAATTATGTGCAATTTCAGTTCGTTATATTTAAAATCATTCATCACATATCGAATGTACTTCTAGTCT

TTTACTCTGCCTCAACATTCTTCATTTATCTCATATTCTCTGAAAAATATCGAAATGTTCTCTCTACATGTGTTACTTGC

AGAAATACTGATGAATTAACAGTATCAAGTCGAGGAAGACAGAATACAAAAACTACCCGATCAAATTCTCAAAATGGTTA

TACAGCGATCAAAAATGAGAAGACATAA

>pR93

ATGCTCGTCGTGAAGACCTATCTGCATCCCGAGCAGGCGATCCATTCCACCTCAATCTACCTAGCAGCGTTAGCGTTCTC

CGACTTTTTTTTGGTGCTTACCGCGATGTTCCTATTTGTGTTGGAGGCCTGGAGGCATCATGATTATCCGACTCTGGCTT

ACCTCTACGTCATCGGAGCACCTATCGTCTTCCCGGTGGCAGCTGTCTTCCAGACGAGCTCTGTCTACTTCTGTGTAGCT

GCTGCAGTTGACTGCTTCATCATGGTAGTTCTACCCGAAAGTGTCAAGCAGCTGTACTGCACGCCGAGACGGGCAAAAAT

CACCTGCGTAGTGCTCATGCTCATCTGCTTCATCTACAACATCCCGCACTTCTTCGAGCTGGAAAAAGTGGACTGTCTAG

ATGAAGACGGTCGGGATTCCATGCAAATCTGCCCGACAGACATCCGGCTTGATCCAGCCTATTACGCGATCTACTACACC

TACATGTACACCACATTCCTGGCGATCGGCCCGCTGACTCTTCTGATCTTGCTGAACATCTGTGTGGTGTTCACTGTAGT

TACCAAGGGCAGCTCTAATGAGAATGGAGAAGACGACACAATATCCCTGATCCTCGTGGTGTTCTTCTTCATTTTTTGCA

ACTTCACCGCGTTGATGGTCAATTTTATGGAGATTATCCTAAATGATCCGTCAATGCTGGTGTACTTCGTGGATCTCTCC

AATCTGCTGGTAGTGGTGAATGGGACGGCCAACTTCTTTTGCTATCTGATCTTTGGTACAAGCTTCCGGGCTACTCTCAA

GAAGGTAGTGCTGGGCTCCCCGGCAAAGCGATCGGCGGTGCTGTGGATCAACGACGAGGAGCACAAGAATCAGCAGTCGC

ATGCCTTGATTTGA

>pR94

ATGTCCCAATGCACAGTGGAATACAATGTGTCTGAAATCACTGAATATGTATTGTCTACTCTAGGTGAACGGTGCCAATC

GGCAGGAATAGTAATTCCAACAGTCATTATTTATGGAACTATTTTTCTACTTGGATTATTCGGAAATATTTGCACATGTA

TCGTAATTGCAGCAAACAAATCTATGCACAATCCAACAAATTACTATTTATTCAGTTTGGCGGTTTCAGATATTATTGCC

TTGATATTAGGGCTCCCCATGGAGTTCTACCAATCCCTTGATTACTCATATCCATATCGATTCAGTGAGGGAATTTGTAA

AGCTCGGGCTTTTCTCATTGAGTTCACTTCATATGCATCAATAATGATTATTTGTTGTTTTTCATTTGAGCGGTGGCTGG

CAATTTGCCATCCACTTCGTTCAAAAATATTCTCAACTCTCTGGAGAGCCAATGTTCTTATTATTCTTGCTTGGACTATT

TCATTCGTTTGCGCTCTACCAATTGCATTCATAGTTCAAATCAACAAACTCCCCCTTCCAGAAGACGCAAAATATCAACC

ATGGACTAACAAGGTCTCAACGGATGGAATATTCGTACTTCACACCGAATTCTGTGCAATGAACCAAAGTCGTCCGGATC

AACAAAAAATGATTATTATATTTGCATTCACTGTGTTTTTTGTTATTCCAGCGATTGCGATCGTTATTATGTACGCGCAT

ATTGCCGTTCAACTAGAATCATCAGAAATTGACTTGAAAGGAGATAAAATGGTAAAGAAAAGAAGGAACAAAAGCAATAG

GACAGTTCTGAAGATGTTGTTATCGGTTGTCATCACATTTTTCATTTGCTGGCTTCCATTCCATATTCAGCGACTTTTAT

CTGTCTATACGACGTGGTCCGAAACCACGACAATATCACCTCCGGTGCAATTCCTATCTATGATTGTCTTCTACATCTCA

GGATTTTGCTATTATTCAAACTCTGCTGCGAACCCAATTCTGTACAACATTCTGTCTCAAAAGTATCGAAGTGCATTTTG

TCGGACCATTCTAGGGGACCATATTGCTAATTTTGTGTTTAAAGGTCATCAGAGACCAGGGCAGTCAAAAAGATGCTCTT

CCTCAACAGAAGCCGAACAAAGAACTCTCATGACACGTGGAAGTGTACGTGTAGATAAGCCCCATCGTAAATTGGAAGTA

CACAACTACTAA

>pR95

ATGTCATTGGACGATGAGTTCAATAAATGTCAAGAAATCGAGTTTACACACGCACAGTTCCTATTTCGATCGTACCTTTT

TCCATTTGTCTATTTATTTGGAATCATATCAAACTCCATTAACATATGTGTCTTCTCACAGAAATCTATGCGGAATCATA

CAGTTAATTGGTTTTTCCTAGCACTTTCCTTCTCCGACCTTTTGACACTTGTCGCATCAATCTTCGTGTTCTCAGTTCCA

GTTTATGCTGAAAACTCCCACAATCCAGAGTACATCGACTTATCTGTTCAATTGATTGTATGGTTCTACCCACTTGCACA

AATTGGACTGACAATGTCTGTTTATGTTACAATTCTTGTTTCAGTACATAGATATCTTGGAGTTTGTCACCCTTTTTTGA

TCCGTCGAATCTCCAATTCAAATGCAGTGAAAGCAGTTATAGTGGCTGCAATTGTTTTTGCATTTGTATTCAATGCGAGT

AGATGGTTTGAACTTCATGCTCAGCCATGTTCATTTGGAACTGAAGGACAGACAAACTCGTCTGTTGTTTATCCAACATC

TCTGATGATGAATCGTATGTATACTCTTATTTTCCGAAATGCAGCTTATACAATTGTCATGTTCTTCCTACCATTTGCTA

TTTTAACTTATGTTAATCTTCGAATAATTGCAACACTCAAACAGTCATATAAAATGAGAAAAGCGATGACAACATCTCGA

TCAAAAAGAAGTGATTCTACAGTACCCACTGACACAATCGTTACAAAAATTGATGGATATTCGGCAGTTGTTCCAGTTGA

AAACGGAGAAAAAGGAAATGGAATTCTCATGGGTGGAGCTAACAATGGAAGTGTGAAAAATGACAAAAAAGAAAATGGAG

TGACTGTAATGCTTGTGGCAATCACAACCGAATTTTTACTCTTCAACCTGATTGCATTTGCCACAAATATTATTGAATTA

TCAAGTATCAGATTTTTCGCTGATCTAGAAACTCTATTGGTGGAGCTATCAACATTTCTGGTGAACGTCAACGGGGCGTC

GACAATTATAATTTATTTAATTTTTGGAAGCAAGTATCGCAATGTGTTCATTCGACTTTTTCGAAAAAATTTGGGGCACA

ATTTTTGTGGTAAATCCCCAAAACGTCCGATTGGGTATTCTCAAGCTCATTTCGAGACAACTCAACTTCTGGATAATTCC

AGATTTGATTTAAATAGAAGCAAGCAAGCGAGCGTTATTCGAAAATCCGATCTTTCACCTTCCACTCGATCATCTTCCTC

GAGAAGACCAACGAATTCATGA

>pR96

ATGAGGGACCGATTCTCGATAGGTCAAGCCATTGCAAATGCAATTGCCAAACGACAAATGCAAACTAGTGAGGTGGAGAA

CATCTTCAATCGCTGCAAGTCAATTGAATGCGTGTATAGAGAAGTTATGGAGACAGATTACGAAGGTGGCATGCCAAATT

GTTCGTATCGATATCCATTTCCACCGGAAGAAGACACAATATCCGCCTACGCATTGCTAATTGATGGTCCTTGCACATGT

GTAGCCGCGTTGCTCTCAATGATCGGAGCCCGATATGCCATCAAATTCCTCTGCCGCGCTGGCCTCAACAAAGAACTCAC

TGCCGCTCTATTCGCCCTTTGTGCCATCGACTCGTTGCTCCTGCTCACCGTCTTTCTCTTCTACGGAATCGAGGCGATGT

CGTTGTTGTTCTTTCGCACAAATATTATGTACGATAAACAAGACTTTACCTATAATTTGCACGGGATCGCGAGCTCACTG

ACCACCGCCTCGACAGCACTAGTGATCTACATCACTTTTCTAAGGTTCATGGTTGTGGTGCAGCCTATGAGATTCGCTAC

AAGCATGAACTCCAACCATCGGAGGACTGGAAGCAAGGCTGTTCGGAAGGGATCGGCTCAGTTGGAGGACTCGGTCACCA

ATAATACCATATTCTCGAGAGGAACCTTCAATAGTTCATCGATCAAACGACATTACAATATTCGAGAGCTAATTCGCCCT

TTTTATATTCCATTTCTCCTAATTCTCATCAGCTTCTCCGTCAACATACCGATTTTCTTCGAGTTTACCACCACAAAATG

TTTCAACGTTGAGCATAATGTCGAGGCGACCAATCCCGAGCCAACAGTCTTTCGATCTTCGTTTGCCAAGTACAAGGCAG

TGCTGATGACGTTGACGCAGACCATCGGGCCGGTATCGATTATTCTTGTCCTATCGATTTTCACAGAGTACAAAGTGCAT

GTCAGTCTGAAGGCTAGAAGAAAGTTATTCGAGAGCCAGCAAAGAAGTCGGTCGGTAGTTCTCAGCGAAGAGTTGAAGGA

GACTGTGTCGAGAACTGTCGCAGTGTTTATTGCAGTCAAGTTTTTGATTTTTAGAACCATGCCGATCTTCTTCGACGTAT

ACGAGAATCTGTATGGAATCGACGAGTGGGGTACTGTAATGAGCATAATGGTGCGAGTATCCGACTTTGGAATTGTGCTC

AACACTGCCACAAACTCGTTGGCATATTTTGGAAAGAAAAGATGGATCGAAAAACGTCTTCGCCTCCGATTGATGAAAAA

AGAAGAAAAACGAAACGCAAAAGCCACCCTAACAGTGCAGGGAAGCTCGTTGAACTCCTCGATGAAGAAGTCACTGCCTC

CGAAAATTGGGTTGCCACACCCGTTGTCCGCCGAGAAAACACCTTTAAATTCAAAGAATATATATCCTGAAATGGTTTAG

>pR97

ATGGAGACAGATTACGAAGGTGGCATGCCAAATTGTTCGTATCGATATCCATTTCCACCGGAAGAAGACACAATATCCGC

CTACGCATTGCTAATTGATGGTCCTTGCACATGTGTAGCCGCGTTGCTCTCAATGATCGGAGCCCGATATGCCATCAAAT

TCCTCTGCCGCGCTGGCCTCAACAAAGAACTCACTGCCGCTCTATTCGCCCTTTGTGCCATCGACTCGTTGCTCCTGCTC

ACCGTCTTTCTCTTCTACGGAATCGAGGCGATGTCGTTGTTGTTCTTTCGCACAAATATTATGTACGATAAACAAGACTT

TACCTATAATTTGCACGGGATCGCGAGCTCACTGACCACCGCCTCGACAGCACTAGTGATCTACATCACTTTTCTAAGGT

TCATGGTTGTGGTGCAGCCTATGAGATTCGCTACAAGCATGAACTCCAACCATCGGAGGACTGGAAGCAAGGCTGTTCGG

AAGGGATCGGCTCAGTTGGAGGACTCGGTCACCAATAATACCATATTCTCGAGAGGAACCTTCAATAGTTCATCGATCAA

ACGACATTACAATATTCGAGAGCTAATTCGCCCTTTTTATATTCCATTTCTCCTAATTCTCATCAGCTTCTCCGTCAACA

TACCGATTTTCTTCGAGTTTACCACCACAAAATGTTTCAACGTTGAGCATAATGTCGAGGCGACCAATCCCGAGCCAACA

GTCTTTCGATCTTCGTTTGCCAAGTACAAGGCAGTGCTGATGACGTTGACGCAGACCATCGGGCCGGTATCGATTATTCT

TGTCCTATCGATTTTCACAGAGTACAAAGTGCATGTCAGTCTGAAGGCTAGAAGAAAGTTATTCGAGAGCCAGCAAAGAA

GTCGGTCGGTAGTTCTCAGCGAAGAGTTGAAGGAGACTGTGTCGAGAACTGTCGCAGTGTTTATTGCAGTCAAGTTTTTG

ATTTTTAGAACCATGCCGATCTTCTTCGACGTATACGAGAATCTGTATGGAATCGACGAGTGGGGTACTGTAATGAGCAT

AATGGTGCGAGTATCCGACTTTGGAATTGTGCTCAACACTGCCACAAACTCGTTGGCATATTTTGGAAAGAAAAGATGGA

TCGAAAAACGTCTTCGCCTCCGATTGATGAAAAAAGAAGAAAAACGAAACGCAAAAGCCACCCTAACAGTGCAGGGAAGC

TCGTTGAACTCCTCGATGAAGAAGTCACTGCCTCCGAAAATTGGGTTGCCACACCCGTTGTCCGCCGAGAAAACACCTTT

AAATTCAAAGAATATATATCCTGAAATGGTTTAG

>pR98

ATGGCGATCCATCATCCGATGTGCTACTACAACTTTACGTGCCGACCGGTGTTCATTGGCATCATGGTTCCGGTTGACTG

GGCCGTTCCAATGTATGGGTATATGATGCCTTTCATTGTTACACTCACGGTGGCAACAAATAGTTTTATTGTGGTTGTTC

TTTCGCACAAATACCTGCGGACGCCAACAAATTACGTGCTGTTGGCAATGGCGGTGACGGAGCTGCTAACAGGGCTGTCA

TGTTTGCCATGGTTCACGTACTATTATACTTTATCAGGATACAAAAACGATTTACAAACCGGGTTGCCCAGCTTCTGGTG

TGACATGATACCCTACATGGCGGCATTTTTACCGTCAATATTTCATACAATGGCAATCTGGCTGACAGTATATCTAGCCA

TTCAACGATACATTTATATATGCGTTCCGTCGCTAGTTCGCAAGTTTTGCACCATTCATAGAAGTAAACAAGTTATATTT

TTCATAATATCAGTTGCTACAATTATGTACACACCTGACCTCATGGCATTCCATAATAAAAGCCACGATGTGTTTGACAG

CAAGAGAAATCGAACTATGAAGCTTTGCTATCGACATCGTGCTCCATTCATGTTGAAACTCGGTGATGACATTTACTACA

AAGTGATGTTCACAACACAAACCGTCGCCGTTCATCTCATCCCGTCTGTGCTGCTTGTCATTTTCACATGGAAACTGGTG

GGCGCCATTAGGGTGGCAGACCGCCGTCACGCCAATCTTCTCTCCAAGTATTCCACGACGACTCGTTCGACGCGACGAAA

GTTCTCCGAGCTGACGAATTCGTCGGAGAACGAAAATAAAATAACTCGATTGTTCAAGCAGCGCGACAGCGTCAGTGTGG

GTAACGAACCTCGTCGAGCTCATGGATTGAAGCAGAATACGCGGATGCTGGTGGTGGTGATCCTGCTATTCCTGATCACT

GAAATACCAGCTGCATTAATTTTTACAATTCATGTGTTGTCAGTGTCTCTGAAGTTCAGCTTTGTAGACTATCAGTTCCT

GAATATTCTACTCATAGTCAGAAACGTCTTGATTGTGGTGTCCTACCCATTCCGCTTTGCGATCTACTGTGGAATGTCCC

AACAGTTCCGAGATGTCGTACGACAAATGTTCACTGGAAAAATGCTCACTCATGCGATCCGAGATAAGGATAACTCAACA

ACAATGCAGCTTGTCCAAGGTGTCACTGATCACAGTGATGATAAACGTCAATCGGTTGTTTTGTGCTCGGCAAACGGAAC

ATTGGTGTCTAGCGTTCCAGAGGAGCGGGCGAAGAAAGACAAGGCCGTTCAATGCAATCCTGACGTTTTCGATGAGGAGA

TTGAGGTTCCATTCGACCCATTCCTTGCAGATCTTGCCATTATTGTCGAAGGACACGATTCACCCAAGCGATTTAAAAAC

TCAGATTCCACATTTGATTGCTCAACACAATGTAGTACCGAAATGGACAATCACATGTTCATCAAGCGGAAAAGTTCACT

TTTGCGGGCAGCGCTGCGAAAAAATCAACAATTATCAGCGATGATGGCTCGTGTTTCTTCTTAG

pR99

ATGATGAACAACACCAACGAACAAAATTTCACGGAGGTCATCGATCCGATCAAAACTGTTGCTCATCTTACATGGTGTCT

GACCGCACTTATTGGCATTCCTGCCAATCTTTTTGTGCTGGCCGCTATTGTGTACTTCCGTGATATGAGAACAATATCTA

ACATCTACATATTCAACCTTGCCGTTGCGGATCTCCTCTTCCTCTGTGGCATTCCAGTGTCCCTATTTGCCCAGTCGTCT

CATGATGGTTGGATATGGGGACCAATAATGTGTAAACTGTACATATCTGGAAATGCAGTATCGCAGTTTGCCTCTGCCGT

CTTTATTGCAATACTCTCGTTCGACAGATATCTGGCGGTCTGTCGCCCGATTCAGTCGAAATCGTTCCGAACAACACAGG

CGGCATTTGCATTGTCTGTGGCTGCATGGATAATGGTCATTCTGGAAATGACACCGTTGTTCCTGTTTGTCAAGCTGATC

AAATCTTCTTCGGCTGGCGAGGCCAGGGGAGGAAGCTGCATGCTATTTGTTGGAAACGTCACTGTACTCGAGTCAGCAAC

CGAAATGAATGAAACATTGCTTAATGAGATCGAGCAGAACATGCTGGCTTCTAGAAGATTCTTCATCAGTTATACATTTG

CCCTTTCGTATCTGATACCATTAGTTGCCGTCTGGTATTTTTATTTCAAAATAATATTGAAAATGTGTCAAAGAAAACGA

CAAATGCAAACTAAAAGAACGGCTACCAAGAAACGAACTACCAAGGTAACCATAATGGGGCTAGCCATTGTCATCTCCTA

CACCCACTGCTGGCTTCCGTTTTGGATTGTTCAATGGTCGATTGAAGCAAATCTATTCGAAAAAAGCAAGTACCTGCTTT

TTTGCTGCACACACTTTGCATTTGCCCTTCAGTACATCAACTCTGCTGCGAACCCGTTCCTCTACGTTTTTCTCTCAGAT

TCTTTCCAGAAAAATATTCAAAAGCTATTACGAACTGCCAAACCGGATAAGCAGATCATGCTCAAAAACTCCACTGATCC

GCAGGACACCAGTAAAATGCTAAACGTCACAAATAATCCGTTGTACTCATCAGCAACCAATTCTCATTACAGTAGCTGCT

GA

>pR100

ATGAGCGAGGCGTGTATGGTGGAGAGCCTGGATCGACGATGGTACCTGGTAGCAGTTTGTGGAACATCATTATCGGTTTT

ATCAATTGTTTCAAATATGTTGATATCAAAAGTCCTACTATCCTCGAAGCACACGCACTTCTACTTCCTAGCCCTACTCG

CCTTATCCGACTGCTTCTTATCTCTAAACTACGGTCCCGTGATAGCAATGGACATAATCAAAGATCGTGTGAAACTGCGT

TGGCTCTCAAATCTTTATATTCAATACGTGGGACCCTTACTTGCTCTCTCACAAACCTCAATGACTTTCTCATGCTACCT

GATCATCCTGGCGACCATCGAGCGATATCTGATCACACTACGCGCAAAATGCCTCGGCACATTTCGACAATGCCGGGGAC

TCTCTGCACTTTGTATGTTCATTTTGGCTCTGTTGCTCAGGGCGCCCATCATTTTCGAAATGCAAATAGAACAGAATCCC

GATTGTATTGGCGAGATTAGCGAGTACTATTTGAACTTGTCGCCGATTGTGAACACGTTTTGGTATGGAACTGTGTACCG

GTTTTATGTCAGAAACATCATGACAGTCCTGGTACCCTTCGTTCTGCTGGCCATTCTTAATTACAAAGTCGTGAAAATTC

TTCGAATGCAGCAGAGATCGGCGCAAATGTTCCGTTTTCTATCAAGTGATCATAAGACGAAAATCCGATCGGCAACAATT

TTGATGGTATGTATCGTTTTCTCGTATCTTCTCGCGAATATTCTAAACGTCGGTGTCACACTGTGGGAGTACGTGGCATT

TCAATCAACACAAACCGGAGACGCTTATGCGTTCTACGAAACTTGCACCGATGTGACGTCAGTACTCTACATTCTGGTGT

GTGCCACGCGTCTTTTTGTATATTATTTGTTCAATCAGGAAATCCGCGAAGCCATCAAACCTCTCGTCTGCCACAGCACG

AGTGCTCCTCGCAAGCAGGACTATGTTACAGTAACCCGAAGCTTGTAA

>pR101

ATGAGCGAGGCGTGTATGGTGGAGAGCCTGGATCGACGATGGTACCTGGTAGCAGTTTGTGGAACATCATTATCGGTTTT

ATCAATTGTTTCAAATATGTTGATATCAAAAGTCCTACTATCCTCGAAGCACACGCACTTCTACTTCCTAGCCCTACTCG

CCTTATCCGACTGCTTCTTATCTCTAAACTACGGTCCCGTGATAGCAATGGACATAATCAAAGATCGTGTGAAACTGCGT

TGGCTCTCAAATCTTTATATTCAATACGTGGGACCCTTACTTGCTCTCTCACAAACCTCAATGACTTTCTCATGCTACCT

GATCATCCTGGCGACCATCGAGCGATATCTGATCACACTACGCGCAAAATGCCTCGGCACATTTCGACAATGCCGGGGAC

TCTCTGCACTTTGTATGTTCATTTTGGCTCTGTTGCTCAGGGCGCCCATCATTTTCGAAATGCAAATAGAACAGAATCCC

GATTGTATTGGCGAGATTAGCGAGTACTATTTGAACTTGTCGCCGATTGTGAACACGTTTTGGTATGGAACTGTGTACCG

GTTTTATGTCAGAAACATCATGACAGTCCTGGTACCCTTCGTTCTGCTGGCCATTCTTAATTACAAAGTCGTGAAAATTC

TTCGAATGCAGCAGAGATCGGCGCAAATGTTCCGTTTTCTATCAAGTGATCATAAGACGAAAATCCGATCGGCAACAATT

TTGATGGTATGTATCGTTTTCTCGTATCTTCTCGCGAATATTCTAAACGTCGGTGTCACACTGTGGGAGTACGTGGCATT

TCAATCAACACAAACCGGAGACGCTTATGCGTTCTACGAAACTTGCACCGATGTGACGTCAGTACTCTACATTCTGGTGT

GTGCCACGCGTCTTTTTGTATATTATTTGTTCAATCAGGAAATCCGCGAAGCCATCAAACCTCTCGTCTGCCACAGCACG

AGTGCTCCTCGCAAGCAGGACTATGTTACAGTAACCCGAAGCTTCGACTCCTCCCGCCAAATCGGAACCGAAATCGATGC

TGTAGCAATTGCCATTGCTCGAAGGCTGCTCTCGAGTGAGCTCGTCTTCGGAGGTGGAGGCACCAAAGAGGCCAACGACT

CATACGTCATCTCAAACCACCACGCGACCACATCTGGAGACGAGGAAACCGACGACGACGACGACGATGAAGAGATTCCA

TATATGGTTTGA

>pR102

ATGAATCGGTACGAGTTCTACGAGCTAAAATCATGGATGTATTTACCTGTAATTTTCATTGGACTTCTTGGAAACTTTGC

ATCATTTTTAGTTTATCGAACAGCGCCGATGAGAAAATCTACAGTTGGCACTCTTCTTCTTATTCTCTCACTCATTGACA

TTCTGCTGCTCATCCTGATAACTCCCATTTTTGTTCTTGTTTTTCTGCCACTCTGGCAGGACCAATGGCAAAAATATTCG

TTTCATATGAGCTTCTTTGCTTACAGTACACGATATGTTTATCCATTATGTATGATGACAAAATCATGTAGTTTATATCT

AATGGTACTTATAACCATTGAACGATGGATTGCTGTATGTAGACCACTTCAGGCAAAAGTATTATGCACAAACAGGAATA

CAATGAAAGCTGGAATTTTCATTATCATATTTTCCATAGTGTTTAATTTTCCAAGATTTTTCGATTATAAAATTGGAGAC

GGGTATCTATCTGAAATGTGGATGTTGGATACAGAGAAACACTGGTGGTACTTCATGTTCTACTTCATCATATTATCAGT

AATCTTCGATTACTTCCTTCCGTTCCTTATCATGACTATCGCCAATTATCATGTCATCAGAGCACTTCAAGAAAGCGACG

AGGTTATCACAGGCTTGGCCGTTCAGAAGCGAAAAGATCAGAAAACAACTGTAATGCTGCTAGTGGTAACCATTTTCTTC

GCGTTCTGCCATCTATTCTCAATGTTTCTCAAAGTTGCTGAAAGTATTTTTGGAGGATTCCTTACACAAACCAGTTTCTA

CTGGGAAGTGTTCGCCGAGTTCACAATCTTTTTGATAATCTTCCATACGTCAAGCACTTTCTTCATATACTACGGATTTT

CGGAGAAATTTCGAACAATATTCCATGACATCATCAGATGTCGCCAAAGGCTGGATGATGTAAAATGCTCGACCTACTTG

CCAGTTAACTCATCTGGAGCTGTTTCTGAATCAAAAACAAGAAAAGCTCCGGCAATTGTTTAA

>pR103

ATGTTGGAGCTTATCAGGATGCAAAGCGAGGAATATCTCAATGCCGATGTCTCAAAATGGGTCGAAAAATTCAGACGTGA

GCATGGTGGTGATGACTCAGCACTTCTCACATTGGTACCTCCAGGACTTTATGTGACTCTCAACAAATATGTTTTCCCAT

TTCAATTCTGCTTTGGAGTTATTGGAAATGTGCTTAATTTATGTGTGCTTCTGAGTAGGAATATGAGAAATGAAGCAAAC

ATTCTTCTCTCGGCAATGGCAATCTGTGATATCATTTTGCTTGTCACAATGCTTCCAAATTCACTCGGTGTCTGGTATCC

AATGTACATGTCCGATTGGTTCAGAAAATTCATATTCAATTCAAATACTTGGACAATTTTTCTGGCAAACTTTTGCTCTT

GCATCACCAGTTGGCTCATTTTGGCGGTATCCGTGGAAAGGTACATGGGAATTCGTTCACCAATTCATTTCCGATATCAT

TGGCGTACAGCTCGTGTGTTTTATCTGATCTTTTCAATAATTCTGGGTTCATTTCTCCTCACTTTCTTCCATACATTTGA

GTATAAATATGGTTATGCAATGATTCGTAATGGTACAAAGTTGTATGGATCACCGGTTAATGTTGACAAGCTCGTTGATG

TTCCAACATGGGTAAAGACAATGATTAAAACCTTCAAAATTCTGCAAGTGGTTGTTGGTGTTGTTCTTCCAACAATTGGA

ATCTTTATCTTCAATATGCTCATTGTTCTCATGCTCCGCAAATCTGAGTACTTCAATTTTCGATCAGCAGAAACAAAGGA

TGAAAACTCAAATCGCAAATACTCGGATTTGGAGATTCGCCAAAAGCGTGACATCAAAGTGACATTCACTGTCTTGGCTA

TCATCTGCTGCTACTTCGTGACTCATATTCCATCGGTCATTCCATTTGTGATAGAGCTTTTCAACTATCATCCAGACTGG

GTTAAAGTCTACGCAATTCCTATTTCTGGATCCTGGTTGATCACTGGAAAGGTAGCCAACTTCGTTCTCTTCTGTATGTC

ATCCGTCTACTTCCGTCGCCGTCTCCGTGATTTGATTCGTGGCCGATTCGACATGTGTGGTGCCAAAAAGAAGTTCAGTG

GTGTCTCTAGCGCACAATCAATTAAAAGTCAAGCTTACAGACTCACCAGATTCCGGGATCGGGACGACATCCAGAGACAA

AGCAGTTGTTTGACTCAAGTCGAGTAA

>pR104

ATGCAAAGCGAGGAATATCTCAATGCCGATGTCTCAAAATGGGTCGAAAAATTCAGACGTGAGCATGGTGGTGATGACTC

AGCACTTCTCACATTGGTACCTCCAGGACTTTATGTGACTCTCAACAAATATGTTTTCCCATTTCAATTCTGCTTTGGAG

TTATTGGAAATGTGCTTAATTTATGTGTGCTTCTGAGTAGGAATATGAGAAATGAAGCAAACATTCTTCTCTCGGCAATG

GCAATCTGTGATATCATTTTGCTTGTCACAATGCTTCCAAATTCACTCGGTGTCTGGTATCCAATGTACATGTCCGATTG

GTTCAGAAAATTCATATTCAATTCAAATACTTGGACAATTTTTCTGGCAAACTTTTGCTCTTGCATCACCAGTTGGCTCA

TTTTGGCGGTATCCGTGGAAAGGTACATGGGAATTCGTTCACCAATTCATTTCCGATATCATTGGCGTACAGCTCGTGTG

TTTTATCTGATCTTTTCAATAATTCTGGGTTCATTTCTCCTCACTTTCTTCCATACATTTGAGTATAAATATGGTTATGC

AATGATTCGTAATGGTACAAAGTTGTATGGATCACCGGTTAATGTTGACAAGCTCGTTGATGTTCCAACATGGGTAAAGA

CAATGATTAAAACCTTCAAAATTCTGCAAGTGGTTGTTGGTGTTGTTCTTCCAACAATTGGAATCTTTATCTTCAATATG

CTCATTGTTCTCATGCTCCGCAAATCTGAGTACTTCAATTTTCGATCAGCAGAAACAAAGGATGAAAACTCAAATCGCAA

ATACTCGGATTTGGAGATTCGCCAAAAGCGTGACATCAAAGTGACATTCACTGTCTTGGCTATCATCTGCTGCTACTTCG

TGACTCATATTCCATCGGTCATTCCATTTGTGATAGAGCTTTTCAACTATCATCCAGACTGGGTTAAAGTCTACGCAATT

CCTATTTCTGGATCCTGGTTGATCACTGGAAAGGTAGCCAACTTCGTTCTCTTCTGTATGTCATCCGTCTACTTCCGTCG

CCGTCTCCGTGATTTGATTCGTGGCCGATTCGACATGTGTGGTGCCAAAAAGAAGTTCAGTGGTGTCTCTAGCGCACAAT

CAATTAAAAGTCAAGCTTACAGACTCACCAGATTCCGGGATCGGGACGACATCCAGAGACAAAGCAGTTGTTTGACTCAA

GTCGAGTAA

>pR105

ATGAATGATTCTGAACACATGAAACTATGGATGTTTATATCTAAATGTCAATTCTGGGTTATTCTAATCACAGTTCTCAT

TTCCTTATTATTAGTGGCCAAAGCATTTTACAATTTAATGAAACGGAAAAGATCAAATTATTTTGTTTTTCTTATCAGTA

TAATAGCTGCAAATGTGTTAACTCTGCTGATAATTCTTTTTGATATTTTCAACTTTTCATTCAAAGGAACTCTCGTGTGC

AAACTTGAACTATTTATCTCAAATAGTGCAGCATGCTTCATAAACTGGATCTGGTTATGCTTATTCACTCAACGATTTTT

TATACTTTTCTATCCTATGAAGAGGTCTTCAAAAGGTTTTTTTGGATTTATGAGGTCTGGAAAGAAGTTGATTCTGGCAA

CTGCTTGCTTTGCAATTCTGACGCAGAGTTGGTCATTGATATTTATCGAAGAAGTTACAATGATGGCTGATGATTTCCAA

TTGATTGGAGTATGTGAGCGGGATGTGGATATTATGAGTGATTTTGGTTATCGAATTCTGGCAATCGGAGAAGCTTTTGT

CACCTATGCATTCCCGTTTCTTTTCACAATTGTCATGGATATTGCTGTACTTTATCAATCAGCTAATTCAAGCTTCGTCG

TAATGTCAGCTGAAAATATCCGATCAGAACATAACACCCTTCTTCATGTAAATGAAACAGTTAAAATACAATCATCTCAA

TCAATAAAACAATCAAATCGTCGACGGCATCAAGCTGTCAGAAGATGTCTAATGATGGCAACAATTCAAGTTTCTCTAAA

TGCTCCATACTATACACTTCAATTATGTGATGAAATATTCAGTTTGAGGAATTCAACATCTTTGTATTTATATTTGGATG

CAATTTTGTATTTTATATATCTCCTTCAATTTTCCATGATCTATTGCTATACGAATCTACTGGTTGCTCCAAGAGGAAAA

AGTTGTCGACAACCTAACAAGATGCCTTTGAGTTGCACAACAAGCCTTCGAAGTGAATACACAACGGTTTAA

>pR106

ATGGCATCCATCAATGCGACCATGTGTCAAGAGCAGCCCGATGGGATTTTGTATTGTCCAAATCACACTGGTGGTCCCGT

CTGGGTACGCAATGTCTATCCACCAATACAGGAACTCCAGCCGAAAGTTCTCATAGTAGTTGTGGTTTTTGTGCTTTGCT

TCATTGTCGGAGTATGCGGAAACTCATCTATCATTACCATTATTCGAGGAGTGGTTGAGGACAGGAGGAAAAGAATGAAA

CGTCATGGCGACAATGCAATTCTTTACATTGGAGCTCTATGCGTGTGCGACTTTGTCATGTCACTCTCCTTACCACCTGC

AATTTTGGATTCAGTAATCGGTTTTTGGATATTCGGAACAACAATGTGCAAAGTGCATCACGTATTCGGAAGTGTTGGGA

GAATAGCCTCAACGTTTTTGATAACTGCAATGTCATTCGACAGATACGTGGCTGTCTGCTATCCACATCATCATCGTCTA

CGTAGCCGTACTTTTGTCATCTCAACGATAGCCTGCCTCTCCTCAATCGCCTTTGTCCTTCTCCTTCCGATGCTGACTTA

CGCAAGTGCAAAGGAAATGGTTCTTCATGAATTAAAAGCTCATGAATCAGCTAACATTACAAGAGTAAGGGTTTTCAAGT

GCTCTGATATGATGCCTGGTCCTATTTTCTACTGGTTTACTTCAACAACCTTCATCCTTGGTTACGTAGTTCCACTAATT

CTTATTGTCTACTTCAATCTCAAACTAATCAACAAACTGTATGCCCATAAGCGAGTACTTCCTCGTTCTGCAATTCCAGT

TCGTCGTGTTGTAGTCTACACAGTTCTGATCGCAGCTGTATATTTTGTTTGTTGGACTCCGTACTGGTTCTCGGTTCTTT

ACGCAATCATCATGTCTCTGCTTGGAAAACCAACCACAAATTCCGAATGGGTGCTATTTGCCATTTATTGTGTTCATTTA

TTGCCATATTTTGGTTCATCAAGCAATTGGATTTTATATGGATTGTTGAACACTCAGCTACAGATGAAAAACGACATTGG

AGATGATGGTCAAAGCATTATGACAACCAACGGAGTTCAAGATCTTCCAAGACGTTCAACATCCCGCCTAATCTGTGATG

CTGGTGCTACGTCATCTCGATCAATACCAACCAACAATACTCCTCTATGGCAAAATGATTCAACTCTGACACGTGGAAAT

TCTGATACAAGTATGGTCTTCTATCAGGATACCGAACAAACACATCTCTAA

>pR107

ATGAATATCACAAATTCAAGCATCGGGTACCCACCTCCACCACTGGTAGGTGCATCATTTGCAAAAACAGCTATACCATA

TTCAATATGTTTTGTTTTTGGAACTCTCGGCAACACGGCCGTTCTCTCCTACGTTTTCTTCATTACACGATCCCTAAAAT

CATCTGTAACTGCACTTGGGAACACATTTATTTACATCGTTGCATTATGCGCAGTCGATCTTCTTGTCACCGTTTCGATT

CCATTCTCCCTATCGTACATGATTCTCAACAACTGGGTATTCGGAGAGCTGGTGTGCAAGATCCACTTTATGCTTGAGCT

TTCTAATAAGATGTGCTCCACATTCATTCTAACCGCACTGGCATTTGATCGTTACATGGCAATATGTCATCCGGAAATAA

AACGAATCCATGAGATGCGTCACACGATTTATATCACGACAATTCTTGCAACATTGTCACTTTTTCTCATATCTCCAGTC

GTTTTGTCTGCCAGAGTGACGAGTTTCAAAAGTGGACAATATTTTGTGAGCGCAAAGAATGAACGGCACGAAGTTATCCG

ACAAATGTGTATTGACGGAATGGCATTAGAGTGGAAGGTTTGGGTGTCTGCATTTCTGATCTTCTTTGCATTCTTACTTC

CATGCACTCTTCTAACCTACTTTTACGCGAAGATCGTCCTTCGTCTGAGGAGACAAAGAAGAACAATGCTCCAATCTCGA

ATTCCCCTCCGCCGCATTACAATATACACAATGGCAGCCACGTTCTTCTATCTTTCGTGTCATATTCCATTTTGGCTCCC

ACAGATCTACAACATTTTCTCGACAGTTCTTGGGCACAAAATGAATCCAAAAGTCATGACATTCACCTATTATTCACATC

TTCTTCCGTTCATATCGGCGGCATTTAACTGGATATTCTATGCTCGACTTAATAGTCAATTCAAAAAAGGATTGGTTCTA

GTTACGGAAAGAATGATCAGAAAGCGAACAAAATCGATGCATGAGAAAGGATACAGTGAGGCAGCTGTTGAGCTTACGAG

CAAGTTCGATGATGTCCCATTGATGTGTCCACACTGTGAAGCTCAACTTTCTATTCGTTCAAGTAGTAATGGAAAGAAGA

ATTCAAGATAA

>pR108

ATGTCGATGGTGAATACACGTGTTGAGGCTCCGGACCTTTTGGCGTTGAAAAACATGCTGCGACAATCAGATGGTACTCG

AATGCTACAAGAAATGAGAAGAAAGTTGATTCAACTGCACTCTTCGCAGATGATCAACGAAACAGAAGAGACATGTGATC

GATATATAGACAAGCATCCAGATATGACAAATGAACCAACAGTCCTTGTCACATTCTCCCTGCTTTATCTTCATATTTTT

CTTCTTGGAATCCTAGGAAATTCGGCTGTTCTATATCTTACAATGAAACATCGTCAATTACAAACTGTTCAAAATATATT

TATTTTGAACTTATGCGCATCGAATGTTTTAATGTGCTTGACGAGTCTTCCAATCACATTTATCACAAATGTCTACAAAC

AATGGTTCTTCTCATCGCCTGTTTGTAAACTTATTCCATTGGTTCAGGGAGCTTCAATCTTTGTCTCCACTTTCTCCCTT

TCCGCCATTGCACTTGACCGATACAACCTCGTGGTTCGTCCTCATAAACAAAAACTCAGTTCACGAAGTGCAATGATGAT

TGCTCTTCTCATTTGGGTTATTAGTGTAGTTGTTTGTATGCCATATGGATGGTACATGGATGTGGAGAAGCTCAATGGAT

TATGTGGAGAGTACTGTTCTGAGCACTGGCCATTAGCAGAAGTACGAAAGGGATATACTTTTTTGGTGTTAATCACTCAA

TTTTTGTTTCCTTTTGCTACGATGGCTTTTTGCTATTATAATATTTTTTCAAGACTCAGACAACGAGTGGAAACGAAACT

GAAGAAATTATCAGAAAGATCTCAACTTTTGGAGAATAGTACAACATGTGGAACCACTAACCACATTGTTAGCATTAACG

CGGAAGCAGTTCAAAATGGTCTGGAAAACAAACAAAGATTAGCTGTTCTTGCTCAACAAAGAAGGACAACTACTATTTTA

TCGTGCATGGTTCTTCTTTTTGCATTTACATGGCTTCCACATAATGTTGTCACTTTGATGATAGAATACGATGGATTCTT

TTTTCATTCTGATGAGACATCGGCAACCAGTACCGATCATACATATATTGTTTCAATGACTGCTCATTTAATATCCATGT

TAACAAATGTAACCAACCCATTTCTGTATGCTTGGCTTAACCCAATGTTCAAAGAAATGCTTATTAAAACTCTCAGAGGT

GGAAGTAAATCCCCGAAACCAGCTGACATCAAGCAAACTTCATTCATTCGAATGCCGAATAGTGGTGCGCCAAGTCAATC

TTCCTACCTGTGA

>pR109

ATGATCAACGAAACAGAAGAGACATGTGATCGATATATAGACAAGCATCCAGATATGACAAATGAACCAACAGTCCTTGT

CACATTCTCCCTGCTTTATCTTCATATTTTTCTTCTTGGAATCCTAGGAAATTCGGCTGTTCTATATCTTACAATGAAAC

ATCGTCAATTACAAACTGTTCAAAATATATTTATTTTGAACTTATGCGCATCGAATGTTTTAATGTGCTTGACGAGTCTT

CCAATCACATTTATCACAAATGTCTACAAACAATGGTTCTTCTCATCGCCTGTTTGTAAACTTATTCCATTGGTTCAGGG

AGCTTCAATCTTTGTCTCCACTTTCTCCCTTTCCGCCATTGCACTTGACCGATACAACCTCGTGGTTCGTCCTCATAAAC

AAAAACTCAGTTCACGAAGTGCAATGATGATTGCTCTTCTCATTTGGGTTATTAGTGTAGTTGTTTGTATGCCATATGGA

TGGTACATGGATGTGGAGAAGCTCAATGGATTATGTGGAGAGTACTGTTCTGAGCACTGGCCATTAGCAGAAGTACGAAA

GGGATATACTTTTTTGGTGTTAATCACTCAATTTTTGTTTCCTTTTGCTACGATGGCTTTTTGCTATTATAATATTTTTT

CAAGACTCAGACAACGAGTGGAAACGAAACTGAAGAAATTATCAGAAAGATCTCAACTTTTGGAGAATAGTACAACATGT

GGAACCACTAACCACATTGTTAGCATTAACGCGGAAGCAGTTCAAAATGGTCTGGAAAACAAACAAAGATTAGCTGTTCT

TGCTCAACAAAGAAGGACAACTACTATTTTATCGTGCATGGTTCTTCTTTTTGCATTTACATGGCTTCCACATAATGTTG

TCACTTTGATGATAGAATACGATGGATTCTTTTTTCATTCTGATGAGACATCGGCAACCAGTACCGATCATACATATATT

GTTTCAATGACTGCTCATTTAATATCCATGTTAACAAATGTAACCAACCCATTTCTGTATGCTTGGCTTAACCCAATGTT

CAAAGAAATGCTTATTAAAACTCTCAGAGGTGGAAGTAAATCCCCGAAACCAGCTGACATCAAGCAAACTTCATTCATTC

GAATGCCGAATAGTGGTGCGCCAAGTCAATCTTCCTACCTGTGA

>pR110

ATGGGAGCCTTCTTTCCTGTCATTCTGCTCACCCCAACACCAATTTCCTCAGCACATAACCTGTATCTTTTTCAGATGCT

CGAGCTACAGGAAAACATCACCGATTCTCAGCCGATGGATCCGCCATCGCTAGAGATCATGATGCTCCATCACCTCATGA

TCATTCTGGTCACCCTTTTCGGTAACACACTTCTGATCTATGTGATCTACAAGAACAATGCGGTGCTGAGGCGGAAACGG

GTCACTCCCGTGCAGATGCTAATGCTTCACATGTGCGCTGCTGACATTCTATTCGCCCTGATCAGCGTGGGCCCGACTAT

GGCCATCACTGCCACCGTCCCCTTCTTCTACGGCCCAAACCTGCTGTGCAAGCTCACCAAATTCCTCCAAGTGATCCCAA

TGTACGCCTCATCCTTCCTGCTCGTGGCGATCAGCGCGGATCGATATCAAGCCATCTGCCGTCCGCTTGCCTCTATGAAA

TCATCCATCTACAACCGCCCGGCCCTCTACTCCGGAATTGCCTGGACCGCGGCGATCCTCTTCAGCACACCTCAACTCTA

CCTCTTCGAGAAACGAAACGGAGATTGCTCAGAAAACTACACGACCGCCTTGCAATACCAGCTCTACGTTTGCTTGTTCA

ACTCAGTGGTATGGCTTCTGCCGAGTGCCATCGCTGGATGGCTCTACTTATGCGTGTGTAAAGCGGTGTGGAAGAGCACC

TCGTTCAGCAGCTCACTGAGGAATAATATGAAGAAGATGGAACATATGAAGCTGACCGAAAAGAACGGCGGCATGCAGGC

TCACCACAAAGGAGCCACGATGCAGTGCGTCGAGTTGGATCGAAGGAGAGTTCAGACCGTGAAGCTCACGTTGACAATTG

TCGCCGCGAACTTTGTGCTCTGGGCTCCGTTTTGCATCACTTCGGTGATTGATGCCGTGTGGCCAACGGCGATAAACTCC

ACATTCGCCACATACATCATGTTCTTCGGCAACCTGAACAGCTGCATGAACCCCTGGCTATGGTTCCATTTCAACCGGAA

ACAATTGAAACGCGCATGCCCGTGCCGAAAATCCTCGGAACCCCTTATCCAATCGTTAGTTTACGTACATGTGATGACCA

GTGAGCAAAGCGATTTTTGA

>pR111

ATGATGTCTCCATATTTTGCCGGTTTTCTTTACATAGCCACAGTTCTTACCGGACTATTTGGAAATATATGGGTCGTTTG

CTCAGTAGCCAGAAGTCGAAAACCAAAATCCCTTATGTCCAGAAGCTCCCCTTCAGATCGATTACGTGCTTATATTTCAG

TGTTAGCAGTGATTGATTTAACAGTTTTAATGGCTCTTTTAGTTAGAGCATTATATCATTTTCTACCACATTTTATGCTC

GATTCAAATAGTTGTCGAGCAATGTTTGTACTTGAGAATAGTGTAAAGATTACATCATTAACAGTTCTTTCGTGTATTTC

AATTGAACGATACATCACAATACGAAAACCATTCTGCAGTGAAGTTCGAAGGCAATTCGTGAATGCAACTCCAATTGGAG

CAAGTATATTCGTTGGGCTTGTCGTTGGTGCAATAATCGTTCAAATCAATTCCGTGACTGTATCATCTGATGGATTGAAT

TGTGTTCGTTCATATCGAGGAAAAGCAATTCCAAGAGTGGCTTCTTATCTAACAGCAGTCGCATTTCTAGTCGATTTAAC

AATTATTTCGTTGAATTATTCACAAATTGTCCGACATGTTAGAAGAAAATTTACAAAGCGAAGAGCTAGAGTCCAAGCCA

ACTCTCGAGTACGTGAATCACTTGTGAATGAGCCCCGTTATATGAGAGAAATGACAGCTGCAATTGTTCGTGTTGGAGTA

TTCCACGTCGTTTGTTGGCTTCCAATGTCTCTGATGCAGTTTATCCCCGATAATACAATTCAATCGGAACTCACAGCTGG

AATCAGGTTATTCAGTAATTTCCGTGACTATAGTATAACTCGTTGGCTAATCTTTATTGCCACATGGCTTACATCCATGA

ATGCAGCTGGAGATTGGATTTTCTACGCGGTCATGAACAGAGACCTTCGAAATCTTATTAGATTTGCCACTGAACGGCGA

AAACGATCAACAATGTCTCATGCAGCATCTCCATCAACAAATCGAAGTCTTCGTCAACAAGTTCCACCATCCATGAAAGT

TCTTCATTCAATATCATATAGAAGTTCAATTGGAGGCTCTATGGATGATGCAGCTGGGGCATCTTTTCTTCAAAGTTCAC

CTAAAGAATCAACTGTATCTCAAGAGCCGAATAGTCCGTATGGAGTTCGAGCGAAAATTAGTATTTTGACAAGAAAAGAA

TCTGATGGAGATGGCGATATGGTTTAG

>pR112

ATGGATTATGAAAGCACGTCGTTCGATCAGTCAATCGCGACTAATGACACCGACCAAGAATGTGCATATTCCGAACCCGT

ATACGTCAAAGAACGTTTTTGGATGGTCGCGATTTTTGGCACCGCTGTGTGCATCATAAATATCCTCGAAAACACATTTC

TCTCCTTTATGCTATTCAAAAAAAAATCATATCGTTCGAGTCATATGCTCTACCTGGCACTTTTAGCCTTTTTTGATGTT

TGGATGGCACTTGCATATATTCCATTGATGAGTTTGAACCTATTTGTCGATTACTACAAGTCTGTAGTCTTACTTCGTGC

TTGGTTTGCCTATATGTTACCCATGATCACAGTATCCCATATCGCTATGACAGCTTCATCGTTTTTAATGGTTGCTGCTT

CTTTAGAAAGATACGTGATAACTTGTCATCCTACGAAAAATCGGTGGCTTTCAAGAAATCGAATGTGGATTGCAGCTTTT

GCTATTTTCCTCGGTACAACTTGTAAATTCTCACAACTGTATGAGATGGAAATTGAATATCTCCCCCAATGCATGGGAAC

AATGAGAGAATATCAACTAAATCTTTCTGCATTGGCTCGCTATGAAATTTATGGATACTGGCGTGTTCATTTTCGAACAA

TAGTAACAATATTGATACCATTCTTCTCATTGGCATTTATCAATATTCGAATAGTCTTAGTACTTTCTAAAAACGAATTC

AAATTTCTTCACTCAACAAAACTAAGCGATGCAAAAAGGAAGTCGGCAGTTCGAAATGCAACAAGAACAATGGTATTCAT

TGTATTCACTTATCTACTATCAAATGTTCTAAATGTTATCATTATCATCTGGGAATATATTGATATGGATATGTTGACTC

AACAATTTGAAACTTTCTACATGTTCGCTGTTGATGTCGTTTCACTTTCTACAATTATTTTTGGAGCACTTCGACTTCCA

ATTTATGTAATTTGTCAAGCCAATTTGAGAAAGGAATTTTTTGCTCAATTAAAAACTATGCTTTCACTCAACAGTTACAC

GGGTTTTCTTTCTTCATCTACCAAGAATATGACTATCGAAGATATGGATATTGATCGAAGGACTCCAGATATAAGTGGAG

ATGGTACCGTCCTGATCGTTGAATCCGATCAGAAAAAGGATTCTTCGGCGACTTGTACCGTGATTCGAAAAAATAGTCTT

CATCGAATGAATGGAGGATTCAGTGATATGAGCGAGAAAGTTCAAATTGTTTAA

>pR113

ATGGATTATGAAAGCACGTCGTTCGATCAGTCAATCGCGACTAATGACACCGACCAAGAATGTGCATATTCCGAACCCGT

ATACGTCAAAGAACGTTTTTGGATGGTCGCGATTTTTGGCACCGCTGTGTGCATCATAAATATCCTCGAAAACACATTTC

TCTCCTTTATGCTATTCAAAAAAAAATCATATCGTTCGAGTCATATGCTCTACCTGGCACTTTTAGCCTTTTTTGATGTT

TGGATGGCACTTGCATATATTCCATTGATGAGTTTGAACCTATTTGTCGATTACTACAAGTCTGTAGTCTTACTTCGTGC

TTGGTTTGCCTATATGTTACCCATGATCACAGTATCCCATATCGCTATGACAGCTTCATCGTTTTTAATGGTTGCTGCTT

CTTTAGAAAGATACGTGATAACTTGTCATCCTACGAAAAATCGGTGGCTTTCAAGAAATCGAATGTGGATTGCAGCTTTT

GCTATTTTCCTCGGTACAACTTGTAAATTCTCACAACTGTATGAGATGGAAATTGAATATCTCCCCCAATGCATGGGAAC

AATGAGAGAATATCAACTAAATCTTTCTGCATTGGCTCGCTATGAAATTTATGGATACTGGCGTGTTCATTTTCGAACAA

TAGTAACAATATTGATACCATTCTTCTCATTGGCATTTATCAATATTCGAATAGTCTTAGTACTTTCTAAAAACGAATTC

AAATTTCTTCACTCAACAAAACTAAGCGATGCAAAAAGGAAGTCGGCAGTTCGAAATGCAACAAGAACAATGGTATTCAT

TGTATTCACTTATCTACTATCAAATGTTCTAAATGTTATCATTATCATCTGGGAATATATTGATATGGATATGTTGACTC

AACAATTTGAAACTTTCTACATGTTCGCTGTTGATGTCGTTTCACTTTCTACAATTATTTTTGGAGCACTTCGACTTCCA

ATTTATGTAATTTGTCAAGCCAATTTGAGAAAGGAATTTTTTGCTCAATTAAAAACTATGCTTTCACTCAACAGTTACAC

GGGTTTTCTTTCTTCATCTACCAAGAATATGACTATCGAAGATATGGATATTGATCGAAGGACTCCAGATATAAGTGGAG

ATGGTGAGAGACAGTATCCGTTATTACTTTCGTTAGGTACCGTCCTGATCGTTGAATCCGATCAGAAAAAGGATTCTTCG

GCGACTTGTACCGTGATTCGAAAAAATAGTCTTCATCGAATGAATGGAGGATTCAGTGATATGAGCGAGAAAGTTCAAAT

TGTTTAA

>pR114

ATGAACTCAACGGACATTATTGCAAATGTCACAAAACCATTTGTAGAAAACTTAACATTAGGAGAAACGGCGTTTTATAT

CAGTTGTGGCATTGTTGGTACAGTTTTCAACGCATTAGTACTATGGATTGCACTCACATATATCAATACGGAAGATAAGC

CTAGACAGATAATTGTTATCAACATGACAGTGGCGGATCTTTTGATGTGTATTGTGTACATGAAAACTCGCCCATGGCTC

TCACATTTTAATCTTTGGCTTTGCCATCCCTACTATGTTATCATTTGGACGTGTCAAATGTGCAGTTGCTTGAATCTCGT

TTGGTTGAATGTTGACAAGCTGATCTACATTCAATTTCCGTTGCATTACTATCAAATTGTGAATCGGAAACGGCTTTTAT

GGATTACTGCAGCGACATGGGGAGGACTTTATGCAATGAACATCGCACTTGTCACCTTTTTGAAAATAACTCGAGGAAGC

TGTCTTGGAGTATCACTTAATCCTTATGTATATCTTTTGAGCCCCATATTCTATGTCGTAATGATTCTTACGTCATTTTC

CTTATCAGCTTTGATCTACTGCATTGCTCACAATTTGACCCATATGGAGGAGCGGCAACGATCGAAACTGTTTCGTCGGC

TGTTCTTCTTGTTTTCCAGCACATTATGGACTTTCTTCACGTGTCTTCCGTATCGGCTCTTGTACCTATTCAGTATTTTT

TGTGGAGAAACATGTCAAATTAACAATTATTACAAAACTGCAACCAACCTTTTTTTCCGTCTTTTAATCGTTGGTATCAT

GATTAATCCAGTGATCACTATTTGGACTCAGAGAATTTATCGCCTACGCCTTATGCGTATGTTTGGGCGACTGCGTGAGA

ATTCATCAACTGAAGTATTGATGGTTAGCAATCGACGAGCATCTGAAAGGCCGCCGGAGCACACCCCCTTACGGTGTGAT

ATGTAG

>pR115

ATGGATGATTCCGTGGATATTTATGAGCAGGCTGCACTGAACATGTCAATACCTGACGGATGCGATGTCAATCGAGTTGT

CTACGCATCAATCAAGCAGTTAATCAGTCATGATATTATCGATCCAAGCTCAGTGGGATTTCAAAACTGTGAACCTTTGT

GTGGAATATGTTACCATGGATCCAAAGAGTGGGATTACATTCGCTTCAATATTATTGTGATCGGAATAATTCTTCCTATT

GTTGGAGTTTTTGGCATTGTGGGAAATGCTATTTCAGCTTTTGTTTATAGTCGTCCAGAAATGCGATGTTCCACTAACTT

GTATCTTTTCTCACTTGCGTGCTCCGATACGGGTGTTGTTCTAACCGGAATTTTTTTATTCTCCCTCGAAACATTCCGTC

CATTTTCGCTGACTGTTGCGAGAATTTCTGGACAGTTGTCTGCAATCGTGTATCCAATGGGCATGATTGCTCAAACGTGC

TCCGTGTATTTCACCGTTTGCGCTGGTGTTGATTGTTTTGTGCAGGTTTGCCTTCCCGAAAAAGTGCGACGAGCATTTTC

CCGCAAAGAAACTGTTCACTTTCTTGCTACATGCGTTGTGATATTCTCAGTTCTCTACAATGTTCCACACTTTTTTGAAG

GATTCGTCATTGATTGTTATCATCAAGAATTGGGTGGTATGAGTAAAGAAGTGTGTCCAGCTACTTTGAGATATAATGAA

ATGTATCAGTCAATATATTATAAATATATGTATGCAATTTTTCTTGCCGTCGGTCCTTTGTTCACATTGATTATTCTCAA

CACATTGATTATTGGTTTCTCAGTGTTCGGGTCAAGTGCCTCTAATATGGATGATACTATGTCATTAATTCTTGTTGTTC

TGCTATTCATTTGCTGCAACACAATTGCCTTGGTCATAAATATATTTGAAAGTTATCTATCGGAGACGCTGGGCAGCAAA

ATTAACTACATAGTGGATTTGTCAAACTTTTTAGTGGTTTTTAATTCGAGTTTTAACATAATCATCTACATCAAATACTC

CCGGCCATTTGCTGACACTCTTTTCTCCTACTTTTGTAACCGAAAACCAATAACCGACAATGATGGACCCGGACCACCGA

TGTTGATCACAGAAAAACCATCTCGGTCAAAAAAGTGTAAAGAGACATTGTCTCGGTTACTAGTCGCTTCTCAACCAGAG

GTACTTATTTGA

>pR116

ATGGCGTCCGCATTGAACTGCACGCAATTCACTGTGCCATATCCGGAGCCGGGCACCGATGAAGTTGGGAATAGCACGAT

TCCATGGTTGATTACTACAGTTGTAAAAGCATACCGGCCGTTTCACTACTATATACTCACTTTTTTAGTTGTTTTTGCAT

TTTTTGCAAATATTATGATTGTGAAAGTTCTTTCCCACAAAGAAATGATTCGTTCTGGTGTGAATGTTACTATGTTGCTT

ATTGCGGTTTGTGATTTTGGGTGTTCAATTGCAGGACTTTTACAATTATTTTTGAGAAGCTACTCAGACAACTATACATC

ATACCCAACTGCTTATGGACAGATAGCGGTTGACTATTTAGCTGTAGCATTCCATGCATCCTCCTTATATCTCGCTGTGG

GAATGGCATTTTGTAGAGTCAAATCGTTAAACATTGCAAATAGAAACAAAGATCTTTGGCAATCACCTAATTATGCAGTA

CGAGTTGCTTGTGCTTTGTGCGCACCTGTTTTCCTAATTGCCACCTTTGTGCTTTTCATAAACGCCGTAAAAAACACCGA

GGAAGAAGGCATTTATTTGGATATTTCTGATTTATCGCTGCTCAGTGAGTGCTTGTTCATGAAAGCTTCTCTCATGGCTA

GTGGGCTTTGTTTTAAGATTGCTCCGTGTCTTCTAATGCTGGTATTGTCGCTGTTCCTTCTGAGACGAATTGACGAGGGA

AAACAAAGTGCGCAGAATGCAGCGGCAAATCAGAAAGGAAAAAAAGATAAAATCGACCGCTCATCTAGATTCATACAATT

TGTTCTAGTCGTATTTCTAATCACTGAGTCACCACAAGGAGTTTTCAGCATTCTTGGTGGCTTTATGATATTAGATTACA

TAAACTATTTTCAAAATCTATCAATTTTTATGAATATTTTGGCATTTTTCAACACTACTACAAGTTTTATCATTTATAGC

TCGTTATCGGCAAAATTTAGAAGAATTTTCGCTCAACTTTTTCTACCAAGCCAGGTTTTGGAGAAAATTTATTTGAAGCG

AAGTACGGTTGTGGTTCATTCAATTGCACATTCTTCACGGACAGTTTGA

>pR117

ATGGAAAGCCAGCAGTTGATGGCATGTGCCATCCTGGTAATTGTGCTGGTAGGGATTTTTGGAAATTCACTTTCGTTTAT

TTTGTTCAGCAGACCGCATATGCGATCTTCATCAGTAAATGTACTTCTATGTGCTTTATCATTCTTCGACTTTTCACTAT

TAACCCTTTCAATTCCAATATTCGTCATCCCAAATCTCGATTTATGGGCTAACGATTTATCACTTTCAACATATATGGCA

TATATTCTCAAGCTCATCTATCCAATCAACTTGATGATGCAGACGTGTTCCGTTTATATAATGGTGATGATAACTCTCGA

ACGTTGGGTTGCCGTGTGTCGACCTTTACAAGTTAGAGTCTGGTGCACACCTCGAAAATCTCGAAACGCGATTCTCGTCA

TCATCGTGTCTGCTTTCTTGTACAATTTCGTTCGGTTTTTCGAATATCGATTTGTAGTGACGGAAAGTGGAGCGCTATAC

GAAAAATGGCTGAGAGATCCAGGAAAGCATAGGTGGTACTACGTTGGCTACTACACAATTCTTTACATTGTCACACATTT

TTTGGTGCCATTTTCTGTTATGGCATTTGCAAATGGGCATGTGATAGTTGCAATGTGCAAACTTAGCAAAACACGGCAGA

TGCTCACAAGACAGCAACAACGGGAACAGTCGACGACAGTAATGCTACTTATCGTCACATTTGTTTTCGCAATTTGCAAC

ACTCTTCCTTTCCTTCTCAACGTCTCAGAGAGCATTTTCCCGACATTATTCCAAGACGAGAGCACGCGGGGACTGGCATA

TTGGCTCAATGATCTATCTAATCTTCTAGTGGTTCTTAACTCGGGTACAACTTTCATTATCTATTTCACTTTCTCAGAGA

AGTACAGGCAAACTTTAGTTTTTATTCTCAAGAATGGATGTTGTGCAACCGTCAGCGACTACAACAACTATACAGCCATG

TCTAGAACGGCTTCGATGAGAATTTCAAGCGAGACGGGCGGTCAAATTCAACGACAGGGATCTAAAATGAGCAATAGCAG

TTATAGAAAGCCAATTCTAAACGCGCATCTGTCGGAGCCGCTCATCGGACGTTCTTCTCAAACCAATTTACATGCAGAAA

AGAAGCGAACGCTTCTCGTCGGAGTACAACGAACGAACTTGCAAACACCTTGCTCCTTTCGAAGAGCACAAATTACCAAA

ATTGCCGAGCGAGAAGAGGAAGAAGAAACTACATAA

>pR118

ATGGAAAGCCAGCAGTTGATGGCATGTGCCATCCTGGTAATTGTGCTGGTAGGGATTTTTGGAAATTCACTTTCGTTTAT

TTTGTTCAGCAGACCGCATATGCGATCTTCATCAGTAAATGTACTTCTATGTGCTTTATCATTCTTCGACTTTTCACTAT

TAACCCTTTCAATTCCAATATTCGTCATCCCAAATCTCGATTTATGGGCTAACGATTTATCACTTTCAACATATATGGCA

TATATTCTCAAGCTCATCTATCCAATCAACTTGATGATGCAGACGTGTTCCGTTTATATAATGGTGATGATAACTCTCGA

ACGTTGGGTTGCCGTGTGTCGACCTTTACAAGTTAGAGTCTGGTGCACACCTCGAAAATCTCGAAACGCGATTCTCGTCA

TCATCGTGTCTGCTTTCTTGTACAATTTCGTTCGGTTTTTCGAATATCGATTTGTAGTGACGGAAAGTGGAGCGCTATAC

GAAAAATGGCTGAGAGATCCAGGAAAGCATAGGTGGTACTACGTTGGCTACTACACAATTCTTTACATTGTCACACATTT

TTTGGTGCCATTTTCTGTTATGGCATTTGCAAATGGGCATGTGATAGTTGCAATGTGCAAACTTAGCAAAACACGGCAGA

TGCTCACAAGACAGCAACAACGGGAACAGTCGACGACAGTAATGCTACTTATCGTCACATTTGTTTTCGCAATTTGCAAC

ACTCTTCCTTTCCTTCTCAACGTCTCAGAGAGCATTTTCCCGACATTATTCCAAGACGAGAGCACGCGGGGACTGGCATA

TTGGCTCAATGATCTATCTAATCTTCTAGTGGTTCTTAACTCGGGTACAACTTTCATTATCTATTTCACTTTCTCAGAGA

AGTACAGGCAAACTTTAGTTTTTATTCTCAAGAATGGATGTTGTGCAACCGTCAGCGACTACAACAACTATACAGCCATG

TCTAGAACGGCTTCGATGAGAATTTCAAGCGAGACGGGCGGTCAAATTCAACGACAGGGATCTAAAATGAGCAATAGCAG

CCGCTCATCGGACGTTCTTCTCAAACCAATTTACATGCAGAAAAGAAGCGAACGCTTCTCGTCGGAGTACAACGAACGAA

CTTGCAAACACCTTGCTCCTTTCGAAGAGCACAAATTACCAAAATTGCCGAGCGAGAAGAGGAAGAAGAAACTACATAAA

ATGTCGGCTGTAGAACATCGTGGGATGCCTGAAATCACTATCACTTTTAGTGAGGACCTTCCAGATGGAGAACCCGATTC

TCCATGTCAACCATGCTAG

>pR119

ATGTACCAAACACCTCCAACATTAACCGCAGCTACAGTAATCCAAATACTCACCTGGGGCTGTGCAATCCCCTCCCTAAT

CGTAAACGTTTTGCTCATAGTCAGCATTAATCGCGAACGTCGAACCGACAATCGATTCCGCTGGTTTAATCCGATTTTTC

TGTACATGTTGTACTGTAACATGGCGCTTGCCGTCCTCTTTCCACTGGTTCACCCAACATACTTTCCCGCTCGTGGCTTG

ATTTATATGGTTTTTGTTGGTCCATTGGCAAGGATTTTAAACCAAATTGTGCAAAAAATTGTCTATGGATTGATCGTCGG

AATCATCACTTTTCAAATGTGCATTGCTCCGATTACGTTGGTTTTGCGATGGATGATAGTGAAGAAACGCATCTCGTGTA

TGACTCTTCACTATAAAACAATACGAAAATTATTCTTTGCAATAAGTTTATTTCTACCTATAACCTTAGAGATATACCTG

ATCTTTTTCACGCCAAACCAGAAAATCGAGTCTTATTCCGAAGTGTCACATCGAGATGTTGTCATGAAACTGAGCGGTCA

CGACTTTTCATATTTTGTGGTTGATTGGAGTTTCTCGTTGCTAGTTTTCACATTTACCATCCCCTACATTACTTGCTTTA

TCGTTAGTCTGGTTTTTCATAAATTTTATATGAAATCGATTCGGAACGTTTCCAACGAGTTCATCGATATGGGATATCAC

GACTCGGCGAACATCAATAGAACCCTCCTATCAATTGCTCTTCAGAATCTCATATCAATTTCCATTCCACACTTTGTGCT

CACAATTACGTCTCTTCTGAAATGGGATCTTGGTGTCCGTGCGTTACCGGCTTCGTTTGGATGTATTACGATTACATTGA

CAACGTCTCTCACAATTTTTGTCAATTTTAGAGCTTACAGACGACAACTCGGCTTCATTATAATCGATTTTTTCCGGCAA

AAAAATGTGAACCTTTCCGTCTTTGTCGTCGATAATCAGCAGCAAGCCGATCAACTTCAAATGTCCACCGTCGGAGCACC

TATTTCGATGATGTGA

>pR120

ATGCAACCGCCAAACTCTACGGGACAATGGACAGAAATTCACTGCGATTGGTTTGCAGAAATGCAAAATTATCAGAACAG

TCTCTACAAAATAAATTTTGATGGATCCCCAATAGTGCCAATGTATGGGTTGGTGTGCTCATTCGGAGCTCTTGCCAACT

TTATCGTTCTCCTCGCATTTGTCAGAACTGCAAATCTTCGAAACTTGAGAAACAGTTTCATTGTGAATTTGGCATTTTCC

GATTTGATTCTGTGTGTGGTCACAGCGCCAGTCACTCTCTATACGAGTTTAAATTTATTTTGGCCATTCGGAGATTGGTC

GTGCAAGTTTCTCGCTGGTGTTCAAGCAGTGAACACATTTGTTTCATCACTTACTCTTGCATTCATTGCAATGGATCGTG

TTCTTCTAACACTTTGTCCAGTTCGATGGAGATTAGCAGCAACTGCACCACTTTTATGCTATGGTGTCGTATGGATCATT

TCGATCATAGTTGCTCTTCCATATGCATTGGCAGTAAGCTCAAAGTTGGCTCCATTTGATCCATGGAGTGATAGAGCAAC

TCCCAAAATGCTAACCTATTGCAACCGTCAAGTGCCAGAAATATGTGCAGAGATGCAAGAAGCCTGGGATAATGCAATTG

TCTCAAAAACCACGTACACATTTGTAGTACTTGGTATTCAATATATTCTACCACTTGCTGCCTTGGCATATGCATATTTC

CAAATCGGGTCAACAATTCAAAAACGATCAAAAGTATCACGTACAGTGGATACAACAAGAAGAATGCAAATGCAAAATAG

AAATCGAAGAGCTCTTTTGTTGCTCTTTTTGTTAGTACTCACATATGCCGTGTGTTGGGCTCCAATGAATATTTATCATG

TTTTGAATGGTCTCGAGATTATTAACTATTCACAAAATATGTACATCTTCTGTCATTTGGTCGGAATCTCTTCAACGTGT

GTTAATCCAATAGTATATGCACTGGTCAATGAATCATTTAGAAATGCTTTACAATCAATGATTCTTCAATTCCGCCCGTG

TTACGTCACAACTACCGGTACAGCAGCCACCAACGTGTATGCATATTCTGCGACTTCGAAGGCGGAAAATGTCACTTTGA

TGCGTGATCCTTTCTCAACAACTCCTCGTCCAAATGAACGCCCAGACAGTGTTTGA

>pR121

ATGCCAAAAGAATGTTCAGTGATAATCAGTCCTTATACAATTATGATATCGTCAATATTTCTACTATGTCTTATACTTTC

ACTACTTGGAAATGCAATTGTGATCCTCACAATCCTCGGAAAATCACATCGATCCAGAAGCATCACAAATTTCTATCTTC

TCAACTTGGCATTTGCAGATCTTCTTCGTAGTATTATATGTATTCCAAGTACACTTCTTGGTGAATTGACACAATGTTGG

CTCCTTGGTGCCGCCATGTGTAAAATTGTCGCATTTCTTCAACCTGTCGGAGTTTGTGCATCCGCGTACACTCTTGCTGT

CATTGCCATTGAACGATACTATGCAATTTGTCGACCATTGGAATCAAGAAAATGGCAGACAAAGAAACGAGCACTGATAA

CAATTTCATTGGTATGGTGCTTCAGCTTCTCAGCCAATTTAACAAGTCTTTTCCTCTACGACGCTAACCCTGGAAAATTC

ACGTGTGACTCCACGAAAGGACCACTCGTTGATTTTATCTACCAACTTTATTTAACTTTTACACTATTATTCGTACCTCT

TGCCCTGATGGTCGGTCTCTATGGAAATGTGATAATCACTCTGAATACTGCCATTAACAGTGATCATCCCACAGTTGAAC

AGCAGATGATTGAAAAAACTCTCCCGTCTCGTGCTTCATTTTCGGATTGGTTTGTTAGTGCAGTGCAACGAGTTCCAAGT

ATGAAAGTTGTTTCAAAAACTTTTCAATTCAAAGAGAAAAACTCTCTGTCAATTCCTCAAACAAGCGGATTGTCTGTTCG

TCCCAGTAGATCATCGTTTTCATCCTTTTTCTCAACACCACGTGGATCGTTTGATGTTACGATGCTTTTAAGGAGCACAA

ATCAAGAAAAGATCCTGATCGCCAAGAAGAAAGTGACCCGAATGTTGATAACACTTGTGATTGTTTTTGCATTTTGTTGG

GTGCCATCTTACCTGTATTGGCTACTACTCAGAATGGCTGAACTAGCGGCGACTGATTTGTGGAATCCTGGACTCAACTC

ATCATTGACAATCATGACATACATCAGCTCTTTGGCAAATCCAATCACCTACTGTTTCATGAATAAATCATTTCGCTCCT

CGGTATTAGCTTACTGTCGTCCAAAACCAAAACGCCCATTGACTCGTTGCTCTGCTCTACCAACTCGTAAGAGCCCTTCA

TCTCCAACTCATGCTATTCCAATGATTAAAATTGATCTTGTACTTGATAGCAGTGTTCACATTTAA

>pR122

ATGGAAAGCGACTCCCCACTTAATCATCCTGTGGTACACTTTTTGCTCTGTGTATCCTACCTTTTTGTCTTTCTGGCCAC

TATAATCGGAAATCTTATCATTCTGCTGGTCTTCTTCTCGCAGAAACGAGTCCGAAGTGTGCCCAACTTCTTTCTGGCAA

ATCTGACGGTTGCCGACCTTTTTGTCGGAATCTTCTGCGTTCTCCAGAACGCTGTGAATTTTGTATTCCAAGGACATGGA

CGGTGGCCGTTTGGCAAAGTGTTATGTTATTCATACATCTACGTACTACACTTCATACCAAATGTGTCGGCTGGAATTCT

GGTGCTGGTCTCGGTTGAAAGATTGATTGCGGTGCTCCGTCCATTGCGGGTGCGAAGAGCATTTTCACAAAAAGTGCTCA

TCACATCGTCAGCTGTTGTTTGGCTAACTAGCGCAGTTATGAACACCCCATATGTAGTGGCTAGCCAGTTTTTAGAGATA

TCAGACGGGAATGAGACATACACAATCTGCACTAGACGTCATGTGGAAATTTATGGATTCAACCTCCTGAAGCTAGTCGC

CACTATCAACTTCATCGTCTGGTACGCCGTCCCTCTAGTAATCCTTCTTTGCATCTACGCTACCATCGGCCTCGTGGTGA

GCAAGGCCGCAACAATCACGAAACACTCCGCCAACAAGCTTTTGCCAAGATCAAGCTGGCGATCCGAAGCCTCAACCGGC

GGTGTGTCAGTGGAGAAGAGGAGAAAGGTTGGTCGTTTGGCAGTCGGAATCGTCGTGGCATTCGCGTTCTTCTCACTCCC

GAGATATGTATATTTTATGTGGACCGTCTGGCGAGATCCGATGGCGCCAAGATGTTTGAACTGCCTTCAAAGTGTACTCC

AACCCATCAGTTTCCTGCTTCTGTTCTTTAATGCCGCCGTCGACCCATTCCTCTACGCATTCCTTTCAACAAGATTTCGT

CAAAGCATCAAAGAAACATTTCATCTTGGAAAGGTGCGTGAAAGCAACATCGAGCAGTCATCACGAGTGCGGCGTAAGGT

GTTAAACTCGGAGTTGGAGACGGGTTACGAATATCATAATCACTCATAA

>pR123

ATGTCCGTAGCGGTGGGCATCCCCTACGTATGTTTCTTCATAATTCTCTCAGTCGTCGGAATTATTGGAAATGTGATTGT

GATCTATGCGATCGCAGGTGATAGGAATATGAGAAAGTCTGTGATGAATATCTTGTTGCTCAATTTGGCAGTGGCAGATC

TTGCCAATTTAATATTTACAATACCTGAATGGATACCGCCCGTCTTTTTTGGAAGTACTGATTGGCTCTTCCCATCATTT

CTGTGTCCAGTATGCAGGTACTTGGAATGTGTTTTCCTATTTGCATCAATCTCCACACAAATGATTGTTTGCATTGAAAG

ATACATTGCAATAGTCCTCCCAATGCAGGCTCGCCAGCTGTGTTCCCGCCGAAATGTTCTGATAACAGTGCTTGTTGATT

GGATCTTTGTCGCCTGTTTTGCATCTCCATATGCTGTGTGGCATTCCGTCAAGACTACCACCTGCTCCAACACAGTTGGA

AAATCCACTTGGTGGCAAGGATACAAGCTGACCGAGTTCCTCGCCTTCTACTTTGTACCATGCTTCATCATCACAGTAGT

TTACACAAAAGTTGCAAAATGTCTGTGGTGCAAGGATCCGACACTTCAATGTGAAACTCGTTCGTGCCTGGACAACAAGT

CATCATCACGAAGCTCCGATGCTCTCCGTACTAGGAGAAATGTCGTCAAAATGTTGATTGCATGTGTGGCAGTGTATTTT

GTGTGTTATTCGCCGATTCAAGTGATATTCTTGTCGAAAGCTGTCCTAAACGTGACAATCCATCCTCCATACGACTTCAT

CCTCCTCATGAATGCTCTTGCAATGACTTGCTCGGCGAGCAACCCTCTTCTTTACACCCTCTTCTCACAAAAATTCCGTA

GAAGGTTACGTGATGTCTTGTACTGCCCGTCTGACGTGGAAAATGAGACAAAAACCTACTATTCAATCAATAACACCTCA

ATCGTCGGTCCACGGGCTAGTTTTAATTGA

>pR124

ATGTTGGACTACAATAACGTTACGTCAGGTTCATCAGAAGAGGACTATGATACAAGTGAAATAATTGGGTTGCAAGTTAA

TACGCACGAAGAATATGAGGATATTATTGCAGAAGCTTTATGGCCAAATGCTGTTGAACAGTGGTTCGTATACATATTGG

CAAGTATGATGGTGATCGGAGTGATTGGAAATACGTTGGTCGTTGTTGTAGTGGCAACTAATAAGTCTATGCGCAACGCT

CTCAACTTGGTACTGATGAACTTGGCAATTGCTGATCTTCTCATATTACTATTCTGTCTTCCATTAACAGTGGTGAATGA

CGTCACCAAGACTTTTTGGTTTTCGGCGGTTTTTTGTAAAAGTGTTAATTTTGTAAATAATACATCAGTGTACGTGAGCA

TAATGTCATTGGTATTCATCACTTGTGAAAGGTGGCGAGCAATTACATATCCCTTAAAAAGCCCATTTGTTAGAACGAGA

TCTGTCATTGGAGGAATTTGGTTCATTGCAATGTTCCTCTCGTCGCCGGAGCCCGTCACGCTACATCTCGCCGGAGCCCC

ATTTGTGCGACCTAATTTTACAACAAAGTGGGGAACACGTTGCAAGGAATCGTGGTCAGAAGAATTCCAGAAAAACTATC

AACTACTACAGACAATTTTCTCATTTGTACTTCCACTTCTTGTCATATCAATTTTATGTCTGCATATGGTCCGAACCTTG

CATTTCTCTGCAAATTACTTGACAGTAGCAAACCGACAAATCAGCATACGAAAAAAAGCGGTCCGGATGCTATGTGCAGT

GGTTTTTCTCTTCTCAATGTCTAATCTACCTGTTCATTTGTACAATATTGCGCTAAACTACGATTTACTGTCAACGGATG

TGAGCACTAATACAATTGCAGTTCGGAAGCTCCTGCCACGCGTTTTTTCATATTCTTCATCATGTCTCAACCCCATTCTG

TACAGCTTTCTAAGTGGAAGTTGTCGAATTCTGGCGGATATCGATTGA

>pR125

ATGTTGGACTACAATAACGTTACGTCAGGTTCATCAGAAGAGGACTATGATACAAGTGAAATAATTGGGTTGCAAGTTAA

TACGCACGAAGAATATGAGGATATTATTGCAGAAGCTTTATGGCCAAATGCTGTTGAACAGTGGTTCGTATACATATTGG

CAAGTATGATGGTGATCGGAGTGATTGGAAATACGTTGGTCGTTGTTGTAGTGGCAACTAATAAGTCTATGCGCAACGCT

CTCAACTTGGTACTGATGAACTTGGCAATTGCTGATCTTCTCATATTACTATTCTGTCTTCCATTAACAGTGGTGAATGA

CGTCACCAAGACTTTTTGGTTTTCGGCGGTTTTTTGTAAAAGTGTTAATTTTGTAAATAATACATCAGTGTACGTGAGCA

TAATGTCATTGGTATTCATCACTTGTGAAAGGTGGCGAGCAATTACATATCCCTTAAAAAGCCCATTTGTTAGAACGAGA

TCTGTCATTGGAGGAATTTGGTTCATTGCAATGTTCCTCTCGTCGCCGGAGCCCGTCACGCTTCATCTCGCCGGAGCCCC

ATTTGTGCGACCTAATTTTACAACAAAGTGGGGAACACGTTGCAAGGAATCGTGGTCAGAAGAATTCCAGAAAAACTATC

AACTACTACAGACAATTTTCTCATTTGTACTTCCACTTCTTGTCATATCAATTTTATGTCTGCATATGGTCCGAACCTTG

CATTTCTCTGCAAATTACTTGACAGTAGCAAACCGACAAATCAGCATACGAAAAAAAGCGGTCCGGATGCTATGTGCAGT

GGTTTTTCTCTTCTCAATGTCTAATCTACCTGTTCATTTGTACAATATTGCGCTAAACTACGATTTACTGTCAACGGATG

TGAGCACTAATACAATTGCAGTTCGGAAGCTCCTGCCACGCGTTTTTTCATATTCTTCATCATGTCTCAACCCCATTCTG

TACAGCTTTCTAAGTGGACGATTCCGCGAAGAGTTCGGCCGAGTAGTATGCTGTCTTCGGGATGAGAAATCAAGCCGTGA

ATTCCGAAGAAAACAGGCTTCACTGTACGCTTCAAGTCAAAGAGTTCGAACGAATAGTTGCTCAACGATGCTTTTGGATC

GTGAAACCAGATCATCACTGATTATTCGGAGAGTGTCGTTGGCACAGGCGATACCTTCATCTCCTCTAATTGCTCATCAG

AATCCACGTCTCTTTTCACTTGATGACGTACGTGAGGAAGTGACAAATTCAGAGGCTCTTGTGCTCCAGAACTGCTGA

>pR126

ATGGATTCGAGTTCACTATGTCTTCCCGAGTTACAGCTCCGGCTGTCGGGGTCTAATTTGGAGGCGTTCCTGTATACGGT

GCTGTTTCCGCCGATTTGTCTGTTCGGAGTGGTGGGCAATGCGTTGACAATTCTTGTGCTGGTTAACAACGATTTTATGT

CAAGAGCCAACATCTTCCTGACATGCCTGGCCGTATGCGATGTCTCCTTCCTGATTCTGATCATTCCGCACTCGCTGGCA

AATTTTGACCGATTCGCGTTCAACTACACCTTCAGGTACCTGTACCTACCGTCTAAAATGCACCTCATTGCATTTGCCAA

TTGGACATCTGCCGTGGCGATATGGCTCGTCGTCGGTGTGTGCTTCGAACGTGTGGCCGGAGTACGGTCTCCGCTCCATC

GGCTGACAACTCCGTCACGTGGCAAACTGACAACCGGCCTACTGACATTGCTGACATGTTGTGCCGCATTGACATTCTAC

AATCATGTCAGCCATCACTGCTTCATCAAGTCGTTTTGCAACGCAACACAGATTATGGCCGTCTGTTTGGACGTCAATTT

GGATGTATGGCCCAACAACCGTACAAACATCTCACCGCCGGCGTTGCGTACCTACGTGGCGGCGACGCGTGCCGCCAACG

CGGCTCTCGTAGTGTTTCTGCCGATGATTCTCCTCGTCGTGCTCAATATGATGCTCTTGTACTATGTAAAAAAGCGATCA

TTTTTCATGTACGCCTCACTGGGAAGGGTCTCGGCACGAATGCGGAAGTCCGGCGGTGACGTGGCACTTCCGTTTGTGGG

GACACTGTTTAGGAGGCACTCGGATCAGATTCTTCAGCAACGAACCGAGCACCGAGTGGCCGTTACGGTATGTGCCGTGG

TGACGTCATTCACAATAACTCAGGCACCGTCCGCTGTGGTGTTAACTGTAAATTCGCTGCTCAAGGAGCGGCTCGACGCG

CATTGGTACCACATGACAACAATTACCTCATTCTTGGTTGTAATCGGAAAAAGTTTGAATTTCGTCGTTTTCTGCCTATC

ATCTTCCAGTTTTCGGAGGCGGTTAACAAATACGCTACGCTCGAAATATGGGTCTCGAGACGAAAAACGTTCCCACTCAA

TCAACACCACCTACACCCGTTGCGATAGCCGAGATTCGCGGATTTTATCCCAGAAAAAATCGTTCCAAAGTATTTAA

>pR127

ATGGATTCGAGTTCACTATGTCTTCCCGAGTTACAGCTCCGGCTGTCGGGGTCTAATTTGGAGGCGTTCCTGTATACGGT

GCTGTTTCCGCCGATTTGTCTGTTCGGAGTGGTGGGCAATGCGTTGACAATTCTTGTGCTGGTTAACAACGATTTTATGT

CAAGAGCCAACATCTTCCTGACATGCCTGGCCGTATGCGATGTCTCCTTCCTGATTCTGATCATTCCGCACTCGCTGGCA

AATTTTGACCGATTCGCGTTCAACTACACCTTCAGGTACCTGTACCTACCGTCTAAAATGCACCTCATTGCATTTGCCAA

TTGGACATCTGCCGTGGCGATATGGCTCGTCGTCGGTGTGTGCTTCGAACGTGTGGCCGGAGTACGGTCTCCGCTCCATC

GGCTGACAACTCCGTCACGTGGCAAACTGACAACCGGCCTACTGACATTGCTGACATGTTGTGCCGCATTGACATTCTAC

AATCATGTCAGCCATCACTGCTTCATCAAGTCGTTTTGCAACGCAACACAGATTATGGCCGTCTGTTTGGACGTCAATTT

GGATGTATGGCCCAACAACCGTACAAACATCTCACCGCCGGCGTTGCGTACCTACGTGGCGGCGACGCGTGCCGCCAACG

CGGCTCTCGTAGTGTTTCTGCCGATGATTCTCCTCGTCGTGCTCAATATGATGCTCTTGTACTATGTAAAAAAGCGATCA

TTTTTCATGTACGCCTCACTGGGAAGGGTCTCGGCACGAATGCGGAAGTCCGGCGGTGACGTGGCACTTCCGTTTGTGGG

GACACTGTTTAGGAGGCACTCGGATCAGATTCTTCAGCAACGAACCGAGCACCGAGTGGCCGTTACGGTATGTGCCGTGG

TGACGTCATTCACAATAACTCAGGCACCGTCCGCTGTGGTGTTAACTGTAAATTCGCTGCTCAAGGAGCGGCTCGACGCG

CATTGGTACCACATGACAACAATTACCTCATTCTTGGTTGTAATCGGAAAAAGTTTGAATTTCGTCGTTTTCTGCCTATC

ATCTTCCAGTTTTCGGAGGCGGTTAACAAATACGCTACGCTCGAAATATGGGTCTCGAGACGAAAAACGTTCCCACTCAA

TCAACACCACCTACACCCGTTGCGATAGCCGAGATTCGCGGATTTTATCCCAGAAAAAATCGTTCCAAAGTAAAGGGTCA

AGACAATTCTGCAGATATCCAGCACAGTGGCGGCCGCTCGAGTCTAGAGGGCCCGCGGTTCGAAGGTAA

>pR128

ATGCTTCTCCCTTCAAACTTGACGACGTCAACATTAATGACGTCATCATCAGAATCGTACGACGCTGATAATCCGGGGCT

TCCGCCTGAGCCAATCCTGTCGGATTACGTGGAAATGTTCACTTTGGTGCTCAATTTTATTGTTGGCGCGCCGTTGAACC

TGGCCGCTTATACACAGCTAAGCGAACGACCTACATCAACGCGGTTAGACCTTCTGAAGCGATCACTCAACTATTCGGAT

CTTCTCGTTCTATTCATCTACGTACCATCTCGTGCCTGCTGGTTATTAACCTACGATTGGCGGGGTGGAGATGCACTCTG

TAAAATTGTCAAGATGTTTCATACGTTCGCGTTTCAGAGCTCCTCCAACGTGATCGTGTGCATCGCCGTGGATCGCCTGC

TATCCGTCCTCTCCCCATCCCATCACAGCCCCAACAAAGCCCTGAAACGGACTAAAATGATGTTAATAGTCGCGTGGATA

GTAGCGCTAGTAATCTCATGCCCACAACTTTTCATCTGGAAAGCATATCTAGCACTTCCCGAGTATAATTGGAGCCAGTG

TCTGCAAATTTGGGAGATTGCACGGATGGAAAAATTCAACAAACCACAGGTAGTGCCAGAGTTTGACGCCGAGTTCTGGT

ACAGCATACTGCATATTAGTCTCGTTTTTTGGATCCCTTGTATCATTATCATGCTATCCTACATCATAGTCATCTCATGG

GTATGGATCAACTCTCGGCCGTCCATCCGTCACACCTCTTCATTTTCCTTCCACACCGGCTGCGATACGGTAGATACAGT

ACTGACTAGAGCCTCTGAATGGAATCCTTTGAAGACATTCTCCCGTCACGTCAACATCAAGGAGCCCGAGAAGCCGATGA

CGACTCCCAGAATCGTGGTCAGCGACGAGACGGAGGTCCCACTGACGCAGCGACCATCGATTTCTCCGTCGGAAGCGTCG

GCGGTGATGAGGACCGGTGTGCACACGAGTACCTCGTATAATGCTAATTTGAATCGATCCCGAGCCCTGCGAGTTTCCTT

GCTACTAGTCGTCGCGTACATCATCTGCTGGCTACCATATAACCTCATAAGTCTTATCCAATTTCTTGATCGGGACTTTT

TTTCGTCATATCTTAAACATGTCCACTTCTGCCAACAACTAATCATTTTTAACTCGGTCGTCAATCCATGGCTCTACGGT

TTCTTCGGTCCCCGCCGCCCGTCTACCACCGGTGCCGGCCGTCACTGA

>pR129

ATGAATATTTCCACAATTATCTCGACTACAACAACTCCACCAGCGGTTTCAAACGAAGATGACACGTCAAGAACCACGTT

GAAATTGGTTTTGGCAATTTCCTATTTGGTTCTTTTTATAATTGGAACAGTCGGAAATGGTACCGTAATTCTCATGATCA

TCAACATTTTGACATCTATGAAGCGGAACGTGCAGGCTGGAAAACGGATGATGAGCAGTACGAATCATGTGTTTATTTAT

GTTCTTGGACTCTCGATTGTTGATCTCCTGGTAATTCTGCATCTTCCGTTTCTGGTCGTAGATCTTTTGAAAGGACAATG

GCTATTTGGAGTGGCAATGTGCAAAGTGTACTGGTTTGGGGAAAGTGTCAACAAATTGTTGAGCTCTTTTTTGATGACAG

TGCTCAGTTGGGACCGGTATATGGCAGTTTGCAGTCCGGTTAAATCTATGAAAATGCGTAGCAATGCAACCGCATTGAAA

GTCCTCCTGGCTTGTACCATATTTGCCACAGTACTCCTTTTACCAGTACTCTACGAGGCCGCCGTATTTAAAATCGATAA

AATGAGAATGGTTCCGTTGTTGGAAGGTACGGAAGAACTGGCAGCGTCGGACCTTGCTGGCACAACAATGTCAAAATGCA

TGTTCGATGCGGATGCCACGTTTACTCTCTACACATTTGTCATCGGATTTGCCGCCCCCGCATTTTTAATTATAATATTC

TACGTACAGGTAATTTATGCTCTTCAAAAATCATCTCGAAATATCCGAGGTGCGCGAGGAATCAGCAAACCGGATGGAAG

CTCGAATAGGGTCAAGAAGGTCACAAAACGAATAGTGGCTGTGATTCTTTTCTACTTCCTGTGTTGGACACCTCAATGGA

CACTGAATATCATGTCACAATTCAATTTGATCGCCGTTTCATGGATGACTCCAGCACTTTCGGCAATGTTTTTTGTGGCG

CATTTGTTGGTATGTTTCAACTCGGCAGCTAATCCAGTTCTCTATGCATACATCAATCGGGAACTTCGTTCCCAACACGT

CATGGCAATGCACAGAAAACGACACTCGTTAGGTCATGAGCAACAGCAACGATTAAATCAGGCGAGACTTTCCATGCAAT

CCCGTCCTTCTGACGCTTCAGCTACAGTTTTCTCGAAACTCGGCGACAGTGACGCCGGTATTTTCAACAGGCTCTCAAAT

TTCCAACAAAACATAATTGCAAGCATTTTACACCGCAAACGAATAATACTGGTGGCGGATAAAAATATTCACATTGAAAA

CACGTCATCATCGACAGACGAGGATAAGGACGATTACTTGTAG

>pR130

ATGAATATTTCCACAATTATCTCGACTACAACAACTCCACCAGCGGTTTCAAACGAAGATGACACGTCAAGAACCACGTT

GAAATTGGTTTTGGCAATTTCCTATTTGGTTCTTTTTATAATTGGAACAGTCGGAAATGGTACCGTAATTCTCATGATCA

TCAACATTTTGACATCTATGAAGCGGAACGTGCAGGCTGGAAAACGGATGATGAGCAGTACGAATCATGTGTTTATTTAT

GTTCTTGGACTCTCGATTGTTGATCTCCTGGTAATTCTGCATCTTCCGTTTCTGGTCGTAGATCTTTTGAAAGGACAATG

GCTATTTGGAGTGGCAATGTGCAAAGTGTACTGGTTTGGGGAAAGTGTCAACAAATTGTTGAGCTCTTTTTTGATGACAG

TGCTCAGTTGGGACCGGTATATGGCAGTTTGCAGTCCGGTTAAATCTATGAAAATGCGTAGCAATGCAACCGCATTGAAA

GTCCTCCTGGCTTGTACCATATTTGCCACAGTACTCCTTTTACCAGTACTCTACGAGGCCGCCGTATTTAAAATCGATAA

AATGAGAATGGTTCCGTTGTTGGAAGACCTTGCTGGCACAACAATGTCAAAATGCATGTTCGATGCGGATGCCACGTTTA

CTCTCTACACATTTGTCATCGGATTTGCCGCCCCCGCATTTTTAATTATAATATTCTACGTACAGGTAATTTATGCTCTT

CAAAAATCATCTCGAAATATCCGAGGTGCGCGAGGAATCAGCAAACCGGATGGAAGCTCGAATAGGGTCAAGAAGGTCAC

AAAACGAATAGTGGCTGTGATTCTTTTCTACTTCCTGTGTTGGACACCTCAATGGACACTGAATATCATGTCACAATTCA

ATTTGATCGCCGTTTCATGGATGACTCCAGCACTTTCGGCAATGTTTTTTGTGGCGCATTTGTTGGTATGTTTCAACTCG

GCAGCTAATCCAGTTCTCTATGCATACATCAATCGGGAACTTCGTTCCCAACACGTCATGGCAATGCACAGAAAACGACA

CTCGTTAGGTCATGAGCAACAGCAACGATTAAATCAGGCGAGACTTTCCATGCAATCCCGTCCTTCTGACGCTTCAGCTA

CAGTTTTCTCGAAACTCGGCGACAGTGACGCCGGTATTTTCAACAGGCTCTCAAATTTCCAACAAAACATAATTGCAAGC

ATTTTACACCGCAAACGAATAATACTGGTGGCGGATAAAAATATTCACATTGAAAACACGTCATCATCGACAGACGAGGA

TAAGGACGATTACTTGTAG

>pR131

ATGAGTTGCATGGGTCTCAACACGACGGAGCTGCAGGAGGCCAGGGACTACTTTCGGCAGCTTCTCCTGCCATTCACCGC

CTCATTCACAATGTTCCACAACTACATATACGTGTTTCTATGTGTCATCGGAGTATTTGCAAACATTTCAATCGTCGTCG

TGCTCCTGCGACCGGCGATGCGAAAATCGCCATTCAACTTGTTTCTAGTAGTGATTGCCGTCTGTGATGCCTCACTAATG

GCCACATATTTGACATATAAACACGTGGAATTGTGCCATCCATGGTATTTCTCGTTTCCATGGGCGATTTATACAAAGTT

CTACGCTATTTTCTCGGTTTTCGTGCATTCGTGCAGTCTATGGCTCACCGTTAATATGGCTATTCTGAGATTCCTGGTAC

TGTACCGCGGAAGTCGTTCCGAGACCCGAATCCCTCAGTGCAATGGATTTGGCGCTGCATTTGTGGCAATTGTGCTAGCA

TTCGCGATAGCTTGCATCGGATGTCTCCCGATTTTCATCCGCTATCGAATAATCGAAGGAGAAGTTGGCGCGGTGCCAGA

TTTATGTCTGGAGGGAAAATATTCAACTGATTGGCAGCCAAATGATATGATTCAGTTCTACGGGTTAAGTCAGCCGCTTT

GGTGGAATTGTGACTGGGAACGTATCAATTATTGGATGGCTGCACTAATCCTAAAACTGATTCCCTGCCTTCTGCTCACA

ATCTTCATGACACTACTCGTCCGAATGCTCATCGAAGCAAGAGAACGAAGGTCCCGACTCTGCGGTGGAATGGGTAATGG

AAACTCGCAAGCCGAGAGGACAACTGCAATGTTAACTGGAATTGTTGCAATCTTCCTTATCACAGAGCTCCCACAAGGTG

TGCTCACATTTGCCGCCGGAGCCAACCCTCGGCTCACATTCCTCACACTTCAAATGAACAATGTTTTCGACTTGCTCTCC

CTAATTAATAGTGCCGTCAATTTTGTGCTCTGCGCTCTGATGTCACATGTTTTTAGAAGAGAATTCCTCCAAACATTCGG

TGTCTGCTGTCCCCAATCGTCGGAAAATCACAGCGGAGCACCAATCAACAAAACCTCAAATCGATCGATTCTCTCCACAT

TCAGCAACATTGCAAAACCAAAGACTTCCAAGAAAAATGGGTTCCTGCCAGTGCCAACTAACTGCCCGGACGACAAGAAT

ACCGAGCAACTAGCGATGAATTGA

>pR132

ATGTACGGCTTCTTCAACTCATCGGAAATCGACGAAGAGTACTACAACTCGACACACACCTCTGCACCATCACCGATTCT

CGCTCTGATATTCACAATTATTTGCATTATCGGAGTGATCGGAAATGCATCACTGCTCGTGTACATCTTCGCCAAGAAGC

TCTACCAAAACTTTATCTCGTCTCGATTCATCGGACACTTGTGCTTCACCAACTTGATCGCGTTGCTCGTATTGGTGCCG

GTGATCATTCATAACGTGTTCACCGGTGTCAACTTGCTTCAGGACAGCAATATGTTGTGTCGGATTCAGGTCTCCATAAC

CGTCACCGTCTGGACCGTCATCGCCATGATGAACCTCTGCATCGCCGGTGTTCATCTGCTAACGTTCGCCCGCATCCACT

ACGAGCAACTTTTCGGCCTGACACCCACCAAACTATGCATCTTGTCGTGGATAATCTCCTGGCTCCTCTCTCTACCGTCT

CTAACCAACGGCCACGTGGCGATCTACGGCCCAGCGGTCCGAACTTGCGTCTTCTCACACTCGGACTCTGGCCTAAAATT

CTTGACGTACACCATGATCTTCGGCGTGTTCATTCCGGCCTTATTCTCGTCGATCGCCTACTTTCGAATTCTACAGACTC

TATTTCATAGTCCAATTGTGTTCCAGTCGCTGGGCTTGTACAAGAGCAGATTCCTCGTGTATTTCTTCCTTTTGGGTCCC

CTCTACGCTCTGCCCTTCTACATCCTAACCGCCTTGGACCCGTCGGATCCCGTCCGAATGAGTCAGGACACACTGTGGAC

CATCGGATGCACCATGTTCGCCTTCGTACCGTGCATCATCGCTCCGTTGCTCTACGGAGCCAGCATTTTCATCATAAAGG

AGGAGGATATGGCTCTGACGGCGAGGACTCACACCAAAACCGGCACCGGAGCCTATCACCACGTCGCACATCAGCACAAT

ATGCAGGCTCAGCTGATTTGA

>pR133

ATGTACGGCTTCTTCAACTCATCGGAAATCGACGAAGAGTACTACAACTCGACACACACCTCTGCACCATCACCGATTCT

CGCTCTGATATTCACAATTATTTGCATTATCGGAGTGATCGGAAATGCATCACTGCTCGTGTACATCTTCGCCAAGAAGC

TCTACCAAAACTTTATCTCGTCTCGATTCATCGGACACTTGTGCTTCACCAACTTGATCGCGTTGCTCGTATTGGTGCCG

GTGATCATTCATAACGTGTTCACCGGTGTCAACTTGCTTCAGGACAGCAATATGTTGTGTCGGATTCAGGTCTCCATAAC

CGTCACCGTCTGGACCGTCATCGCCATGATGAACCTCTGCATCGCCGGTGTTCATCTGCTAACGTTCGCCCGCATCCACT

ACGAGCAACTTTTCGGCCTGACACCCACCAAACTATGCATCTTGTCGTGGATAATCTCCTGGCTCCTCTCTCTACCGTCT

CTAACCAACGGCCACGTGGCGATCTACGGCCCAGCGGTCCGAACTTGCGTCTTCTCACACTCGGACTCTGGCCTAAAATT

CTTGACGTACACCATGATCTTCGGCGTGTTCATTCCGGCCTTATTCTCGTCGATCGCCTACTTTCGAATTCTACAGACTC

TATTTCATAGTCCAATTGTGTTCCAGTCGCTGGGCTTGTACAAGAGCAGATTCCTCGTGTATTTCTTCCTTTTGGGTCCC

CTCTACGCTCTGCCCTTCTACATCCTAACCGCCTTGGACCCGTCGGATCCCGTCCGAATGAGTCAGGACACACTGTGGAC

CATCGGATGCACCATGTTCGCCTTCGTACCGTGCATCATCGCTCCGTTGCTCTACGGAGCCAGCATTTTCATCATAAAGG

AGGAGGATATGGCTCTGACGGCGAGGACTCACACCAAAACCGGCACCGGAGCCTATCACCACGTCGCACATCAGCACAAT

ATGCAGGCTCAGCTGATTAAGGGTCAAGACCAATTCTGCAGATATCCAGCACAGTGGCGGCCGCTCGAGTCTAGAGGGCC

CGCGGTTCGAAAGGTAAGCCTATCCCTAACCCTCTCCTCGGTCTCGATTCTACGCGTACCGGGTCATCATTCACATTCAC

CATTGGAGTTTAAACCCCGCTGA

>pR134

ATGGAGGAGCCCATAAGTTGTCTAACGTCTCCCGAGCAGCGGTTAATTGAGGAACGATTCTACATAAATGGCGTGTTCGG

TACAGTAATTGCAGTTTTCGGGGTCATCGCCAACGGATTCCTAGCCACCCTATTTTTAACCCGAACAATTTATAGGAATT

CTCCGTTCTTTTTTCTCGGTTTTGTTGCCCTTTTCGACACACTGGTGGATGCCACCTTTTTATTACTTTTGCCTCTAGAA

GTCAGCTCGGTGTACTACGAAGAAAGTGTGGCCTATCAAATTTGGCTGAAATACGTCAGACATGTGTATTTATTCGCGCA

AATTGTGAAGATCTCCTCTGTGTTTTCATTGATCATGGCGAGTATTGAGAGATACTGCATGACGAAGCATTGGACATTTG

TTGGCTTTGAGATGAGAACCCGTTGGCTCGTGCTCTTTTCACTTGTCTGTGCAGCTGTATTTATAAAATATATGTCAGAA

GAAGCTGTAGTTGTCACTCGAGAAAACTGTTTCGGCTTCCGGAGATGGGCGGTGCGAATTCAACGCGGAGAGCTCGGCCA

TATGGGGTGGCTCCACTCACTAGCGATTTTTCTCCCATTCTTCACACTAATCTTTCTTAATGGAGGAATTGTTCGAATGC

TCCGGAAACAGAATGTTCAACAGCTCCGGAGCCTGATTGCAGAGCTCACTTTGGGACACGACCTGTCGAAAATTCGGAAA

AAGAACTTGAGAAGTGCAACGAGGACCCTGATTGTTATCATCTCGGCGTATTTGATTTCGAACTTGCTCAGCTTGATTTT

GATCATTGTAGAGTACTTTAATCCGGATTACCTCCACGTCTACCATCCTGACTTAAACCGCCTGGCAACCGACTCTGCAG

CTCTTCTGACAGTAGTCGGAAATGCGATTCGATGCCCCGCTCATATTTTCAGCAATTCTGAAATTCGAACACAATTTCGA

ATAATGTTATGTGGCGAGAGAGAGAAAAAGGTAGTGAAAAAGCTCTCGGAACGCCGCCGAACACAGGAGAAGCTCGACAA

TCCGTGGATGGCTCTGATGTGGGTTTCAGCCAATAATCAGGCGTCTGATGATGATAAAGCCGCATTTATGCCACGTGCAT

TCATGAAACGACACTCGGCAATCGCTATTGTATAA

>pR135

ATGTCAACGGTACATCAGGAGGAAATATGCCGGTTTCGTGGTACAACAGAAAATTACACAATCGCCGTGACGTTTTTCAT

GATTTTTTTGCTATCAGTCGTCGGAAATTCGGTGGTTTTGATAGTGATTATTAAGCAACGTGCAATGCGATCAATAACCA

ATATATATTTAATGAATCTGGCGGCATCGGATATGATGTTATCAGTGGTTTGTATGCCGCCGACACTTGTCTCCATGGTT

ATGAATTGTTGGATGTTCGGCAATTATATGTGCAAAATTTTGGCTTATTTGCAACCCGTAGTAGTCACTGCTTCAGCGTA

CACCCTAGCAGTAATCGCATTCGAGAGGTACTTTGCAATTTGCAAGCCGCTTCACTCAAGGATTTGGCAAACCAGATCCC

ACGCCTACGCAATGATCACACTTGTGTGGGTGATCGCCATTGCCGCCAATATTCTGATGCTTTTTATGTACGAACAACAA

ACGTATAGCTCGAATGGATACACGTGTGCTCCAATTCATCCGCCGATTTATCATTTTGCTTATCAGGTATACATGACCGT

CGTCCTACTTGTAATCCCGCTAGTTGTGATGGCGGGTTTGTACGGCAACGTCATTACCTCCCTAAAATCCGGCATCAAAC

TGGAAATCGCCTCTGTGGATCCGCCGCTCGCCACCGCCACAACCACAGCAATTGTTGCGTCGATGACCGACGAGCAAAAG

TTATCGTTCTGGAATAAGCTGTCGAACAAGCTGACTTTTAGTCAGCAAGATAAAACCGTGCAACACCCAAACTTTGGGCA

TCGCAAATCCGACACATCAATTTGCCTCGAAAATCCTAGTTTAAGGTCTACGCACACTCAGAAAAGTGCAATGGCAAAGC

AAAGAGTGATTAAAATGCTCATTGTTGTCGTAATTATCTTTTTCTGTTGTTGGACGCCTTCCTACATCTGGTGGTTGTTA

TTGATCGCGGGAGACTCGTTTCAAAGCCTCAACTTATCTGTCTGGAACAGCGATATCAACACATTCATCACTCTTCTTAC

CTATATTTCTTCATGCACCAATCCCATCACATATTGCTTCCTGAACAAAAAATTTAGGAATGCGGTGTATGCAACATTTG

GCAGGAAGAAAAATATGCGACATCATTTTCAGAAGGTCGGTCTCGTCGACATTCACTCCACAACATCAAGTCGTTTTTAA

>pR136

ATGGATAGTAACAATATCAGCTTCGTAGAAGAAGTATTCGAGGATATAGAAGATGAGTGTGCCTACTCTCCACCAATATT

TCTCGAGTGGAAAATGGTTTTTGTGGGTGTGATTGGATTAATAGTTGCACTCATCAGTATTGTTCACAATTCACTTCTCT

TCTACACATTTAGTACATCTACAGTACTCCGGAAACGAAACCTGACATATTTGATGTGGATTTCCGGATGCGATGTTTTC

ATCTCGATTTGTTATATTGCAATTATGTGTGTTCAGGTGTACACCGACTATTTCGTATCATTCAAGTTATTAGTTTTATG

GCATGACTATTTACGGACTGCTTTCACAGTGTCTCATATCACCTTATCGGCTGCTTCATTTCTTCTTATGGCCGCAACAG

TTGAAAGATATCTTCAAAGTACTGCCGATAACAGGACTCTGAAGTTATTTCAAATTCTCGCAACCCACAGGACATATGTT

GTTTTTTGTTGTTTCCTAGCGAGTCTCATATTTCGAGGAACCGTGTTTTTTGAGATTGAGGTTGTTACTCAACCGAATTG

TACGGGATTTTCATCAATGGGTTTGGCAGTTGTACCAATGTTTGGAGAAAGTATGGATGTTGTTTGGAGATTTTTTATTC

GAAAAATTGTTACCGTTTTCTTGCCATTTGCAGTTTTGGCATACTTCAATGCAGCAATTGTAATGAATGTGCGAAGGACT

GATCGGGATCAAACAGTCAAAGCATTGGTTTTATTTATTACTGTTGGAACAAGAGGAGAAGTTACACGATTGAGATCACG

GCTGAGAGCTGTTACAAGAATGCTTGTTATGGTTGTTACTGGATATCTACTTGCGAATATTTTAGATATTATTATTGCGT

TTTGGGAGACAATCAATATTCAAAGTCTTCAAGAACTTCCTTCATTATATACAGTACTTTCGGACATTTCATCATTTTTA

CCAATTGCTGCATGTGCTCTTCGTCTCCCTATCTACACAATTAATGATCGTCAAATTAGAGTCGAGGTTCGAAGAAAGTT

TTGTTATCTCATAACGAGATGTTGCACATGTTGTCCAGGAGTTCTTTGCGATACTCATCAAAGAAAGAAACTTCATGATA

ATCAGTTGGAATCGGAAAGACTTTGGCCCAAGAATGAGCCAACTCCAAAACTTGAGCATGAAATGACACAACTATGGACA

AAGTCTACTGAAAGACTTGGCTGA

>pR137

ATGGATAGTAACAATATCAGCTTCGTAGAAGAAGTATTCGAGGATATAGAAGATGAGTGTGCCTACTCTCCACCAATATT

TCTCGAGTGGAAAATGGTTTTTGTGGGTGTGATTGGATTAATAGTTGCACTCATCAGTATTGTTCACAATTCACTTCTCT

TCTACACATTTAGTACATCTACAGTACTCCGGAAACGAAACCTGACATATTTGATGTGGATTTCCGGATGCGATGTTTTC

ATCTCGATTTGTTATATTGCAATTATGTGTGTTCAGGTGTACACCGACTATTTCGTATCATTCAAGTTATTAGTTTTATG

GCATGACTATTTACGGACTGCTTTCACAGTGTCTCATATCACCTTATCGGCTGCTTCATTTCTTCTTATGGCCGCAACAG

TTGAAAGATATCTTCAAAGTACTGCCGATAACAGGACTCTGAAGTTATTTCAAATTCTCGCAACCCACAGGACATATGTT

GTTTTTTGTTGTTTCCTAGCGAGTCTCATATTTCGAGGAACCGTGTTTTTTGAGATTGAGGTTGTTACTCAACCGAATTG

TACGGGATTTTCATCAATGGGTTTGGCAGTTGTACCAATGTTTGGAGAAAGTATGGATGTTGTTTGGAGATTTTTTATTC

GAAAAATTGTTACCGTTTTCTTGCCATTTGCAGTTTTGGCATACTTCAATGCAGCAATTGTAATGAATGTGCGAAGGACT

GATCGGGATCAAACAGTCAAAGCATTGGTTTTATTTATTACTGTTGGAACAAGAGGAGAAGTTACACGATTGAGATCACG

GCTGAGAGCTGTTACAAGAATGCTTGTTATGGTTGTTACTGGATATCTACTTGCGAATATTTTAGATATTATTATTGCGT

TTTGGGAGACAATCAATATTCAAAGTCTTCAAGAACTTCCTTCATTATATACAGTACTTTCGGACATTTCATCATTTTTA

CCAATTGCTGCATGTGCTCTTCGTCTCCCTATCTACACAATTAATGATCGTCAAATTAGAGTCGAGGTTCGAAGAAAGTT

TTGTTATCTCATAACGAGATGTTGCACATGTTGTCCAGGAGTTCTTTGCGATACTCATCAAAGAAAGAAACTTCATGATA

ATCAGTTGGAATCGGAAAGACTTTGGCCCAAGAATGAGCCAACTCCAAAACTTGAGCATGAAATGACACAACTATGGACA

AAGTCTACTGAAAGACTTGGCTGAAAAAAGATCAGAAATTACGGCCTTCAATCCCTAATAATGGCTCGAGCTTCTCTCTC

CACAAAGGAAATCTACGACATTCACCGGTTCCGAGAAACTTATGATGTTTGA

>pR138

ATGGATAGTAACAATATCAGCTTCGTAGAAGAAGTATTCGAGGATATAGAAGATGAGTGTGCCTACTCTCCACCAATATT

TCTCGAGTGGAAAATGGTTTTTGTGGGTGTGATTGGATTAATAGTTGCACTCATCAGTATTGTTCACAATTCACTTCTCT

TCTACACATTTAGTACATCTACAGTACTCCGGAAACGAAACCTGACATATTTGATGTGGATTTCCGGATGCGATGTTTTC

ATCTCGATTTGTTATATTGCAATTATGTGTGTTCAGGTGTACACCGACTATTTCGTATCATTCAAGTTATTAGTTTTATG

GCATGACTATTTACGGACTGCTTTCACAGTGTCTCATATCACCTTATCGGCTGCTTCATTTCTTCTTATGGCCGCAACAG

TTGAAAGATATCTTCAAAGTACTGCCGATAACAGGACTCTGAAGTTATTTCAAATTCTCGCAACCCACAGGACATATGTT

GTTTTTTGTTGTTTCCTAGCGAGTCTCATATTTCGAGGAACCGTGTTTTTTGAGATTGAGGTTGTTACTCAACCGAATTG

TACGGGATTTTCATCAATGGGTTTGGCAGTTGTACCAATGTTTGGAGAAAGTATGGATGTTGTTTGGAGATTTTTTATTC

GAAAAATTGTTACCGTTTTCTTGCCATTTGCAGTTTTGGCATACTTCAATGCAGCAATTGTAATGAATGTGCGAAGGACT

GATCGGGATCAAACAGTCAAAGCATTGGTTTTATTTATTACTGTTGGAACAAGAGGAGAAGTTACACGATTGAGATCACG

GCTGAGAGCTGTTACAAGAATGCTTGTTATGGTTGTTACTGGATATCTACTTGCGAATATTTTAGATATTATTATTGCGT

TTTGGGAGACAATCAATATTCAAAGTCTTCAAGAACTTCCTTCATTATATACAGTACTTTCGGACATTTCATCATTTTTA

CCAATTGCTGCATGTGCTCTTCGTCTCCCTATCTACACAATTAATGATCGTCAAATTAGAGTCGAGGTTCGAAGAAAGTT

TTGTTATCTCATAACGAGATGTTGCACATGTTGTCCAGGAGTTCTTTGCGATACTCATCAAAGAAAGAAACTTCATGATA

ATCAGTTGGAATCGGAAAGACTTTGGCCCAAGAATGAGCCAACTCCAAAACTTGAGATCAGAAATTACGGCCTTCAATCC

CTAATAATGGCTCGAGCTTCTCTCTCCACAAAGGAAATCTACGACATTCACCGGTTCCGAGAAACTTATGATGTTTGA

>pR139

ATGACGACGGACGCTAATTTCACTGGCGACCCGTCGCCGCTCGTTTCCGTACTTGGTTTGATCAAGTCACCTGATTTGAT

GATTGGCTCAACAGAGGTGCCGTTGTTTGAGGAGGAGGATATGTGTACACCACTCAACATTTCTTGTGCATGTCATACGG

ACTACTCAAGTTATGAAGGTTTGGAGAGAGTATTTCTTGGCCTCGTCGCAGTTCCAGTAATCATGTTCGGCATTCTGGCA

AACGTCACATCAATGCGCATCTTTACACACCGTATCATGTCGGGATCCTCAATCAACTGGTATCTTGCCGTACTCTCCGC

GTCTGACACGTTGATTCTCTTCTCCGCGTTTTTTGTGCTCTCGTTGCCGCGTTTCGGAGAATATCTCACCTGGTGGCGGG

CGAATTATATCAGCTATTCGGTGACTCCGTGCATGTACGGTCTAATGATGACCGCCCAGACGTGCAGTGTCTTCATGACG

GTCGGTGTCTCCGTTCATCGCTACATCGGCGTCTGTCATCCGTACAAATCGGTCGAGTGGCTGCCCAAGAAACGGGTCAC

CACTTTCATCATTGGTCTACTCACGTTCTCCGTGATTTTCAACACGACTCGGTTCTTTGAGGTCCACATTTCCAACGTGT

GCTACCGAGAAAACATCGACTACTACATGCCATCACTTCAGCCGACCGCTCTTCGAATGAACGATCTGTATAGGCAAATA

TTCTTCGGATGGGCGTACACCATTGTGATGTATGTGGTGCCGTTCACACTGCTGATCGCGCTCAACTCGATGGTGCTGTC

GGCGGTGAGACGGTCGAGGCGGATGCATATGGTTTCGCAGTGTGGCGCCGAAAGTGAGGAGTTCTCGAAGAAAGCCGAAC

GAAAGGAGCGACAAACTTCTATTATGCTCATCTCAATCGTGCTCCTGTTCATCGCCTGTAACACTCTTGCATTCGTTTGC

AATATTATGGAGAACATGGGCACTGAGGGAGCAATCTATCAAACTCTTGTCACTTTTAACAATTTATTGGTGATCATCAA

TGCTTCATGCAACATCTGTGTCTACATGCTGTTCTCTGGCAAATACCGAATGCTTCTCCGCTACTATCTGTTCTGTGATT

GGTCCCGACAGGGAGAGCTACTCGTTAGTTCCGTACTCAATTAG

>pR140

ATGACGACGGACGCTAATTTCACTGGCGACCCGTCGCCGCTCGTTTCCGTACTTGGTTTGATCAAGTCACCTGATTTGAT

GATTGGCTCAACAGAGGTGCCGTTGTTTGAGGAGGAGGATATGTGTACACCACTCAACATTTCTTGTGCATGTCATACGG

ACTACTCAAGTTATGAAGGTTTGGAGAGAGTATTTCTTGGCCTCGTCGCAGTTCCAGTAATCATGTTCGGCATTCTGGCA

AACGTCACATCAATGCGCATCTTTACACACCGTATCATGTCGGGATCCTCAATCAACTGGTATCTTGCCGTACTCTCCGC

GTCTGACACGTTGATTCTCTTCTCCGCGTTTTTTGTGCTCTCGTTGCCGCGTTTCGGAGAATATCTCACCTGGTGGCGGG

CGAATTATATCAGCTATTCGGTGACTCCGTGCATGTACGGTCTAATGATGACCGCCCAGACGTGCAGTGTCTTCATGACG

GTCGGTGTCTCCGTTCATCGCTACATCGGCGTCTGTCATCCGTACAAATCGGTCGAGTGGCTGCCCAAGAAACGGGTCAC

CACTTTCATCATTGGTCTACTCACGTTCTCCGTGATTTTCAACACGACTCGGTTCTTTGAGGTCCACATTTCCAACGTGT

GCTACCGAGAAAACATCGACTACTACATGCCATCACTTCAGCCGACCGCTCTTCGAATGAACGATCTGTATAGGCAAATA

TTCTTCGGATGGGCGTACACCATTGTGATGTATGTGGTGCCGTTCACACTGCTGATCGCGCTCAACTCGATGGTGCTGTC

GGCGGTGAGACGGTCGAGGCGGATGCATATGGTTTCGCAGTGTGGCGCCGAAAGTGAGGAGTTCTCGAAGAAAGCCGAAC

GAAAGGAGCGACAAACTTCTATTATGCTCATCTCAATCGTGCTCCTGTTCATCGCCTGTAACACTCTTGCATTCGTTTGC

AATATTATGGAGAACATGGGCACTGAGGGAGCAATCTATCAAACTCTTGTCACTTTTAACAATTTATTGGTGATCATCAA

TGCTTCATGCAACATCTGTGTCTACATGCTGTTCTCTGGCAAATACCGAATGCTTCTCCGCTACTATCTGTTCTGTGATT

GGTCCCGACAGGGAGAGCTACTCATTGATCGTCTCGAGAAGAGTCAAACTTGA

>pR141

ATGGATTTACCCCTCAACTCCACAATACCGTCCACTTCGGGAAATGAGACAGAAATTGAAGATGACTGTATCTATGAAGT

CATGCAACCCCATATGCTTGTCATGAGATTCTGGATGGTTTCGATATTCGGCTCGACTATTTCATTTATTTCAATGATTG

AGAATATTTTCCTGTTTTGGTTATTTGTCACAAGCCGCCGCCACCGCCGTGAAAACCTATTCATGATGCTACTTGCCTTT

TTTGATATTTTCGTCTCAATGGCCTATATAATGCTAATGTCGGTTAATGTATTATCCGATGTTCTAATGTCACCAATATT

GGTCAATATTTGGTACACCTATATGATCCCCATATTGACAGTATCCCATATTGCGATGACGTCATCCTCCTTTCTTATTG

TTGCGGCCTCGTTTGAACGTTACTGTGCCACGCTAAACTCTCCGTATCTTAGATTTGCACAGAAAAATCGATCATTCATT

GCGCTATGTGCAGTATTTCTTGGAGTTGTGAGCAAAGGGACCATTTCCATTGAGTTTGAGCTCGTGCACCATGAGCAATG

TGAGGGTCAAATGATGGCAACCACCATGAAATTCCGCCCGTTCGTACTGGAAACTCAATACCATTACCTTTTCCGTTTCT

GGTACCGGAACTTTGTGACAGTCTTTGCCCCATTCTTTATCCTTCTCTACCTGAACGTCCGAATTGTGAAAGCACTGACG

ACCTACACAACTGCCACAGTATGCCTTGTCACTAATGGGAATACTGAGGATTTGCAGAAGAGAAAGGCAAGTGCCCGTGC

TGCTACTAGGACACTGGTGATGGTTGGATGTTGTTATCTTGTGTCGAATATTATCAATGTGACACTGACATCGCTGGAGC

AGACTAATAATGAGCTTGTCGTGGACTATCCAGACCTATACATCGTACTAATCGATTTGGTATCCCTGCTCACAACAATG

GCTTGCGCCGCCCGCCTTCCAATTTACCTGTACTGTCAGCCAGTTCTTCGCCGTGAAATTCTGTGTCGTTTCAAACGACT

CTGTGTTCCATGTAACCCGAGAGCAGACGAAAAATACGTGGAACGTCTGAAATCCGTGGAAACCGACACGAGCTTTTTGA

GTAGAGGAGATGCAAATCATTCTGAAACAATGCCGATTGCCACAATTTCGGAGCCTCGGAATAAGCACAAATGCTCCTTA

AATTCTCCTTACGAGACTCTTTTGTGA

>pR142

ATGTGCACACAGGATCTACCACTTTTCGATTTGGAAAATAATAGTACAATTTATTTTTTCGATCAGTTGATCGCTTTCAA

TCAGGTCTACACAGTATTGCATCGGTATTTATGTCTTTTTGTGTGTTTTTTCGGTGTTCTCCTCAACAGCCTTCATTTTT

ATGTGTTAACAAGAAAAGCCATGCGAGTGTATATAATAAACGCCTTACTATGTGCAATGTCAATTTGTGACATAATCACC

ATGACGTCATACTTCATATATATCCTTCGGTTCCGAATATTCGACACCCCATCCACCATAATCGGATACTCATATCCATG

GTTAATATTTCTTATCACCCATGTGACGTCATCCATTGCACTCCATACTACATCACTATACCTATCCGTAATCATGGCAT

ACATTCGATGGACGGCACTTGATCGACTTGATGCGAAATGGATCAATCATGGTGCATTGAAACAAATTTTAATATTCACT

GCGCTAATCGTGTCAGTGATATCAATACCAACAGTGATGGTACACAAAATTGTACCGGTCACAGAGGTTTTGGGGATGAA

TGAGACTGATTTGTTGGAGGGTGCAAAGTTCGACGGCCTCTACACAGTTCAACTGGACGAGACCAAGATCAACGGATGTG

CACTTTTTCGTGTAAATTTATGGATTACCGGGATCATGTTCAAGGCTCTTCCTTGCCTTCTGATCCTCTGGTTCACCATT

GCCTTAATCTATAAACTGGTTCAAATGAGCGAGAAGCGGAAGATTCTACGTGGTGAAAAGCGGGAAAACGTGGAATTTCA

GCTGCTTGCCACGTCAGCAACACCCACGACGACTACAAGAAATGAAGCTTCACCACGATTGCGAAAAGTGTCCCAATGCA

GCAGAAATGTCTCAATTGACCGAACGACATTGATGCTCATCATAATGCTTGTGGTATTCCTGTGCACCGAAATGCCGCAA

GGCCTACTTTCCATTTTATCGGCAATCTACCCGACACATGTGCATACAATGATCTATGTGAATGTTGGAGAGGTTCTCGA

TTTAATGTCACTAATCAATTGTCTGACGTCTTTCATTGTATATTGCGTGATGAGTACCACGTACAGAGCTACGGTAAAGT

CTGTTCTGTGCCGATCGACTTCGCGACGGACTGTCTTCAAAAATACGCATCTCTTGCTCGATTCTTATAGAATACAGGTT

TCATAG

>pR143

ATGAGCAGCAATCTAACCGACACGACGACAACGGTATTCGAGACGTTTCTCCTCGAATATGGGAGGATAATTCACCCGCC

GATGGTCCTACTTTTATGTATTTCCGGGGCTCTCGGGCATATGCTCACAATCACAACACTCTCATCAATGCTGAACCCAA

CGAATATGTTCTTGATTTCTATGAGCTGCAGTCAACTGGCCTTATGCATCAATTTTCTCTACTCAACATTCTTCAAATTC

ATGTCAGATCAGCTGTGTCAGCCGTTCTTCTTCTCCTATTACCTAGCCACTACTATGCATTTTTCCGTCACAGTTTCGGT

ACTCGTGCATATGTCAGCGGTATTCCACGTAGTTGCTCTTTCGTTAATACGATTCTTTTCGCTTGCACAGCTCTCTAGCG

TCAATTCGAATGTGCAATGGTTTACGTGGCAGAAATCCCGCGTGGCAATCATAATGATCTACGTGAGCGTGATCTTCCTG

TGCATCCCACTACACTTCACATCACAAGTGACCGAGGTCTCCGAAAACGAGGGATGTGCTGAGCGATATCCTGCTCTCAG

GAATAAAGTGGCCTACCAGTTGAGCTACACGCAAAGCATTTGGCTCAGGAATTTGAATTTTTGGCTCTTCTACCTGGTTG

CCAAGGTGGTTCCATCGATAATCCTGTGCATAATGACCTGCCTAATTTTGGACCAGCTCAAGAAGATTCAAGTTTTAAGC

GCGCGCTTTTCGTCAGTGGAGCGCGATAAGCAACACTCGAGAACGACAAATATGATACTTGCGATAATGATTCTCTTCAT

TATTGTCGAGCTTCCACAGGGTGTGCTCGCCGTGCTCTCCACCGTCACCAGTGTTAAGCTGATCTATGAGCTCGGTGATT

TGACGGAGCTCTTCACATTGCTCACTTCTATCATTATTTTCACACTCTTGTGCTCGATGAATGGAAAGATTCGATCAGCA

TTCAAAGAGTTGTCGTGTGTTCGATCGATTGGCAGGATCTTTGCGACAATGTGTCCGCCGCCGAGCCCGGTCGCTGCGAG

CATCTCTGGACACGATGCTCTTCTGGAGCACAACACGATTATCATCACCAATTGTCATATTGACAAATCGGAGAATTATG

CGGTTCTTTGA

>pR144

ATGTCCGAAGACGGAGATGATCCGTCATGCTACATGGTGCAGGAGACGACGTTCGATTACGAGTATGACGAAACGTTCGA

CTGGAGTAACATGACTGCGATGGACTGCAACTGCACACATCCGTTGCACAAGTATCTTGAGGTTTGCATTTCTCGCTGCA

CTGTACCCGATGACACTGTTTTCTTCTCAATGACCGACGAGGAGTTGTTCGAGATCGCTCTTCCCGGCTTCTTATATCTC

ACTGTTTTTCTAGTTGGAACTATTGGAAACTCTATGGTAATATTTGTTGTAAACCGATTCAAACGAATGCGCAACGTAAC

GAACATCTTTCTCGCCTCGCTCAGCACCGCCGACTTGTGCCTTATCTGGTTTTGTGTGCCTATAATGTTCATGAAGTATA

TGTCGCATACGTGGTCGATGGGCAGGTTTGCGTGCTATTCTGTGCACTACATTCAACAGTTTACATGCTTCTGCTCGGTG

CTCACGATGACTATGATTAGTTTTGAGAGGTTTCTTGCAATAGCATATCCTATGCGAAATATCTGGTTTTCATCCATAGG

ACGAGCTAAAAAAGTAATCCTACTCATCTGGATGTCATCCGCGGTACTGGCCGTCCCAACCGCTGTTCGGATGGATTATG

AGACCAATCTTTCATTGTCCGGGCAGAGGGTGCATTGGTGTCGAAGACGGTTTCCCGCGCAATTTTTAGGATATCCTAGG

ACATCGTTGAATAAAGCATATGCCATGTATCAGTTGTTGCTACTTATCATATTCCCGGTGCTCACAATGTCCATATGTTA

TGCTCGTGTATCTGCAATAGTGTACAAGTCGTCGAAAGATCGTGTAATACTTTCCCAAGCTATGGTCGCATTTTCAAAGG

CCGCCACTGACGCCGTTACCTTCTCCGGCTACTCTGCAATCCCCATGATTACAACTTCCCGGAACTTGAAAACCGCCAAC

ACGACCATCAAGTCCTACAGTAATCACAGAAATAATCGCGTGGCAGAGGCGAATAAGAAACAGATAGTTCAAATGTTAAT

CTCCATCGTCTGCATGTACACGGTTTGCTGGCTCCCAACGATCGTCGACGAGCTTCTGACGTCCTTCGGGTACATTTGTC

GAACATCAAATACGCAGACATTGAAGCATATGCGGATGGGATTTAATGCGCTGACCTATTGCCAATCATGCATTAATCCT

ATCCTGTATGCCTTCATTTCTCAGAATTTCCGGTCCACGTTCAAGACCGCCTATTCGAGAATGAAAAGTCGTCTTCAGGG

CGTGGAAGAAATGCGCTCTCGTATGGGATCGTGCTCTTCGGCGAGCATGATGTCGACCCGCAACCAGCACCGAATCTACG

GGTCCAGCTTCAACACACTCACAGTGCCCGGTCGCTCCATGATCACTCCGAACATGTCACGCGACGTCTCACAGTTGTCA

TTGTGCAGGCCGACGTCGCAAATGTCGTTCTGCCGACCGAAGAGTCCAATGGCCGGAGACTCTCCGTCGACTTTCATGAG

ACATCGTCCTAGATCGCCAACCGACGTGTCCCATGATTCCGGAAGACCACGTAGCCCCACGGATCTTTCGCAGTCGACAA

AGCCTTCGCGACGATCCTCCTCTATAAGACCCCGTAGCCCAACATCGACATCTCAAATGTCTACCATAGTCCGATCGAGA

AGTCCTACAGGCGCATCGGACACCTCTTCATTGTTCCCATCAAGAACAAGAAGTCCAACTCTCCAATCAAACACATCTGG

TCAATCCATGGAACGAACCACGGACCGACTGAGTGTCCGTGACGCTGTCCGACCAAAAACACCGCCAGCAGTGGTTGTCT

AG

>pR145

ATGGAATGCAACGACTCGAGATTATTCGATACAAACAACAATTTGACATATTTTGTGATAGAGCAATTTTATCGATTTAG

ATACCTATACTCTAACGTTCATCCTTTCTTGTCATTCATCCTATGTGTTCTTGGATTAGCTGCCAATGCCATACACATCA

CTGTCCTCACAAGACCACGAATGCGCCACTCATCTGTTCATACCGTTTTAGTGTGCATTGCAATTTCTGATATGGGAACA

ATGACCTCCTATTTGATGTACATAAGTCGATTTGAATTTCTATCGGACAAAGAAGGATACTCATATTTTTGGGCATTGTT

TCTAAAGTGTCATGCAATGTTATCTATTGCACTACACGCTATCACTCTGTATTTGGTAGTTTTGATGGCATTCATTAGAT

TATCAGCAATGAAACTGACTACATCTAGATGGTTGGATCATACGAGAGCTTTAACAAGTGCAATATTTATAGCTCTCTTC

GTTTTTATAATGTGTGTACCTACGTTATTGGCACACCAAATTGATGAAACCACTAGAGGTGTGACTATGAATGGAATGTA

TTATAAATATAGCGTCGGATTTTCTACATTGATGATGCAGAATGGATGTTCCTTAATGAAAGGCAACCTGTGGCTGACTG

GAATTTTTTTGAAGGCTATACCATGTCTTCTCCTGCTAACATTCACCATTGCTCTAATCAACCGTCTTCGTGAAAACAAC

GAAAAACGAAAGATATTGATAAAAGAGGAACGCGCCAAAAAGCGCGGAGACTTCACCACCTACATGCTTCTTCTCATGGT

TACTGTCTTTTTGTTCACAGAATTACCACAGGGGATCATGGCAATTTTGAACGCTCTGTTCACTACTCAATTCCACCAGA

TGGTCTACTTGAACCTGGCCGACGTTTTAGACCTCCTGTCGTTAATCAATTGCTACGTGGCATTTTTGGTGTACAGTTTC

ACATCATCCAGATACCGACAGACGCTGTTCAGCTTGTTACCATTGACAAAAATTTCCTATTCCGGAATCTCCACCCGTCA

AGGAACCTTGAAATCTCATCAACACCCTGGAGCAAAAACGTTGGTGCAGCGAGCCAACTCAGTTGAGGTGGCTTCTGCTC

GATCACCGCTGGTCGACAAAGCAGCAATAACACGGCCGAATTCCGCACAGCCCACAGATTTTTAA

>pR146

ATGGTTAGTTCGGCGGCCACCATTTCGACCATTTCAACCACAACGACTCCCTCCACCATCAGCAACGTTATCACAAGTCA

TTCGAACAATGGCTCGTGCATTCAGATCGCTGAGGCGATTGCGGCACAAGGCATCGATGATATTACTGTAGACTTTTACA

TCCGATCAATCTTCACATTCCTCTACGGGTTCCTGTTTGTATTAGGCATTTTTGGAAACGGCGGCGTACTATGGGCGGTG

GCGAGAAACAAGCGGCTCCAATCGGCTCGCAACGTATTTCTGCTCAACTTGATCTTCACCGATTTGATATTGGTGTTCAC

AGCGATTCCAGTCACACCATGGTACGCGATGACCAAAGACTGGGCATTCGGGTCAGTGATGTGCCATTTAGTTCCTTTGT

CAAATTCGTGTTCGGTGTTTGTGACGAGTTGGAGCCTCACTGCAATCTCCTTAGATAAATTTCTGCATATCAACGATCCC

ACCAAACAACCAGTTTCTATTCGTCAAGCGTTGGCAATAACATTTCTTATCTGGATAGTCTCAACACTGATAAATCTACC

GTATCTTATGTCTTTCGAGCACGTCGATGGAAGCTTTTACGTTCAGCCCGGAGAAACTCCATACTGCGGGCACTTTTGCG

ACGAGGCGAATTGGCAGAGCGAAAATAGTCGAAAGATTTACGGAACTACGGTTATGTTGTTACAGTTCGTCGTGCCGATG

GCAGTGATCACGTATTGCTACTTCAAAATCTTGCAAAAAGTGTCAAAAGACATGATCATCCAAAATGCTCAATTCTGTCA

ATCACTGACACAAAAGCAGAGAAGTGATGCGACGTCACGAAAGAAGAAAGTGAATTATATTCTAATTGCAATGGTTGTCA

CATTTATCGGGTGTTGGTTGCCTTTAACATTACTCAATTTGGTCAAAGATTTTAAAAAAGAGCCCGAATGGCTAAAACGT

CAGCCGTTCTTCTGGGCAATAAATGCTCACGTCATAGCCATGTCCTTAGTCGTCTGGAACCCTCTGCTATTCTTTTGGCT

GACACGAAAACAAAAACGTTCCGGACTGTCAAAAATACTCAACTCAACAGAGGGTTCGAAAAAAGCAGGTGGTTCTGGAT

TGCGAGGGATCCAGCTACACGACCTCCTCCCGACCTCTACTCATTCGGACAGATGTGCAGGCAACTCTTTCTAA

>pR147

ATGGTTAGTTCGGCGGCCACCATTTCGACCATTTCAACCACAACGACTCCCTCCACCATCAGCAACGTTATCACAAGTCA

TTCGAACAATGGCTCGTGCATTCAGATCGCTGAGGCGATTGCGGCACAAGGCATCGATGATATTACTGTAGACTTTTACA

TCCGATCAATCTTCACATTCCTCTACGGGTTCCTGTTTGTATTAGGCATTTTTGGAAACGGCGGCGTACTATGGGCGGTG

GCGAGAAACAAGCGGCTCCAATCGGCTCGCAACGTATTTCTGCTCAACTTGATCTTCACCGATTTGATATTGGTGTTCAC

AGCGATTCCAGTCACACCATGGTACGCGATGACCAAAGACTGGGCATTCGGGTCAGTGATGTGCCATTTAGTTCCTTTGT

CAAATTCGTGTTCGGTGTTTGTGACGAGTTGGAGCCTCACTGCAATCTCCTTAGATAAATTTCTGCATATCAACGATCCC

ACCAAACAACCAGTTTCTATTCGTCAAGCGTTGGCAATAACATTTCTTATCTGGATAGTCTCAACACTGATAAATCTACC

GTATCTTATGTCTTTCGAGCACGTCGATGGAAGCTTTTACGTTCAGCCCGGAGAAACTCCATACTGCGGGCACTTTTGCG

ACGAGGCGAATTGGCAGAGCGAAAATAGTCGAAAGATTTACGGAACTACGGTTATGTTGTTACAGTTCGTCGTGCCGATG

GCAGTGATCACGTATTGCTACTTCAAAATCTTGCAAAAAGTGTCAAAAGACATGATCATCCAAAATGCTCAATTCTGTCA

ATCACTGACACAAAAGCAGAGAAGTGATGCGACGTCACGAAAGAAGAAAGTGAATTATATTCTAATTGCAATGGTTGTCA

CATTTATCGGGTGTTGGTTGCCTTTAACATTACTCAATTTGGTCAAAGATTTTAAAAAAGAGCCCGAATGGCTAAAACGT

CAGCCGTTCTTCTGGGCAATAAATGCTCACGTCATAGCCATGTCCTTAGTCGTCTGGAACCCTCTGCTATTCTTTTGGCT

GACACGAAAACAAAAACGTTCCGGACTGTCAAAAATACTCAACTCAACAGAGGTGAGTATTGTAAACTTCTTCAAGTCCC

CTTGTATAAAGTTTGCTCCTAGCCGTGTGCTCCTGGTGTTCTTCCCGTTTTCCACTGCTTCCTTTGTGTCCCCCCTCTCT

TCCTCTGAGCCGCCTTTTGTTTTCAGATTGTGTCCTCGTTTGCCAGTAGAGTGA

>pR148

ATGGTTAGTTCGGCGGCCACCATTTCGACCATTTCAACCACAACGACTCCCTCCACCATCAGCAACGTTATCACAAGTCA

TTCGAACAATGGCTCGTGCATTCAGATCGCTGAGGCGATTGCGGCACAAGGCATCGATGATATTACTGTAGACTTTTACA

TCCGATCAATCTTCACATTCCTCTACGGGTTCCTGTTTGTATTAGGCATTTTTGGAAACGGCGGCGTACTATGGGCGGTG

GCGAGAAACAAGCGGCTCCAATCGGCTCGCAACGTATTTCTGCTCAACTTGATCTTCACCGATTTGATATTGGTGTTCAC

AGCGATTCCAGTCACACCATGGTACGCGATGACCAAAGACTGGGCATTCGGGTCAGTGATGTGCCATTTAGTTCCTTTGT

CAAATTCGTGTTCGGTGTTTGTGACGAGTTGGAGCCTCACTGCAATCTCCTTAGATAAATTTCTGCATATCAACGATCCC

ACCAAACAACCAGTTTCTATTCGTCAAGCGTTGGCAATAACATTTCTTATCTGGATAGTCTCAACACTGATAAATCTACC

GTATCTTATGTCTTTCGAGCACGTCGATGGAAGCTTTTACGTTCAGCCCGGAGAAACTCCATACTGCGGGCACTTTTGCG

ACGAGGCGAATTGGCAGAGCGAAAATAGTCGAAAGATTTACGGAACTACGGTTATGTTGTTACAGTTCGTCGTGCCGATG

GCAGTGATCACGTATTGCTACTTCAAAATCTTGCAAAAAGTGTCAAAAGACATGATCATCCAAAATGCTCAATTCTGTCA

ATCACTGACACAAAAGCAGAGAAGTGATGCGACGTCACGAAAGAAGAAAGTGAATTATATTCTAATTGCAATGGTTGTCA

CATTTATCGGGTGTTGGTTGCCTTTAACATTACTCAATTTGGTCAAAGATTTTAAAAAAGAGCCCGAATGGCTAAAACGT

CAGCCGTTCTTCTGGGCAATAAATGCTCACGTCATAGCCATGTCCTTAGTCGTCTGGAACCCTCTGCTATTCTTTTGGCT

GACACGAAAACAAAAACGTTCCGGACTGTCAAAAATACTCAACTCAACAGAGATTGTGTCCTCGTTTGCCAGTAGAGTGA

GTAACTCGATTCGGCGGTCAACGTTTCGGAGAAACAATATTGACAGGGTTCGAAAAAAGCAGGTGGTTCTGGATTGCGAG

GGATCCAGCTACACGACCTCCTCCCGACCTCTACTCATTCGGACAGATGTGCAGGCAACTCTTTCTAATGGCTCGACGAG

TACTACCCGCGAGATGCTGTAA

>pR149

ATGGACGAAGGAGGGGGTATTGGAAGCAGTTTGCTCTCCAGAATCACGACGACAGCCTCTGAAATTATGATGCGAAACGA

ACCGACGACAACTGAAAATCCAGCTGTTCAAGAAATGAATCATATTTATCATTTGACACCAAGTATGAAGATGTTATGTA

TTCTTTTCTATAGTATACTTTGCGTATGCTGTGTCTATGGAAATGTGCTCGTTATTCTGGTTATTGTTTATTTTAAACGA

CTTCGAACGGCGACTAATATTTTGATATTGAACTTGGCAGTTGCTGATCTTCTCATATCAGTATTCTGCATTCCGTTCAG

CTATTGGCAAGTATTGATTTATGATGATCAACGTTGGCTCTTCGGCTCAATGATGTGCTCTTTATTAGCATTCCTTCAAG

CAATGGCTGTATTTTTATCAGCTTGGACACTTGTCGTTATCAGTTTTGATCGGTGGATGGCTATAATGTTCCTTTTAACT

CCAAATATTCGAATTACAAGACGAAGAGCTCTTTATCTGGTAGCTGCCACGTGGATATTCAGTATCCTAATGGCGTTGCC

GTTACTCTTCACAACGAGATTTTTCGAAGACCAAGACGGTTTACCGAATTGTGGAGAAAATTGGACGTATTTTGGAGATT

CTGGAGAACAAGTGAGAAAAGTGTATTCTTCAATGGTCTTAATTCTACAATATGTTGTACCTCAAGCAGTTTTAATAATA

ACTTACACACATATTGGAATTAAAATGTGGAATAGTCGAGTACCAGGAATGCAGAATGGAGCAACAAAGAAAATGATCGT

TGATCGACATGAAAGTGTCAAAAAGCTGGTCCCAATGGTGATTCTCATTTCGGCACTCTTCGCACTTTGTTGGCTTCCTT

TACTTATACTGATCAACGTCATTCCAGAATTCTATCCAGATATCAACAGTTGGGGATATATTCTGTATTTGTGGTGGTTT

GCTCATGGACTTGCCATGTCTCATTCAATGGTCAACCCAATTATCTATTTCATTCGAAATGCCCGTTTCCGTGAAGGATT

CTGTTTCTTCTCTTCAAAACTTCTTCCATGTATATCATTTAAAGAACTTCGTCTTTTAACTGATAATACTAGCAGAAGAC

ATCGCTTACGAGATATTCACGAAGTGGAGTCCTTGACAGGCAAACATGTCGTTCGGCACGTTTCTTCGAAGCCCGACCAC

TCGTCGTCGTCCGAAACAACTCTGCCAATTCTCTCGCGTAGCTTTTCCCGTATCATTAAGAAAATTGATCTACCATGTAC

TTGA

>pR150

ATGAATTCCACAAAGAATCTTGTTCCAACTCCAATGATATATTCAACTATGGCATTATCAATTATAGGATTAATCGGGAA

TATGACTATTGTACTTGCCACTGTTATTAGTAAACGTCTCCAATCGAGATGTAATATATTAATTGGACTTTTGGCACTTT

CTGATGTTGTAGTTTGTACTTATTTGATTCATTTACGAGTTTTGATGTTCTTGGACATGTACATGTTGACTTCAACCCAG

TGTTTCCTACTTTCTGCCTATGGTTTATTTGCTCTAAATATGCAAAGTTCATTGTGCCTTATTATAGGATTAGATAGACT

TTATAATGTCACCTACCCTTTGCGATATTCCCAACTTCCAAACTCATTTTACATTGCCCTTTTCCTTTTTTGCGTGGGAT

TTTCCGTTCTTATCACTTATTCCGGATATTCTTATGCTTCGGATTCAGAAATAGTTCTAGTATGCTTACCACCAACAGCT

TATACCGATAAATCTAGAATTATATGGATTGGTTCGAATTTTATTATTGCAATTTTAGTCATTTTTGTGTATGGTGGTGC

ACAAATTCGTTGTCGGATTTTGAGAGCCCGTCACGTGCACGAACAATCTATGGATGCAGTGAATCGACTGCTTAAATCGT

TGAGCATTGTTGTAACAATATATGTCTGCACGTGGTTTCTCACGATCAGTTCACTTGTTGTTTCTCAGTTGATTACCTTT

TCACCGCCTGTCACCGCAGAAATTAATCGACAACTTGGCTGGCTGGTGATTATCAATGCTTCACTCAATTTCTTTGTTTA

TCTTTGGAGAAATTCAGAATATCGAAAAGCATTCATCCGACTTTATGGACTTTCTCGACTCTTCAGCGTTGAAAATACCT

CGATTGATGCTGCAACACGAGTAATGAAGTCTTCTCATGTTCGAGTTGCATAA

>pR151

ATGACCGATTTGGAGATTTGCGATTATGACGAGACCACCAACAATAGGCAAAACTGTACCGAAGACTACGGCGTATACGA

TGGATATACACAGACATGCGTTCCTCTACAACTTATACAATTTGAAGATGTGGAAAATGATGTGAAATTGCAAATAATCT

ATGGAATTATATTGCCAGTTCTTGCGGTTCTTGTTGTTGTTTCCAATAGTGTTGTTATATTGGTCTTGAGCCAGCAAAAA

TCGAAAAGAGCCAGCGTGGAGCCTTTACTATGGATGGCAATATGCTCATTATTAATGGCAATTTCACCACTACCATTTAC

TATCTATTACTACAATTTAAGCCATCATTTGGACTTCAATCAGACACTTTTCCTATGCTATTTGCAAAAAGTTTGCATGG

AAATTATGCCGTTTTTCTTCAATAACTTAGTTCTGTTGTTCACAATTTTATTAGGGGTTCAAAGATTCATTGTTGTTCAG

TACCCTCTTCAATCCATCAGATGGTGCACACCAAAAATGGTTCGAAGATATTCAAAATTAATATTAATACTGGCCACATT

TCTAACCGGTTTTCATTCTCTTTATGATATCCGTTTATTGTACCATTTTTGTATAAAATATCGAGATGAATCATTCTGGG

TGGCGAGATGTTTTATTGGGTACTCTTCGTTAACTTCTGCTTTTGGCGCAGACGCTTTTTCGGCGCTCTTCGACTATTTT

CGAATTACCATCAACCTACTGGCCAGTGGGTTATTATTCGTTGTGACCATTTTGCTGATCCAGACAATAAGAACACATGA

TCATCCAAAACAGGGAGTTCACCGTCACAAAAATCGGAAAACTTCAGCGAACACGACTATAATGTTGACAGTAATCATTA

TTATTTATATGCTGGCGAGAGTACCTACCACACTGCTATTCCTACTTGTAAAACTGATGGACTACATTTCAGTGCCAACA

ATAGCATTTGAAGTGATGAATAATATCTATCTACGGGTATTTGCAAATATCACCACAATATCTCTACATCCCATATCATT

TGCGATTTATATGTTCATGTCGAGGAAATTTCGAGTTTCAATGAGAAGGTTGTTAGGTTGGCGATTTCTTGCATCCGACG

AAGAATTTAATGCATTCACATCATCCACCAGAATCCACACGGCACATCCAAAAATGACTCTAATGGAGAAGGAAAGCATA

GGCAGCAGTGGGAAAGGTACTCCATCAGTCGGACAAAAACGTGGAATGTCCGTCGATTTTAAGATTTCGTTTGACGTTGA

GCCCCAACGTCGTCGCCGAGCAACCACCGCCGACACGAATCATGTAAATTTCCGAAGATTCGCTGAAAAATCGGAAAAAA

CGACGGAAATTTTGAAGATTCGAAGAGAAAATTCGTAG

>pR152

ATGGCAGTGGCTCACCATTTGTGTCAATTTCTGCGGAAAAATGAGACGATTGATTTCTCGACGTTATTGAGTCATTCGCC

ACGGGAAGAAGCCACTTCAAACATAAATAAGATTACATATGCTTACATTGCGCCGTTCATAATCATATTTGGAATAGTGG

GTGATGTGTTAACAGTTGTCACTTTAACACACCCATTATTACGGAAATCATCAATCATATACACATACCTGACACTGCTG

GCTATGACTGATTTGCTCACACAATTCTCTGTTATACCAATGATCATGTGGCTGCTCGACATTCGTGCCTGCTCCAAATC

ATCGGCGTTTTTCTATGCGCACATTGGGTTTCCTTTGGCGAACGCTTTGATGGGCTCAAGTGTGTGGATTGTTGTATTTC

TGACGTTATCACAGTATATGGCGGTGTGTAAACCGTTTGCATATGGTTTAAGGTCACGGAAAATCTGTTATGTATTATTT

GCACTTGCTTATATGTTCAATTTCTGTATCTATGCACCATGGGCGGTAAAGAAGAATGTTCATGACATTTCTGATCTGGT

ACCGGAGAGCTTCCTGGTATCCGTATGCCCATACGTAGTGTGTGACGCAAAGCGACCGGATTGGTTCGTGATTTATGAAA

CTGCTCGCGAGCTCATATCCAGAATATTTCCATTCTTCCTTGTCGCATTTCTCAACATCAAGATTCTTATAACGTATAGA

AACACCAAGCGGGATAGGATGGAGCGGCTGGCAAACTCTCAGAAAAAGTTTATGTTCGAAAAGAGCGAGAAGGAGGAGAA

GCGACTTTTTATATTGCTCTTTGCAATTGTCATTGTGTTCTTTGTGTGCACTATTCCAGCAGCTCCACTCACTATTTTAG

TGGCGGATACGAAAAATAACAATGTCGGATTTCAAATCTTCCGAGCAGTTGTGAACTTACTAGAATTTACCAAGTTCGCA

ATGAATTTCTACTTTTACTGTCTCATCAATCCAGACATCCGCAGTATTTGTGCGCACGTAATTACATGTAAGAAGATCAC

AAAGCCTGCGAGAGTAAAGGGTCAACCAACTACTCCACTTTCGAAATATACAAGATCTACAAAAAATTCCACAATTCGCG

ACGCAAACAAGAATGGTGAAAAGCGAGGAATCGACGGCAACGATAATGATTCAAAGAAATTCGATGACTCTAGAAATAGC

TCGTTGAAATCTGGAAAAGATTCCGTTCGGCGGGGAGTACCTGCGGACAGTTTAATCGAGAAATTAACTGTCATTAAAGA

AAACGAGTCAGTCACTTCCGATACCGGAGATCACGTGTAA

>pR153

ATGGCAGTGGCTCACCATTTGTGTCAATTTCTGCGGAAAAATGAGACGATTGATTTCTCGACGTTATTGAGTCATTCGCC

ACGGGAAGAAGCCACTTCAAACATAAATAAGATTACATATGCTTACATTGCGCCGTTCATAATCATATTTGGAATAGTGG

GTGATGTGTTAACAGTTGTCACTTTAACACACCCATTATTACGGAAATCATCAATCATATACACATACCTGACACTGCTG

GCTATGACTGATTTGCTCACACAATTCTCTGTTATACCAATGATCATGTGGCTGCTCGACATTCGTGCCTGCTCCAAATC

ATCGGCGTTTTTCTATGCGCACATTGGGTTTCCTTTGGCGAACGCTTTGATGGGCTCAAGTGTGTGGATTGTTGTATTTC

TGACGTTATCACAGTATATGGCGGTGTGTAAACCGTTTGCATATGGTTTAAGGTCACGGAAAATCTGTTATGTATTATTT

GCACTTGCTTATATGTTCAATTTCTGTATCTATGCACCATGGGCGGTAAAGAAGAATGTTCATGACATTTCTGATCTGGT

ACCGGAGAGCTTCCTGGTATCCGTATGCCCATACGTAGTGTGTGACGCAAAGCGACCGGATTGGTTCGTGATTTATGAAA

CTGCTCGCGAGCTCATATCCAGAATATTTCCATTCTTCCTTGTCGCATTTCTCAACATCAAGATTCTTATAACGTATAGA

AACACCAAGCGGGATAGGATGGAGCGGCTGGCAAACTCTCAGAAAAAGTTTATGTTCGAAAAGAGCGAGAAGGAGGAGAA

GCGACTTTTTATATTGCTCTTTGCAATTGTCATTGTGTTCTTTGTGTGCACTATTCCAGCAGCTCCACTCACTATTTTAG

TGGCGGATACGAAAAATAACAATGTCGGATTTCAAATCTTCCGAGCAGTTGTGAACTTACTAGAATTTACCAAGTTCGCA

ATGAATTTCTACTTTTACTGTCTCATCAATCCAGACATCCGCAGTATTTGTGCGCACGTAATTACATGTAAGAAGATCAC

AAAGCCTGCGAGAGTAAAGGGTCAACCAACTACTCCACTTTCGAAATATACAAGGTAG

>pR154

ATGAACAACTCAACATCGGATACGTTTGTCATGATGCATTCGACTTCAGAAGTTATCGAAATGTTTTACCAGCTTGCATT

TTTCATAATTGGTACTCCAATCAATATTTTTGCTTTTGTCAAAACAAGTCGAAATGTTCGAGAAGGAGGCGTGGAGTCCA

GACTGGTCAAGCTTAGTCGACAGCTACTTATTGCCCACATAATGGTTCTTTTTATGTACGGGATCTGGAGAAGTTATTGG

ATATATAATATCGTCTGGACTCAAGGAGACATTATGTGTCGAGTATTCAGTTTTCTTTGTGCTCTGCCCTTTCATCTTTG

GTCGAACATGGTAGCAGCTATTGCAATTGACATGCTTTGCTGTATTACATCTCCACTGAGCAGCTATCGAACAGGGGCAA

ACCGAGTTAATTGGCTTATTACTCTTGCGTGGGGCTTTGCAGTTGTATGTGCATCACCAATGAGCATTCTGCGTGGAACC

ATTCAGATCAACGATGAAGATATGTATCAATGCTACCCAAGAACTGATGTATTCAATGATGATGTGTTAATTGCATTCAA

CCTGTTTCATGTCATAACTTCGTTTTACATTCCATTGTTGATTGTTATCATATGCTATTTGGCTATCGGGGTATCGATAC

GGAAACAAATGGCGGAACGTCGTTTATTACAGGACGGAGGACAAAGTGGACAAAAAACGAACAACACGAAAGCAAGATTT

CTCAGAGCTTCTGTGGCAATTATTTGTACGTTCCTGTTCACATGGCTTCCATATCAAGTTTTGGCTCTTCTTCGAATTGT

CTGTGTTTCTGAAAATTGCCAGGAAATTGTGAGCAAACTGAATTGGTTACAGGCAATTATTATAGCAAGCACGTGCATCA

ATCCATTCCTATATCGTTTTGGCACTGAGCCAAAAAGAAACTCGTATTACTGCAGTACTATGGACACTGCTGGTAAGGAG

GAAGTCGGTCTATCCGCCGATCGACGACGGTCCGGGAAGCCGAGGCCCAGCGGATTGTTGCCGCAGCAAATCGCGCACAC

CGAGAACAAAACTCCTCCCAAGTGTCGCCTCGAGGTACACGCGCCACGGTTCGAGAACAGACGCGCGTCGCAAATCGACT

CAAAAACACCAAAAGTAATGAGGTTTAGATGCACCGGATCACTTGACGGAAAAGCTGGCAGATGCTGA

>pR155

ATGGGTGATGCTGAATCTCATCATTGTATAGATGTGAACGCCATTCTTCAGCAGTTCAATGATTGGACAGTCCTCTTTGA

AGTTCGGCTTGGATATTCAGTACTATACTTTCTCATATTAATAATCGGATTGGTTGGAAATGGGCTATTGATCACTTCAA

TTTTAATGCGAAAGAAACTTTCCGTGGCAAACATATTCTTGATAAACCTGGCAGTTTCTGATTTGCTTCTTTGCATCACG

GCGGTGCCGATCACTCCAGTATTGGCGTTTATGAAGCGATGGATATTTGGAATAATTATGTGTAAATTGGTTCCAACTTG

TCAGGCGTTTTCGGTGCTCATTTCTTCATGGTCTTTGTGTTACATCGCAATTGATAGATATCGAAGTATTGTGACGCCAC

TCCGGGAACCATGGTCTGATAGGCATGCAAGGTGGCTTCTGATGTTCACATGGGTGGTCGCCTTCCTTGCTAGTTATCCT

CTATATTACTCACAGAACTTGAAAACAATGGTTATTGAAAATGTGACATTATGTGGAGATTTTTGCGGCGAGTTCAATTG

GCAGTCGGATGAAATATCCAAGTTGACATATACTACGAGTTTATTGATTATTCAGCTGATTATTCCAGCAATTATCATGT

CTTTTTGTTATTTAATGATTCTACAAAAGGTACAAACCGACTGGCTTGTCGACGAGGGATCCATGTTGACTGCCGCACAA

CAGGCTCAAACAGCAGTTCGAAAGCGACGAGTGATGTACGTGTTGATTCTAATGGTTATTGTTTTTATGGCTTGCTGGTT

CCCGTTGTCCGCCGTGAATTTGTTCAGAGATCTCGGAATGCGATTCGAGTTCTGTCAAACTGTTTACAAGGTTTTAATGA

TGGACCAAATGTATTTCAAGTTGCTCAATGTGCACGTCATCGCGATGACTTCGATCGTATGGAATCCGGTGCTCTATTTC

TGGATGAGCAAGCGTCATCGACGAGCCCTGAAAGACGACATGACGTGGCTCACCAATGCTCGCCGTCATACAAACGTCGG

CGTTCTGTCTCGCTTCACACCTTCTCCATCAGTTTCAGTGGTTTACAGACGAACTCTGGAGCGACATCTAGGTGTCAATC

ATTTCCGCCGTGGCACACTTGCGGACCCGACATGCACTTCACGTGAACGAAGTCTTCCGCGAGAACTTCAATCAAATTGT

TTCCTTCTTGTTCCGCTTATGCCATTATGTCAATCTGTGACGAGGAAGAATAGTCATCTAGCAATCAATCGAGACGGGAT

CGTCATTCCACAAGCCAATGGCTCAAGTCGTCGGCCGAGCAGCGTGAATACCAATTCAACTCGAGACTGGTGA

>pR156

ATGGACTTTACGGAAAATGAAGAGGAGTACGAGCATTGGACACATATTGAACGACGAGTCCCGTTTCAAGGATACCGAAC

GTATGTGGCTTCCACCTACATTAGTTTCAACGTCGTTGGCTTTGTGATCAATGCCTGGGTACTTTATGTGGTGGCTCCAT

TGTTATTTGCACCAGCTATCAAAGTTCCAAAAAGCATACTTTTCTACATATTTGCGTTATGTGTTGGAGACTTAATGACC

ATGATAGCTATGTTACTTCTAGTCATTGAACTAGTTTTTGGCACGTGGCAGTTTTCATCAATGGTTTGCACTAGCTATCT

AATTTTTGATTCTATGAATAAATTTATGGCTCCAATGATAGTTTTTTTGATTAGTAGAACTTGCTATTCAACTGTCTGTC

TTGATAAAACAAGAGGAGAAAAAGCAGCAACACTGAAGTATGCAATAATTCAATTCTGCATTGCCTTTGCATTTGTCATG

ATCCTCCTTTGGCCAGTGTTTGCCTATTCCCAAGTTTTTACATTCTACATGAATCCAAACTCAACTGCCCAGGAAGTAAC

AGTTATGAGGAAATGTGGTTTCTTCCCGCCACCACAAATTGAGTTTTGGTTCAACTTGATCGCTTGCATCACTTCCTATG

CAGTCCCTCTTTTCGGAATTATTTACTGGTATGTCTCGGTACCCTTTTTCCTGAAACGGCGGGCGCTGACGACACTGGTC

GCCTCAAGTTCAATGGATGCAGCTCTTCGAAAAGTGATCACAACTGTTCTCCTTCTCACAGTCATTTATGTTCTCTGTTG

GACACCTTACTGGGTCAGTATGTTTGCTAATCGCATCTGGATTATGGAGAAAAAGAGCATCATCATCATTTCCTATTTCA

TCCATCTTCTTCCATACATCAGTTGTGTCGCTTATCCATTGATCTTTACATTGCTCAATCGAGGGATTCGATCGGCTCAC

GCGAAGATTGTAGCTGATCAGAGAAGACGCTTCAGAAGTCTTACTGATGAAGCAAGCTCGCAAATTCGGACGGCAATAAG

AACCATCCCTGGCACGAAAATGAAGAAAAACGAATTTTTAACTCGTACAGAAGAGATATCTTCCGACAAGATTGCTTCAG

AGGCCGTATCAGAGTTCCGAGAAGGCCTACCATCTCAAACATCGTTTCCCGATGAGACACTTTTATGA

>pR157

ATGAGCAGTAGCACGGAAATTGCTGATACATCTGAAAATGTGCTCAGCAGCTCTGATGGTTATTCATATGAGGATGAGTG

GTGGGAAATGAATTGTTCAATTATACCGCCCGAGTTTAACAGACTAATGTTGGTGGGTGTAGTTGGTGCAGTGATTGCCT

TCATCTCGGTGTGCTTCAACACATTTCTCTTCGTCGTTCTCTGCAGGAATCCACGTCATAGACATTCTCACTTGGTGTAT

TTGATGTTTCTATCAGTAGCTGATATATTTTTATCAGCAGCCTATATTCTATTGTTTCCTGTCAATTTGTACATGGACTA

TTTTGCATCTGAACTGTTGGCTGCAGCATGGTGGTCTTATATGAGAATTATGATCACAATATCTCATGTATTCATTTCAA

CGTCAGCATTTTTAATTTGTGCAGCAGCATTTGAAAGGTACATAACAATTTCAAAAATTGCGTGCCAGTTTGCTCGTCAT

CACCGGCTTATAATTATTGGGGCATGTATTTTCATTGCAATTATTGCCAAGGGGCCAATGTATTTTGAGTTTGAGGTTGT

ACCAAATGCGAATTGCACCGGAGTGACGTCACTAACTGCAATTCCCTCTGAATTCTCAGAGTCCGAGCCTTACAAAACTG

CATATAAGTTCTGGTTCCGAAATCTGCTCACAGTCGCTCTTCCCTTCATAATGTGTTTCTACCTCAACTTTGCAATAATG

CATCGTCTCCGAATTCAACACCTCGGTGCGAAACTCTTTCGATTCGCCACATCAGAGCATCGTCAGAATATCCGAGCCGC

CACGCTGATGCTTGTTGCAGTTACATGTTCATATCTGGCATCAAACCTTCTCAATGTTGTAGTATACACTTGGGAACTGG

TAGATAAAGAGTCATTACTATCAGAAAACATTCGGCCTCTTTACACTTTATCTTCGGATTTGGTATCATTGCTGACTGTG

GTCGCAAGTGCTTGTCGACTTCCGATTTATTTGGTTTGCAATGCAAGGATTCGATGTGAAATTTTGGATTATGTCGATAA

TTGTGTGCTCACTCATTTACATATCAAACCGTATTCGGGTCTCGGAAGTAAACGATGTCGTGCAACAACTGTAAGATATT

GTGACACTGGTAGCGGTTATATGGTCTATGATGCTCCCAATAAAAAAGGCGAGCGACGAGTTCGGTCGGTTGGAACTGGA

TTAGATCGAGTCGTATTATCAGTTGCCATGGGAAGTATCAAAGCATCACAGAGTACGACACATTTGACTATTCCACTTAA

TAATAATGCAATGATAATTCAAGAAGAGTGA

>pR158

ATGAGCAGTAGCACGGAAATTGCTGATACATCTGAAAATGTGCTCAGCAGCTCTGATGGTTATTCATATGAGGATGAGTG

GTGGGAAATGAATTGTTCAATTATACCGCCCGAGTTTAACAGACTAATGTTGGTGGGTGTAGTTGGTGCAGTGATTGCCT

TCATCTCGGTGTGCTTCAACACATTTCTCTTCGTCGTTCTCTGCAGGAATCCACGTCATAGACATTCTCACTTGGTGTAT

TTGATGTTTCTATCAGTAGCTGATATATTTTTATCAGCAGCCTATATTCTATTGTTTCCTGTCAATTTGTACATGGACTA

TTTTGCATCTGAACTGTTGGCTGCAGCATGGTGGTCTTATATGAGAATTATGATCACAATATCTCATGTATTCATTTCAA

CGTCAGCATTTTTAATTTGTGCAGCAGCATTTGAAAGGTACATAACAATTTCAAAAATTGCGTGCCAGTTTGCTCGTCAT

CACCGGCTTATAATTATTGGGGCATGTATTTTCATTGCAATTATTGCCAAGGGGCCAATGTATTTTGAGTTTGAGGTTGT

ACCAAATGCGAATTGCACCGGAGTGACGTCACTAACTGCAATTCCCTCTGAATTCTCAGAGTCCGAGCCTTACAAAACTG

CATATAAGTTCTGGTTCCGAAATCTGCTCACAGTCGCTCTTCCCTTCATAATGTGTTTCTACCTCAACTTTGCAATAATG

CATCGTCTCCGAATTCAACACCTCGGTGCGAAACTCTTTCGATTCGCCACATCAGAGCATCGTCAGAATATCCGAGCCGC

CACGCTGATGCTTGTTGCAGTTACATGTTCATATCTGGCATCAAACCTTCTCAATGTTGTAGTATACACTTGGGAACTGG

TAGATAAAGAGTCATTACTATCAGAAAACATTCGGCCTCTTTACACTTTATCTTCGGATTTGGTATCATTGCTGACTGTG

GTCGCAAGTGCTTGTCGACTTCCGATTTATTTGGTTTGCAATGCAAGGATTCGATGTGAAATTTTGGATTATGTCGATAA

TTGTGTGCTCACTCATTTACATATCAAACCGTATTCGGGTCTCGGAAGTAAACGATGTCGTGCAACAACTGTAAGATATT

GTGACACTGGTAGCGGTTATATGGTCTATGATACAAAACAAGAGGAAGTGATACCGAGTCCAGTGGCTCCCAATAAAAAA

GGCGAGCGACGAGTTCGGTCGGTTGGAACTGGATTAGATCGAGTCGTATTATCAGTTGCCATGGGAAGTATCAAAGCATC

ACAGAGTACGACACATTTGACTATTCCACTTAATAATAATGCAATGATAATTCAAGAAGAGTGA

>pR159

ATGAATCCATCGATATCCACGGCGGGAGCAGTCGTCGACGATTTTCAGATGTCATGCAGATTTTCAAATTTTGGAGATCA

TGCACCTGAATCATTTGACTCACTTTCCTGTGGAGCATGCTTCTATTTTATTTTTATGCTCGTAGAGCACTATAAAGATA

GATACGTTGCAAGTCTTTGTGAAGACACTTTCGGAGTACTAGTATGCCCAGTTGATGTCAGTGACAAAATTTGCAATTGC

TATAATCGAGAATATGTCTACAATATTCCGGAGTTCGCACTTAATTCATCTCGAATAAAACCTCTGAATCATCTGCGACT

TGATCCACCGTTACCTTTTCAATTCGATTTAATTGTCAAAGATTGCTGCTCGGCCGCACGATATTGCTGTAGAAATACAC

TTGTGAAATATCATCATCGTGTCGATGATTCTCCATGTCCGCCTACTTGGGATGGATGGAATTGTTTTGATTCAGCAACT

CCTGGTGTTGTTTTTAAACAATGTCCTAATTATATTTATGGTGGATCCAACATTAAAACTGATTATGATAGATTATCTCA

AAAAGTATGCAGATCTAACGGATGGGCGACGCCAGAAGTCAATGCTGCAGCACGAGAGCACACTGATTACACGGGTTGCA

TGTCTAATGGGGATGTTGAAGCTCGTATTCTAGCTGGATTGCTTACTTACTCTGCTAGCGTCATATTTCTCATTCCAGCG

GTTTTCTTACTGACACTACTCAGACCAATTCGTTGTCAACCAATGTTCATTCTGCATCGTCATCTGCTCATCTCATGTCT

CTTATACGGAGCATTCTATCTGATCACCGTGTCTCTTTTCGTTGTCAACGATGCTCCACTCTCTTCACAAGTCTTCCAGA

ATCATCTGTTCTGTCGTCTTCTCTTCTCCATTCAATTACGATACCTTCGACTGACCAATTTCACATGGATGCTTGCAGAA

GCTGTTTACTTATGGAGACTATTACATACTGCTCAACATTCTGAAGGAGAGACTCTCCGAAGCTACAAAGTTATATGTTG

GGGTGTCCCTGGAGTGATCACAGTTGTCTACATATTTGTCCGAAGTCTGAATGACGACGTTGGAATGTGTTGGATTGAAA

ACTCGACGGTCGCCTGGATCGAATGGATGATCATAACACCATCTCTTCTTGCAATGGGTGTCAATTTGCTCTTGCTCGGG

CTAATCGTCTACATTCTCGTCAAAAAACTCCGATGTGATCCGCACTTGGAGAGAATTCAATACAGAAAAGCTGTCCGCGG

AGCACTCATGCTTATTCCAGTATTCGGTGTTCAACAGCTTCTTACAATATATCGATTCAGTAACACATACTATCAGGTTA

CGGATCAATCACTTAATGGATTACAGGGAATGTTTGTCTCATTTATCGTTTGTTACACAAATCGTTCTGTCGTAGAATGT

GTTCTGAAATTCTGGTCATCACATCAGGAGAAGCGGGCTCTCGGAGCTGAATGCCGACAAAGAATGAGCATTCAAGAAAG

CGGAAAACTTATTCTCAGGCCGCCACAAACCGAACACGTCGTACTCTAA

>pR160

ATGTTCCCATTCTTGGCAATACTGGGTGTTTTTGGTAATTCTATCACATTATTTGTCTTGCTTAGTAGATCAATGAGAAA

CAGCACAAATGAAATGCTGGCGGCAGCAGCTTTTAGTGACATCCTCTACATTATTTGTATGGTTCCAAATCAGTTGAGTC

GTTGGCCGTCAATGGTTCTTTCCGACTGTCCCGATAACCCAGAGAGAAAATGTGCATCGGAATATCACATGTGGTTTGCA

GTTCACAAACATCATCTAGCTTTTCTTGCAAACTGGTTTTCTGCGGCTAGCACTTGGTTCATTGTGACGGTATCATTTGA

CCGACTATACGCGATAAAGGCTCCATTTTCTGCTCGACTGCAAACAATTTGCCCAAGACATAACCTTGTTGCGATTCCAC

TAATCCTTTTCTTCACGGGTGCAAGCTGCTTTCATATGAATTTCAAACTGCTTGATGACCATGCAGGCTCTGGAAATTTT

ACGAATTCAACGATGGAAAATATTGAAGATAACAGAATCACTTTGATCCAAGTAATGACAGTTTTCATGTTCATCTTCCA

CATTCTGATCCCAATGATTCTTTTAATATCATTTCATACTTGCTTGCTGTATTATCTTCGTAACCGATTTAGGCACTTCT

TTCCGGCTCGAACAAGAAGTGCAAGAAACTCGACAAGATCCGATGACGTCCCTGCTCCACTTTTATCCAACGTAACCGAT

CGATCGGAAGTAGTGCGACATCATTCTTCTAACAGTGGTGTTTGGAATCGCCATGTGAACAAAGCCGAACGTCATGTAAC

TTATACGGTCATAGCAATTGTCACATGCTACATTGTTTCACATATTCCATCTGCATGCCTATATGTTTACATCAACCTGT

TTCATAACGTTCTTTACTCAACAAGATGGATGTACACAAGTGTTCAAGTTTCCACAACCGTTGTCACATGCTCCAAAGTT

GCAAATTTCATACTTTTTTGCATGAGCAGTAAACATTTTCGCAAAGAAATGAAAAAGAAGTTGTGCTTTCTGTCGTGCAA

ATACAAAGATGGAATCAGTGGAAAAGATAGTGCAAATCAGCCGAGAACAAGGTTCAGATTCAGATCCCTCCCTCTGAACA

TCATTGGCGATTCAAGTAACAACTGTTCTCACGATTGA

>pR161

ATGAGCTCGGCAGAGGAAGAACAGTTTTTACGGGAATGGGCATTCTCATCCCGTCTTTTGTTCTACTCAATCATTGCTGT

CACTTCAATTACTCTGCCAGTACTTGTGATAATTATGGTTTTGATAAGACCACATCGGAATTCACAATACTTTCCATACA

TGTGTGGGATCATCGCCGGCAATTTTGTGCTACTATCTACTGTCTTCACATTGGTTGTCTCGGAAAATGTTAATTTTGTT

TATAGTTATCTTCCTGGCTTTATGATCTGCAAATTTTCTGCATTCCTTGTCAACTCGGCATCTTGTTTCATACATTGGAC

ATGGGTTGGAATGTATTCACAACGTTTTCTTCATGTTTTCTTTCCACTTCGTTCTAGACGAAGACTCCCACGTAGTACCT

GGAATACACTACTTATAATTTTGTTAATCTCTTGTACGTGCCAAATCTGGGCTCCAATTACAATAACCGAGCTTAGCTTC

TCTCCGGGAAACAATAGCCAAGGCACGTATTGTGCAGAGGATCCGTACTTCAGCAGCTCATCAAAGATCATTGTTCTCGT

TGAAAGCACAATTACATTCTTCATTCCATTCATGTTAACTGTTCTAGCGGATATCTCAGTTTTGGTGTCAAAAGTTCCAT

GGCAAACACCTTTCAAATTAATTTCTGCTGATGATTTAAGAAACAATAATAGAAGTGATAACATGAAAATTGTTTCAAAA

TTAAATTTAGAGAGCGCTGAAAAACGAAGAATAAATGCAATCAAAAGATGTCTCATTTCTGCAACGCTCACTTTAGTACT

AAATTTGCCAAATTATGTCCTTCAGGTCGTTGATGAGTTCTACTCATTGCGAGAGCATGGAAATATCTCCATTCGACGAG

TTTTTCTTCAGGCGGACGCAGCTGTTTATATTTTGTATCTTCTGCAGTTCCCATTGGTTCCAGTGAACATGTATTTCCTT

CGGCAGAACATTACAAGAAGCTCAAGGAAACTTCGAGAGAAGACAACATTACAACAAGAGCACAGCAAATCGTTCCAAGT

GCTCGAAATGAGTAATGATCAGAGCCGAATTCCGTTGAGCACCACGATTCCATCTCCAATGTCTCATGCGTAG
